# PG-MLD: Physics-Guided Molecular Representation Learning via Dynamic 3D Trajectory Distillation

**DOI:** 10.64898/2026.07.29.741404

**Authors:** Zihe Liu, Zhonghan Wu, Zhuo Chen, Xin Gao, Bin Yu

## Abstract

Molecular representation learning underpins molecular property prediction and drug design by capturing molecular structure–property relationships. SMILES-based molecular language models learn chemical semantics from large-scale unlabeled data and support efficient inference. However, the one-dimensional nature of SMILES constrains their ability to capture 3D geometry and conformational evolution, whereas 3D molecular models require conformer generation and substantial computational resources. To bridge this gap, we propose PG-MLD, a dynamic 3D-to-1D physical knowledge distillation paradigm for molecular representation learning. PG-MLD constructs a dynamic 3D physical teacher by combining equivariant geometric encoding with Liquid Time-Constant modeling to capture 3D geometry, atom-level electronic descriptors, and conformational evolution. PG-MLD subsequently distills the learned trajectory knowledge into SMILES-based students through atom- and molecule-level representation alignment and cross-modal contrastive learning, with masked language modeling retained where supported. The distilled students perform downstream tasks using only SMILES, without conformer generation or molecular dynamics simulations. Experiments on MoleculeNet show that PG-MLD improves overall property prediction performance across three molecular language student architectures while maintaining SMILES-only inference. The learned representations also encode 3D geometry and conformational dynamics more effectively, demonstrating that dynamic 3D physical knowledge can be transferred to SMILES-based molecular language models with different architectures.

## Introduction

Molecular representation learning plays a central role in computational chemistry, drug discovery, and materials science (Masood, Kaski, and Cui 2025; Gaudelet et al. 2021; Reiser et al. 2022). Its goal is to learn representations that accurately capture molecular structure–property relationships and generalize well (Sheshanarayana and You 2025). In recent years, the field has evolved from conventional molecular descriptors (Rogers and Hahn 2010) and two-dimensional graph neural networks (Reiser et al. 2022) toward SMILES-based molecular language pretraining (Chithrananda, Grand, and Ramsundar 2020; Ahmad et al. 2022; Ross et al. 2022; Zhang et al. 2021), 3D geometric modeling (Stärk et al. 2022), and multimodal representation learning (Zhou et al. 2025; Jiang et al. 2025). Molecular language models extract chemical semantics from large collections of SMILES strings, offering scalable pretraining and efficient inference. By contrast, 3D molecular models use atomic coordinates and geometric constraints to better encode spatial structure. Although recent studies have introduced 3D information into low-dimensional molecular models through cross-modal alignment or knowledge distillation (Cho et al. 2025), most existing approaches still rely primarily on static conformations (Zhu et al. 2024; Wang et al. 2024a). This reliance leaves dynamic 3D physical knowledge largely underexplored. Consequently, SMILES-based molecular language models remain limited in their ability to explicitly capture 3D geometry, conformational evolution, and local physicochemical environments (Li et al. 2023a).

Molecules are inherently dynamic, with their structures evolving continuously over time. Atomic positions, bond angles, dihedral angles, and local charge environments are shaped by thermal fluctuations, intramolecular interactions, and external conditions (Zhu et al. 2024; Thölke and De Fabritiis 2022). Molecular dynamics trajectories characterize time-resolved conformational evolution, thereby providing richer physical supervision than static conformers (Zhang and Vitalis 2025; Jing et al. 2024). However, most existing 3D representation methods operate on a single conformer or a small conformer ensemble (Hamakawa and Miyao 2025). They therefore struggle to capture temporal dependencies, continuous conformational evolution, or the coupling between 3D geometry and local physicochemical environments along a trajectory. Moreover, molecular dynamics simulations and 3D structure processing are computationally expensive. Three-dimensional inputs are also difficult to obtain at scale for molecular screening and practical deployment. A key challenge is therefore to transfer dynamic 3D physical knowledge to SMILES-based molecular language models during training, enabling physically informed representations and efficient SMILES-only inference.

To address this challenge, we propose PG-MLD, a distillation paradigm that transfers dynamic 3D physical knowledge to one-dimensional molecular language models. PG-MLD consists of two stages. First, the 3D teacher combines equivariant geometric encoding with continuous-time modeling to capture 3D geometry, conformational evolution, and atom-level electronic descriptors along each trajectory. Second, PG-MLD transfers the learned dynamic 3D physical knowledge to SMILES-based molecular language models with different architectures through atom- and molecule-level representation alignment and cross-modal contrastive learning, together with masked language modeling where supported. As a result, the student models acquire physically informed representations from SMILES alone, avoiding conformer generation and costly 3D computation during downstream inference.

Our contributions are threefold:

- We propose PG-MLD, a dynamic 3D physical knowledge distillation paradigm for molecular language models that transfers 3D geometry and conformational dynamics from molecular dynamics trajectories to SMILES representations.
- We develop a 3D teacher that combines equivariant geometric encoding with Liquid Time-Constant modeling. We further introduce atom- and molecule-level representation alignment to transfer dynamic 3D physical knowledge across modalities.
- Extensive experiments across multiple molecular language models demonstrate that PG-MLD improves overall property prediction performance while enabling physically informed, SMILES-only inference.

## Related Work

### Molecular Representation Learning

Molecular representation learning aims to acquire transferable representations from large-scale unlabeled molecular data. SMILES-based approaches formulate molecules as chemical sequences and adopt self-supervised pretraining strategies from natural language processing. MoLFormer (Ross et al. 2022) learns contextual molecular representations through masked language modeling on large-scale SMILES corpora. More recently, SMI-Editor (Zheng et al. 2025) introduces edit-based pretraining with fragment-level supervision to capture chemically meaningful substructures, while MLM-FG (Peng et al. 2025) employs functional-group-aware masking to incorporate structural priors into SMILES representations. Despite these advances, sequence-only models remain limited in explicitly representing 3D geometry and conformational dynamics.

Beyond sequence modeling, existing studies have explored geometric priors and cross-modal knowledge transfer to enhance molecular representations. SCAGE (Qiao et al. 2025) combines pretraining objectives based on molecular fingerprints, functional groups, 2D interatomic distances, and 3D bond angles, together with multi-scale conformational learning, to improve molecular property prediction and substructure interpretability. D&D (Cho et al. 2025) trains a denoising teacher on 3D conformers and transfers the learned representations to a 2D graph encoder through cross-modal distillation, thereby removing the need for 3D conformer input in downstream tasks. However, these methods derive geometric supervision primarily from a single conformer or a limited set of static conformers, leaving trajectory-level molecular dynamics insufficiently explored (Zhu et al. 2024; Hamakawa and Miyao 2025).

### Dynamic 3D Molecular Representation Learning

Three-dimensional molecular representation learning encodes 3D geometry from atomic coordinates and geometric constraints. Uni-Mol (Zhou et al. 2023), HAGO-Net (Pei et al. 2024), ViSNet (Wang et al. 2024b), and MPerformer (Wang et al. 2023) capture interatomic interactions through geometric features and equivariant mechanisms. These methods have advanced quantum-chemical property prediction, molecular energy estimation, and atomic force prediction.

Molecular dynamics (MD) trajectories consist of sequences of continuously sampled 3D conformations. They record atomic motions, conformational transitions, and state dependencies across time. Compared with static conformers, MD trajectories provide richer physical information for modeling flexible molecules and conformation-sensitive properties (Zhang and Vitalis 2025; Jing et al. 2024). Existing studies encode multi-frame trajectories using time-lagged representation learning, graph neural networks, spatiotemporal attention, or generative modeling (Mardt et al. 2018; Huang et al. 2024; Wu et al. 2023; Jing et al. 2024; Zou, Wang, and Tiwary 2025). These approaches extract molecular dynamical states and temporal features. Despite modeling spatiotemporal information, these methods remain limited in capturing long-range dependencies and conformational evolution across multiple time scales (Mardt et al. 2018; Zou, Wang, and Tiwary 2025). In particular, atoms and local structures exhibit distinct motion rates and response time scales, making their cross-frame state evolution difficult to represent adequately using a uniform discrete update mechanism.

Liquid Time-Constant Networks (LTCs) (Hasani et al. 2021) use learnable time constants to adaptively regulate state-update rates, making them suitable for nonstationary sequences and multi-scale dynamical processes. This property may enable the joint modeling of long-term molecular scaffold stability and rapid local changes in flexible regions. However, LTCs have not yet been systematically applied to dynamic 3D molecular trajectory representation or to the transfer of physical knowledge from such trajectories to molecular language models.

## Methods

### Overall Architecture

As shown in Fig. 1, PG-MLD adopts a teacher–student architecture comprising a 3D teacher and a SMILES-based molecular language student. Given an input SMILES string, PG-MLD constructs a dynamic 3D physical view from the initial 3D conformer, molecular dynamics trajectories, conformational perturbations, and atom-level electronic descriptors. This view captures 3D geometry, conformational dynamics, and local electronic environments. The 3D teacher first encodes the spatial structure of each frame using an equivariant geometric encoder. A Liquid Time-Constant network then models conformational evolution across frames. After training, the teacher is frozen. Its dynamic 3D physical knowledge is then distilled into the student through atom- and molecule-level representation alignment and cross-modal contrastive learning, together with masked language modeling where supported. The resulting student performs downstream property prediction from SMILES alone, without conformer generation or molecular dynamics simulations.

**Figure 1.**
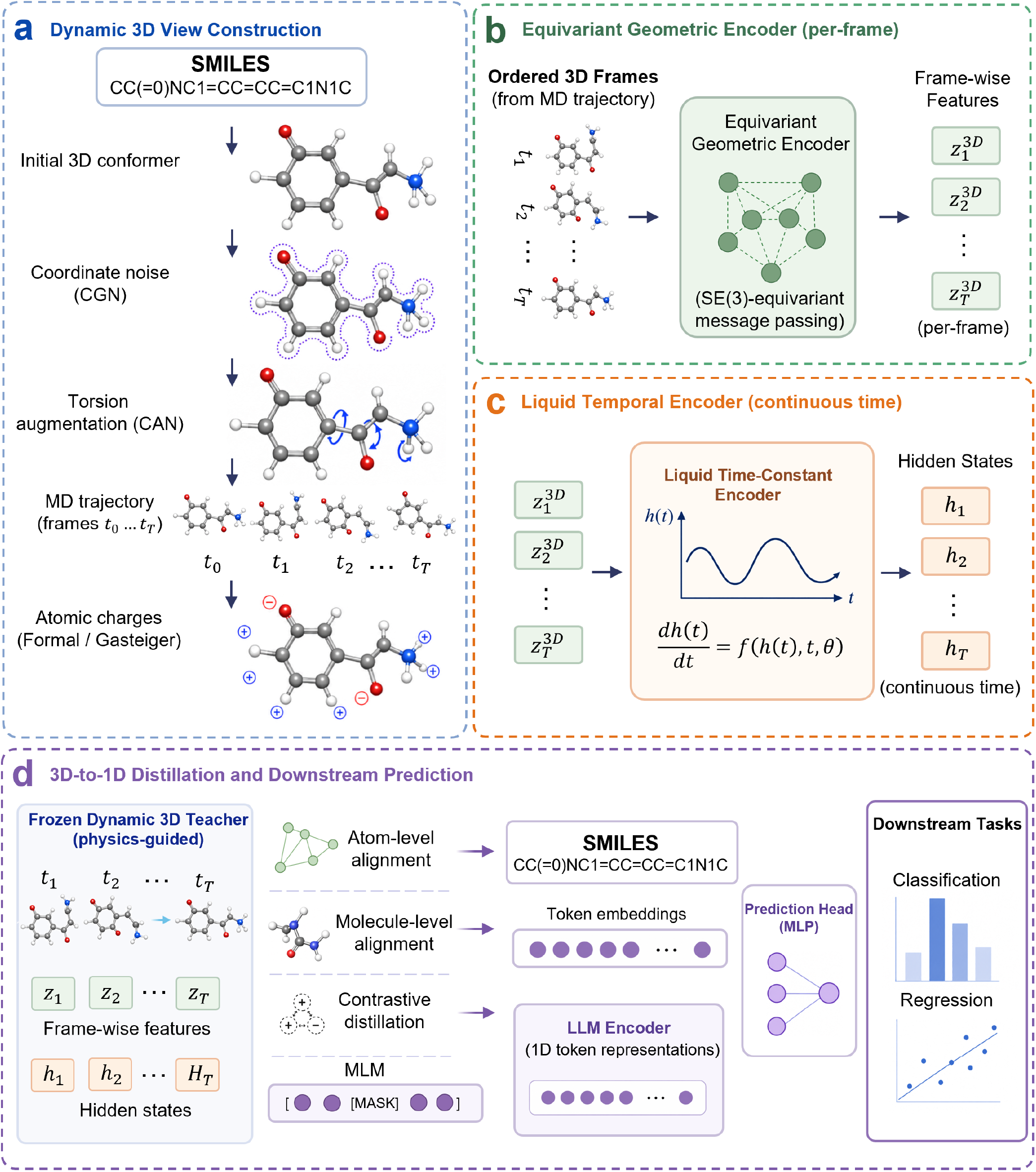
Overview of PG-MLD. (a) Construction of a dynamic 3D physical view from molecular dynamics trajectories, conformational perturbations, and atom-level electronic descriptors. (b) Frame-wise equivariant geometric encoding. (c) Cross-frame continuous-time modeling with a Liquid Time-Constant network. (d) Distillation of dynamic 3D physical knowledge into a SMILES-based molecular language model through atom- and molecule-level representation alignment, followed by downstream property prediction.

### 3D Teacher Encoder

As shown in Fig. 2, the 3D teacher in PG-MLD comprises four components: dynamic 3D physical view construction, frame-wise equivariant geometric encoding, Liquid Time-Constant modeling, and dynamic representation generation. Together, these components model 3D geometry, conformational evolution, and local electronic environments.

**Figure 2.**
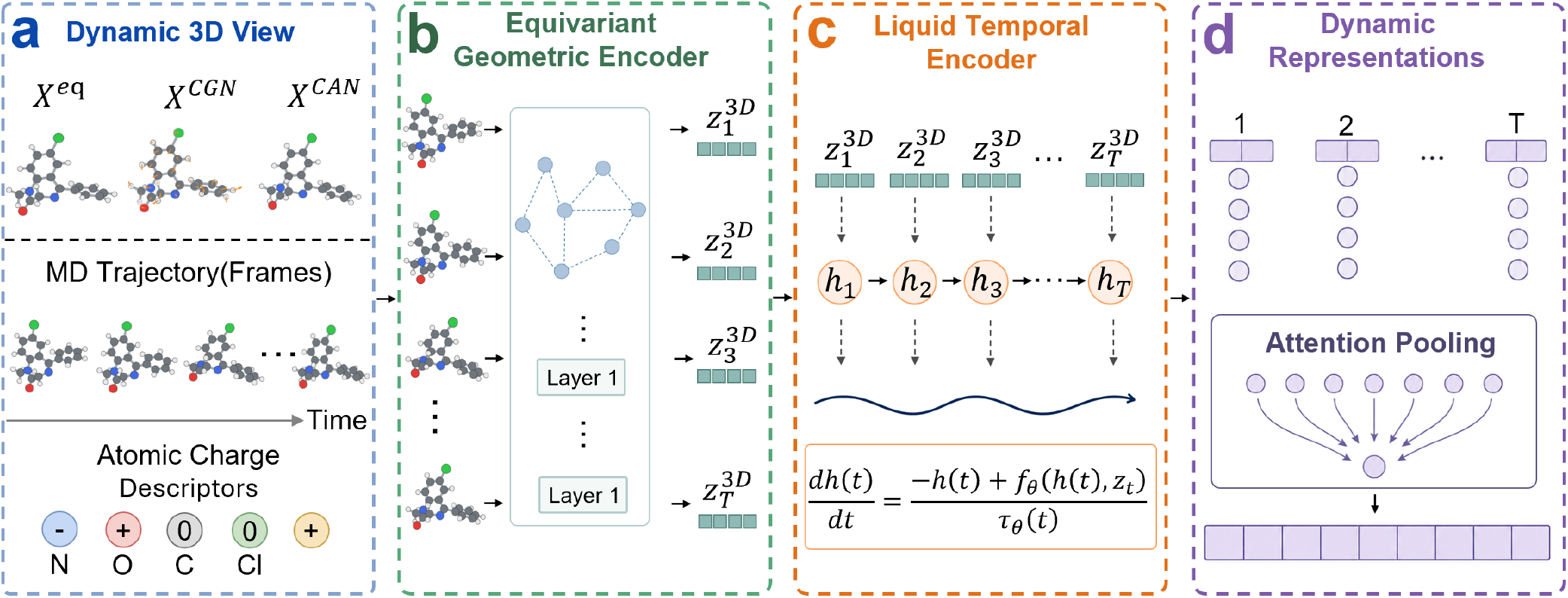
Architecture of the 3D teacher encoder. (a) Dynamic 3D physical view construction. (b) Frame-wise equivariant geometric encoding. (c) Cross-frame continuous-time modeling. (d) Generation of dynamic atom-level representations and attention pooling into a molecule-level representation.

During dynamic 3D physical view construction (Fig. 2a), PG-MLD integrates the initial conformer with coordinate Gaussian noise (CGN), conformational angle noise (CAN), and temporally ordered molecular dynamics trajectories. Formal charges and Gasteiger partial charges are included as atom-level electronic descriptors. CGN captures local coordinate perturbations, whereas CAN represents conformational variations. MD trajectories encode the temporal evolution of molecular structure, while atomic charges provide complementary information about local electronic environments.

The equivariant geometric encoder processes each trajectory frame to obtain the atom-level spatial representation 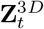for frame *t* (Fig. 2b). The Liquid Time-Constant encoder then performs continuous-time modeling over the temporally ordered frame-wise representations (Fig. 2c). We express the hidden-state evolution in the generic form

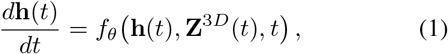

where **h**(*t*) denotes the continuous-time hidden state and *f*_*θ*_(*·*) denotes an LTC state-update function parameterized by learnable time constants. By adaptively regulating the state-update rate, this mechanism captures cross-frame dependencies and conformational evolution across multiple time scales. Following continuous-time encoding and aggregation, the teacher produces two outputs: the atom-level representation 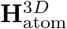 and the molecule-level representation.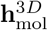 These outputs serve as teacher signals for subsequent atom- and molecule-level representation alignment.

### Molecular Language Student Encoder

PG-MLD employs a pretrained molecular language model as the student encoder. Given the SMILES representation *S* of a molecule *M*, the student model tokenizes *S* into a sequence of length *L* and encodes it as a token-level language representation 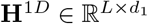

To enable atom- and molecule-level representation alignment with the 3D teacher, PG-MLD uses a token–atom mapping *a*(*ℓ*) to aggregate the token features associated with each atom. The resulting atom-level representations are then pooled into a molecule-level representation:

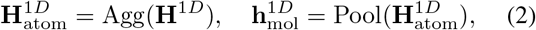

where 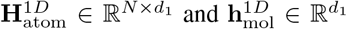denote the atom-and molecule-level language representations, respectively, and *N* is the number of atoms. Agg(·) aggregates token features mapped to the same atom, whereas Pool(·) reduces the atom-level representations to a molecule-level representation. The two representations are used for subsequent atom- and molecule-level representation alignment during 3D-to-1D knowledge distillation.

### 3D-to-1D Physical Knowledge Distillation

During distillation, the 3D teacher is frozen. Atom- and molecule-level representation alignment is used to transfer dynamic 3D physical knowledge, including 3D geometry, conformational dynamics, and local electronic environments, to the molecular language student. Because the teacher and student representations differ in dimensionality and feature distribution, PG-MLD applies separate projection heads at each level to map them into the corresponding shared latent spaces:

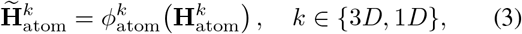

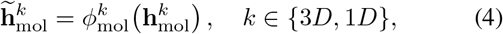

where *k* ∈ {3*D*, 1*D*} indexes the teacher and student, respectively; 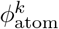 (*·*) and 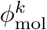 (*·*) denote the corresponding atom- and molecule-level projection functions. A tilde denotes a representation projected into the corresponding shared latent space.

Using the token–atom mapping, atom-level representation alignment matches corresponding atom representations produced by the 3D teacher and the 1D student. This alignment transfers information about local geometry, atomic neighborhoods, and electronic environments. Molecule-level representation alignment matches their global representations to transfer overall conformational dynamics and global physical information. The corresponding atom- and molecule-level representation alignment losses are defined as

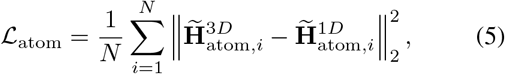

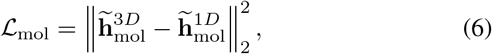

where *N* denotes the number of atoms in the molecule.

PG-MLD further employs cross-modal contrastive learning to bring the 3D and 1D representations of the same molecule closer while improving discrimination between different molecules. For student architectures that support masked language modeling, the corresponding objective is retained to preserve the modeling of SMILES fragments, sequence syntax, and contextual semantics. The overall distillation objective is given by

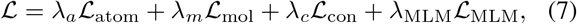

where *L*_con_ and *L*_MLM_ correspond to cross-modal contrastive learning and masked language modeling, respectively. The coefficients *λ*_*a*_, *λ*_*m*_, *λ*_*c*_, and *λ*_MLM_ control the contribution of each loss. For a student without an MLM objective, *λ*_MLM_ is set to zero. After distillation, the teacher is discarded. The student encoder is retained and paired with a task-specific prediction head. They are fine-tuned using a classification or regression objective, depending on the downstream task. Inference requires only SMILES input, without 3D conformer generation or molecular dynamics simulations.

## Experiments

This section systematically evaluates the effectiveness of PG-MLD. We first describe the datasets, baseline methods, and experimental settings, and then address the following five research questions:

- **Q1:** Does PG-MLD achieve competitive performance across MoleculeNet classification and regression tasks?
- **Q2:** Does PG-MLD consistently improve different student models through dynamic 3D physical knowledge transfer while preserving SMILES-only inference?
- **Q3:** Does PG-MLD improve agreement between similarity in the representation space and structural similarity between molecules?
- **Q4:** Does PG-MLD improve the identification of property-relevant atoms and key substructures?
- **Q5:** How does PG-MLD affect the global distribution and class separability of the molecular representations learned by the student model?

## Datasets and Baselines

### Datasets

We evaluate PG-MLD on nine molecular property prediction datasets from MoleculeNet (Wu et al. 2018). BBBP, BACE, ClinTox, Tox21, ToxCast, and HIV are classification tasks evaluated using ROC-AUC. ESOL, Free-Solv, and Lipophilicity are regression tasks evaluated using RMSE. All methods follow the same random splits and evaluation protocol. We report the mean and standard deviation over five runs using seeds 42–46.

### Baselines

We compare PG-MLD with representative methods from three molecular representation learning paradigms. Molecular graph pretraining baselines include GraphMVP (Liu et al. 2022) and MolCLR (Wang et al. 2022). Geometry-based or structure-enhanced baselines include GEM (Fang et al. 2022), KPGT (Li et al. 2023b), Galformer (Bai et al.2026), KANO (Fang et al. 2023), MoleculeFormer (Qin et al. 2025), and MolGramTreeNet (Zhang et al. 2026). ChemBERTa (Chithrananda, Grand, and Ramsundar 2020), ChemBERTa-2 (Ahmad et al. 2022), and MoLFormer (Ross et al. 2022) represent SMILES-based molecular language pretraining methods. Detailed baseline descriptions and implementation settings are provided in Appendices B and D of the supplementary material, respectively.

## Results and Analysis

### Property Prediction Performance on MoleculeNet (Q1)

Table 1 compares PG-MLD with representative methods on the MoleculeNet benchmark. PG-MLD achieves the best performance on all six classification datasets, with ROC-AUC scores of 0.966, 0.936, 0.997, 0.851, 0.744, and 0.825on BBBP, BACE, ClinTox, Tox21, ToxCast, and HIV, respectively. For the regression tasks, PG-MLD achieves the lowest RMSE on ESOL and FreeSolv, with scores of 0.627 and 0.945, respectively. These results indicate that dynamic 3D physical knowledge distillation improves the ability of molecular language models to represent complex structure– property relationships while preserving SMILES-only inference.

**Table 1.** Performance comparison on molecular property prediction tasks from MoleculeNet. Results are reported as mean (standard deviation) where available, using ROC-AUC for classification tasks (higher is better) and RMSE for regression tasks (lower is better). The best and second-best results are shown in **bold** and <u>underlined</u>, respectively.

| Method | BBBP↑ | BACE↑ | ClinTox↑ | Tox21↑ | ToxCast↑ | HIV↑ | ESOL↓ | FreeSolv↓ | Lipo↓ |
| --- | --- | --- | --- | --- | --- | --- | --- | --- | --- |
| GraphMVP | 0.918(.019) | 0.866(.028) | 0.705(.092) | 0.832(.017) | 0.719(.017) | 0.791(.004) | 0.995(.113) | 1.806(.164) | 0.736(.051) |
| KPGT | 0.927(.028) | 0.893(.032) | 0.915(.027) | 0.847(.013) | 0.715(.012) | 0.768(.007) | 0.802(.096) | 1.230(.231) | 0.621(.018) |
| Galformer | 0.933(.027) | 0.897(.026) | 0.910(.030) | 0.845(.015) | 0.720(.015) | 0.796(.017) | 0.741(.087) | 1.213(.200) | 0.628(.015) |
| MolGramTreeNet | 0.952(.021) | 0.873(.025) | <u>0.991(.012)</u> | 0.832(.021) | 0.734(.031) | <u>0.823(.015)</u> | 0.667(.013) | 1.29(.257) | 0.915(.016) |
| MoleculeFormer | 0.925(.015) | 0.916(.012) | <u>0.901(.010)</u> | 0.819(.019) | 0.729(.016) | <u>0.821(.019)</u> | 0.645(.021) | 1.132(.219) | 0.679(.018) |
| MolCLR | 0.733(.010) | 0.828(.007) | 0.898(.053) | 0.741(.021) | 0.659(.021) | 0.821(.021) | <u>1.113(.023)</u> | <u>2.301(.247)</u> | 0.789(.009) |
| GEM | 0.953(.007) | 0.925(.010) | 0.977(.003) | 0.849(.003) | 0.742(.004) | 0.819(.019) | – | – | – |
| KANO | <u>0.960(.016)</u> | <u>0.931(.021)</u> | 0.944(.013) | 0.837(.013) | 0.732(.016) | 0.816(.017) | 0.670(.019) | 1.142(.258) | **0.566(.007)** |
| PG-MLD | **0.966(.004)** | **0.936(.006)** | **0.997(.002)** | **0.851(.003)** | **0.744(.013)** | **0.825(.011)** | **0.627(.011)** | **0.945(.148)** | <u>0.620(.017)</u> |

### Cross-Architecture Physical Knowledge Transfer (Q2)

To evaluate the transferability of PG-MLD across architectures, we use three representative molecular language models—ChemBERTa, ChemBERTa-2, and MoLFormer— as students. Table 2 compares their performance before and after PG-MLD distillation.

**Table 2.** Performance of molecular language models before and after PG-MLD distillation. Higher values are better for classification tasks, and lower values are better for regression tasks. The best and second-best results are shown in **bold** and <u>underlined</u>, respectively.

| Method | BBBP↑ | BACE↑ | ClinTox↑ | Tox21↑ | ToxCast↑ | HIV↑ | ESOL↓ | FreeSolv↓ | Lipo↓ |
| --- | --- | --- | --- | --- | --- | --- | --- | --- | --- |
| Dataset size | 2039 | 1513 | 1478 | 7831 | 8575 | 41,127 | 1427 | 642 | 4200 |
| ChemBERTa | 0.952(.015) | 0.901(.011) | 0.983(.010) | 0.831(.015) | 0.712(.022) | 0.810(.026) | 0.804(.031) | 1.637(.205) | 0.743(.030) |
| + PG-MLD | 0.953(.007) | 0.927(.008) | <u>0.995(.012)</u> | 0.825(.010) | <u>0.742(.015)</u> | 0.821(.021) | 0.706(.022) | 1.403(.152) | 0.734(.019) |
| ChemBERTa-2 | 0.947(.015) | 0.915(.014) | 0.989(.005) | 0.841(.017) | 0.723(.024) | 0.812(.021) | 0.706(.029) | 1.692(.225) | 0.685(.022) |
| + PG-MLD | **0.966(.004)** | **0.936(.006)** | 0.993(.003) | **0.851(.003)** | 0.733(.015) | <u>0.824(.010)</u> | **0.627(.011)** | 1.114(.153) | 0.686(.019) |
| MoLFormer | 0.957(.012) | 0.913(.011) | 0.986(.006) | 0.839(.016) | 0.724(.025) | 0.794(.019) | 0.677(.026) | 1.011(.214) | 0.662(.025) |
| + PG-MLD | <u>0.964(.005)</u> | <u>0.931(.008)</u> | **0.997(.002)** | <u>0.850(.005)</u> | **0.744(.013)** | **0.825(.011)** | <u>0.653(.019)</u> | **0.945(.148)** | **0.620(.017)** |

As shown in Table 2, PG-MLD improves 25 of the 27 model–task settings across the three students. Representative gains include an increase in ChemBERTa’s BACE ROCAUC from 0.901 to 0.927 and a reduction in MoLFormer’s Lipophilicity RMSE from 0.662 to 0.620. The only exceptions are a slight decrease for ChemBERTa on Tox21 and a marginal increase in ChemBERTa-2’s Lipophilicity RMSE. Overall, these results demonstrate that PG-MLD transfers dynamic 3D physical knowledge across different molecular language model architectures.

## Representation and Interpretability Analysis

### Chemical Neighborhood Consistency Analysis (Q3)

Fig. 3 compares the top-three nearest neighbors retrieved for the same query molecule by the original molecular language model and its PG-MLD-distilled counterpart. For the original model, the retrieved neighbors have a mean cosine similarity of 0.930 and a mean ECFP-based Tanimoto similarity of 0.393 (Rogers and Hahn 2010). After distillation, these values increase to 0.995 and 0.631, respectively. The ECFP-based Tanimoto similarities of the first-, second-, and third-ranked neighbors increase from 0.388, 0.371, and 0.420 to 0.762, 0.575, and 0.556, respectively. Thus, PG-MLD improves the structural relevance of all three retrieved neighbors while increasing their similarity in the learned representation space. This case study suggests that dynamic 3D physical knowledge transfer improves the correspondence between latent-space neighborhoods and those defined by molecular structure.

**Figure 3.**
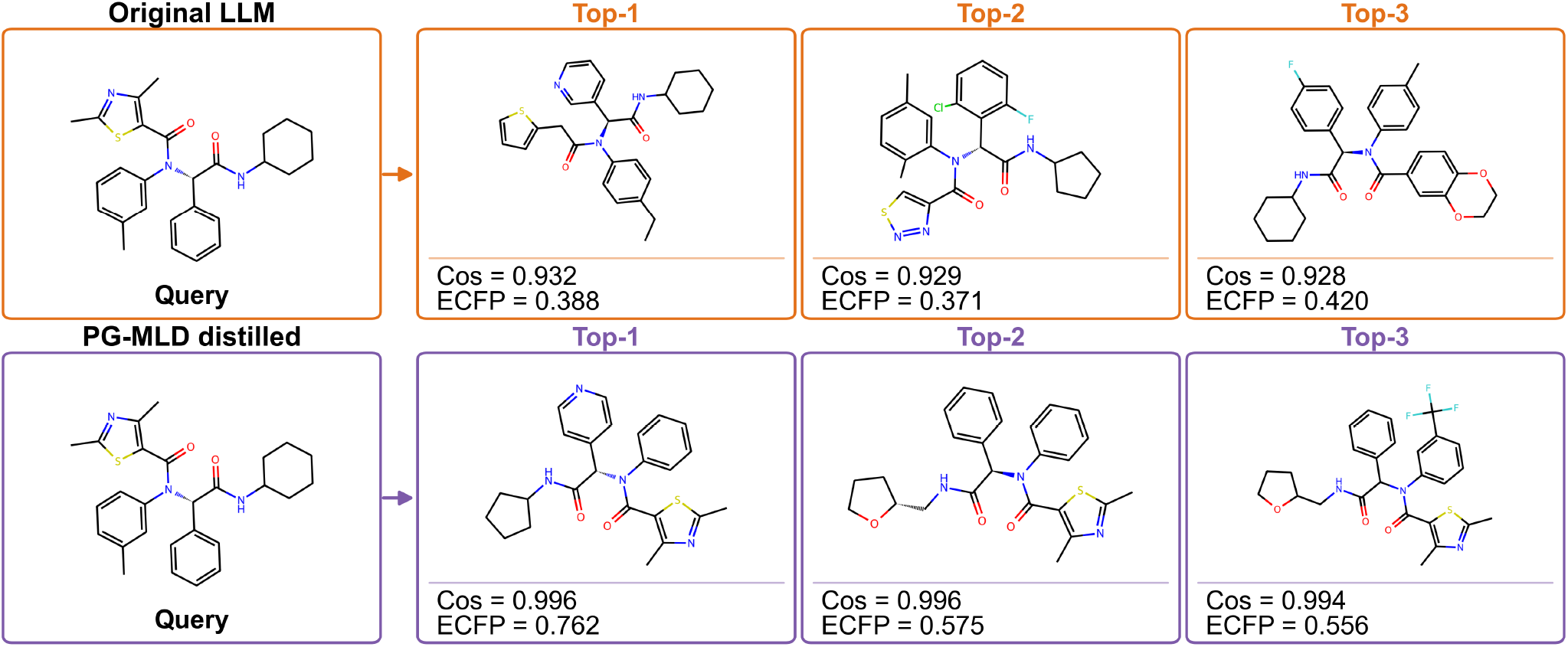
Top-3 neighbor retrieval before and after PG-MLD distillation. Cos denotes cosine similarity in the molecular representation space, whereas ECFP denotes Tanimoto similarity between ECFP fingerprints.

### Token-Level SHAP Analysis (Q4)

Fig. 4 compares the normalized SHAP attributions (Lundberg and Lee 2017) for the same molecule before and after PG-MLD distillation. Blue and orange bars denote the negative and positive contributions of individual SMILES tokens to the prediction of class 1, respectively. Both models correctly classify the sample, whose ground-truth label is 0. PG-MLD reduces the predicted probability of class 1 from 0.092 to 0.032, indicating greater confidence in the correct class.

**Figure 4.**
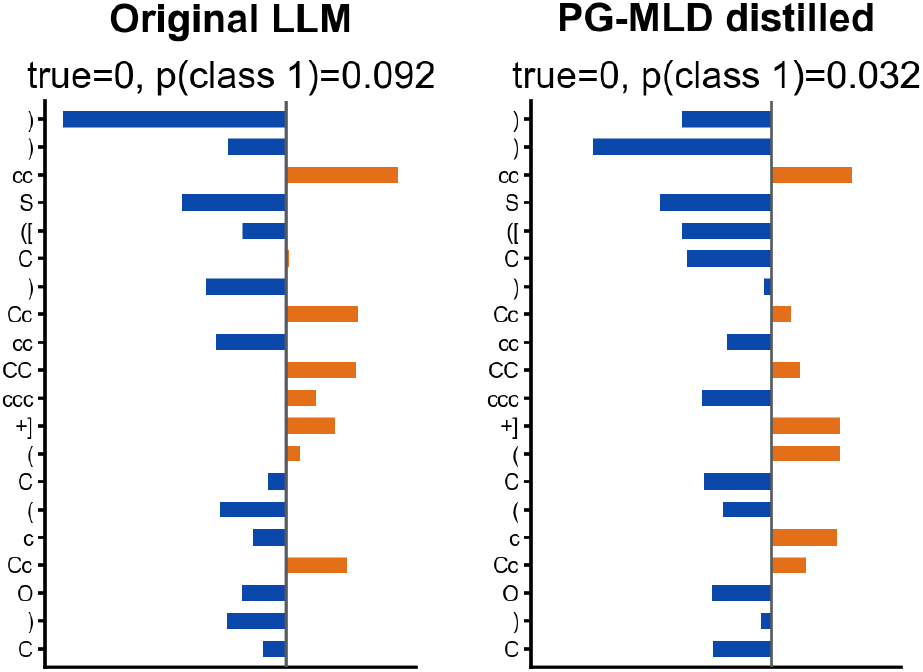
Normalized SHAP attributions before and after PG-MLD distillation. Blue and orange indicate negative and positive contributions to the prediction of class 1, respectively.

The original model is dominated by a small number of tokens with large absolute SHAP values, with positive and negative contributions partially offsetting each other. After distillation, the negative contributions are distributed across more tokens, reducing the dominance of any single token. This example demonstrates that PG-MLD changes how the student model aggregates local token-level evidence during property prediction.

### PCA of Global Representation Space (Q5)

Fig. 5 compares the PCA projections of molecular representations on BACE and BBBP before and after PG-MLD distillation. For BACE, PG-MLD distillation reduces the DBI from 0.95 to 0.85 and the PCA-space class-overlap score from 0.66 to 0.60. The corresponding scores on BBBP decrease from 1.27 and 0.73 to 0.81 and 0.58, respectively. After distillation, the two classes exhibit less overlap and clearer separation. The decreases in both metrics indicate improved within-class compactness and between-class separability. These results suggest that dynamic 3D physical knowledge transfer improves the global structure of the student representation space.

**Figure 5.**
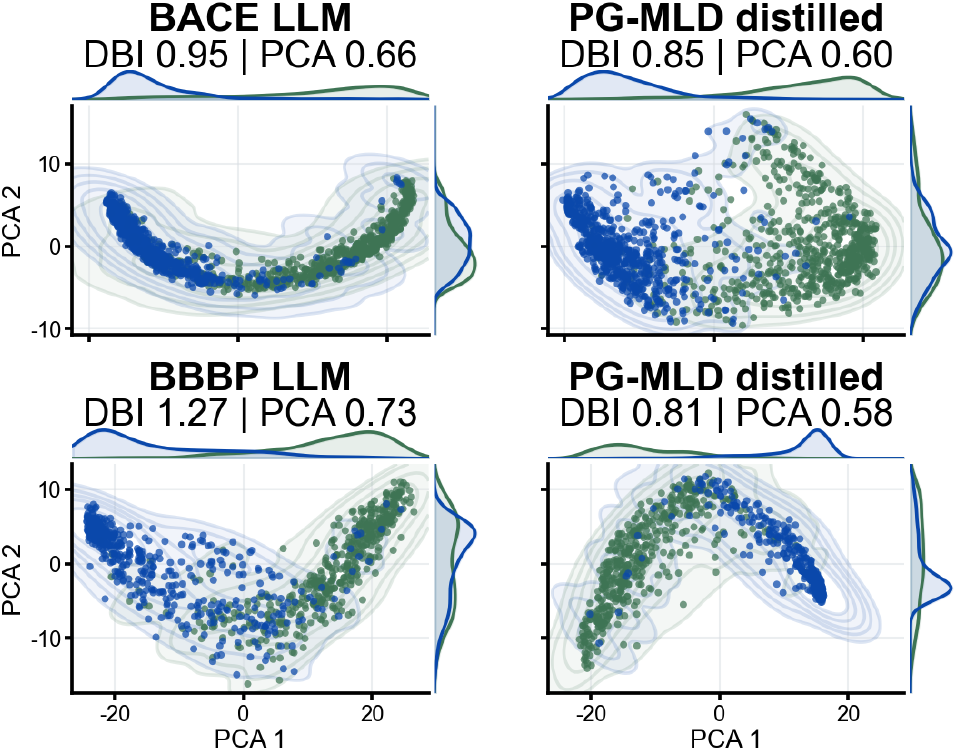
PCA projections of molecular representations before and after PG-MLD distillation. Colors denote classes, and marginal curves show class densities. Lower DBI and PCA-space class-overlap scores indicate better separation.

## Conclusion

We present PG-MLD, a dynamic 3D physical knowledge distillation paradigm for molecular language models. The 3D teacher combines equivariant geometric encoding with Liquid Time-Constant modeling. PG-MLD transfers the teacher’s trajectory-derived knowledge through atom- and molecule-level representation alignment and cross-modal contrastive learning, together with masked language modeling where supported. Across MoleculeNet tasks and three student architectures, PG-MLD consistently improves property prediction while retaining SMILES-only inference. Representation analyses further indicate that the distilled models capture 3D geometry and conformational dynamics more effectively. Future work will incorporate richer quantum-chemical and multiscale dynamical information and extend PG-MLD to broader downstream tasks in drug discovery, complex biological system modeling, and materials design.

