## Supplementary Material for "PG-MLD: Physics-Guided Molecular Representation Learning via Dynamic 3D Trajectory Distillation"

**Zihe Liu**[1], **Zhonghan Wu**[1], **Zhuo Chen**[1,*],
**Xin Gao**[2], **Bin Yu**[1,3,*]
[1]School of Data Science, Qingdao University of Science and Technology, Qingdao 266061, China
[2]Computational Bioscience Research Center (CBRC), Computer, Electrical and Mathematical Sciences and Engineering Division, King Abdullah University of Science and Technology (KAUST), Thuwal 23955, Saudi Arabia
[3]School of Artificial Intelligence and Data Science, University of Science and Technology of China, Hefei 230027, China
[*]Corresponding authors

#### A Dataset and Evaluation Details

We evaluate PG-MLD on nine molecular property prediction datasets from MoleculeNet (Wu et al. 2018), a widely used benchmark spanning diverse classification and regression tasks. Table 1 summarizes the dataset size, task type, and evaluation metric. BBBP, BACE, ClinTox, Tox21, ToxCast, and HIV are classification datasets evaluated using ROC-AUC, whereas ESOL, FreeSolv, and Lipophilicity are regression datasets evaluated using RMSE. ROC-AUC measures classification performance across decision thresholds, while RMSE quantifies regression error and penalizes larger deviations. For every dataset, molecules are randomly divided into training, validation, and test sets using an 80%/10%/10% ratio. For each seed, the same split is used when comparing an original student with its PG-MLD-distilled counterpart. Model selection and early stopping are performed exclusively on the validation set, and the test set is used only for final evaluation. Each experiment is repeated with five random seeds (42, 43, 44, 45, and 46), and we report the mean and standard deviation of the test-set performance.

| Dataset | Molecules | Task type | Metric |
|---|---|---|---|
| BBBP | 2,039 | Classification | ROC-AUC ↑ |
| BACE | 1,513 | Classification | ROC-AUC ↑ |
| ClinTox | 1,478 | Classification | ROC-AUC ↑ |
| Tox21 | 7,831 | Classification | ROC-AUC ↑ |
| ToxCast | 8,575 | Classification | ROC-AUC ↑ |
| HIV | 41,127 | Classification | ROC-AUC ↑ |
| ESOL | 1,427 | Regression | RMSE ↓ |
| FreeSolv | 642 | Regression | RMSE ↓ |
| Lipophilicity | 4,200 | Regression | RMSE ↓ |

Table 1: Statistics and evaluation protocols for the MoleculeNet datasets used in this study.

#### B Baseline Details

To evaluate PG-MLD, we select representative molecular property prediction methods spanning graph self-supervised learning, geometry-aware pretraining, knowledge-enhanced learning, hybrid graph–Transformer architectures, and SMILES-based molecular language modeling. The comparisons follow the random-split protocol and evaluation metrics described above.

**GraphMVP** (Liu et al. 2022). GraphMVP is a multi-view molecular pretraining framework that exploits the correspondence between 2D molecular topology and 3D geometry. It transfers geometric information into a 2D graph encoder by encouraging consistency between representations learned from the two views.

**MolCLR** (Wang et al. 2022). MolCLR is a graph contrastive learning framework pretrained on large-scale unlabeled molecular data. It constructs positive pairs through molecular graph augmentations, including atom masking, bond deletion, and subgraph removal, and learns representations by maximizing agreement between augmented views of the same molecule.

**GEM** (Fang et al. 2022). GEM combines a geometry-based graph neural network with geometry-level self-supervised objectives. Its GeoGNN jointly models atoms, bonds, and bond angles, allowing molecular topology and coarse 3D geometry to contribute to property prediction.

**KPGT** (Li et al. 2023). KPGT is a knowledge-guided graph Transformer pretraining framework built on a line-graph backbone. It uses molecular descriptors and fingerprints as supervision in addition to masked-node prediction, thereby incorporating explicit chemical knowledge into the learned representation.

**Galformer** (Bai et al. 2026). Galformer employs a geometry-aware line-graph Transformer to encode complementary 2D topological and 3D geometric views. Its self-supervised objectives model both cross-modal correspondence and within-modal structural information.

**KANO** (Fang et al. 2023). KANO integrates chemical knowledge graphs with molecular contrastive learning. Element-level knowledge guides graph augmentation during pretraining, while functional-group prompts adapt the pretrained encoder to downstream molecular property prediction tasks.

**MoleculeFormer** (Qin et al. 2025). MoleculeFormer is a GCN–Transformer architecture that extracts multi-scale features from atom- and bond-level molecular graphs. It further incorporates 3D geometric constraints and molecular fingerprints to combine local structural information with long-range dependencies.

**MolGramTreeNet** (Zhang et al. 2026). MolGramTreeNet is a multimodal dual-path framework that combines molecular graphs with grammar-tree representations. The grammar-tree branch explicitly captures chemical construction rules and hierarchical substructures, complementing the local connectivity encoded by the graph branch.

The following pretrained molecular language models are additionally used as student architectures to evaluate whether PG-MLD transfers dynamic 3D physical knowledge across different sequence-modeling backbones.

**ChemBERTa** (Chithrananda, Grand, and Ramsundar 2020). ChemBERTa treats SMILES strings as molecular language and applies Transformer-based self-supervised pretraining to learn contextual representations that can be fine-tuned for molecular property prediction.

**ChemBERTa-2** (Ahmad et al. 2022). ChemBERTa-2 extends ChemBERTa through larger-scale and optimized chemical pretraining. It investigates both masked language modeling and multitask property-prediction objectives for learning transferable SMILES representations.

**MoLFormer** (Ross et al. 2022). MoLFormer is a large-scale molecular language model that combines linear attention with rotary positional embeddings. It is pretrained by masked language modeling on large SMILES corpora to provide efficient and transferable molecular representations.

### C Training Procedure

Algorithm 1 summarizes the two-stage optimization procedure of PG-MLD, including dynamic 3D teacher pretraining and subsequent 3D-to-1D knowledge distillation.

**Algorithm 1** PG-MLD Dynamic 3D-to-1D Distillation

**Require:** SMILES $S$, molecular graph $G$, trajectory $\{\mathbf{X}_t\}_{t=1}^{T}$, atomic descriptors $\mathbf{E}$
**Ensure:** Distilled molecular language model $f_\psi$
Initialize 3D teacher $f_\theta$ and SMILES student $f_\psi$
**for** each teacher pretraining batch **do**
  **for** $t = 1, \ldots, T$ **do**
    $\mathbf{Z}_t^{3D} \leftarrow f_{\text{eq}}(G, \mathbf{X}_t, \mathbf{E})$
    $\mathbf{c}_t \leftarrow [\mathbf{W}_{\text{in}}\mathbf{Z}_t^{3D} \| \mathbf{h}_{t-1}]$
    $\bar{\mathbf{h}}_t \leftarrow g_\theta(\mathbf{c}_t)$,
    $\boldsymbol{\tau}_t \leftarrow \text{softplus}(q_\theta(\mathbf{c}_t)) + \epsilon$
    $\mathbf{h}_t \leftarrow \bar{\mathbf{h}}_t + (\mathbf{h}_{t-1} - \bar{\mathbf{h}}_t) \odot e^{-\Delta t/\boldsymbol{\tau}_t}$
  **end for**
  Compute teacher self-supervised loss $\mathcal{L}_{\text{3D}}$
  Update teacher parameters $\theta$
**end for**
Freeze teacher parameters $\theta$
**for** each distillation batch **do**
  $(\mathbf{H}_{\text{atom}}^{3D}, \mathbf{h}_{\text{mol}}^{3D}) \leftarrow f_\theta(G, \mathbf{X}_{1:T}, \mathbf{E})$
  $\mathbf{H}^{1D} \leftarrow f_\psi(S)$
  $\mathbf{H}_{\text{atom}}^{1D} \leftarrow \text{TokenAtomAgg}(\mathbf{H}^{1D})$
  $\mathbf{h}_{\text{mol}}^{1D} \leftarrow \text{Pool}(\mathbf{H}_{\text{atom}}^{1D})$
  $\mathcal{L}_{\text{distill}} \leftarrow \lambda_a \mathcal{L}_{\text{atom}} + \lambda_m \mathcal{L}_{\text{mol}} + \lambda_c \mathcal{L}_{\text{con}} + \lambda_{\text{MLM}} \mathcal{L}_{\text{MLM}}$
  Update student $\psi$ and projection heads
**end for**
**return** $f_\psi$

### D Implementation and Hyperparameters

All models are optimized using AdamW, and teacher pretraining and distillation use a cosine learning-rate schedule with linear warm-up. The 3D teacher contains four equivariant geometric layers with eight attention heads; its geometric and LTC representations are both 384-dimensional. ChemBERTa and ChemBERTa-2 have a hidden size of 384, whereas MoLFormer has a hidden size of 768. Architecture-specific projection heads map the student representations into the shared teacher space. The main optimization and architecture settings are summarized in Table 2.

For all students, we set $\lambda_a = \lambda_m = 1.0$. For ChemBERTa and ChemBERTa-2, $\lambda_c = 0.1$, the contrastive temperature is 0.1, $\lambda_{\text{MLM}} = 0.1$, and the masking probability is 0.15. For MoLFormer, the corresponding values are 0.05, 0.2, 0, and 0, respectively. Teacher pretraining and distillation use seed 2026. During downstream fine-tuning, the full student encoder and a task-specific prediction head are jointly optimized, with model selection based on validation performance over five runs using seeds 42–46.

### E Dynamic 3D Trajectory Construction

The dynamic 3D pretraining data are constructed from 250,000 unlabeled molecules sampled from ZINC15 (Sterling and Irwin 2015). For each molecule, RDKit (RDKit contributors 2026) generates and optimizes an initial conformer, after which OpenMM (Eastman et al. 2017) and the OpenFF force field (Boothroyd et al. 2023) are used to produce an MD trajectory. At training time, PG-MLD samples at most eight temporally ordered frames to preserve conformational evolution while controlling computational cost.

CGN and CAN provide complementary coordinate- and dihedral-level perturbations. Formal charges and Gasteiger partial charges describe the local electronic environment of each atom. The trajectory and perturbation configurations are summarized in Table 3. Trajectory generation uses deterministic molecule-specific seeds derived from seed 2026.

### F Component Ablation Results

Table 4 evaluates the contributions of molecular dynamics trajectories, atom-level electronic descriptors, and CGN/CAN geometric perturbations using MoLFormer as the student model. Here, "w/o" denotes that the corresponding component is removed while the remaining training configuration is kept unchanged.

Removing MD trajectories causes the largest overall degradation across the nine tasks, including increases in FreeSolv and Lipophilicity RMSE from 0.945 to 0.988 and from 0.620 to 0.644, respectively. These results indicate that temporally ordered conformations contribute most strongly among the evaluated components. Removing either the electronic descriptors or CGN/CAN produces smaller but consistent performance decreases across all tasks, suggesting stable yet comparatively modest complementary gains. The full model performs best on every task in this comparison, supporting the joint use of dynamic trajectories, electronic descriptors, and geometric perturbations.

| Stage | Learning rate | Batch size | Epochs | Weight decay | Hidden size | Seed |
|---|---|---|---|---|---|---|
| 3D teacher pretraining | $2 \times 10^{-4}$ | 4 | 10 | 0.01 | 384 | 2026 |
| 3D-to-1D distillation | $2 \times 10^{-5}$ | 8 | 10 | 0.01 | 384/768 | 2026 |
| Downstream fine-tuning | $5 \times 10^{-6}$–$2 \times 10^{-5}$ | 16 | $\leq 60$ | 0.01 | 256 (head) | 42–46 (5 runs) |

Table 2: Main optimization and architecture hyperparameters. The 384/768 entry denotes the native hidden sizes of the molecular language students.

| Component | Setting |
|---|---|
| Initial conformer | RDKit ETKDGv3; MMFF94 optimization with UFF fallback |
| MD engine and force field | OpenMM; OpenFF unconstrained 2.1.0 |
| Integrator | Langevin middle, 300 K, 1 $\text{ps}^{-1}$ friction, 1 fs time step |
| Simulation length | 1,000 equilibration steps + 1,000 production steps |
| Trajectory sampling | Every 100 steps; at most 8 ordered frames during training |
| CGN | Gaussian coordinate noise, $\sigma = 0.02$ Å; 4 views |
| CAN | Rotatable dihedral perturbation, maximum 30°; 4 views |
| Electronic descriptors | Formal charge and Gasteiger partial charge |

Table 3: Configuration used to construct the dynamic 3D physical views.

| Method | BBBP↑ | BACE↑ | ClinTox↑ | Tox21↑ | ToxCast↑ | HIV↑ | ESOL↓ | FreeSolv↓ | Lipo.↓ |
|---|---|---|---|---|---|---|---|---|---|
| Dataset size | 2,039 | 1,513 | 1,478 | 7,831 | 8,575 | 41,127 | 1,427 | 642 | 4,200 |
| Original student | 0.957(.012) | 0.913(.011) | 0.986(.006) | 0.839(.016) | 0.724(.025) | 0.794(.019) | 0.677(.026) | 1.011(.214) | 0.662(.025) |
| w/o MD trajectories | 0.959(.008) | 0.915(.010) | 0.988(.005) | 0.842(.010) | 0.735(.018) | 0.810(.016) | 0.665(.025) | 0.988(.175) | 0.644(.021) |
| w/o electronic descriptors | 0.960(.010) | 0.919(.012) | 0.990(.007) | 0.846(.006) | 0.741(.014) | 0.819(.012) | 0.660(.021) | 0.958(.155) | 0.635(.021) |
| w/o CGN/CAN | 0.961(.006) | 0.917(.011) | 0.991(.005) | 0.845(.006) | 0.742(.012) | 0.818(.015) | 0.659(.023) | 0.954(.157) | 0.631(.019) |
| Full PG-MLD | **0.964(.005)** | **0.931(.008)** | **0.997(.002)** | **0.850(.005)** | **0.744(.013)** | **0.825(.011)** | **0.653(.019)** | **0.945(.148)** | **0.620(.017)** |

Table 4: Component ablation results on the MoleculeNet property prediction tasks using MoLFormer as the student model. Results are reported as mean (standard deviation), using ROC-AUC for classification tasks (higher is better) and RMSE for regression tasks (lower is better). The best and second-best results are shown in **bold** and underlined, respectively.
