## Supplementary material for "PG-MLD: Physics-Guided Molecular Representation Learning via Dynamic 3D Trajectory Distillation": Code and Data: dynamic_md_dataset.py

```
from __future__ import annotations

import glob
import random
import warnings
from pathlib import Path
from typing import Any

import torch
from rdkit import Chem
from torch.utils.data import Dataset, IterableDataset, get_worker_info

from comld.data.featurizer import atom_features, bond_features


def load_dynamic_md_items(path: str | Path) -> list[dict[str, Any]]:
    path = Path(path)
    if not path.exists():
        raise FileNotFoundError(f"Dynamic MD dataset not found: {path}")
    try:
        items = torch.load(path, map_location="cpu", weights_only=False)
    except TypeError:
        items = torch.load(path, map_location="cpu")
    if not isinstance(items, list) or not items:
        raise ValueError(f"No dynamic MD molecules found in {path}")
    return items


def _heavy_atom_indices(atom_symbols: list[str]) -> torch.Tensor:
    indices = [idx for idx, symbol in enumerate(atom_symbols) if str(symbol).upper() != "H"]
    return torch.tensor(indices, dtype=torch.long)


def _build_heavy_atom_graph(
    smiles: str,
    electronic_feats: torch.Tensor,
) -> tuple[torch.Tensor, torch.Tensor, torch.Tensor]:
    mol = Chem.MolFromSmiles(smiles)
    if mol is None:
        raise ValueError(f"Invalid SMILES in dynamic MD dataset: {smiles}")
    if mol.GetNumAtoms() != electronic_feats.shape[0]:
        raise ValueError(
            f"Heavy-atom mismatch for {smiles}: RDKit={mol.GetNumAtoms()} "
            f"trajectory={electronic_feats.shape[0]}"
        )

    node_rows = []
    for atom_idx, atom in enumerate(mol.GetAtoms()):
        gasteiger = float(electronic_feats[atom_idx, 1].item())
        atom.SetProp("_GasteigerCharge", str(gasteiger))
        node_rows.append(atom_features(atom))
    node_feats = torch.tensor(node_rows, dtype=torch.float)
    node_feats = torch.cat([node_feats, electronic_feats.float()], dim=-1)

    edge_pairs: list[list[int]] = []
    edge_rows: list[list[float]] = []
    for bond in mol.GetBonds():
        begin = int(bond.GetBeginAtomIdx())
        end = int(bond.GetEndAtomIdx())
        features = bond_features(bond)
        edge_pairs.extend([[begin, end], [end, begin]])
        edge_rows.extend([features, features])

    if edge_pairs:
        edge_index = torch.tensor(edge_pairs, dtype=torch.long).t().contiguous()
        edge_attr = torch.tensor(edge_rows, dtype=torch.float)
    else:
        edge_index = torch.empty((2, 0), dtype=torch.long)
        edge_dim = len(bond_features(Chem.MolFromSmiles("CC").GetBondWithIdx(0)))
        edge_attr = torch.empty((0, edge_dim), dtype=torch.float)
    return node_feats, edge_index, edge_attr


class DynamicMDTrajectoryDataset(Dataset):
    """Loads real OpenMM trajectories and keeps only SMILES-aligned heavy atoms."""

    def __init__(
        self,
        path: str | Path,
        max_molecules: int | None = None,
        max_frames: int | None = None,
        frame_stride: int = 1,
        random_frame_crop: bool = False,
    ) -> None:
        self.path = Path(path)
        self.max_frames = int(max_frames) if max_frames is not None else None
        self.random_frame_crop = bool(random_frame_crop)
        items = load_dynamic_md_items(self.path)
        if max_molecules is not None:
            items = items[: int(max_molecules)]

        self.graphs: list[dict[str, Any]] = []
        stride = max(int(frame_stride), 1)
        for item in items:
            coords = torch.as_tensor(item["coords"], dtype=torch.float)
            electronic = torch.as_tensor(item["electronic_feats"], dtype=torch.float)
            symbols = [str(symbol) for symbol in item["atom_symbols"]]
            if coords.ndim != 3 or coords.shape[-1] != 3:
                raise ValueError(f"Expected trajectory [T,N,3], got {tuple(coords.shape)}")
            if electronic.ndim != 2 or electronic.shape[-1] != 2:
                raise ValueError(f"Expected electronic features [N,2], got {tuple(electronic.shape)}")
            if coords.shape[1] != len(symbols) or electronic.shape[0] != len(symbols):
                raise ValueError(f"Atom dimensions disagree for row {item.get('row_idx')}")

            heavy_indices = _heavy_atom_indices(symbols)
            heavy_symbols = [symbols[int(idx)] for idx in heavy_indices.tolist()]
            coords = coords.index_select(1, heavy_indices)[::stride]
            electronic = electronic.index_select(0, heavy_indices)
            if self.max_frames is not None and not self.random_frame_crop:
                coords = coords[: self.max_frames]
            if coords.shape[0] == 0:
                continue
            if not torch.isfinite(coords).all() or not torch.isfinite(electronic).all():
                continue

            smiles = str(item["smiles"])
            rdkit_mol = Chem.MolFromSmiles(smiles)
            rdkit_symbols = (
                [atom.GetSymbol() for atom in rdkit_mol.GetAtoms()]
                if rdkit_mol is not None
                else []
            )
            if rdkit_symbols != heavy_symbols:
                raise ValueError(
                    f"Heavy-atom order mismatch for row {item.get('row_idx')}: "
                    f"SMILES={rdkit_symbols}, trajectory={heavy_symbols}"
                )
            node_feats, edge_index, edge_attr = _build_heavy_atom_graph(smiles, electronic)
            timestep_fs = float(item.get("timestep_fs", 1.0))
            report_interval = int(item.get("report_interval", 1))
            delta_time_ps = timestep_fs * report_interval * stride / 1000.0
            self.graphs.append(
                {
                    "row_idx": int(item.get("row_idx", len(self.graphs))),
                    "smiles": smiles,
                    "num_atoms": int(node_feats.shape[0]),
                    "num_frames": int(coords.shape[0]),
                    "coords": coords.contiguous(),
                    "node_feats": node_feats,
                    "edge_index": edge_index,
                    "edge_attr": edge_attr,
                    "electronic_feats": electronic,
                    "delta_time_ps": float(delta_time_ps),
                }
            )

        if not self.graphs:
            raise ValueError(f"No valid dynamic MD trajectories found in {self.path}")

    def __len__(self) -> int:
        return len(self.graphs)

    def __getitem__(self, index: int) -> dict[str, Any]:
        graph = self.graphs[index]
        if (
            self.max_frames is None
            or not self.random_frame_crop
            or graph["coords"].shape[0] <= self.max_frames
        ):
            return graph

        max_start = int(graph["coords"].shape[0]) - self.max_frames
        start = int(torch.randint(0, max_start + 1, (1,)).item())
        cropped = dict(graph)
        cropped["coords"] = graph["coords"][start : start + self.max_frames]
        cropped["num_frames"] = int(cropped["coords"].shape[0])
        return cropped


def resolve_dynamic_md_shards(path: str | Path) -> list[Path]:
    path_text = str(path)
    candidate = Path(path_text)
    if candidate.is_file():
        return [candidate]
    if candidate.is_dir():
        direct = sorted(candidate.glob("md_cgn_can_electronic.pt"))
        nested = sorted(candidate.glob("shard_*/md_cgn_can_electronic.pt"))
        shards = direct + nested
    elif any(token in path_text for token in "*?[]"):
        shards = sorted(Path(match) for match in glob.glob(path_text))
    else:
        shards = []
    shards = [shard for shard in shards if shard.is_file() and shard.stat().st_size > 0]
    if not shards:
        raise FileNotFoundError(
            f"No dynamic MD shard found for '{path}'. Expected a .pt file, a directory "
            "containing shard_*/md_cgn_can_electronic.pt, or a glob pattern."
        )
    return shards


class ShardedDynamicMDTrajectoryDataset(IterableDataset):
    """Streams one packed trajectory shard at a time to keep memory bounded."""

    def __init__(
        self,
        path: str | Path,
        max_molecules: int | None = None,
        max_frames: int | None = None,
        frame_stride: int = 1,
        random_frame_crop: bool = False,
        seed: int = 2026,
        shuffle_shards: bool = True,
        shuffle_molecules: bool = True,
    ) -> None:
        super().__init__()
        self.path = str(path)
        self.shards = resolve_dynamic_md_shards(path)
        self.max_molecules = int(max_molecules) if max_molecules is not None else None
        self.max_frames = max_frames
        self.frame_stride = int(frame_stride)
        self.random_frame_crop = bool(random_frame_crop)
        self.seed = int(seed)
        self.shuffle_shards = bool(shuffle_shards)
        self.shuffle_molecules = bool(shuffle_molecules)
        self.epoch = 0
        self._sample: dict[str, Any] | None = None

    def set_epoch(self, epoch: int) -> None:
        self.epoch = int(epoch)

    def sample_item(self) -> dict[str, Any]:
        if self._sample is None:
            errors = []
            for shard in self.shards:
                try:
                    dataset = DynamicMDTrajectoryDataset(
                        shard,
                        max_molecules=1,
                        max_frames=self.max_frames,
                        frame_stride=self.frame_stride,
                        random_frame_crop=False,
                    )
                    self._sample = dataset[0]
                    break
                except (OSError, RuntimeError, EOFError, ValueError) as exc:
                    errors.append(f"{shard}: {exc}")
            if self._sample is None:
                preview = "\n".join(errors[:5])
                raise RuntimeError(
                    "No readable dynamic MD shard is available. "
                    f"First errors:\n{preview}"
                )
        return self._sample

    def __len__(self) -> int:
        if self.max_molecules is not None:
            return self.max_molecules
        # Preprocessing uses 1,000 molecules per shard except the final shard.
        # This is an estimate used only for progress reporting.
        return len(self.shards) * 1000

    def __iter__(self):
        worker = get_worker_info()
        worker_id = worker.id if worker is not None else 0
        num_workers = worker.num_workers if worker is not None else 1
        rng = random.Random(self.seed + self.epoch * 1009 + worker_id)
        shards = list(self.shards)
        if self.shuffle_shards:
            rng.shuffle(shards)
        shards = shards[worker_id::num_workers]

        yielded = 0
        worker_limit = None
        if self.max_molecules is not None:
            worker_limit = (self.max_molecules + num_workers - 1) // num_workers

        for shard in shards:
            try:
                dataset = DynamicMDTrajectoryDataset(
                    shard,
                    max_frames=self.max_frames,
                    frame_stride=self.frame_stride,
                    random_frame_crop=self.random_frame_crop,
                )
            except (OSError, RuntimeError, EOFError, ValueError) as exc:
                warnings.warn(
                    f"Skipping unreadable dynamic MD shard '{shard}': {exc}",
                    RuntimeWarning,
                )
                continue
            indices = list(range(len(dataset)))
            if self.shuffle_molecules:
                rng.shuffle(indices)
            for index in indices:
                if worker_limit is not None and yielded >= worker_limit:
                    return
                yielded += 1
                yield dataset[index]


def build_dynamic_md_dataset(
    path: str | Path,
    max_molecules: int | None = None,
    max_frames: int | None = None,
    frame_stride: int = 1,
    random_frame_crop: bool = False,
    seed: int = 2026,
) -> Dataset | IterableDataset:
    shards = resolve_dynamic_md_shards(path)
    if len(shards) == 1 and Path(path).is_file():
        return DynamicMDTrajectoryDataset(
            shards[0],
            max_molecules=max_molecules,
            max_frames=max_frames,
            frame_stride=frame_stride,
            random_frame_crop=random_frame_crop,
        )
    return ShardedDynamicMDTrajectoryDataset(
        path=path,
        max_molecules=max_molecules,
        max_frames=max_frames,
        frame_stride=frame_stride,
        random_frame_crop=random_frame_crop,
        seed=seed,
    )


def dynamic_md_sample(dataset: Dataset | IterableDataset) -> dict[str, Any]:
    if isinstance(dataset, ShardedDynamicMDTrajectoryDataset):
        return dataset.sample_item()
    return dataset[0]


def collate_dynamic_md(items: list[dict[str, Any]]) -> dict[str, Any]:
    frame_node_feats = []
    frame_coords = []
    frame_edge_indices = []
    frame_edge_attrs = []
    frame_batch = []
    atom_batch = []
    electronic_feats = []
    trajectory_specs = []
    trajectory_frame_ptr = [0]
    smiles = []
    atom_counts = []
    delta_times_ps = []

    node_offset = 0
    frame_index = 0
    for mol_idx, graph in enumerate(items):
        num_atoms = int(graph["num_atoms"])
        num_frames = int(graph["num_frames"])
        trajectory_specs.append((num_frames, num_atoms))
        trajectory_frame_ptr.append(trajectory_frame_ptr[-1] + num_frames)
        smiles.append(graph["smiles"])
        atom_counts.append(num_atoms)
        delta_times_ps.append(float(graph["delta_time_ps"]))
        atom_batch.append(torch.full((num_atoms,), mol_idx, dtype=torch.long))
        electronic_feats.append(graph["electronic_feats"])

        for time_idx in range(num_frames):
            frame_node_feats.append(graph["node_feats"])
            frame_coords.append(graph["coords"][time_idx])
            frame_batch.append(torch.full((num_atoms,), frame_index, dtype=torch.long))
            if graph["edge_index"].numel() > 0:
                frame_edge_indices.append(graph["edge_index"] + node_offset)
                frame_edge_attrs.append(graph["edge_attr"])
            node_offset += num_atoms
            frame_index += 1

    if frame_edge_indices:
        edge_index = torch.cat(frame_edge_indices, dim=1)
        edge_attr = torch.cat(frame_edge_attrs, dim=0)
    else:
        edge_index = torch.empty((2, 0), dtype=torch.long)
        edge_attr = torch.empty((0, items[0]["edge_attr"].shape[-1]), dtype=torch.float)

    return {
        "smiles": smiles,
        "atom_counts": torch.tensor(atom_counts, dtype=torch.long),
        "node_feats": torch.cat(frame_node_feats, dim=0).float(),
        "coords": torch.cat(frame_coords, dim=0).float(),
        "edge_index": edge_index.long(),
        "edge_attr": edge_attr.float(),
        "frame_batch": torch.cat(frame_batch, dim=0).long(),
        "atom_batch": torch.cat(atom_batch, dim=0).long(),
        "electronic_feats": torch.cat(electronic_feats, dim=0).float(),
        "trajectory_specs": trajectory_specs,
        "trajectory_frame_ptr": torch.tensor(trajectory_frame_ptr, dtype=torch.long),
        "delta_times_ps": torch.tensor(delta_times_ps, dtype=torch.float),
    }
```
