## Supplementary material for "PG-MLD: Physics-Guided Molecular Representation Learning via Dynamic 3D Trajectory Distillation": Code and Data: featurizer.py

```
from __future__ import annotations

import math

from rdkit import Chem


COMMON_ATOMIC_NUMBERS = [1, 5, 6, 7, 8, 9, 14, 15, 16, 17, 35, 53]
FORMAL_CHARGES = [-2, -1, 0, 1, 2]
DEGREES = [0, 1, 2, 3, 4, 5]
TOTAL_HS = [0, 1, 2, 3, 4]
HYBRIDIZATIONS = [
    Chem.rdchem.HybridizationType.SP,
    Chem.rdchem.HybridizationType.SP2,
    Chem.rdchem.HybridizationType.SP3,
    Chem.rdchem.HybridizationType.SP3D,
    Chem.rdchem.HybridizationType.SP3D2,
]
BOND_TYPES = [
    Chem.rdchem.BondType.SINGLE,
    Chem.rdchem.BondType.DOUBLE,
    Chem.rdchem.BondType.TRIPLE,
    Chem.rdchem.BondType.AROMATIC,
]
STEREO_TYPES = [
    Chem.rdchem.BondStereo.STEREONONE,
    Chem.rdchem.BondStereo.STEREOANY,
    Chem.rdchem.BondStereo.STEREOZ,
    Chem.rdchem.BondStereo.STEREOE,
]


def one_hot_with_unknown(value, choices) -> list[float]:
    values = [float(value == choice) for choice in choices]
    values.append(float(value not in choices))
    return values


def safe_float(value: str, default: float = 0.0) -> float:
    try:
        number = float(value)
    except (TypeError, ValueError):
        return default
    if math.isnan(number) or math.isinf(number):
        return default
    return number


def atom_features(atom: Chem.Atom) -> list[float]:
    charge = safe_float(
        atom.GetProp("_GasteigerCharge")
        if atom.HasProp("_GasteigerCharge")
        else "0"
    )
    charge = max(min(charge, 2.0), -2.0)
    features: list[float] = []
    features += one_hot_with_unknown(atom.GetAtomicNum(), COMMON_ATOMIC_NUMBERS)
    features += one_hot_with_unknown(atom.GetTotalDegree(), DEGREES)
    features += one_hot_with_unknown(atom.GetFormalCharge(), FORMAL_CHARGES)
    features += one_hot_with_unknown(atom.GetHybridization(), HYBRIDIZATIONS)
    features += one_hot_with_unknown(atom.GetTotalNumHs(), TOTAL_HS)
    features += [
        float(atom.GetIsAromatic()),
        float(atom.IsInRing()),
        atom.GetMass() * 0.01,
        charge,
    ]
    return features


def bond_features(bond: Chem.Bond) -> list[float]:
    features: list[float] = []
    features += one_hot_with_unknown(bond.GetBondType(), BOND_TYPES)
    features += one_hot_with_unknown(bond.GetStereo(), STEREO_TYPES)
    features += [
        float(bond.GetIsConjugated()),
        float(bond.IsInRing()),
    ]
    return features
```
