## Supplementary material for "PG-MLD: Physics-Guided Molecular Representation Learning via Dynamic 3D Trajectory Distillation": Code and Data: chemberta_1d.py

```
from __future__ import annotations

from pathlib import Path
from typing import Sequence

import torch
from torch import nn

from comld.models.chemberta_teacher import mean_pool_tokens, resolve_model_name_or_path


def attention_mean_pool(hidden: torch.Tensor, attention_mask: torch.Tensor) -> torch.Tensor:
    weights = attention_mask.bool().unsqueeze(-1).to(hidden.dtype)
    pooled = (hidden * weights).sum(dim=1)
    denom = weights.sum(dim=1).clamp_min(1.0)
    return pooled / denom


class ChemBERTaMoleculeEncoder(nn.Module):
    def __init__(
        self,
        model_name: str = "DeepChem/ChemBERTa-77M-MLM",
        trust_remote_code: bool | None = None,
    ) -> None:
        super().__init__()
        from transformers import AutoModel, AutoTokenizer

        model_name_or_path = resolve_model_name_or_path(model_name)
        model_path = Path(model_name_or_path)
        local_files_only = model_path.exists()
        if trust_remote_code is None:
            lower_name = str(model_name_or_path).lower()
            trust_remote_code = "molformer" in lower_name or (
                local_files_only and any(model_path.glob("modeling_*.py"))
            )
        self.tokenizer = AutoTokenizer.from_pretrained(
            model_name_or_path,
            use_fast=True,
            local_files_only=local_files_only,
            trust_remote_code=trust_remote_code,
        )
        self.model = AutoModel.from_pretrained(
            model_name_or_path,
            local_files_only=local_files_only,
            trust_remote_code=trust_remote_code,
        )
        self.model_name = model_name_or_path
        self.trust_remote_code = bool(trust_remote_code)

    @property
    def hidden_size(self) -> int:
        for name in ("hidden_size", "n_embd", "d_model", "embedding_size"):
            value = getattr(self.model.config, name, None)
            if value is not None:
                return int(value)
        raise AttributeError(f"Cannot infer hidden size from model config: {self.model.config}")

    def forward(self, smiles: Sequence[str], device: torch.device | str | None = None) -> torch.Tensor:
        if device is None:
            device = next(self.model.parameters()).device
        device = torch.device(device)
        token_kwargs = {
            "padding": True,
            "truncation": True,
            "return_tensors": "pt",
        }
        if getattr(self.tokenizer, "is_fast", False):
            token_kwargs["return_offsets_mapping"] = True
        tokens = self.tokenizer(list(smiles), **token_kwargs)
        offsets = tokens.pop("offset_mapping", None)
        if offsets is not None:
            offsets = offsets.to(device)
        model_inputs = {key: value.to(device) for key, value in tokens.items()}
        output = self.model(**model_inputs)
        token_hidden = output.last_hidden_state if hasattr(output, "last_hidden_state") else output[0]
        if offsets is not None:
            return mean_pool_tokens(token_hidden, model_inputs["attention_mask"], offsets)
        return attention_mean_pool(token_hidden, model_inputs["attention_mask"])


class ChemBERTaMoleculeClassifier(nn.Module):
    def __init__(
        self,
        encoder: ChemBERTaMoleculeEncoder,
        output_dim: int = 1,
        head_hidden_dim: int = 256,
        dropout: float = 0.1,
    ) -> None:
        super().__init__()
        self.encoder = encoder
        self.head = nn.Sequential(
            nn.Linear(encoder.hidden_size, head_hidden_dim),
            nn.SiLU(),
            nn.Dropout(dropout),
            nn.Linear(head_hidden_dim, int(output_dim)),
        )

    def forward(self, smiles: Sequence[str], device: torch.device | str | None = None) -> torch.Tensor:
        mol_embeddings = self.encoder(smiles, device=device)
        return self.head(mol_embeddings)
```
