## Supplementary material for "PG-MLD: Physics-Guided Molecular Representation Learning via Dynamic 3D Trajectory Distillation": Code and Data: chemberta_teacher.py

```
from __future__ import annotations

import re
from pathlib import Path
from typing import Sequence

import torch
from torch import nn
import torch.nn.functional as F


ATOM_SPAN_PATTERN = re.compile(r"\[[^\[\]]+\]|Br|Cl|B|C|N|O|P|S|F|I|b|c|n|o|p|s")


def find_smiles_atom_spans(smiles: str) -> list[tuple[int, int]]:
    return [match.span() for match in ATOM_SPAN_PATTERN.finditer(smiles)]


def mean_pool_tokens(hidden: torch.Tensor, attention_mask: torch.Tensor, offsets: torch.Tensor) -> torch.Tensor:
    valid = attention_mask.bool() & (offsets[..., 1] > offsets[..., 0]).to(attention_mask.device)
    weights = valid.unsqueeze(-1).to(hidden.dtype)
    pooled = (hidden * weights).sum(dim=1)
    denom = weights.sum(dim=1).clamp_min(1.0)
    return pooled / denom


def pool_tokens_to_atoms(
    token_hidden: torch.Tensor,
    offsets: torch.Tensor,
    attention_mask: torch.Tensor,
    smiles: Sequence[str],
    atom_counts: torch.Tensor,
    mol_embeddings: torch.Tensor,
) -> torch.Tensor:
    atom_embeddings: list[torch.Tensor] = []
    offsets_cpu = offsets.detach().cpu()
    attention_cpu = attention_mask.detach().cpu().bool()
    for batch_idx, smi in enumerate(smiles):
        spans = find_smiles_atom_spans(smi)
        n_atoms = int(atom_counts[batch_idx].item())
        for atom_idx in range(n_atoms):
            if atom_idx >= len(spans):
                atom_embeddings.append(mol_embeddings[batch_idx])
                continue

            start, end = spans[atom_idx]
            token_hits = []
            for token_idx, (tok_start, tok_end) in enumerate(offsets_cpu[batch_idx].tolist()):
                if not attention_cpu[batch_idx, token_idx]:
                    continue
                if tok_end <= tok_start:
                    continue
                if tok_start < end and tok_end > start:
                    token_hits.append(token_idx)

            if token_hits:
                hit_idx = torch.tensor(token_hits, device=token_hidden.device, dtype=torch.long)
                atom_embeddings.append(token_hidden[batch_idx].index_select(0, hit_idx).mean(dim=0))
            else:
                atom_embeddings.append(mol_embeddings[batch_idx])
    return torch.stack(atom_embeddings, dim=0)


class ChemBERTaDistillationStudent(nn.Module):
    """ChemBERTa encoder with an auxiliary masked-language-modeling objective."""

    def __init__(self, model_name: str = "DeepChem/ChemBERTa-77M-MLM") -> None:
        super().__init__()
        from transformers import AutoModelForMaskedLM, AutoTokenizer

        model_name_or_path = resolve_model_name_or_path(model_name)
        local_files_only = Path(model_name_or_path).exists()
        self.tokenizer = AutoTokenizer.from_pretrained(
            model_name_or_path,
            use_fast=True,
            local_files_only=local_files_only,
        )
        if not getattr(self.tokenizer, "is_fast", False):
            raise ValueError("ChemBERTaDistillationStudent requires a fast tokenizer.")
        self.model = AutoModelForMaskedLM.from_pretrained(
            model_name_or_path,
            local_files_only=local_files_only,
        )
        self.model_name = model_name_or_path

    @property
    def base_model(self) -> nn.Module:
        return self.model.base_model

    @property
    def hidden_size(self) -> int:
        return int(self.model.config.hidden_size)

    def encoder_state_dict(self) -> dict[str, torch.Tensor]:
        return self.base_model.state_dict()

    def _mask_inputs(
        self,
        input_ids: torch.Tensor,
        attention_mask: torch.Tensor,
        mlm_probability: float,
    ) -> tuple[torch.Tensor, torch.Tensor]:
        labels = input_ids.clone()
        probability = torch.full(
            labels.shape,
            float(mlm_probability),
            device=input_ids.device,
            dtype=torch.float,
        )
        special_masks = [
            self.tokenizer.get_special_tokens_mask(row, already_has_special_tokens=True)
            for row in labels.detach().cpu().tolist()
        ]
        special_mask = torch.tensor(special_masks, device=input_ids.device, dtype=torch.bool)
        probability.masked_fill_(special_mask | ~attention_mask.bool(), 0.0)
        masked = torch.bernoulli(probability).bool()

        for row_idx in range(masked.shape[0]):
            if masked[row_idx].any():
                continue
            candidates = torch.nonzero(probability[row_idx] > 0, as_tuple=False).view(-1)
            if candidates.numel() > 0:
                masked[row_idx, candidates[0]] = True

        labels[~masked] = -100
        corrupted = input_ids.clone()
        replace_with_mask = torch.bernoulli(
            torch.full(labels.shape, 0.8, device=input_ids.device)
        ).bool() & masked
        corrupted[replace_with_mask] = int(self.tokenizer.mask_token_id)

        replace_random = torch.bernoulli(
            torch.full(labels.shape, 0.5, device=input_ids.device)
        ).bool() & masked & ~replace_with_mask
        random_tokens = torch.randint(
            len(self.tokenizer),
            labels.shape,
            dtype=torch.long,
            device=input_ids.device,
        )
        corrupted[replace_random] = random_tokens[replace_random]
        return corrupted, labels

    def forward(
        self,
        smiles: Sequence[str],
        atom_counts: torch.Tensor,
        device: torch.device | str | None = None,
        mlm_probability: float = 0.15,
    ) -> dict[str, torch.Tensor]:
        if device is None:
            device = next(self.parameters()).device
        device = torch.device(device)
        tokens = self.tokenizer(
            list(smiles),
            padding=True,
            truncation=True,
            return_tensors="pt",
            return_offsets_mapping=True,
        )
        offsets = tokens.pop("offset_mapping").to(device)
        model_inputs = {key: value.to(device) for key, value in tokens.items()}

        encoder_output = self.base_model(**model_inputs)
        token_hidden = encoder_output.last_hidden_state
        mol_embeddings = mean_pool_tokens(
            token_hidden,
            model_inputs["attention_mask"],
            offsets,
        )
        atom_embeddings = pool_tokens_to_atoms(
            token_hidden=token_hidden,
            offsets=offsets,
            attention_mask=model_inputs["attention_mask"],
            smiles=smiles,
            atom_counts=atom_counts,
            mol_embeddings=mol_embeddings,
        )

        corrupted_ids, mlm_labels = self._mask_inputs(
            input_ids=model_inputs["input_ids"],
            attention_mask=model_inputs["attention_mask"],
            mlm_probability=mlm_probability,
        )
        mlm_output = self.model(
            input_ids=corrupted_ids,
            attention_mask=model_inputs["attention_mask"],
        )
        mlm_loss = F.cross_entropy(
            mlm_output.logits.view(-1, mlm_output.logits.shape[-1]),
            mlm_labels.view(-1),
            ignore_index=-100,
        )
        return {
            "atom_embeddings": atom_embeddings,
            "mol_embeddings": mol_embeddings,
            "mlm_loss": mlm_loss,
        }


def resolve_model_name_or_path(model_name: str) -> str:
    path = Path(model_name)
    looks_local = path.is_absolute() or "/" in model_name or "\\" in model_name or model_name.startswith(".")
    if not looks_local:
        return model_name

    resolved = path.expanduser().resolve()
    if resolved.exists() and (resolved / "config.json").exists():
        return str(resolved)

    discovered = discover_local_chemberta_path()
    if discovered is not None:
        print(f"Configured ChemBERTa path is invalid: {resolved}")
        print(f"Using discovered local ChemBERTa path: {discovered}")
        return str(discovered)

    if not resolved.exists():
        raise FileNotFoundError(
            f"ChemBERTa local path does not exist: {resolved}\n"
            "Check configs/pretrain_zinc.yaml teacher.model_name, or run:\n"
            "  find . -maxdepth 8 -type f -name config.json | grep -i chemberta"
        )

    raise FileNotFoundError(
        f"ChemBERTa local path exists but config.json is missing: {resolved}\n"
        "teacher.model_name must point to the exact folder containing config.json, tokenizer.json, "
        "and pytorch_model.bin. Locate it with:\n"
        "  find . -maxdepth 8 -type f -name config.json | grep -i chemberta"
    )


def discover_local_chemberta_path() -> Path | None:
    cwd = Path.cwd()
    search_roots = [cwd / "models", cwd / "comld" / "models", cwd]
    seen: set[Path] = set()

    for root in search_roots:
        if not root.exists() or root in seen:
            continue
        seen.add(root)
        for config_file in root.rglob("config.json"):
            model_dir = config_file.parent
            lower_path = str(model_dir).lower()
            if "chemberta" not in lower_path:
                continue
            has_tokenizer = (model_dir / "tokenizer.json").exists() or (
                (model_dir / "vocab.json").exists() and (model_dir / "merges.txt").exists()
            )
            has_weights = (model_dir / "pytorch_model.bin").exists() or any(
                model_dir.glob("model*.safetensors")
            )
            if has_tokenizer and has_weights:
                return model_dir.resolve()
    return None
```
