## Supplementary material for "PG-MLD: Physics-Guided Molecular Representation Learning via Dynamic 3D Trajectory Distillation": Code and Data: chemblm_distill.py

```
from __future__ import annotations

from typing import Sequence

import torch
from torch import nn

from comld.models.molformer_distill import (
    MoLFormerDistillationStudent,
    MoLFormerEncoder,
)


class ChemBLMEncoder(MoLFormerEncoder):
    """Generic ChemBLM/HuggingFace molecular language encoder.

    ChemBLM-like models are treated as SMILES language encoders.  The wrapper
    intentionally reuses the robust token/atom pooling code used by the
    MoLFormer branch, but keeps a separate class name and checkpoint namespace
    so ChemBLM experiments do not overwrite the ChemBERTa or MoLFormer lines.
    """


class ChemBLMMoleculeClassifier(nn.Module):
    def __init__(
        self,
        encoder: ChemBLMEncoder,
        output_dim: int,
        head_hidden_dim: int = 256,
        dropout: float = 0.1,
    ) -> None:
        super().__init__()
        self.encoder = encoder
        self.head = nn.Sequential(
            nn.Linear(encoder.hidden_size, head_hidden_dim),
            nn.SiLU(),
            nn.Dropout(dropout),
            nn.Linear(head_hidden_dim, int(output_dim)),
        )

    def forward(self, smiles: Sequence[str], device: torch.device | str | None = None) -> torch.Tensor:
        return self.head(self.encoder(smiles, device=device))


class ChemBLMDistillationStudent(MoLFormerDistillationStudent):
    def __init__(
        self,
        model_name: str,
        teacher_dim: int,
        trust_remote_code: bool = True,
        use_projection: bool = True,
        use_safetensors: bool = True,
    ) -> None:
        nn.Module.__init__(self)
        self.encoder = ChemBLMEncoder(
            model_name=model_name,
            trust_remote_code=trust_remote_code,
            use_safetensors=use_safetensors,
        )
        self.model_name = self.encoder.model_name
        self.teacher_dim = int(teacher_dim)
        self.use_projection = bool(use_projection)
        if self.use_projection or self.encoder.hidden_size != self.teacher_dim:
            self.atom_projector = nn.Linear(self.encoder.hidden_size, self.teacher_dim)
            self.mol_projector = nn.Linear(self.encoder.hidden_size, self.teacher_dim)
        else:
            self.atom_projector = nn.Identity()
            self.mol_projector = nn.Identity()
```
