## Supplementary material for "PG-MLD: Physics-Guided Molecular Representation Learning via Dynamic 3D Trajectory Distillation": Code and Data: dynamic_se3_liquid.py

```
from __future__ import annotations

from typing import Sequence

import torch
from torch import nn
import torch.nn.functional as F

from comld.models.se3_transformer import (
    SE3TransformerDistillationStudent,
    segment_softmax,
)


class LiquidTimeConstantCell(nn.Module):
    """Closed-form liquid time-constant update under piecewise-constant input."""

    def __init__(self, input_dim: int, hidden_dim: int) -> None:
        super().__init__()
        self.input_proj = nn.Linear(input_dim, hidden_dim)
        self.target_net = nn.Sequential(
            nn.Linear(hidden_dim * 2, hidden_dim),
            nn.SiLU(),
            nn.Linear(hidden_dim, hidden_dim),
            nn.Tanh(),
        )
        self.tau_net = nn.Sequential(
            nn.Linear(hidden_dim * 2, hidden_dim),
            nn.SiLU(),
            nn.Linear(hidden_dim, hidden_dim),
        )

    def forward(
        self,
        inputs: torch.Tensor,
        state: torch.Tensor,
        delta_time: torch.Tensor,
    ) -> torch.Tensor:
        projected = self.input_proj(inputs)
        context = torch.cat([projected, state], dim=-1)
        target = self.target_net(context)
        tau = F.softplus(self.tau_net(context)) + 1e-3
        while delta_time.ndim < state.ndim:
            delta_time = delta_time.unsqueeze(-1)
        decay = torch.exp(-delta_time.clamp_min(1e-6) / tau)
        return target + (state - target) * decay


class LiquidTrajectoryEncoder(nn.Module):
    def __init__(self, input_dim: int, hidden_dim: int) -> None:
        super().__init__()
        self.hidden_dim = int(hidden_dim)
        self.cell = LiquidTimeConstantCell(input_dim=input_dim, hidden_dim=hidden_dim)
        self.output_norm = nn.LayerNorm(hidden_dim)

    def forward(
        self,
        sequence: torch.Tensor,
        delta_time: float | torch.Tensor,
    ) -> tuple[torch.Tensor, torch.Tensor]:
        if sequence.ndim != 3:
            raise ValueError(f"Liquid input must be [entities,T,D], got {tuple(sequence.shape)}")
        num_entities, num_steps, _ = sequence.shape
        state = sequence.new_zeros((num_entities, self.hidden_dim))
        states = []
        dt = torch.as_tensor(delta_time, device=sequence.device, dtype=sequence.dtype)
        for time_idx in range(num_steps):
            state = self.cell(sequence[:, time_idx], state, dt)
            states.append(self.output_norm(state))
        state_sequence = torch.stack(states, dim=1)
        return state_sequence[:, -1], state_sequence


class DynamicSE3LiquidTeacher(nn.Module):
    def __init__(
        self,
        node_feat_dim: int,
        edge_feat_dim: int,
        output_dim: int = 384,
        hidden_dim: int = 384,
        liquid_hidden_dim: int = 384,
        num_layers: int = 4,
        message_hidden_dim: int = 256,
        num_heads: int = 8,
        rbf_dim: int = 16,
        radial_cutoff: float = 10.0,
        coord_update_scale: float = 0.1,
        dropout: float = 0.1,
        electronic_dim: int = 2,
    ) -> None:
        super().__init__()
        self.frame_encoder = SE3TransformerDistillationStudent(
            node_feat_dim=node_feat_dim,
            edge_feat_dim=edge_feat_dim,
            teacher_dim=output_dim,
            hidden_dim=hidden_dim,
            num_layers=num_layers,
            message_hidden_dim=message_hidden_dim,
            num_heads=num_heads,
            rbf_dim=rbf_dim,
            radial_cutoff=radial_cutoff,
            coord_update_scale=coord_update_scale,
            dropout=dropout,
        )
        self.liquid = LiquidTrajectoryEncoder(
            input_dim=hidden_dim,
            hidden_dim=liquid_hidden_dim,
        )
        self.atom_projector = (
            nn.Identity()
            if int(liquid_hidden_dim) == int(output_dim)
            else nn.Linear(liquid_hidden_dim, output_dim)
        )
        self.pool_score = nn.Linear(liquid_hidden_dim, 1)
        self.mol_projector = (
            nn.Identity()
            if int(liquid_hidden_dim) == int(output_dim)
            else nn.Linear(liquid_hidden_dim, output_dim)
        )
        self.charge_head = nn.Sequential(
            nn.Linear(liquid_hidden_dim, liquid_hidden_dim),
            nn.SiLU(),
            nn.Linear(liquid_hidden_dim, electronic_dim),
        )
        self.temporal_predictor = nn.Sequential(
            nn.Linear(hidden_dim, hidden_dim),
            nn.SiLU(),
            nn.Linear(hidden_dim, hidden_dim),
        )
        self.frame_projector = nn.Linear(hidden_dim, output_dim)

    def forward(
        self,
        node_feats: torch.Tensor,
        coords: torch.Tensor,
        edge_index: torch.Tensor,
        edge_attr: torch.Tensor,
        frame_batch: torch.Tensor,
        atom_batch: torch.Tensor,
        trajectory_specs: Sequence[tuple[int, int]],
        delta_times_ps: torch.Tensor,
    ) -> dict[str, torch.Tensor]:
        frame_out = self.frame_encoder(
            node_feats=node_feats,
            coords=coords,
            edge_index=edge_index,
            edge_attr=edge_attr,
            batch=frame_batch,
        )
        frame_atom_hidden = frame_out["atom_hidden"]
        frame_mol_hidden = frame_out["mol_hidden"]

        dynamic_atom_hidden = []
        liquid_state_sequences = []
        node_start = 0
        for mol_idx, (num_frames, num_atoms) in enumerate(trajectory_specs):
            node_count = int(num_frames) * int(num_atoms)
            molecule_frames = frame_atom_hidden[node_start : node_start + node_count]
            molecule_frames = molecule_frames.view(int(num_frames), int(num_atoms), -1)
            atom_sequences = molecule_frames.transpose(0, 1).contiguous()
            final_state, state_sequence = self.liquid(
                atom_sequences,
                delta_time=delta_times_ps[mol_idx],
            )
            dynamic_atom_hidden.append(final_state)
            liquid_state_sequences.append(state_sequence)
            node_start += node_count

        dynamic_atom_hidden_out = torch.cat(dynamic_atom_hidden, dim=0)
        num_molecules = len(trajectory_specs)
        scores = self.pool_score(dynamic_atom_hidden_out)
        weights = segment_softmax(scores, atom_batch, num_molecules)
        dynamic_mol_hidden = dynamic_atom_hidden_out.new_zeros(
            (num_molecules, dynamic_atom_hidden_out.shape[-1])
        )
        dynamic_mol_hidden.index_add_(0, atom_batch, dynamic_atom_hidden_out * weights)

        return {
            "atom_embeddings": self.atom_projector(dynamic_atom_hidden_out),
            "mol_embeddings": self.mol_projector(dynamic_mol_hidden),
            "atom_hidden": dynamic_atom_hidden_out,
            "mol_hidden": dynamic_mol_hidden,
            "frame_mol_hidden": frame_mol_hidden,
            "frame_mol_embeddings": self.frame_projector(frame_mol_hidden),
            "liquid_state_sequences": liquid_state_sequences,
            "denoised_coords": frame_out["denoised_coords"],
            "charge_pred": self.charge_head(dynamic_atom_hidden_out),
        }
```
