## Supplementary material for "PG-MLD: Physics-Guided Molecular Representation Learning via Dynamic 3D Trajectory Distillation": Code and Data: molformer_distill.py

```
from __future__ import annotations

from pathlib import Path
from typing import Sequence

import torch
from torch import nn

from comld.models.chemberta_teacher import (
    find_smiles_atom_spans,
    mean_pool_tokens,
    pool_tokens_to_atoms,
    resolve_model_name_or_path,
)


def attention_mean_pool(hidden: torch.Tensor, attention_mask: torch.Tensor) -> torch.Tensor:
    weights = attention_mask.bool().unsqueeze(-1).to(hidden.dtype)
    pooled = (hidden * weights).sum(dim=1)
    denom = weights.sum(dim=1).clamp_min(1.0)
    return pooled / denom


def infer_hidden_size(config: object) -> int:
    for name in ("hidden_size", "n_embd", "d_model", "embedding_size"):
        value = getattr(config, name, None)
        if value is not None:
            return int(value)
    raise AttributeError(f"Cannot infer hidden size from config: {config}")


class MoLFormerEncoder(nn.Module):
    def __init__(
        self,
        model_name: str,
        trust_remote_code: bool = True,
        use_safetensors: bool = False,
    ) -> None:
        super().__init__()
        from transformers import AutoModel, AutoTokenizer

        model_name_or_path = resolve_model_name_or_path(model_name)
        local_files_only = Path(model_name_or_path).exists()
        self.tokenizer = AutoTokenizer.from_pretrained(
            model_name_or_path,
            use_fast=True,
            local_files_only=local_files_only,
            trust_remote_code=trust_remote_code,
        )
        self.model = AutoModel.from_pretrained(
            model_name_or_path,
            local_files_only=local_files_only,
            trust_remote_code=trust_remote_code,
            use_safetensors=use_safetensors,
        )
        self.model_name = model_name_or_path

    @property
    def hidden_size(self) -> int:
        return infer_hidden_size(self.model.config)

    def tokenize(self, smiles: Sequence[str], device: torch.device) -> tuple[dict[str, torch.Tensor], torch.Tensor | None]:
        kwargs = {
            "padding": True,
            "truncation": True,
            "return_tensors": "pt",
        }
        if getattr(self.tokenizer, "is_fast", False):
            kwargs["return_offsets_mapping"] = True
        tokens = self.tokenizer(list(smiles), **kwargs)
        offsets = tokens.pop("offset_mapping", None)
        model_inputs = {key: value.to(device) for key, value in tokens.items()}
        if offsets is not None:
            offsets = offsets.to(device)
        return model_inputs, offsets

    def forward_tokens(self, smiles: Sequence[str], device: torch.device) -> tuple[torch.Tensor, torch.Tensor, torch.Tensor | None]:
        model_inputs, offsets = self.tokenize(smiles, device)
        output = self.model(**model_inputs)
        token_hidden = output.last_hidden_state if hasattr(output, "last_hidden_state") else output[0]
        return token_hidden, model_inputs["attention_mask"], offsets

    def forward(self, smiles: Sequence[str], device: torch.device | str | None = None) -> torch.Tensor:
        if device is None:
            device = next(self.parameters()).device
        device = torch.device(device)
        token_hidden, attention_mask, offsets = self.forward_tokens(smiles, device)
        if offsets is not None:
            return mean_pool_tokens(token_hidden, attention_mask, offsets)
        return attention_mean_pool(token_hidden, attention_mask)


class MoLFormerMoleculeClassifier(nn.Module):
    def __init__(
        self,
        encoder: MoLFormerEncoder,
        output_dim: int,
        head_hidden_dim: int = 256,
        dropout: float = 0.1,
    ) -> None:
        super().__init__()
        self.encoder = encoder
        self.head = nn.Sequential(
            nn.Linear(encoder.hidden_size, head_hidden_dim),
            nn.SiLU(),
            nn.Dropout(dropout),
            nn.Linear(head_hidden_dim, int(output_dim)),
        )

    def forward(self, smiles: Sequence[str], device: torch.device | str | None = None) -> torch.Tensor:
        return self.head(self.encoder(smiles, device=device))


class MoLFormerDistillationStudent(nn.Module):
    def __init__(
        self,
        model_name: str,
        teacher_dim: int,
        trust_remote_code: bool = True,
        use_projection: bool = True,
    ) -> None:
        super().__init__()
        self.encoder = MoLFormerEncoder(model_name=model_name, trust_remote_code=trust_remote_code)
        self.model_name = self.encoder.model_name
        self.teacher_dim = int(teacher_dim)
        self.use_projection = bool(use_projection)
        if self.use_projection or self.encoder.hidden_size != self.teacher_dim:
            self.atom_projector = nn.Linear(self.encoder.hidden_size, self.teacher_dim)
            self.mol_projector = nn.Linear(self.encoder.hidden_size, self.teacher_dim)
        else:
            self.atom_projector = nn.Identity()
            self.mol_projector = nn.Identity()

    @property
    def hidden_size(self) -> int:
        return self.encoder.hidden_size

    @property
    def base_model(self) -> nn.Module:
        return self.encoder.model

    def encoder_state_dict(self) -> dict[str, torch.Tensor]:
        return self.encoder.model.state_dict()

    def forward(
        self,
        smiles: Sequence[str],
        atom_counts: torch.Tensor,
        device: torch.device | str | None = None,
        mlm_probability: float = 0.0,
    ) -> dict[str, torch.Tensor]:
        del mlm_probability
        if device is None:
            device = next(self.parameters()).device
        device = torch.device(device)
        token_hidden, attention_mask, offsets = self.encoder.forward_tokens(smiles, device)
        if offsets is not None:
            mol_embeddings_raw = mean_pool_tokens(token_hidden, attention_mask, offsets)
            atom_embeddings_raw = pool_tokens_to_atoms(
                token_hidden=token_hidden,
                offsets=offsets,
                attention_mask=attention_mask,
                smiles=smiles,
                atom_counts=atom_counts,
                mol_embeddings=mol_embeddings_raw,
            )
        else:
            mol_embeddings_raw = attention_mean_pool(token_hidden, attention_mask)
            atom_embeddings_raw = []
            for batch_idx, smi in enumerate(smiles):
                n_atoms = int(atom_counts[batch_idx].item())
                spans = find_smiles_atom_spans(smi)
                for atom_idx in range(n_atoms):
                    if atom_idx < len(spans):
                        atom_embeddings_raw.append(mol_embeddings_raw[batch_idx])
                    else:
                        atom_embeddings_raw.append(mol_embeddings_raw[batch_idx])
            atom_embeddings_raw = torch.stack(atom_embeddings_raw, dim=0)

        atom_embeddings = self.atom_projector(atom_embeddings_raw)
        mol_embeddings = self.mol_projector(mol_embeddings_raw)
        return {
            "atom_embeddings": atom_embeddings,
            "mol_embeddings": mol_embeddings,
            "mlm_loss": token_hidden.new_tensor(0.0),
        }
```
