## Supplementary material for "PG-MLD: Physics-Guided Molecular Representation Learning via Dynamic 3D Trajectory Distillation": Code and Data: se3_transformer.py

```
from __future__ import annotations

import math

import torch
from torch import nn


def segment_mean(values: torch.Tensor, segment_ids: torch.Tensor, num_segments: int) -> torch.Tensor:
    out = values.new_zeros((num_segments, values.shape[-1]))
    out.index_add_(0, segment_ids, values)
    counts = values.new_zeros((num_segments, 1))
    counts.index_add_(0, segment_ids, torch.ones((values.shape[0], 1), device=values.device, dtype=values.dtype))
    return out / counts.clamp_min(1.0)


def segment_softmax(scores: torch.Tensor, segment_ids: torch.Tensor, num_segments: int) -> torch.Tensor:
    if scores.numel() == 0:
        return scores
    # Reductions are kept in FP32 under autocast. Some PyTorch/CUDA versions
    # promote exp() to FP32 while retaining an FP16 accumulation buffer, which
    # makes scatter_add_ fail with a dtype mismatch.
    output_dtype = scores.dtype
    stable_scores = scores.float()
    index = segment_ids.view(-1, 1).expand_as(stable_scores)
    max_values = stable_scores.new_full(
        (num_segments, stable_scores.shape[-1]),
        -torch.inf,
    )
    max_values.scatter_reduce_(
        0,
        index,
        stable_scores,
        reduce="amax",
        include_self=True,
    )
    shifted = stable_scores - max_values.index_select(0, segment_ids)
    exp_scores = torch.exp(shifted)
    denom = stable_scores.new_zeros((num_segments, stable_scores.shape[-1]))
    denom.scatter_add_(0, index, exp_scores)
    probabilities = exp_scores / denom.index_select(0, segment_ids).clamp_min(1e-9)
    return probabilities.to(output_dtype)


class GaussianRadialBasis(nn.Module):
    def __init__(self, num_basis: int = 16, cutoff: float = 10.0) -> None:
        super().__init__()
        centers = torch.linspace(0.0, float(cutoff), int(num_basis))
        self.register_buffer("centers", centers)
        self.gamma = 1.0 / max((float(cutoff) / max(int(num_basis) - 1, 1)) ** 2, 1e-6)

    def forward(self, distances: torch.Tensor) -> torch.Tensor:
        return torch.exp(-self.gamma * (distances.unsqueeze(-1) - self.centers) ** 2)


class SE3TransformerLayer(nn.Module):
    def __init__(
        self,
        hidden_dim: int,
        edge_feat_dim: int,
        num_heads: int = 8,
        rbf_dim: int = 16,
        message_hidden_dim: int = 256,
        coord_update_scale: float = 0.1,
        dropout: float = 0.1,
    ) -> None:
        super().__init__()
        if hidden_dim % num_heads != 0:
            raise ValueError(f"hidden_dim={hidden_dim} must be divisible by num_heads={num_heads}")
        self.hidden_dim = int(hidden_dim)
        self.num_heads = int(num_heads)
        self.head_dim = self.hidden_dim // self.num_heads
        self.coord_update_scale = float(coord_update_scale)

        edge_input_dim = int(edge_feat_dim) + int(rbf_dim)
        self.edge_encoder = nn.Sequential(
            nn.Linear(edge_input_dim, message_hidden_dim),
            nn.SiLU(),
            nn.Dropout(dropout),
            nn.Linear(message_hidden_dim, hidden_dim),
            nn.SiLU(),
        )
        self.q_proj = nn.Linear(hidden_dim, hidden_dim)
        self.k_proj = nn.Linear(hidden_dim, hidden_dim)
        self.v_proj = nn.Linear(hidden_dim, hidden_dim)
        self.edge_value_proj = nn.Linear(hidden_dim, hidden_dim)
        self.edge_bias_proj = nn.Linear(hidden_dim, num_heads)
        self.out_proj = nn.Sequential(
            nn.Linear(hidden_dim, hidden_dim),
            nn.Dropout(dropout),
        )
        self.coord_mlp = nn.Sequential(
            nn.Linear(hidden_dim, message_hidden_dim),
            nn.SiLU(),
            nn.Linear(message_hidden_dim, 1),
        )
        self.ffn = nn.Sequential(
            nn.Linear(hidden_dim, message_hidden_dim),
            nn.SiLU(),
            nn.Dropout(dropout),
            nn.Linear(message_hidden_dim, hidden_dim),
            nn.Dropout(dropout),
        )
        self.attn_norm = nn.LayerNorm(hidden_dim)
        self.ffn_norm = nn.LayerNorm(hidden_dim)

    def forward(
        self,
        h: torch.Tensor,
        coords: torch.Tensor,
        edge_index: torch.Tensor,
        edge_attr: torch.Tensor,
        rbf: torch.Tensor,
    ) -> tuple[torch.Tensor, torch.Tensor]:
        if edge_index.numel() == 0:
            h = self.attn_norm(h)
            h = self.ffn_norm(h + self.ffn(h))
            return h, coords

        src, dst = edge_index[0], edge_index[1]
        num_nodes = h.shape[0]
        edge_context = self.edge_encoder(torch.cat([edge_attr, rbf], dim=-1))

        q = self.q_proj(h).view(num_nodes, self.num_heads, self.head_dim)
        k = self.k_proj(h).view(num_nodes, self.num_heads, self.head_dim)
        v = self.v_proj(h).view(num_nodes, self.num_heads, self.head_dim)
        edge_value = self.edge_value_proj(edge_context).view(-1, self.num_heads, self.head_dim)

        scores = (q[dst] * k[src]).sum(dim=-1) / math.sqrt(self.head_dim)
        scores = scores + self.edge_bias_proj(edge_context)
        attn = segment_softmax(scores, dst, num_nodes)

        messages = attn.unsqueeze(-1) * (v[src] + edge_value)
        messages = messages.reshape(messages.shape[0], self.hidden_dim)
        aggregated = h.new_zeros(h.shape)
        aggregated.index_add_(0, dst, messages)
        h = self.attn_norm(h + self.out_proj(aggregated))
        h = self.ffn_norm(h + self.ffn(h))

        coord_diff = coords[src] - coords[dst]
        coord_gate = torch.tanh(self.coord_mlp(edge_context)) * self.coord_update_scale
        coord_weight = attn.mean(dim=-1, keepdim=True) * coord_gate
        coord_updates = coord_diff * coord_weight
        coord_agg = coords.new_zeros(coords.shape)
        coord_agg.index_add_(0, dst, coord_updates)
        coords = coords + coord_agg
        return h, coords


class SE3TransformerDistillationStudent(nn.Module):
    def __init__(
        self,
        node_feat_dim: int,
        edge_feat_dim: int,
        teacher_dim: int,
        hidden_dim: int = 384,
        num_layers: int = 4,
        message_hidden_dim: int = 256,
        num_heads: int = 8,
        rbf_dim: int = 16,
        radial_cutoff: float = 10.0,
        coord_update_scale: float = 0.1,
        dropout: float = 0.1,
    ) -> None:
        super().__init__()
        self.node_encoder = nn.Sequential(
            nn.Linear(node_feat_dim, hidden_dim),
            nn.SiLU(),
            nn.LayerNorm(hidden_dim),
        )
        self.radial_basis = GaussianRadialBasis(num_basis=rbf_dim, cutoff=radial_cutoff)
        self.layers = nn.ModuleList(
            [
                SE3TransformerLayer(
                    hidden_dim=hidden_dim,
                    edge_feat_dim=edge_feat_dim,
                    num_heads=num_heads,
                    rbf_dim=rbf_dim,
                    message_hidden_dim=message_hidden_dim,
                    coord_update_scale=coord_update_scale,
                    dropout=dropout,
                )
                for _ in range(num_layers)
            ]
        )
        self.atom_projector = nn.Linear(hidden_dim, teacher_dim)
        self.mol_projector = nn.Linear(hidden_dim, teacher_dim)

    def forward(
        self,
        node_feats: torch.Tensor,
        coords: torch.Tensor,
        edge_index: torch.Tensor,
        edge_attr: torch.Tensor,
        batch: torch.Tensor,
    ) -> dict[str, torch.Tensor]:
        h = self.node_encoder(node_feats)
        x = coords
        for layer in self.layers:
            if edge_index.numel() > 0:
                src, dst = edge_index[0], edge_index[1]
                distances = torch.linalg.norm(x[src] - x[dst], dim=-1)
                rbf = self.radial_basis(distances)
            else:
                rbf = edge_attr.new_empty((0, self.radial_basis.centers.numel()))
            h, x = layer(h, x, edge_index, edge_attr, rbf)

        num_graphs = int(batch.max().item()) + 1 if batch.numel() else 0
        mol_hidden = segment_mean(h, batch, num_graphs)
        return {
            "atom_embeddings": self.atom_projector(h),
            "mol_embeddings": self.mol_projector(mol_hidden),
            "denoised_coords": x,
            "atom_hidden": h,
            "mol_hidden": mol_hidden,
        }
```
