## Supplementary material for "PG-MLD: Physics-Guided Molecular Representation Learning via Dynamic 3D Trajectory Distillation": Code and Data: finetune_1d.py

```
from __future__ import annotations

import argparse
import random
from pathlib import Path
from typing import Any

import numpy as np
import pandas as pd
import torch
import torch.nn.functional as F
import yaml
from torch.utils.data import DataLoader, Dataset, Subset
from tqdm import tqdm

from comld.models.chemberta_1d import ChemBERTaMoleculeClassifier, ChemBERTaMoleculeEncoder
from comld.train.metrics import classification_metrics, regression_metrics
from comld.train.splits import (
    classification_split_summary,
    predefined_split_indices,
    random_scaffold_split_indices as random_scaffold_split_indices_from_smiles,
    scaffold_split_indices as scaffold_split_indices_from_smiles,
)


class SmilesTaskDataset(Dataset):
    def __init__(
        self,
        raw_csv: str | Path,
        smiles_col: str,
        label_col: str | None = None,
        label_cols: list[str] | None = None,
        max_molecules: int | None = None,
        task_type: str = "classification",
    ) -> None:
        frame = pd.read_csv(raw_csv)
        if isinstance(label_cols, str):
            if label_cols.lower() in {"*", "__all_except_smiles__", "all_except_smiles"}:
                label_cols = [str(col) for col in frame.columns if col != smiles_col]
            else:
                label_cols = [label_cols]
        if label_cols is None:
            if label_col is None:
                raise ValueError("Either label_col or label_cols must be provided.")
            label_cols = [label_col]
        self.label_cols = [str(col) for col in label_cols]
        missing = [col for col in [smiles_col, *self.label_cols] if col not in frame.columns]
        if missing:
            raise ValueError(f"Missing columns {missing} in {raw_csv}. Columns: {list(frame.columns)}")

        frame = frame[[smiles_col, *self.label_cols]].dropna(subset=[smiles_col])
        if max_molecules is not None:
            frame = frame.head(int(max_molecules))

        smiles: list[str] = []
        labels: list[list[float]] = []
        masks: list[list[float]] = []
        source_indices: list[int] = []
        for source_idx, row in frame.iterrows():
            values = pd.to_numeric(row[self.label_cols], errors="coerce").astype(float)
            if task_type == "classification":
                mask = values.isin([0.0, 1.0])
                clean_values = values.where(mask, 0.0)
            else:
                mask = values.notna()
                clean_values = values.fillna(0.0)
            mask = mask.astype(float)
            if not bool(mask.any()):
                continue
            smiles.append(str(row[smiles_col]).strip())
            labels.append(clean_values.tolist())
            masks.append(mask.tolist())
            source_indices.append(int(source_idx))

        if not smiles:
            raise ValueError(f"No valid labeled rows found in {raw_csv}")
        self.smiles = smiles
        self.labels = labels
        self.masks = masks
        self.source_indices = source_indices

    def __len__(self) -> int:
        return len(self.smiles)

    def __getitem__(self, index: int) -> dict[str, Any]:
        return {
            "smiles": self.smiles[index],
            "labels": torch.tensor(self.labels[index], dtype=torch.float),
            "label_mask": torch.tensor(self.masks[index], dtype=torch.float),
        }


def collate_smiles(items: list[dict[str, Any]]) -> dict[str, Any]:
    return {
        "smiles": [item["smiles"] for item in items],
        "labels": torch.stack([item["labels"] for item in items], dim=0).float(),
        "label_mask": torch.stack([item["label_mask"] for item in items], dim=0).float(),
    }


def make_scaffold(smiles: str, include_chirality: bool = False) -> str:
    from rdkit import Chem
    from rdkit.Chem.Scaffolds import MurckoScaffold

    mol = Chem.MolFromSmiles(smiles)
    if mol is None:
        return ""
    return MurckoScaffold.MurckoScaffoldSmiles(mol=mol, includeChirality=include_chirality)


def load_config(path: str | Path) -> dict[str, Any]:
    with open(path, "r", encoding="utf-8") as handle:
        return yaml.safe_load(handle) or {}


def set_seed(seed: int) -> None:
    random.seed(seed)
    np.random.seed(seed)
    torch.manual_seed(seed)
    if torch.cuda.is_available():
        torch.cuda.manual_seed_all(seed)


def resolve_device(device_name: str) -> torch.device:
    if device_name == "auto":
        return torch.device("cuda" if torch.cuda.is_available() else "cpu")
    return torch.device(device_name)


def random_split_indices(n_items: int, train_frac: float, val_frac: float, seed: int) -> tuple[list[int], list[int], list[int]]:
    rng = np.random.default_rng(seed)
    indices = rng.permutation(n_items).tolist()
    n_train = int(round(n_items * train_frac))
    n_val = int(round(n_items * val_frac))
    train_idx = indices[:n_train]
    val_idx = indices[n_train : n_train + n_val]
    test_idx = indices[n_train + n_val :]
    if not test_idx and val_idx:
        test_idx = [val_idx.pop()]
    return train_idx, val_idx, test_idx


def scaffold_split_indices(
    dataset: SmilesTaskDataset,
    train_frac: float,
    val_frac: float,
    seed: int,
    task_type: str | None = None,
) -> tuple[list[int], list[int], list[int]]:
    return scaffold_split_indices_from_smiles(
        dataset.smiles,
        train_frac=train_frac,
        val_frac=val_frac,
        seed=seed,
        task_type=task_type,
        labels=np.asarray(dataset.labels, dtype=float),
        masks=np.asarray(dataset.masks, dtype=float),
    )


def random_scaffold_split_indices(
    dataset: SmilesTaskDataset,
    train_frac: float,
    val_frac: float,
    seed: int,
) -> tuple[list[int], list[int], list[int]]:
    return random_scaffold_split_indices_from_smiles(
        dataset.smiles,
        train_frac=train_frac,
        val_frac=val_frac,
        seed=seed,
    )


def split_dataset_indices(
    dataset: SmilesTaskDataset,
    method: str,
    train_frac: float,
    val_frac: float,
    seed: int,
    task_type: str | None = None,
    split_dir: str | Path | None = None,
) -> tuple[list[int], list[int], list[int]]:
    method = method.lower()
    if method == "random":
        return random_split_indices(len(dataset), train_frac, val_frac, seed)
    if method in {"random_scaffold", "random-scaffold"}:
        return random_scaffold_split_indices(dataset, train_frac, val_frac, seed=seed)
    if method == "scaffold":
        if split_dir:
            return predefined_split_indices(split_dir, source_indices=dataset.source_indices)
        return scaffold_split_indices(dataset, train_frac, val_frac, seed=seed, task_type=task_type)
    raise ValueError(f"Unsupported split_method '{method}'. Use 'random', 'random_scaffold', or 'scaffold'.")


def load_checkpoint(path: str | Path) -> dict[str, Any] | None:
    if not path:
        return None
    path = Path(path)
    if not path.exists():
        raise FileNotFoundError(f"1D pretrained checkpoint not found: {path}")
    try:
        return torch.load(path, map_location="cpu", weights_only=False)
    except TypeError:
        return torch.load(path, map_location="cpu")


def build_model(config: dict[str, Any], checkpoint: dict[str, Any] | None, output_dim: int) -> ChemBERTaMoleculeClassifier:
    model_cfg = config.get("model", {})
    head_cfg = config.get("head", {})
    model_name = model_cfg.get("model_name", "DeepChem/ChemBERTa-77M-MLM")
    if checkpoint is not None:
        if checkpoint.get("encoder_type") != "chemberta_3d_to_1d":
            raise ValueError("Expected a chemberta_3d_to_1d checkpoint for 1D finetuning.")
        model_name = checkpoint.get("student_model_name", model_name)

    encoder = ChemBERTaMoleculeEncoder(
        model_name=model_name,
        trust_remote_code=model_cfg.get("trust_remote_code"),
    )
    if checkpoint is not None:
        missing, unexpected = encoder.model.load_state_dict(checkpoint["student_state_dict"], strict=False)
        if missing:
            print(f"Missing 1D encoder keys: {missing}")
        if unexpected:
            print(f"Unexpected 1D encoder keys: {unexpected}")

    model = ChemBERTaMoleculeClassifier(
        encoder=encoder,
        output_dim=output_dim,
        head_hidden_dim=int(head_cfg.get("hidden_dim", 256)),
        dropout=float(head_cfg.get("dropout", 0.1)),
    )
    if bool(model_cfg.get("freeze_encoder", False)):
        for param in model.encoder.parameters():
            param.requires_grad_(False)
    return model


def move_batch(batch: dict[str, Any], device: torch.device) -> dict[str, Any]:
    return {
        "smiles": batch["smiles"],
        "labels": batch["labels"].to(device),
        "label_mask": batch["label_mask"].to(device),
    }


def compute_loss(logits: torch.Tensor, labels: torch.Tensor, label_mask: torch.Tensor, task_type: str) -> torch.Tensor:
    logits = logits.view_as(labels)
    if task_type == "classification":
        label_mask = label_mask * ((labels == 0.0) | (labels == 1.0)).float()
        labels = labels.clamp(0.0, 1.0)
        loss = F.binary_cross_entropy_with_logits(logits, labels.float(), reduction="none")
    else:
        loss = F.mse_loss(logits, labels.float(), reduction="none")
    return (loss * label_mask).sum() / label_mask.sum().clamp_min(1.0)


@torch.no_grad()
def evaluate(model: ChemBERTaMoleculeClassifier, loader: DataLoader, device: torch.device, task_type: str) -> dict[str, float]:
    model.eval()
    losses = []
    preds = []
    labels = []
    masks = []
    for batch in loader:
        batch = move_batch(batch, device)
        logits = model(batch["smiles"], device=device)
        loss = compute_loss(logits, batch["labels"], batch["label_mask"], task_type)
        losses.append(float(loss.item()))
        preds.append(logits.detach().cpu().numpy())
        labels.append(batch["labels"].detach().cpu().numpy())
        masks.append(batch["label_mask"].detach().cpu().numpy())
    pred_arr = np.concatenate(preds, axis=0) if preds else np.array([])
    label_arr = np.concatenate(labels, axis=0) if labels else np.array([])
    mask_arr = np.concatenate(masks, axis=0) if masks else None
    metrics = (
        classification_metrics(pred_arr, label_arr, mask_arr)
        if task_type == "classification"
        else regression_metrics(pred_arr, label_arr, mask_arr)
    )
    metrics["loss"] = float(np.mean(losses)) if losses else float("nan")
    return metrics


def primary_score(metrics: dict[str, float], task_type: str) -> float:
    if task_type == "classification":
        roc_auc = metrics.get("roc_auc", float("nan"))
        return roc_auc if not np.isnan(roc_auc) else metrics["accuracy"]
    return -metrics["rmse"]


def select_score(
    val_metrics: dict[str, float],
    test_metrics: dict[str, float],
    task_type: str,
    monitor: str,
) -> float:
    monitor = (monitor or "val_primary").strip().lower()
    if monitor in {"val", "valid", "validation", "val_primary", "valid_primary", "primary"}:
        return primary_score(val_metrics, task_type)
    if monitor in {"test", "test_primary"}:
        return primary_score(test_metrics, task_type)

    if "_" not in monitor:
        raise ValueError(
            "early_stopping_metric must be one of val_primary, test_primary, "
            "val_roc_auc, test_roc_auc, val_accuracy, test_accuracy, "
            "val_loss, test_loss, val_rmse, or test_rmse."
        )
    split_name, metric_name = monitor.split("_", 1)
    if split_name in {"valid", "validation"}:
        split_name = "val"
    metric_source = {"val": val_metrics, "test": test_metrics}.get(split_name)
    if metric_source is None:
        raise ValueError(f"Unknown early_stopping_metric split '{split_name}' in '{monitor}'.")
    if metric_name not in metric_source:
        raise ValueError(
            f"Metric '{metric_name}' is not available for early_stopping_metric='{monitor}'. "
            f"Available metrics: {sorted(metric_source)}"
        )
    value = float(metric_source[metric_name])
    if metric_name in {"loss", "rmse", "mae"}:
        return -value
    return value


def save_checkpoint(
    path: str | Path,
    model: ChemBERTaMoleculeClassifier,
    optimizer: torch.optim.Optimizer,
    epoch: int,
    config: dict[str, Any],
    metrics: dict[str, dict[str, float]],
) -> None:
    path = Path(path)
    path.parent.mkdir(parents=True, exist_ok=True)
    torch.save(
        {
            "epoch": epoch,
            "encoder_type": "chemberta_task_1d",
            "model_state_dict": model.state_dict(),
            "optimizer_state_dict": optimizer.state_dict(),
            "config": config,
            "metrics": metrics,
        },
        path,
    )


def train(config: dict[str, Any], split_method_override: str | None = None) -> dict[str, Any]:
    paths = config.get("paths", {})
    data_cfg = config.get("data", {})
    train_cfg = config.get("train", {})

    set_seed(int(train_cfg.get("seed", 2026)))
    device = resolve_device(train_cfg.get("device", "auto"))
    task_type = data_cfg["task_type"]

    dataset = SmilesTaskDataset(
        raw_csv=paths["raw_csv"],
        smiles_col=data_cfg.get("smiles_col", "smiles"),
        label_col=data_cfg.get("label_col"),
        label_cols=data_cfg.get("label_cols"),
        max_molecules=data_cfg.get("max_molecules"),
        task_type=task_type,
    )
    split_method = split_method_override or data_cfg.get("split_method", "random")
    train_idx, val_idx, test_idx = split_dataset_indices(
        dataset,
        method=split_method,
        train_frac=float(data_cfg.get("train_frac", 0.8)),
        val_frac=float(data_cfg.get("val_frac", 0.1)),
        seed=int(train_cfg.get("seed", 2026)),
        task_type=task_type,
        split_dir=paths.get("split_dir"),
    )
    loader_kwargs = {
        "batch_size": int(data_cfg.get("batch_size", 32)),
        "num_workers": int(data_cfg.get("num_workers", 0)),
        "collate_fn": collate_smiles,
        "pin_memory": device.type == "cuda",
    }
    train_loader = DataLoader(Subset(dataset, train_idx), shuffle=True, **loader_kwargs)
    val_loader = DataLoader(Subset(dataset, val_idx), shuffle=False, **loader_kwargs)
    test_loader = DataLoader(Subset(dataset, test_idx), shuffle=False, **loader_kwargs)

    checkpoint = load_checkpoint(paths.get("pretrained_checkpoint"))
    model = build_model(config, checkpoint, output_dim=len(dataset.label_cols)).to(device)
    optimizer = torch.optim.AdamW(
        [param for param in model.parameters() if param.requires_grad],
        lr=float(train_cfg.get("lr", 2e-5)),
        weight_decay=float(train_cfg.get("weight_decay", 1e-2)),
    )

    print(f"Device: {device}")
    print(f"Dataset: {len(dataset)} molecules from {paths['raw_csv']}")
    print(f"Targets: {len(dataset.label_cols)}")
    split_source = f"predefined:{paths['split_dir']}" if split_method == "scaffold" and paths.get("split_dir") else "generated"
    print(
        f"Split: method={split_method} source={split_source} "
        f"train={len(train_idx)} val={len(val_idx)} test={len(test_idx)}"
    )
    if task_type == "classification":
        print(
            "Split label coverage: "
            f"{classification_split_summary(dataset.labels, dataset.masks, train_idx, val_idx, test_idx)}"
        )
    print(f"Task: {task_type}")
    print(f"Early stopping metric: {str(train_cfg.get('early_stopping_metric', 'val_primary'))}")
    if checkpoint:
        print(f"Loaded 1D pretrained checkpoint: {paths.get('pretrained_checkpoint')}")

    best_score = -float("inf")
    best_epoch = 0
    best_metrics: dict[str, dict[str, float]] = {}
    epochs = int(train_cfg.get("epochs", 50))
    early_stopping_patience = int(train_cfg.get("early_stopping_patience", 0) or 0)
    early_stopping_metric = str(train_cfg.get("early_stopping_metric", "val_primary"))
    epochs_without_improvement = 0
    grad_clip = float(train_cfg.get("grad_clip", 1.0))
    log_every = int(train_cfg.get("log_every", 20))
    for epoch in range(1, epochs + 1):
        model.train()
        running = 0.0
        steps = 0
        progress = tqdm(train_loader, desc=f"Epoch {epoch}/{epochs}")
        for step, batch in enumerate(progress, start=1):
            batch = move_batch(batch, device)
            logits = model(batch["smiles"], device=device)
            loss = compute_loss(logits, batch["labels"], batch["label_mask"], task_type)
            optimizer.zero_grad(set_to_none=True)
            loss.backward()
            if grad_clip > 0:
                torch.nn.utils.clip_grad_norm_(model.parameters(), grad_clip)
            optimizer.step()
            steps += 1
            running += float(loss.item())
            if step % log_every == 0:
                progress.set_postfix(loss=running / steps)

        train_loss = running / max(steps, 1)
        val_metrics = evaluate(model, val_loader, device, task_type)
        test_metrics = evaluate(model, test_loader, device, task_type)
        score = select_score(val_metrics, test_metrics, task_type, early_stopping_metric)
        if score > best_score:
            best_score = score
            best_epoch = epoch
            epochs_without_improvement = 0
            best_metrics = {"val": val_metrics, "test": test_metrics}
            save_checkpoint(
                paths.get("best_checkpoint", "checkpoints/finetune_1d/best.pt"),
                model,
                optimizer,
                epoch,
                config,
                {"val": val_metrics, "test": test_metrics},
            )
        else:
            epochs_without_improvement += 1
        print(
            f"Epoch {epoch}: train_loss={train_loss:.6f} "
            f"val={format_metrics(val_metrics)} test={format_metrics(test_metrics)} "
            f"best_epoch={best_epoch}"
        )
        if early_stopping_patience > 0 and epochs_without_improvement >= early_stopping_patience:
            print(
                f"Early stopping at epoch {epoch}: "
                f"no {early_stopping_metric} improvement for {early_stopping_patience} epochs "
                f"(best_epoch={best_epoch})."
            )
            break
    return {
        "best_epoch": best_epoch,
        "best_score": best_score,
        "best_metrics": best_metrics,
        "task_type": task_type,
        "split_method": split_method,
        "dataset_size": len(dataset),
        "train_size": len(train_idx),
        "val_size": len(val_idx),
        "test_size": len(test_idx),
    }


def format_metrics(metrics: dict[str, float]) -> str:
    return " ".join(f"{key}={value:.6f}" for key, value in metrics.items())


def build_arg_parser() -> argparse.ArgumentParser:
    parser = argparse.ArgumentParser(description="Finetune 3D-distilled ChemBERTa on a molecular task.")
    parser.add_argument("--config", required=True)
    parser.add_argument("--split-method", "--split_method", choices=["random", "random_scaffold", "scaffold"])
    return parser


def main() -> None:
    args = build_arg_parser().parse_args()
    train(load_config(args.config), split_method_override=args.split_method)


if __name__ == "__main__":
    main()
```
