## Supplementary material for "PG-MLD: Physics-Guided Molecular Representation Learning via Dynamic 3D Trajectory Distillation": Code and Data: finetune_bace_1d.py

```
from __future__ import annotations

import argparse

from comld.train.finetune_1d import load_config, train


def build_arg_parser() -> argparse.ArgumentParser:
    parser = argparse.ArgumentParser(description="Finetune 3D-distilled ChemBERTa on BACE.")
    parser.add_argument("--config", default="configs/finetune/scaffold/finetune_bace_1d.yaml")
    parser.add_argument("--split-method", "--split_method", choices=["random", "scaffold"])
    return parser


def main() -> None:
    args = build_arg_parser().parse_args()
    train(load_config(args.config), split_method_override=args.split_method)


if __name__ == "__main__":
    main()
```
