## Supplementary material for "PG-MLD: Physics-Guided Molecular Representation Learning via Dynamic 3D Trajectory Distillation": Code and Data: finetune_cliff_1d.py

```
from __future__ import annotations

import argparse
import json
import math
import random
from pathlib import Path
from typing import Any

import numpy as np
import pandas as pd
import torch
import torch.nn.functional as F
import yaml
from torch.utils.data import DataLoader, Dataset, Subset
from tqdm import tqdm

from comld.train.finetune_1d import load_checkpoint, resolve_device
from comld.train.finetune_qm9_1d import (
    atomic_json_save,
    atomic_torch_save,
    build_model,
    linear_warmup_decay,
    make_grad_scaler,
)


DEFAULT_DATASET = "CHEMBL2034_Ki"


class MoleculeACEDataset(Dataset):
    """Official MoleculeACE regression data with its fixed train/test split."""

    def __init__(
        self,
        raw_csv: str | Path,
        smiles_col: str = "smiles",
        target_col: str = "y [pEC50/pKi]",
        cliff_col: str = "cliff_mol",
        split_col: str = "split",
    ) -> None:
        frame = pd.read_csv(raw_csv)
        if target_col not in frame.columns and target_col == "y [pEC50/pKi]" and "y" in frame.columns:
            print(
                "Warning: 'y [pEC50/pKi]' is absent; using the original MoleculeACE "
                "'y' column. RMSE values are unaffected by this constant label shift."
            )
            target_col = "y"

        required = [smiles_col, target_col, cliff_col, split_col]
        missing = [column for column in required if column not in frame.columns]
        if missing:
            raise ValueError(
                f"Missing MoleculeACE columns {missing} in {raw_csv}. "
                f"Columns: {list(frame.columns)}"
            )

        labels = pd.to_numeric(frame[target_col], errors="coerce")
        splits = frame[split_col].astype(str).str.strip().str.lower()
        cliffs = parse_boolean_series(frame[cliff_col])
        valid = (
            frame[smiles_col].notna()
            & labels.notna()
            & np.isfinite(labels.to_numpy(dtype=float))
            & splits.isin(["train", "test"])
            & cliffs.notna()
        )
        removed = int((~valid).sum())
        if removed:
            print(f"Removed {removed} invalid MoleculeACE rows.")

        self.smiles = frame.loc[valid, smiles_col].astype(str).str.strip().tolist()
        self.labels = labels.loc[valid].astype(np.float32).to_numpy()
        self.cliff_flags = cliffs.loc[valid].astype(bool).to_numpy()
        self.splits = splits.loc[valid].to_numpy(dtype=str)
        self.target_col = target_col
        if not self.smiles:
            raise ValueError(f"No valid MoleculeACE rows found in {raw_csv}")
        if not np.any(self.splits == "train") or not np.any(self.splits == "test"):
            raise ValueError("The official MoleculeACE CSV must contain both train and test rows.")

    def __len__(self) -> int:
        return len(self.smiles)

    def __getitem__(self, index: int) -> dict[str, Any]:
        return {
            "smiles": self.smiles[index],
            "label": torch.tensor(self.labels[index], dtype=torch.float32),
            "cliff": torch.tensor(self.cliff_flags[index], dtype=torch.bool),
        }


def parse_boolean_series(values: pd.Series) -> pd.Series:
    if pd.api.types.is_bool_dtype(values):
        return values.astype("boolean")
    normalized = values.astype(str).str.strip().str.lower()
    mapping = {
        "true": True,
        "1": True,
        "yes": True,
        "false": False,
        "0": False,
        "no": False,
    }
    return normalized.map(mapping).astype("boolean")


def collate_cliff(items: list[dict[str, Any]]) -> dict[str, Any]:
    return {
        "smiles": [item["smiles"] for item in items],
        "labels": torch.stack([item["label"] for item in items]),
        "cliffs": torch.stack([item["cliff"] for item in items]),
    }


def load_config(path: str | Path) -> dict[str, Any]:
    with open(path, "r", encoding="utf-8") as handle:
        return yaml.safe_load(handle) or {}


def set_seed(seed: int) -> None:
    random.seed(seed)
    np.random.seed(seed)
    torch.manual_seed(seed)
    if torch.cuda.is_available():
        torch.cuda.manual_seed_all(seed)


def split_official_train_validation(
    dataset: MoleculeACEDataset,
    val_frac: float,
    seed: int,
) -> tuple[list[int], list[int], list[int]]:
    """Keep the official test set fixed and stratify validation by cliff label."""

    if not 0.0 < val_frac < 0.5:
        raise ValueError(f"validation fraction must be in (0, 0.5), got {val_frac}")
    official_train = np.flatnonzero(dataset.splits == "train")
    official_test = np.flatnonzero(dataset.splits == "test")
    rng = np.random.RandomState(seed)
    train_parts: list[np.ndarray] = []
    val_parts: list[np.ndarray] = []

    for cliff_value in (False, True):
        group = official_train[dataset.cliff_flags[official_train] == cliff_value].copy()
        rng.shuffle(group)
        if len(group) <= 1:
            train_parts.append(group)
            continue
        val_count = max(1, int(round(len(group) * val_frac)))
        val_count = min(val_count, len(group) - 1)
        val_parts.append(group[:val_count])
        train_parts.append(group[val_count:])

    train_idx = np.concatenate(train_parts)
    val_idx = np.concatenate(val_parts) if val_parts else np.empty(0, dtype=np.int64)
    rng.shuffle(train_idx)
    rng.shuffle(val_idx)
    if not len(val_idx):
        raise ValueError("Unable to create a non-empty validation split from official training rows.")
    return train_idx.tolist(), val_idx.tolist(), official_test.tolist()


def target_statistics(dataset: MoleculeACEDataset, indices: list[int]) -> tuple[float, float]:
    values = dataset.labels[np.asarray(indices, dtype=np.int64)].astype(np.float64)
    mean = float(values.mean())
    std = float(values.std())
    if not math.isfinite(mean) or not math.isfinite(std) or std < 1e-12:
        raise ValueError(f"Invalid training target statistics: mean={mean}, std={std}")
    return mean, std


def normalized_mse(
    predictions: torch.Tensor,
    labels: torch.Tensor,
    target_mean: float,
    target_std: float,
) -> torch.Tensor:
    normalized_labels = (labels.reshape(-1) - target_mean) / target_std
    return F.mse_loss(predictions.reshape(-1), normalized_labels)


def subset_error_metrics(
    true: np.ndarray,
    prediction: np.ndarray,
    mask: np.ndarray,
    prefix: str,
) -> dict[str, float | int]:
    count = int(mask.sum())
    result: dict[str, float | int] = {f"{prefix}_count": count}
    if count == 0:
        result[f"{prefix}_rmse"] = float("nan")
        result[f"{prefix}_mae"] = float("nan")
        return result
    error = prediction[mask] - true[mask]
    result[f"{prefix}_rmse"] = float(np.sqrt(np.mean(np.square(error))))
    result[f"{prefix}_mae"] = float(np.mean(np.abs(error)))
    return result


@torch.no_grad()
def evaluate(
    model: torch.nn.Module,
    loader: DataLoader,
    device: torch.device,
    target_mean: float,
    target_std: float,
) -> dict[str, float | int]:
    model.eval()
    predictions: list[np.ndarray] = []
    labels: list[np.ndarray] = []
    cliffs: list[np.ndarray] = []
    for batch in loader:
        pred_norm = model(batch["smiles"], device=device).reshape(-1)
        pred = pred_norm * target_std + target_mean
        predictions.append(pred.detach().cpu().numpy())
        labels.append(batch["labels"].numpy())
        cliffs.append(batch["cliffs"].numpy().astype(bool))

    if not predictions:
        raise ValueError("Cannot evaluate an empty data loader.")
    prediction = np.concatenate(predictions)
    true = np.concatenate(labels)
    cliff_mask = np.concatenate(cliffs)
    metrics: dict[str, float | int] = {}
    metrics.update(subset_error_metrics(true, prediction, np.ones_like(cliff_mask), "all"))
    metrics.update(subset_error_metrics(true, prediction, cliff_mask, "cliff"))
    metrics.update(subset_error_metrics(true, prediction, ~cliff_mask, "noncliff"))
    return metrics


def resolve_data_path(config: dict[str, Any], dataset_name: str) -> Path:
    paths = config.get("paths", {})
    if paths.get("raw_csv"):
        raw = str(paths["raw_csv"]).format(dataset=dataset_name)
        return Path(raw)
    return Path(paths.get("data_root", "output/molecular/reg/moleculeace")) / f"{dataset_name}.csv"


def resolve_output_paths(
    config: dict[str, Any],
    dataset_name: str,
    seed: int,
) -> tuple[Path, Path]:
    paths = config.get("paths", {})
    checkpoint_root = Path(paths.get("checkpoint_root", "checkpoints/cliff"))
    result_root = Path(paths.get("result_root", "logs/cliff"))
    leaf = Path(dataset_name) / f"seed_{seed}"
    return checkpoint_root / leaf / "best.pt", result_root / leaf / "result.json"


def count_cliffs(dataset: MoleculeACEDataset, indices: list[int]) -> int:
    return int(dataset.cliff_flags[np.asarray(indices, dtype=np.int64)].sum())


def train(
    config: dict[str, Any],
    dataset_override: str | None = None,
    seed_override: int | None = None,
    validate_only: bool = False,
) -> dict[str, Any]:
    paths = config.get("paths", {})
    data_cfg = config.get("data", {})
    model_cfg = config.get("model", {})
    train_cfg = config.get("train", {})
    experiment_cfg = config.get("experiment", {})

    dataset_name = str(dataset_override or data_cfg.get("dataset", DEFAULT_DATASET))
    seed = int(seed_override if seed_override is not None else train_cfg.get("seed", 42))
    set_seed(seed)
    raw_csv = resolve_data_path(config, dataset_name)
    if not raw_csv.exists():
        raise FileNotFoundError(
            f"MoleculeACE dataset not found: {raw_csv}. "
            f"Run: python scripts/download_moleculeace.py --datasets {dataset_name}"
        )

    dataset = MoleculeACEDataset(
        raw_csv=raw_csv,
        smiles_col=str(data_cfg.get("smiles_col", "smiles")),
        target_col=str(data_cfg.get("target_col", "y [pEC50/pKi]")),
        cliff_col=str(data_cfg.get("cliff_col", "cliff_mol")),
        split_col=str(data_cfg.get("split_col", "split")),
    )
    train_idx, val_idx, test_idx = split_official_train_validation(
        dataset,
        val_frac=float(data_cfg.get("val_frac_from_train", 0.1)),
        seed=int(data_cfg.get("validation_seed", 42)),
    )
    target_mean, target_std = target_statistics(dataset, train_idx)
    checkpoint_path, result_path = resolve_output_paths(config, dataset_name, seed)

    summary: dict[str, Any] = {
        "benchmark": "MoleculeACE",
        "dataset": dataset_name,
        "experiment": str(experiment_cfg.get("name", model_cfg.get("family", "model"))),
        "variant": str(experiment_cfg.get("variant", "original")),
        "model_family": str(model_cfg.get("family", "chemblm")),
        "seed": seed,
        "target_column": dataset.target_col,
        "dataset_size": len(dataset),
        "train_size": len(train_idx),
        "validation_size": len(val_idx),
        "test_size": len(test_idx),
        "train_cliffs": count_cliffs(dataset, train_idx),
        "validation_cliffs": count_cliffs(dataset, val_idx),
        "test_cliffs": count_cliffs(dataset, test_idx),
        "target_mean": target_mean,
        "target_std": target_std,
        "checkpoint": str(checkpoint_path),
    }

    print(f"MoleculeACE dataset: {dataset_name}")
    print(f"Data: {len(dataset)} molecules from {raw_csv}")
    print(
        "Split: official MoleculeACE train/test; validation is taken only from train "
        f"(train={len(train_idx)}, val={len(val_idx)}, test={len(test_idx)})"
    )
    print(
        "Cliff molecules: "
        f"train={summary['train_cliffs']} val={summary['validation_cliffs']} "
        f"test={summary['test_cliffs']}"
    )
    print(f"Train normalization: mean={target_mean:.8g} std={target_std:.8g}")
    if validate_only:
        print("Validation only: dataset columns, official split, and cliff labels are valid.")
        return summary

    device = resolve_device(str(train_cfg.get("device", "auto")))
    pretrained = load_checkpoint(paths.get("pretrained_checkpoint"))
    model = build_model(config, pretrained).to(device)
    trainable_parameters = [parameter for parameter in model.parameters() if parameter.requires_grad]
    if not trainable_parameters:
        raise ValueError("No trainable parameters remain after applying freeze_encoder.")

    batch_size = int(data_cfg.get("batch_size", 32))
    loader_kwargs = {
        "batch_size": batch_size,
        "num_workers": int(data_cfg.get("num_workers", 0)),
        "collate_fn": collate_cliff,
        "pin_memory": device.type == "cuda",
    }
    generator = torch.Generator().manual_seed(seed)
    train_loader = DataLoader(
        Subset(dataset, train_idx),
        shuffle=True,
        generator=generator,
        **loader_kwargs,
    )
    val_loader = DataLoader(Subset(dataset, val_idx), shuffle=False, **loader_kwargs)
    test_loader = DataLoader(Subset(dataset, test_idx), shuffle=False, **loader_kwargs)

    optimizer = torch.optim.AdamW(
        trainable_parameters,
        lr=float(train_cfg.get("lr", 1e-5)),
        weight_decay=float(train_cfg.get("weight_decay", 1e-2)),
    )
    epochs = int(train_cfg.get("epochs", 60))
    accumulation = max(1, int(train_cfg.get("gradient_accumulation_steps", 1)))
    updates_per_epoch = math.ceil(len(train_loader) / accumulation)
    total_updates = max(1, updates_per_epoch * epochs)
    warmup_steps = train_cfg.get("warmup_steps")
    if warmup_steps is None:
        warmup_steps = round(total_updates * float(train_cfg.get("warmup_ratio", 0.05)))
    scheduler = linear_warmup_decay(optimizer, int(warmup_steps), total_updates)
    amp_enabled = bool(train_cfg.get("amp", True)) and device.type == "cuda"
    scaler = make_grad_scaler(amp_enabled)

    print(f"Device: {device}")
    print(
        f"Model: {experiment_cfg.get('name', model_cfg.get('family'))} "
        f"variant={experiment_cfg.get('variant', 'original')}"
    )
    if pretrained is None:
        print("Initialization: original language model (no PG-MLD checkpoint)")
    else:
        print(f"Loaded PG-MLD checkpoint: {paths.get('pretrained_checkpoint')}")
    print(
        f"Optimization: epochs={epochs} batch={batch_size} accumulation={accumulation} "
        f"lr={float(train_cfg.get('lr', 1e-5)):.3g} amp={amp_enabled} "
        f"warmup_steps={int(warmup_steps)}"
    )
    print("Model selection: minimum validation RMSE; official test set is evaluated once.")

    best_val_rmse = float("inf")
    best_epoch = 0
    no_improvement = 0
    patience = max(0, int(train_cfg.get("early_stopping_patience", 10)))
    grad_clip = float(train_cfg.get("grad_clip", 1.0))
    log_every = max(1, int(train_cfg.get("log_every", 20)))

    for epoch in range(1, epochs + 1):
        model.train()
        optimizer.zero_grad(set_to_none=True)
        running_loss = 0.0
        steps = 0
        progress = tqdm(train_loader, desc=f"MoleculeACE {dataset_name} epoch {epoch}/{epochs}")
        for step, batch in enumerate(progress, start=1):
            labels = batch["labels"].to(device, non_blocking=True)
            with torch.autocast(
                device_type=device.type,
                dtype=torch.float16,
                enabled=amp_enabled,
            ):
                prediction = model(batch["smiles"], device=device)
                loss = normalized_mse(prediction, labels, target_mean, target_std)
                scaled_loss = loss / accumulation
            scaler.scale(scaled_loss).backward()

            should_update = step % accumulation == 0 or step == len(train_loader)
            if should_update:
                if grad_clip > 0:
                    scaler.unscale_(optimizer)
                    torch.nn.utils.clip_grad_norm_(trainable_parameters, grad_clip)
                scaler.step(optimizer)
                scaler.update()
                optimizer.zero_grad(set_to_none=True)
                scheduler.step()

            running_loss += float(loss.item())
            steps += 1
            if step % log_every == 0:
                progress.set_postfix(loss=running_loss / steps, lr=scheduler.get_last_lr()[0])

        val_metrics = evaluate(model, val_loader, device, target_mean, target_std)
        val_rmse = float(val_metrics["all_rmse"])
        improved = val_rmse < best_val_rmse
        if improved:
            best_val_rmse = val_rmse
            best_epoch = epoch
            no_improvement = 0
            atomic_torch_save(
                {
                    "epoch": epoch,
                    "encoder_type": "moleculeace_task_1d",
                    "model_state_dict": model.state_dict(),
                    "config": config,
                    "dataset": dataset_name,
                    "target_mean": target_mean,
                    "target_std": target_std,
                    "val_metrics": val_metrics,
                },
                checkpoint_path,
            )
        else:
            no_improvement += 1

        print(
            f"Epoch {epoch}: train_normalized_mse={running_loss / max(steps, 1):.6f} "
            f"val_RMSE={val_rmse:.6f} val_cliff_RMSE={float(val_metrics['cliff_rmse']):.6f} "
            f"best_epoch={best_epoch}"
        )
        if patience > 0 and no_improvement >= patience:
            print(
                f"Early stopping at epoch {epoch}: no validation RMSE improvement "
                f"for {patience} epochs (best_epoch={best_epoch})."
            )
            break

    if best_epoch == 0 or not checkpoint_path.exists():
        raise RuntimeError("Training finished without producing a valid best checkpoint.")
    try:
        best = torch.load(checkpoint_path, map_location=device, weights_only=False)
    except TypeError:
        best = torch.load(checkpoint_path, map_location=device)
    model.load_state_dict(best["model_state_dict"], strict=True)
    test_metrics = evaluate(model, test_loader, device, target_mean, target_std)
    summary.update({"best_epoch": best_epoch, "val": best["val_metrics"], "test": test_metrics})
    best["test_metrics"] = test_metrics
    atomic_torch_save(best, checkpoint_path)
    atomic_json_save(summary, result_path)

    print(
        f"Best epoch {best_epoch}: test_RMSE={float(test_metrics['all_rmse']):.6f} "
        f"test_cliff_RMSE={float(test_metrics['cliff_rmse']):.6f}"
    )
    print(f"Checkpoint: {checkpoint_path}")
    print(f"Result JSON: {result_path}")
    return summary


def build_arg_parser() -> argparse.ArgumentParser:
    parser = argparse.ArgumentParser(
        description="MoleculeACE activity-cliff regression for original or PG-MLD models."
    )
    parser.add_argument("--config", required=True)
    parser.add_argument("--dataset", default=None)
    parser.add_argument("--seed", type=int)
    parser.add_argument(
        "--validate-only",
        action="store_true",
        help="Validate data columns and official splits without loading a language model.",
    )
    return parser


def main() -> None:
    args = build_arg_parser().parse_args()
    train(
        load_config(args.config),
        dataset_override=args.dataset,
        seed_override=args.seed,
        validate_only=args.validate_only,
    )


if __name__ == "__main__":
    main()
```
