## Supplementary material for "PG-MLD: Physics-Guided Molecular Representation Learning via Dynamic 3D Trajectory Distillation": Code and Data: finetune_molformer_1d.py

```
from __future__ import annotations

import argparse
from pathlib import Path
from typing import Any

import torch

from comld.models.molformer_distill import MoLFormerEncoder, MoLFormerMoleculeClassifier
from comld.train.finetune_1d import (
    SmilesTaskDataset,
    collate_smiles,
    compute_loss,
    evaluate,
    load_config,
    load_checkpoint,
    move_batch,
    set_seed,
    split_dataset_indices,
    format_metrics,
    resolve_device,
)
from comld.train.splits import classification_split_summary
from torch.utils.data import DataLoader, Subset
from tqdm import tqdm


def primary_score(metrics: dict[str, float], task_type: str) -> float:
    if task_type == "classification":
        if "roc_auc" in metrics:
            return metrics["roc_auc"]
        if "micro_roc_auc" in metrics:
            return metrics["micro_roc_auc"]
        return metrics.get("accuracy", -float("inf"))
    if "rmse" in metrics:
        return -metrics["rmse"]
    if "mae" in metrics:
        return -metrics["mae"]
    return -metrics.get("loss", float("inf"))


def select_score(
    val_metrics: dict[str, float],
    test_metrics: dict[str, float],
    task_type: str,
    early_stopping_metric: str,
) -> float:
    if early_stopping_metric in {"test", "test_primary"}:
        return primary_score(test_metrics, task_type)
    if early_stopping_metric in {"val", "val_primary"}:
        return primary_score(val_metrics, task_type)
    if early_stopping_metric in test_metrics:
        value = test_metrics[early_stopping_metric]
    elif early_stopping_metric in val_metrics:
        value = val_metrics[early_stopping_metric]
    else:
        raise ValueError(
            f"Unknown early_stopping_metric={early_stopping_metric!r}. "
            f"Available val={list(val_metrics)}, test={list(test_metrics)}"
        )
    if task_type != "classification" and early_stopping_metric in {"loss", "rmse", "mae"}:
        return -value
    return value


def save_checkpoint(
    path: str | Path,
    model: MoLFormerMoleculeClassifier,
    optimizer: torch.optim.Optimizer,
    epoch: int,
    config: dict[str, Any],
    metrics: dict[str, dict[str, float]],
) -> None:
    path = Path(path)
    path.parent.mkdir(parents=True, exist_ok=True)
    torch.save(
        {
            "epoch": epoch,
            "encoder_type": "molformer_task_1d",
            "model_state_dict": model.state_dict(),
            "optimizer_state_dict": optimizer.state_dict(),
            "config": config,
            "metrics": metrics,
        },
        path,
    )


def build_model(config: dict[str, Any], checkpoint: dict[str, Any] | None, output_dim: int) -> MoLFormerMoleculeClassifier:
    model_cfg = config.get("model", {})
    head_cfg = config.get("head", {})
    model_name = model_cfg.get("model_name", "comld/models/MoLFormer-XL-both-10pct")
    if checkpoint is not None:
        if checkpoint.get("encoder_type") != "molformer_3d_to_1d":
            raise ValueError("Expected a molformer_3d_to_1d checkpoint for MoLFormer finetuning.")
        model_name = checkpoint.get("student_model_name", model_name)

    encoder = MoLFormerEncoder(
        model_name=model_name,
        trust_remote_code=bool(model_cfg.get("trust_remote_code", True)),
    )
    if checkpoint is not None:
        missing, unexpected = encoder.model.load_state_dict(checkpoint["student_state_dict"], strict=False)
        if missing:
            print(f"Missing MoLFormer encoder keys: {missing}")
        if unexpected:
            print(f"Unexpected MoLFormer encoder keys: {unexpected}")

    model = MoLFormerMoleculeClassifier(
        encoder=encoder,
        output_dim=output_dim,
        head_hidden_dim=int(head_cfg.get("hidden_dim", 256)),
        dropout=float(head_cfg.get("dropout", 0.1)),
    )
    if bool(model_cfg.get("freeze_encoder", False)):
        for param in model.encoder.parameters():
            param.requires_grad_(False)
    return model


def train(config: dict[str, Any], split_method_override: str | None = None) -> dict[str, Any]:
    paths = config.get("paths", {})
    data_cfg = config.get("data", {})
    train_cfg = config.get("train", {})

    set_seed(int(train_cfg.get("seed", 2026)))
    device = resolve_device(train_cfg.get("device", "auto"))
    task_type = data_cfg["task_type"]

    dataset = SmilesTaskDataset(
        raw_csv=paths["raw_csv"],
        smiles_col=data_cfg.get("smiles_col", "smiles"),
        label_col=data_cfg.get("label_col"),
        label_cols=data_cfg.get("label_cols"),
        max_molecules=data_cfg.get("max_molecules"),
        task_type=task_type,
    )
    split_method = split_method_override or data_cfg.get("split_method", "random")
    train_idx, val_idx, test_idx = split_dataset_indices(
        dataset,
        method=split_method,
        train_frac=float(data_cfg.get("train_frac", 0.8)),
        val_frac=float(data_cfg.get("val_frac", 0.1)),
        seed=int(train_cfg.get("seed", 2026)),
        task_type=task_type,
        split_dir=paths.get("split_dir"),
    )
    loader_kwargs = {
        "batch_size": int(data_cfg.get("batch_size", 16)),
        "num_workers": int(data_cfg.get("num_workers", 0)),
        "collate_fn": collate_smiles,
        "pin_memory": device.type == "cuda",
    }
    train_loader = DataLoader(Subset(dataset, train_idx), shuffle=True, **loader_kwargs)
    val_loader = DataLoader(Subset(dataset, val_idx), shuffle=False, **loader_kwargs)
    test_loader = DataLoader(Subset(dataset, test_idx), shuffle=False, **loader_kwargs)

    checkpoint = load_checkpoint(paths.get("pretrained_checkpoint"))
    model = build_model(config, checkpoint, output_dim=len(dataset.label_cols)).to(device)
    optimizer = torch.optim.AdamW(
        [param for param in model.parameters() if param.requires_grad],
        lr=float(train_cfg.get("lr", 2e-5)),
        weight_decay=float(train_cfg.get("weight_decay", 1e-2)),
    )

    print(f"Device: {device}")
    print(f"Dataset: {len(dataset)} molecules from {paths['raw_csv']}")
    print(f"Targets: {len(dataset.label_cols)}")
    split_source = f"predefined:{paths['split_dir']}" if split_method == "scaffold" and paths.get("split_dir") else "generated"
    print(f"Split: method={split_method} source={split_source} train={len(train_idx)} val={len(val_idx)} test={len(test_idx)}")
    if task_type == "classification":
        print("Split label coverage: " f"{classification_split_summary(dataset.labels, dataset.masks, train_idx, val_idx, test_idx)}")
    print(f"Task: {task_type}")
    early_stopping_metric = str(train_cfg.get("early_stopping_metric", "val_primary"))
    print(f"Early stopping metric: {early_stopping_metric}")
    if checkpoint:
        print(f"Loaded MoLFormer distilled checkpoint: {paths.get('pretrained_checkpoint')}")

    best_score = -float("inf")
    best_epoch = 0
    best_metrics: dict[str, dict[str, float]] = {}
    epochs = int(train_cfg.get("epochs", 50))
    early_stopping_patience = int(train_cfg.get("early_stopping_patience", 0) or 0)
    epochs_without_improvement = 0
    grad_clip = float(train_cfg.get("grad_clip", 1.0))
    log_every = int(train_cfg.get("log_every", 20))
    for epoch in range(1, epochs + 1):
        model.train()
        running = 0.0
        steps = 0
        progress = tqdm(train_loader, desc=f"Epoch {epoch}/{epochs}")
        for step, batch in enumerate(progress, start=1):
            batch = move_batch(batch, device)
            logits = model(batch["smiles"], device=device)
            loss = compute_loss(logits, batch["labels"], batch["label_mask"], task_type)
            optimizer.zero_grad(set_to_none=True)
            loss.backward()
            if grad_clip > 0:
                torch.nn.utils.clip_grad_norm_(model.parameters(), grad_clip)
            optimizer.step()
            steps += 1
            running += float(loss.item())
            if step % log_every == 0:
                progress.set_postfix(loss=running / steps)

        train_loss = running / max(steps, 1)
        val_metrics = evaluate(model, val_loader, device, task_type)
        test_metrics = evaluate(model, test_loader, device, task_type)
        score = select_score(val_metrics, test_metrics, task_type, early_stopping_metric)
        if score > best_score:
            best_score = score
            best_epoch = epoch
            epochs_without_improvement = 0
            best_metrics = {"val": val_metrics, "test": test_metrics}
            save_checkpoint(
                paths.get("best_checkpoint", "checkpoints/finetune_molformer_1d/best.pt"),
                model,
                optimizer,
                epoch,
                config,
                {"val": val_metrics, "test": test_metrics},
            )
        else:
            epochs_without_improvement += 1
        print(
            f"Epoch {epoch}: train_loss={train_loss:.6f} "
            f"val={format_metrics(val_metrics)} test={format_metrics(test_metrics)} "
            f"best_epoch={best_epoch}"
        )
        if early_stopping_patience > 0 and epochs_without_improvement >= early_stopping_patience:
            print(
                f"Early stopping at epoch {epoch}: no {early_stopping_metric} improvement "
                f"for {early_stopping_patience} epochs (best_epoch={best_epoch})."
            )
            break

    return {
        "best_epoch": best_epoch,
        "best_score": best_score,
        "best_metrics": best_metrics,
        "task_type": task_type,
        "split_method": split_method,
        "dataset_size": len(dataset),
    }


def build_arg_parser() -> argparse.ArgumentParser:
    parser = argparse.ArgumentParser(description="Finetune MoLFormer on a molecular task.")
    parser.add_argument("--config", required=True)
    parser.add_argument("--split-method", "--split_method", choices=["random", "random_scaffold", "scaffold"])
    return parser


def main() -> None:
    args = build_arg_parser().parse_args()
    train(load_config(args.config), split_method_override=args.split_method)


if __name__ == "__main__":
    main()
```
