## Supplementary material for "PG-MLD: Physics-Guided Molecular Representation Learning via Dynamic 3D Trajectory Distillation": Code and Data: finetune_qm9_1d.py

```
from __future__ import annotations

import argparse
import json
import math
import random
from pathlib import Path
from typing import Any

import numpy as np
import pandas as pd
import torch
import torch.nn.functional as F
import yaml
from torch.utils.data import DataLoader, Dataset, Subset
from tqdm import tqdm

from comld.train.finetune_1d import load_checkpoint, resolve_device


QM9_TARGETS = ("mu", "alpha", "homo", "lumo", "gap", "zpve", "u0", "u", "h")
QM9_REPORT_UNITS = {
    "mu": "D",
    "alpha": "a0^3",
    "homo": "meV",
    "lumo": "meV",
    "gap": "meV",
    "zpve": "meV",
    "u0": "meV",
    "u": "meV",
    "h": "meV",
}
QM9_REPORT_FACTORS = {
    "mu": 1.0,
    "alpha": 1.0,
    "homo": 1000.0,
    "lumo": 1000.0,
    "gap": 1000.0,
    "zpve": 1000.0,
    "u0": 1000.0,
    "u": 1000.0,
    "h": 1000.0,
}
HARTREE_TO_EV = 27.211386246
QM9_SOURCE_ALIASES = {
    "mu": ("mu",),
    "alpha": ("alpha",),
    "homo": ("homo",),
    "lumo": ("lumo",),
    "gap": ("gap",),
    "zpve": ("zpve",),
    "u0": ("u0", "U0"),
    "u": ("u298", "U"),
    "h": ("h298", "H"),
}
QM9_ENERGY_TARGETS = {"homo", "lumo", "gap", "zpve", "u0", "u", "h"}


class QM9SmilesDataset(Dataset):
    """A single-target QM9 SMILES dataset loaded from the official CSV.

    DeepChem's CSV stores energy targets in Hartree. They are converted to eV
    for training and then to meV only when metrics are reported.
    """

    def __init__(
        self,
        raw_csv: str | Path,
        target: str,
        smiles_col: str = "smiles",
        source_energy_unit: str = "hartree",
        uncharacterized_txt: str | Path | None = None,
        max_molecules: int | None = None,
    ) -> None:
        if target not in QM9_TARGETS:
            raise ValueError(f"Unsupported QM9 target {target!r}. Choose from {QM9_TARGETS}.")

        frame = pd.read_csv(raw_csv)
        lower_to_original = {str(column).lower(): str(column) for column in frame.columns}
        source_target = next(
            (
                lower_to_original[alias.lower()]
                for alias in QM9_SOURCE_ALIASES[target]
                if alias.lower() in lower_to_original
            ),
            None,
        )
        if smiles_col not in frame.columns or source_target is None:
            raise ValueError(
                f"Cannot resolve smiles={smiles_col!r} and target={target!r} in {raw_csv}. "
                f"Columns: {list(frame.columns)}"
            )

        if uncharacterized_txt:
            skip_path = Path(uncharacterized_txt)
            if not skip_path.exists():
                raise FileNotFoundError(
                    f"QM9 uncharacterized list not found: {skip_path}. "
                    "Download it or remove paths.uncharacterized_txt to use all CSV rows."
                )
            lines = skip_path.read_text(encoding="utf-8", errors="replace").splitlines()
            skip = {
                int(fields[0]) - 1
                for line in lines[9:]
                if (fields := line.split()) and fields[0].isdigit()
            }
            if len(skip) != 3054:
                raise ValueError(
                    f"Expected 3054 uncharacterized QM9 indices, parsed {len(skip)} from {skip_path}."
                )
            frame = frame.drop(index=[index for index in skip if index in frame.index])
            frame = frame.reset_index(drop=True)

        values = pd.to_numeric(frame[source_target], errors="coerce")
        source_energy_unit = source_energy_unit.strip().lower()
        if target in QM9_ENERGY_TARGETS:
            if source_energy_unit == "hartree":
                values = values * HARTREE_TO_EV
            elif source_energy_unit != "ev":
                raise ValueError(
                    f"Unsupported source_energy_unit={source_energy_unit!r}; use hartree or ev."
                )
        valid = frame[smiles_col].notna() & values.notna() & np.isfinite(values.to_numpy(dtype=float))
        smiles_values = frame.loc[valid, smiles_col].astype(str).str.strip().tolist()
        label_values = values.loc[valid].astype(np.float32).to_numpy()
        if max_molecules is not None:
            limit = int(max_molecules)
            smiles_values = smiles_values[:limit]
            label_values = label_values[:limit]
        if not smiles_values:
            raise ValueError(f"No valid rows for target {target!r} in {raw_csv}")

        self.smiles = smiles_values
        self.labels = label_values
        self.target = target

    def __len__(self) -> int:
        return len(self.smiles)

    def __getitem__(self, index: int) -> dict[str, Any]:
        return {
            "smiles": self.smiles[index],
            "label": torch.tensor(self.labels[index], dtype=torch.float32),
        }


def collate_qm9(items: list[dict[str, Any]]) -> dict[str, Any]:
    return {
        "smiles": [item["smiles"] for item in items],
        "labels": torch.stack([item["label"] for item in items], dim=0),
    }


def load_config(path: str | Path) -> dict[str, Any]:
    with open(path, "r", encoding="utf-8") as handle:
        return yaml.safe_load(handle) or {}


def set_seed(seed: int) -> None:
    random.seed(seed)
    np.random.seed(seed)
    torch.manual_seed(seed)
    if torch.cuda.is_available():
        torch.cuda.manual_seed_all(seed)


def load_fixed_split(path: str | Path, dataset_size: int) -> tuple[list[int], list[int], list[int]]:
    split_path = Path(path)
    if not split_path.exists():
        raise FileNotFoundError(
            f"QM9 split not found: {split_path}. Remove paths.split_npz to use "
            "the deterministic in-memory split."
        )
    with np.load(split_path) as data:
        required = ("train", "val", "test")
        missing = [key for key in required if key not in data]
        if missing:
            raise ValueError(f"Split file {split_path} is missing arrays: {missing}")
        arrays = [np.asarray(data[key], dtype=np.int64) for key in required]

    all_indices = np.concatenate(arrays)
    if all_indices.size != dataset_size:
        raise ValueError(
            f"Split contains {all_indices.size} rows but dataset contains {dataset_size}. "
            "Do not use max_molecules with the full QM9 split."
        )
    if all_indices.min(initial=0) < 0 or all_indices.max(initial=-1) >= dataset_size:
        raise ValueError(f"Split {split_path} contains out-of-range indices.")
    if np.unique(all_indices).size != dataset_size:
        raise ValueError(f"Split {split_path} has duplicate or missing indices.")
    return tuple(array.tolist() for array in arrays)  # type: ignore[return-value]


def make_fixed_random_split(
    dataset_size: int,
    train_size: int,
    val_size: int,
    split_seed: int,
) -> tuple[list[int], list[int], list[int]]:
    if train_size <= 0 or val_size <= 0 or train_size + val_size >= dataset_size:
        raise ValueError(
            f"Invalid QM9 split sizes: dataset={dataset_size}, train={train_size}, val={val_size}"
        )
    permutation = np.random.RandomState(split_seed).permutation(dataset_size)
    train = permutation[:train_size]
    val = permutation[train_size : train_size + val_size]
    test = permutation[train_size + val_size :]
    return train.tolist(), val.tolist(), test.tolist()


def target_statistics(dataset: QM9SmilesDataset, indices: list[int]) -> tuple[float, float]:
    values = dataset.labels[np.asarray(indices, dtype=np.int64)].astype(np.float64)
    mean = float(values.mean())
    std = float(values.std())
    if not math.isfinite(mean) or not math.isfinite(std) or std < 1e-12:
        raise ValueError(f"Invalid train statistics for {dataset.target}: mean={mean}, std={std}")
    return mean, std


def build_model(config: dict[str, Any], checkpoint: dict[str, Any] | None) -> torch.nn.Module:
    model_cfg = config.get("model", {})
    family = str(model_cfg.get("family", "chemberta")).strip().lower()
    if checkpoint is not None and model_cfg.get("model_name"):
        # Distillation checkpoints may contain an absolute path from the
        # training server. The QM9 config is the portable source of truth for
        # the same backbone on the current machine.
        checkpoint = dict(checkpoint)
        checkpoint["student_model_name"] = str(model_cfg["model_name"])
    if family in {"chemberta", "chemberta2", "chemberta-2"}:
        from comld.train.finetune_1d import build_model as build_chemberta

        return build_chemberta(config, checkpoint, output_dim=1)
    if family == "chemblm":
        from comld.train.finetune_chemblm_1d import build_model as build_chemblm

        return build_chemblm(config, checkpoint, output_dim=1)
    if family in {"molformer", "molformer-xl"}:
        from comld.train.finetune_molformer_1d import build_model as build_molformer

        return build_molformer(config, checkpoint, output_dim=1)
    raise ValueError(
        f"Unsupported model.family={family!r}. "
        "Use chemberta, chemberta2, chemblm, or molformer."
    )


def normalized_mse(
    predictions: torch.Tensor,
    labels: torch.Tensor,
    target_mean: float,
    target_std: float,
) -> torch.Tensor:
    predictions = predictions.reshape(-1)
    normalized_labels = (labels.reshape(-1) - target_mean) / target_std
    return F.mse_loss(predictions, normalized_labels)


@torch.no_grad()
def evaluate(
    model: torch.nn.Module,
    loader: DataLoader,
    device: torch.device,
    target: str,
    target_mean: float,
    target_std: float,
) -> dict[str, float]:
    model.eval()
    predictions: list[np.ndarray] = []
    labels: list[np.ndarray] = []
    normalized_losses: list[float] = []
    for batch in loader:
        y = batch["labels"].to(device, non_blocking=True)
        pred_norm = model(batch["smiles"], device=device).reshape(-1)
        loss = normalized_mse(pred_norm, y, target_mean, target_std)
        pred_native = pred_norm * target_std + target_mean
        predictions.append(pred_native.detach().cpu().numpy())
        labels.append(y.detach().cpu().numpy())
        normalized_losses.append(float(loss.item()))

    if not predictions:
        raise ValueError("Cannot evaluate an empty loader.")
    pred = np.concatenate(predictions)
    true = np.concatenate(labels)
    error = pred - true
    native_mae = float(np.mean(np.abs(error)))
    native_rmse = float(np.sqrt(np.mean(np.square(error))))
    factor = QM9_REPORT_FACTORS[target]
    return {
        "mae": native_mae * factor,
        "rmse": native_rmse * factor,
        "normalized_mae": native_mae / target_std,
        "normalized_mse": float(np.mean(normalized_losses)),
    }


def atomic_torch_save(payload: dict[str, Any], path: str | Path) -> None:
    destination = Path(path)
    destination.parent.mkdir(parents=True, exist_ok=True)
    temporary = destination.with_suffix(destination.suffix + ".tmp")
    try:
        torch.save(payload, temporary)
        temporary.replace(destination)
    except BaseException:
        temporary.unlink(missing_ok=True)
        raise


def atomic_json_save(payload: dict[str, Any], path: str | Path) -> None:
    destination = Path(path)
    destination.parent.mkdir(parents=True, exist_ok=True)
    temporary = destination.with_suffix(destination.suffix + ".tmp")
    try:
        temporary.write_text(
            json.dumps(payload, indent=2, ensure_ascii=False),
            encoding="utf-8",
        )
        temporary.replace(destination)
    except BaseException:
        temporary.unlink(missing_ok=True)
        raise


def linear_warmup_decay(
    optimizer: torch.optim.Optimizer,
    warmup_steps: int,
    total_steps: int,
) -> torch.optim.lr_scheduler.LambdaLR:
    warmup_steps = max(0, int(warmup_steps))
    total_steps = max(1, int(total_steps))

    def multiplier(step: int) -> float:
        if warmup_steps > 0 and step < warmup_steps:
            return float(step + 1) / float(warmup_steps)
        remaining = max(total_steps - step, 0)
        decay_steps = max(total_steps - warmup_steps, 1)
        return float(remaining) / float(decay_steps)

    return torch.optim.lr_scheduler.LambdaLR(optimizer, multiplier)


def resolve_output_paths(
    config: dict[str, Any],
    target: str,
    seed: int,
) -> tuple[Path, Path]:
    paths = config.get("paths", {})
    checkpoint_root = Path(paths.get("checkpoint_root", "checkpoints/qm9"))
    result_root = Path(paths.get("result_root", "logs/qm9"))
    leaf = Path(target) / f"seed_{seed}"
    return checkpoint_root / leaf / "best.pt", result_root / leaf / "result.json"


def make_grad_scaler(enabled: bool) -> torch.cuda.amp.GradScaler:
    try:
        return torch.amp.GradScaler("cuda", enabled=enabled)  # type: ignore[return-value]
    except (AttributeError, TypeError):
        return torch.cuda.amp.GradScaler(enabled=enabled)


def train(
    config: dict[str, Any],
    target_override: str | None = None,
    seed_override: int | None = None,
    validate_only: bool = False,
) -> dict[str, Any]:
    paths = config.get("paths", {})
    data_cfg = config.get("data", {})
    train_cfg = config.get("train", {})
    model_cfg = config.get("model", {})
    experiment_cfg = config.get("experiment", {})

    target = str(target_override or data_cfg.get("target", "mu")).lower()
    if target not in QM9_TARGETS:
        raise ValueError(f"Unsupported target {target!r}; choose from {QM9_TARGETS}")
    seed = int(seed_override if seed_override is not None else train_cfg.get("seed", 42))
    set_seed(seed)

    dataset = QM9SmilesDataset(
        raw_csv=paths["raw_csv"],
        target=target,
        smiles_col=data_cfg.get("smiles_col", "smiles"),
        source_energy_unit=data_cfg.get("source_energy_unit", "hartree"),
        uncharacterized_txt=paths.get("uncharacterized_txt"),
        max_molecules=data_cfg.get("max_molecules"),
    )
    split_path = paths.get("split_npz")
    if split_path:
        train_idx, val_idx, test_idx = load_fixed_split(split_path, len(dataset))
        split_description = str(split_path)
    else:
        split_seed = int(data_cfg.get("split_seed", 42))
        train_idx, val_idx, test_idx = make_fixed_random_split(
            len(dataset),
            train_size=int(data_cfg.get("train_size", 110000)),
            val_size=int(data_cfg.get("val_size", 10000)),
            split_seed=split_seed,
        )
        split_description = f"in-memory random seed={split_seed}"
    target_mean, target_std = target_statistics(dataset, train_idx)

    checkpoint_path, result_path = resolve_output_paths(config, target, seed)
    summary = {
        "experiment": str(experiment_cfg.get("name", model_cfg.get("family", "model"))),
        "variant": str(experiment_cfg.get("variant", "original")),
        "model_family": str(model_cfg.get("family", "chemberta")),
        "target": target,
        "unit": QM9_REPORT_UNITS[target],
        "seed": seed,
        "dataset_size": len(dataset),
        "train_size": len(train_idx),
        "val_size": len(val_idx),
        "test_size": len(test_idx),
        "target_mean_native": target_mean,
        "target_std_native": target_std,
        "checkpoint": str(checkpoint_path),
    }

    print(f"QM9 target: {target} ({QM9_REPORT_UNITS[target]})")
    print(f"Dataset: {len(dataset)} molecules from {paths['raw_csv']}")
    print(
        f"Fixed split: train={len(train_idx)} val={len(val_idx)} test={len(test_idx)} "
        f"from {split_description}"
    )
    print(f"Train normalization: mean={target_mean:.8g} std={target_std:.8g}")
    if validate_only:
        print("Validation only: dataset, target, and fixed split are valid.")
        return summary

    device = resolve_device(str(train_cfg.get("device", "auto")))
    pretrained = load_checkpoint(paths.get("pretrained_checkpoint"))
    model = build_model(config, pretrained).to(device)
    trainable_parameters = [parameter for parameter in model.parameters() if parameter.requires_grad]
    if not trainable_parameters:
        raise ValueError("No trainable parameters remain after applying freeze_encoder.")

    batch_size = int(data_cfg.get("batch_size", 32))
    loader_kwargs = {
        "batch_size": batch_size,
        "num_workers": int(data_cfg.get("num_workers", 0)),
        "collate_fn": collate_qm9,
        "pin_memory": device.type == "cuda",
    }
    generator = torch.Generator()
    generator.manual_seed(seed)
    train_loader = DataLoader(
        Subset(dataset, train_idx),
        shuffle=True,
        generator=generator,
        **loader_kwargs,
    )
    val_loader = DataLoader(Subset(dataset, val_idx), shuffle=False, **loader_kwargs)
    test_loader = DataLoader(Subset(dataset, test_idx), shuffle=False, **loader_kwargs)

    optimizer = torch.optim.AdamW(
        trainable_parameters,
        lr=float(train_cfg.get("lr", 2e-5)),
        weight_decay=float(train_cfg.get("weight_decay", 1e-2)),
    )
    epochs = int(train_cfg.get("epochs", 30))
    accumulation = max(1, int(train_cfg.get("gradient_accumulation_steps", 1)))
    updates_per_epoch = math.ceil(len(train_loader) / accumulation)
    total_updates = max(1, updates_per_epoch * epochs)
    warmup_steps = train_cfg.get("warmup_steps")
    if warmup_steps is None:
        warmup_steps = round(total_updates * float(train_cfg.get("warmup_ratio", 0.05)))
    scheduler = linear_warmup_decay(optimizer, int(warmup_steps), total_updates)
    amp_enabled = bool(train_cfg.get("amp", True)) and device.type == "cuda"
    scaler = make_grad_scaler(amp_enabled)

    print(f"Device: {device}")
    print(
        f"Model: {experiment_cfg.get('name', model_cfg.get('family'))} "
        f"variant={experiment_cfg.get('variant', 'original')}"
    )
    if pretrained is not None:
        print(f"Loaded PG-MLD checkpoint: {paths.get('pretrained_checkpoint')}")
    else:
        print("Initialization: original language model (no PG-MLD checkpoint)")
    print(
        f"Optimization: epochs={epochs} batch={batch_size} accumulation={accumulation} "
        f"lr={float(train_cfg.get('lr', 2e-5)):.3g} amp={amp_enabled} "
        f"warmup_steps={int(warmup_steps)} total_updates={total_updates}"
    )
    print("Model selection: minimum validation MAE; test set is evaluated only once after selection.")

    best_val_mae = float("inf")
    best_epoch = 0
    patience = max(0, int(train_cfg.get("early_stopping_patience", 8)))
    no_improvement = 0
    grad_clip = float(train_cfg.get("grad_clip", 1.0))
    log_every = max(1, int(train_cfg.get("log_every", 100)))

    for epoch in range(1, epochs + 1):
        model.train()
        optimizer.zero_grad(set_to_none=True)
        running_loss = 0.0
        steps = 0
        progress = tqdm(train_loader, desc=f"QM9 {target} epoch {epoch}/{epochs}")
        for step, batch in enumerate(progress, start=1):
            labels = batch["labels"].to(device, non_blocking=True)
            with torch.autocast(
                device_type=device.type,
                dtype=torch.float16,
                enabled=amp_enabled,
            ):
                pred_norm = model(batch["smiles"], device=device)
                loss = normalized_mse(pred_norm, labels, target_mean, target_std)
                scaled_loss = loss / accumulation
            scaler.scale(scaled_loss).backward()

            should_update = step % accumulation == 0 or step == len(train_loader)
            if should_update:
                if grad_clip > 0:
                    scaler.unscale_(optimizer)
                    torch.nn.utils.clip_grad_norm_(trainable_parameters, grad_clip)
                scaler.step(optimizer)
                scaler.update()
                optimizer.zero_grad(set_to_none=True)
                scheduler.step()

            running_loss += float(loss.item())
            steps += 1
            if step % log_every == 0:
                progress.set_postfix(loss=running_loss / steps, lr=scheduler.get_last_lr()[0])

        val_metrics = evaluate(model, val_loader, device, target, target_mean, target_std)
        train_loss = running_loss / max(steps, 1)
        improved = val_metrics["mae"] < best_val_mae
        if improved:
            best_val_mae = val_metrics["mae"]
            best_epoch = epoch
            no_improvement = 0
            atomic_torch_save(
                {
                    "epoch": epoch,
                    "encoder_type": "qm9_task_1d",
                    "model_state_dict": model.state_dict(),
                    "config": config,
                    "target": target,
                    "target_mean_native": target_mean,
                    "target_std_native": target_std,
                    "report_unit": QM9_REPORT_UNITS[target],
                    "val_metrics": val_metrics,
                },
                checkpoint_path,
            )
        else:
            no_improvement += 1

        print(
            f"Epoch {epoch}: train_normalized_mse={train_loss:.6f} "
            f"val_mae={val_metrics['mae']:.6f} {QM9_REPORT_UNITS[target]} "
            f"val_normalized_mae={val_metrics['normalized_mae']:.6f} "
            f"best_epoch={best_epoch}"
        )
        if patience > 0 and no_improvement >= patience:
            print(
                f"Early stopping at epoch {epoch}: no validation MAE improvement "
                f"for {patience} epochs (best_epoch={best_epoch})."
            )
            break

    if best_epoch == 0 or not checkpoint_path.exists():
        raise RuntimeError("Training finished without producing a valid best checkpoint.")
    try:
        best = torch.load(checkpoint_path, map_location=device, weights_only=False)
    except TypeError:
        best = torch.load(checkpoint_path, map_location=device)
    model.load_state_dict(best["model_state_dict"], strict=True)
    test_metrics = evaluate(model, test_loader, device, target, target_mean, target_std)
    summary.update(
        {
            "best_epoch": best_epoch,
            "val": best["val_metrics"],
            "test": test_metrics,
        }
    )
    best["test_metrics"] = test_metrics
    atomic_torch_save(best, checkpoint_path)
    atomic_json_save(summary, result_path)

    print(
        f"Best epoch {best_epoch}: test_MAE={test_metrics['mae']:.6f} "
        f"{QM9_REPORT_UNITS[target]}"
    )
    print(f"Checkpoint: {checkpoint_path}")
    print(f"Result JSON: {result_path}")
    return summary


def build_arg_parser() -> argparse.ArgumentParser:
    parser = argparse.ArgumentParser(
        description="Single-property QM9 finetuning for original or PG-MLD-distilled language models."
    )
    parser.add_argument("--config", required=True)
    parser.add_argument("--target", choices=QM9_TARGETS)
    parser.add_argument("--seed", type=int)
    parser.add_argument(
        "--validate-only",
        action="store_true",
        help="Validate CSV columns and the fixed split without loading a language model.",
    )
    return parser


def main() -> None:
    args = build_arg_parser().parse_args()
    train(
        load_config(args.config),
        target_override=args.target,
        seed_override=args.seed,
        validate_only=args.validate_only,
    )


if __name__ == "__main__":
    main()
```
