## Supplementary material for "PG-MLD: Physics-Guided Molecular Representation Learning via Dynamic 3D Trajectory Distillation": Code and Data: losses.py

```
from __future__ import annotations

import torch
import torch.nn.functional as F


def edge_distance_loss(
    pred_coords: torch.Tensor,
    target_coords: torch.Tensor,
    edge_index: torch.Tensor,
) -> torch.Tensor:
    if edge_index.numel() == 0:
        return pred_coords.new_tensor(0.0)
    src, dst = edge_index[0], edge_index[1]
    pred_dist = torch.linalg.norm(pred_coords[src] - pred_coords[dst], dim=-1)
    target_dist = torch.linalg.norm(target_coords[src] - target_coords[dst], dim=-1)
    return F.mse_loss(pred_dist, target_dist)
```
