## Supplementary material for "PG-MLD: Physics-Guided Molecular Representation Learning via Dynamic 3D Trajectory Distillation": Code and Data: metrics.py

```
from __future__ import annotations

import math

import numpy as np


def binary_roc_auc(y_true: np.ndarray, y_score: np.ndarray) -> float:
    """Compute binary ROC-AUC with average ranks, including tied predictions."""
    y_true = np.asarray(y_true).reshape(-1)
    y_score = np.asarray(y_score, dtype=float).reshape(-1)
    if y_true.shape != y_score.shape:
        raise ValueError(
            f"ROC-AUC shape mismatch: labels={y_true.shape}, scores={y_score.shape}"
        )
    finite = np.isfinite(y_score)
    y_true = y_true[finite]
    y_score = y_score[finite]
    positives = y_true == 1
    negatives = y_true == 0
    n_pos = int(positives.sum())
    n_neg = int(negatives.sum())
    if n_pos == 0 or n_neg == 0:
        return float("nan")

    order = np.argsort(y_score, kind="mergesort")
    sorted_scores = y_score[order]
    ranks = np.empty(y_score.shape[0], dtype=float)
    start = 0
    while start < sorted_scores.shape[0]:
        end = start + 1
        while end < sorted_scores.shape[0] and sorted_scores[end] == sorted_scores[start]:
            end += 1
        average_rank = 0.5 * ((start + 1) + end)
        ranks[order[start:end]] = average_rank
        start = end

    positive_rank_sum = float(ranks[positives].sum())
    return (
        positive_rank_sum - n_pos * (n_pos + 1) / 2.0
    ) / float(n_pos * n_neg)


def sigmoid(logits: np.ndarray) -> np.ndarray:
    logits = np.asarray(logits)
    positive = logits >= 0
    out = np.empty_like(logits, dtype=float)
    out[positive] = 1.0 / (1.0 + np.exp(-logits[positive]))
    exp_logits = np.exp(logits[~positive])
    out[~positive] = exp_logits / (1.0 + exp_logits)
    return out


def regression_metrics(pred: np.ndarray, target: np.ndarray, mask: np.ndarray | None = None) -> dict[str, float]:
    pred = np.asarray(pred)
    target = np.asarray(target)
    if mask is not None:
        valid = np.asarray(mask).astype(bool)
        pred = pred[valid]
        target = target[valid]
    else:
        pred = pred.reshape(-1)
        target = target.reshape(-1)
    if pred.size == 0:
        return {"mse": float("nan"), "rmse": float("nan"), "mae": float("nan"), "r2": float("nan")}
    mse = float(np.mean((pred - target) ** 2))
    mae = float(np.mean(np.abs(pred - target)))
    rmse = math.sqrt(mse)
    denom = float(np.sum((target - target.mean()) ** 2))
    r2 = float(1.0 - np.sum((pred - target) ** 2) / denom) if denom > 0 else float("nan")
    return {"mse": mse, "rmse": rmse, "mae": mae, "r2": r2}


def classification_metrics(logits: np.ndarray, target: np.ndarray, mask: np.ndarray | None = None) -> dict[str, float]:
    logits = np.asarray(logits)
    target = np.asarray(target)
    prob = sigmoid(logits)
    pred = (prob >= 0.5).astype(int)

    if mask is None:
        valid = np.ones_like(target, dtype=bool)
    else:
        valid = np.asarray(mask).astype(bool)
    valid = valid & ((target == 0) | (target == 1))
    if valid.sum() == 0:
        return {"accuracy": float("nan"), "roc_auc": float("nan"), "valid_tasks": 0.0}

    target_int = target.astype(int)
    acc = float(np.mean(pred[valid] == target_int[valid]))
    metrics = {"accuracy": acc, "roc_auc": float("nan"), "valid_tasks": 0.0}

    if target_int.ndim == 1 or target_int.shape[-1] == 1:
        y_true = target_int[valid].reshape(-1)
        y_prob = prob[valid].reshape(-1)
        if len(np.unique(y_true)) < 2:
            return metrics
        auc = binary_roc_auc(y_true, y_prob)
        if np.isfinite(auc):
            metrics["roc_auc"] = float(auc)
            metrics["valid_tasks"] = 1.0
        return metrics

    aucs = []
    for task_idx in range(target_int.shape[-1]):
        task_valid = valid[:, task_idx]
        if task_valid.sum() == 0:
            continue
        y_true = target_int[:, task_idx][task_valid]
        if len(np.unique(y_true)) < 2:
            continue
        y_prob = prob[:, task_idx][task_valid]
        auc = binary_roc_auc(y_true, y_prob)
        if np.isfinite(auc):
            aucs.append(float(auc))
    if aucs:
        metrics["roc_auc"] = float(np.mean(aucs))
        metrics["valid_tasks"] = float(len(aucs))
    flat_true = target_int[valid].reshape(-1)
    flat_prob = prob[valid].reshape(-1)
    if len(np.unique(flat_true)) >= 2:
        micro_auc = binary_roc_auc(flat_true, flat_prob)
        if np.isfinite(micro_auc):
            metrics["micro_roc_auc"] = float(micro_auc)
    return metrics
```
