## Supplementary material for "PG-MLD: Physics-Guided Molecular Representation Learning via Dynamic 3D Trajectory Distillation": Code and Data: pretrain_dynamic_3d.py

```
from __future__ import annotations

import argparse
import math
import random
from pathlib import Path
from typing import Any

import numpy as np
import torch
import torch.nn.functional as F
import yaml
from torch.utils.data import DataLoader, IterableDataset
from tqdm import tqdm

from comld.data.dynamic_md_dataset import (
    ShardedDynamicMDTrajectoryDataset,
    build_dynamic_md_dataset,
    collate_dynamic_md,
    dynamic_md_sample,
)
from comld.models.dynamic_se3_liquid import DynamicSE3LiquidTeacher
from comld.train.losses import edge_distance_loss


def load_config(path: str | Path) -> dict[str, Any]:
    with open(path, "r", encoding="utf-8") as handle:
        return yaml.safe_load(handle) or {}


def set_seed(seed: int) -> None:
    random.seed(seed)
    np.random.seed(seed)
    torch.manual_seed(seed)
    if torch.cuda.is_available():
        torch.cuda.manual_seed_all(seed)


def resolve_device(device_name: str) -> torch.device:
    if device_name == "auto":
        return torch.device("cuda" if torch.cuda.is_available() else "cpu")
    return torch.device(device_name)


def move_batch(batch: dict[str, Any], device: torch.device) -> dict[str, Any]:
    moved = dict(batch)
    for key in [
        "node_feats",
        "coords",
        "edge_index",
        "edge_attr",
        "frame_batch",
        "atom_batch",
        "electronic_feats",
        "atom_counts",
        "trajectory_frame_ptr",
        "delta_times_ps",
    ]:
        moved[key] = batch[key].to(device)
    return moved


def temporal_prediction_loss(
    model: DynamicSE3LiquidTeacher,
    frame_hidden: torch.Tensor,
    frame_ptr: torch.Tensor,
) -> torch.Tensor:
    losses = []
    ptr = frame_ptr.detach().cpu().tolist()
    for start, end in zip(ptr[:-1], ptr[1:]):
        sequence = frame_hidden[int(start) : int(end)]
        if sequence.shape[0] < 2:
            continue
        prediction = model.temporal_predictor(sequence[:-1])
        losses.append(F.mse_loss(prediction, sequence[1:].detach()))
    if not losses:
        return frame_hidden.new_tensor(0.0)
    return torch.stack(losses).mean()


def trajectory_consistency_loss(
    outputs: dict[str, torch.Tensor],
    frame_ptr: torch.Tensor,
) -> torch.Tensor:
    ptr = frame_ptr.detach().cpu().tolist()
    mean_frame_embeddings = []
    for start, end in zip(ptr[:-1], ptr[1:]):
        mean_frame_embeddings.append(
            outputs["frame_mol_embeddings"][int(start) : int(end)].mean(dim=0)
        )
    target = torch.stack(mean_frame_embeddings, dim=0).detach()
    return F.mse_loss(outputs["mol_embeddings"], target)


def mask_electronic_inputs(
    node_feats: torch.Tensor,
    trajectory_specs: list[tuple[int, int]],
    probability: float,
) -> tuple[torch.Tensor, torch.Tensor]:
    """Mask charge-related node channels consistently across all frames of an atom."""
    total_atoms = sum(int(num_atoms) for _, num_atoms in trajectory_specs)
    atom_mask = torch.rand(total_atoms, device=node_feats.device) < float(probability)
    if total_atoms > 0 and not atom_mask.any():
        atom_mask[torch.randint(total_atoms, (1,), device=node_feats.device)] = True

    frame_masks = []
    atom_start = 0
    for num_frames, num_atoms in trajectory_specs:
        molecule_mask = atom_mask[atom_start : atom_start + int(num_atoms)]
        frame_masks.append(molecule_mask.repeat(int(num_frames)))
        atom_start += int(num_atoms)
    frame_mask = torch.cat(frame_masks, dim=0)

    masked_feats = node_feats.clone()
    # Feature layout from comld.data.featurizer.atom_features followed by
    # appended [formal_charge, Gasteiger_charge]:
    # formal-charge one-hot = [20:26], base Gasteiger = 41, appended charges = [42:44].
    masked_feats[frame_mask, 20:26] = 0.0
    masked_feats[frame_mask, 41:44] = 0.0
    return masked_feats, atom_mask


def save_checkpoint(
    checkpoint_dir: str | Path,
    epoch: int,
    model: DynamicSE3LiquidTeacher,
    optimizer: torch.optim.Optimizer,
    config: dict[str, Any],
    model_config: dict[str, Any],
    metrics: dict[str, float],
    scheduler: torch.optim.lr_scheduler.LRScheduler | None = None,
    scaler: torch.cuda.amp.GradScaler | None = None,
    keep_epoch_checkpoint: bool = False,
) -> None:
    checkpoint_dir = Path(checkpoint_dir)
    checkpoint_dir.mkdir(parents=True, exist_ok=True)
    payload = {
        "epoch": int(epoch),
        "encoder_type": "dynamic_se3_liquid_teacher",
        "model_state_dict": model.state_dict(),
        "optimizer_state_dict": optimizer.state_dict(),
        "model_config": model_config,
        "config": config,
        "metrics": metrics,
    }
    if scheduler is not None:
        payload["scheduler_state_dict"] = scheduler.state_dict()
    if scaler is not None:
        payload["scaler_state_dict"] = scaler.state_dict()
    latest_path = checkpoint_dir / "latest.pt"
    temporary_path = checkpoint_dir / "latest.pt.tmp"
    torch.save(payload, temporary_path)
    temporary_path.replace(latest_path)
    if keep_epoch_checkpoint:
        torch.save(payload, checkpoint_dir / f"epoch_{epoch:03d}.pt")


def load_training_checkpoint(path: str | Path | None) -> dict[str, Any] | None:
    if not path:
        return None
    path = Path(path)
    if not path.exists():
        return None
    try:
        return torch.load(path, map_location="cpu", weights_only=False)
    except TypeError:
        return torch.load(path, map_location="cpu")


def cosine_with_warmup(step: int, warmup_steps: int, total_steps: int) -> float:
    if step < warmup_steps:
        return float(step + 1) / max(warmup_steps, 1)
    progress = (step - warmup_steps) / max(total_steps - warmup_steps, 1)
    return 0.5 * (1.0 + math.cos(math.pi * min(max(progress, 0.0), 1.0)))


def train(config: dict[str, Any]) -> None:
    paths = config.get("paths", {})
    data_cfg = config.get("data", {})
    model_cfg = config.get("model", {})
    noise_cfg = config.get("noise", {})
    loss_cfg = config.get("loss", {})
    train_cfg = config.get("train", {})

    set_seed(int(train_cfg.get("seed", 2026)))
    device = resolve_device(train_cfg.get("device", "auto"))
    dataset = build_dynamic_md_dataset(
        path=paths["dynamic_md_pt"],
        max_molecules=data_cfg.get("max_molecules"),
        max_frames=data_cfg.get("max_frames"),
        frame_stride=int(data_cfg.get("frame_stride", 1)),
        random_frame_crop=bool(data_cfg.get("random_frame_crop", True)),
        seed=int(train_cfg.get("seed", 2026)),
    )
    loader = DataLoader(
        dataset,
        batch_size=int(data_cfg.get("batch_size", 4)),
        shuffle=not isinstance(dataset, IterableDataset),
        num_workers=int(data_cfg.get("num_workers", 0)),
        collate_fn=collate_dynamic_md,
        pin_memory=device.type == "cuda",
    )

    sample = dynamic_md_sample(dataset)
    model_config = {
        "node_feat_dim": int(sample["node_feats"].shape[-1]),
        "edge_feat_dim": int(sample["edge_attr"].shape[-1]),
        "output_dim": int(model_cfg.get("output_dim", 384)),
        "hidden_dim": int(model_cfg.get("hidden_dim", 384)),
        "liquid_hidden_dim": int(model_cfg.get("liquid_hidden_dim", 384)),
        "num_layers": int(model_cfg.get("num_layers", 4)),
        "message_hidden_dim": int(model_cfg.get("message_hidden_dim", 256)),
        "num_heads": int(model_cfg.get("num_heads", 8)),
        "rbf_dim": int(model_cfg.get("rbf_dim", 16)),
        "radial_cutoff": float(model_cfg.get("radial_cutoff", 10.0)),
        "coord_update_scale": float(model_cfg.get("coord_update_scale", 0.1)),
        "dropout": float(model_cfg.get("dropout", 0.1)),
        "electronic_dim": int(model_cfg.get("electronic_dim", 2)),
    }
    model = DynamicSE3LiquidTeacher(**model_config).to(device)
    optimizer = torch.optim.AdamW(
        model.parameters(),
        lr=float(train_cfg.get("lr", 3e-4)),
        weight_decay=float(train_cfg.get("weight_decay", 1e-2)),
    )

    epochs = int(train_cfg.get("epochs", 10))
    accumulation_steps = max(int(train_cfg.get("gradient_accumulation_steps", 1)), 1)
    updates_per_epoch = math.ceil(
        math.ceil(len(dataset) / int(data_cfg.get("batch_size", 4))) / accumulation_steps
    )
    total_updates = max(updates_per_epoch * epochs, 1)
    warmup_steps = int(train_cfg.get("warmup_steps", max(int(total_updates * 0.05), 1)))
    scheduler = torch.optim.lr_scheduler.LambdaLR(
        optimizer,
        lr_lambda=lambda step: cosine_with_warmup(step, warmup_steps, total_updates),
    )
    amp_enabled = bool(train_cfg.get("amp", True)) and device.type == "cuda"
    scaler = torch.cuda.amp.GradScaler(enabled=amp_enabled)
    resume_path = train_cfg.get(
        "resume_checkpoint",
        str(Path(paths.get("checkpoint_dir", "checkpoints/pretrain_zinc_dynamic_3d_liquid")) / "latest.pt"),
    )
    resume = load_training_checkpoint(resume_path)
    start_epoch = 1
    if resume is not None:
        if resume.get("encoder_type") != "dynamic_se3_liquid_teacher":
            raise ValueError(f"Invalid resume checkpoint type in {resume_path}")
        model.load_state_dict(resume["model_state_dict"], strict=True)
        optimizer.load_state_dict(resume["optimizer_state_dict"])
        if "scheduler_state_dict" in resume:
            scheduler.load_state_dict(resume["scheduler_state_dict"])
        if "scaler_state_dict" in resume:
            scaler.load_state_dict(resume["scaler_state_dict"])
        start_epoch = int(resume["epoch"]) + 1
        print(f"Resuming 3D teacher from {resume_path} at epoch {start_epoch}")
    log_every = int(train_cfg.get("log_every", 20))
    save_every = int(train_cfg.get("save_every", 1))
    grad_clip = float(train_cfg.get("grad_clip", 5.0))
    extra_cgn_sigma = float(noise_cfg.get("extra_cgn_sigma", 0.01))
    electronic_mask_probability = float(
        noise_cfg.get("electronic_mask_probability", 0.15)
    )

    print(f"Device: {device}")
    print(f"Dynamic MD dataset: {len(dataset)} molecules from {paths['dynamic_md_pt']}")
    if isinstance(dataset, ShardedDynamicMDTrajectoryDataset):
        print(f"Streaming {len(dataset.shards)} trajectory shards (one shard in memory at a time)")
    print("3D teacher: SE(3)-Transformer + atom-level Liquid Time-Constant Network")
    print("Input: OpenMM MD frames + CAN + CGN + formal/Gasteiger charge")
    print(
        f"Optimization: amp={amp_enabled} accumulation={accumulation_steps} "
        f"warmup_steps={warmup_steps} total_updates≈{total_updates}"
    )

    for epoch in range(start_epoch, epochs + 1):
        if isinstance(dataset, ShardedDynamicMDTrajectoryDataset):
            dataset.set_epoch(epoch)
        model.train()
        totals = {
            "total": 0.0,
            "denoise": 0.0,
            "distance": 0.0,
            "charge": 0.0,
            "temporal": 0.0,
            "consistency": 0.0,
        }
        steps = 0
        optimizer.zero_grad(set_to_none=True)
        progress = tqdm(loader, desc=f"Dynamic 3D epoch {epoch}/{epochs}")
        for step, batch in enumerate(progress, start=1):
            batch = move_batch(batch, device)
            clean_coords = batch["coords"]
            noisy_coords = clean_coords + torch.randn_like(clean_coords) * extra_cgn_sigma
            masked_node_feats, charge_atom_mask = mask_electronic_inputs(
                batch["node_feats"],
                batch["trajectory_specs"],
                probability=electronic_mask_probability,
            )
            with torch.autocast(device_type=device.type, dtype=torch.float16, enabled=amp_enabled):
                outputs = model(
                    node_feats=masked_node_feats,
                    coords=noisy_coords,
                    edge_index=batch["edge_index"],
                    edge_attr=batch["edge_attr"],
                    frame_batch=batch["frame_batch"],
                    atom_batch=batch["atom_batch"],
                    trajectory_specs=batch["trajectory_specs"],
                    delta_times_ps=batch["delta_times_ps"],
                )

                denoise = F.mse_loss(outputs["denoised_coords"], clean_coords)
                distance = edge_distance_loss(
                    outputs["denoised_coords"],
                    clean_coords,
                    batch["edge_index"],
                )
                charge = F.mse_loss(
                    outputs["charge_pred"][charge_atom_mask],
                    batch["electronic_feats"][charge_atom_mask],
                )
                temporal = temporal_prediction_loss(
                    model,
                    outputs["frame_mol_hidden"],
                    batch["trajectory_frame_ptr"],
                )
                consistency = trajectory_consistency_loss(
                    outputs,
                    batch["trajectory_frame_ptr"],
                )
                total = (
                    float(loss_cfg.get("lambda_denoise", 1.0)) * denoise
                    + float(loss_cfg.get("lambda_distance", 0.1)) * distance
                    + float(loss_cfg.get("lambda_charge", 0.25)) * charge
                    + float(loss_cfg.get("lambda_temporal", 0.5)) * temporal
                    + float(loss_cfg.get("lambda_consistency", 0.25)) * consistency
                )

            scaler.scale(total / accumulation_steps).backward()
            should_update = step % accumulation_steps == 0
            if should_update:
                scaler.unscale_(optimizer)
                if grad_clip > 0:
                    torch.nn.utils.clip_grad_norm_(model.parameters(), grad_clip)
                scaler.step(optimizer)
                scaler.update()
                scheduler.step()
                optimizer.zero_grad(set_to_none=True)

            steps += 1
            values = {
                "total": total,
                "denoise": denoise,
                "distance": distance,
                "charge": charge,
                "temporal": temporal,
                "consistency": consistency,
            }
            for key, value in values.items():
                totals[key] += float(value.detach().item())
            if step % log_every == 0:
                progress.set_postfix(
                    total=totals["total"] / steps,
                    denoise=totals["denoise"] / steps,
                    charge=totals["charge"] / steps,
                    temporal=totals["temporal"] / steps,
                    lr=optimizer.param_groups[0]["lr"],
                )

        if steps % accumulation_steps != 0:
            scaler.unscale_(optimizer)
            if grad_clip > 0:
                torch.nn.utils.clip_grad_norm_(model.parameters(), grad_clip)
            scaler.step(optimizer)
            scaler.update()
            scheduler.step()
            optimizer.zero_grad(set_to_none=True)

        metrics = {key: value / max(steps, 1) for key, value in totals.items()}
        print(
            f"Epoch {epoch}: total={metrics['total']:.6f} "
            f"denoise={metrics['denoise']:.6f} distance={metrics['distance']:.6f} "
            f"charge={metrics['charge']:.6f} temporal={metrics['temporal']:.6f} "
            f"consistency={metrics['consistency']:.6f}"
        )
        if save_every > 0 and epoch % save_every == 0:
            save_checkpoint(
                checkpoint_dir=paths.get(
                    "checkpoint_dir",
                    "checkpoints/pretrain_zinc_dynamic_3d_liquid",
                ),
                epoch=epoch,
                model=model,
                optimizer=optimizer,
                config=config,
                model_config=model_config,
                metrics=metrics,
                scheduler=scheduler,
                scaler=scaler,
                keep_epoch_checkpoint=bool(train_cfg.get("keep_epoch_checkpoints", False)),
            )


def build_arg_parser() -> argparse.ArgumentParser:
    parser = argparse.ArgumentParser(
        description="Pretrain a dynamic SE(3)-Transformer + Liquid 3D teacher."
    )
    parser.add_argument("--config", default="configs/pretrain/pretrain_dynamic_3d_zinc.yaml")
    return parser


def main() -> None:
    args = build_arg_parser().parse_args()
    train(load_config(args.config))


if __name__ == "__main__":
    main()
```
