## Supplementary material for "PG-MLD: Physics-Guided Molecular Representation Learning via Dynamic 3D Trajectory Distillation": Code and Data: pretrain_dynamic_3d_to_1d.py

```
from __future__ import annotations

import argparse
import math
import random
from pathlib import Path
from typing import Any

import numpy as np
import torch
import torch.nn.functional as F
import yaml
from torch.utils.data import DataLoader, IterableDataset
from tqdm import tqdm

from comld.data.dynamic_md_dataset import (
    ShardedDynamicMDTrajectoryDataset,
    build_dynamic_md_dataset,
    collate_dynamic_md,
    dynamic_md_sample,
)
from comld.models.chemberta_teacher import ChemBERTaDistillationStudent
from comld.models.dynamic_se3_liquid import DynamicSE3LiquidTeacher


def load_config(path: str | Path) -> dict[str, Any]:
    with open(path, "r", encoding="utf-8") as handle:
        return yaml.safe_load(handle) or {}


def set_seed(seed: int) -> None:
    random.seed(seed)
    np.random.seed(seed)
    torch.manual_seed(seed)
    if torch.cuda.is_available():
        torch.cuda.manual_seed_all(seed)


def resolve_device(device_name: str) -> torch.device:
    if device_name == "auto":
        return torch.device("cuda" if torch.cuda.is_available() else "cpu")
    return torch.device(device_name)


def load_checkpoint(path: str | Path) -> dict[str, Any]:
    path = Path(path)
    if not path.exists():
        raise FileNotFoundError(f"Dynamic 3D teacher checkpoint not found: {path}")
    try:
        return torch.load(path, map_location="cpu", weights_only=False)
    except TypeError:
        return torch.load(path, map_location="cpu")


def build_dynamic_teacher(
    checkpoint: dict[str, Any],
    sample: dict[str, Any],
    device: torch.device,
) -> DynamicSE3LiquidTeacher:
    if checkpoint.get("encoder_type") != "dynamic_se3_liquid_teacher":
        raise ValueError(
            "Expected a dynamic_se3_liquid_teacher checkpoint. "
            "Run comld.train.pretrain_dynamic_3d first."
        )
    model_config = dict(checkpoint["model_config"])
    expected_node_dim = int(sample["node_feats"].shape[-1])
    expected_edge_dim = int(sample["edge_attr"].shape[-1])
    if int(model_config["node_feat_dim"]) != expected_node_dim:
        raise ValueError(
            f"Teacher node feature dim={model_config['node_feat_dim']} "
            f"but dataset dim={expected_node_dim}"
        )
    if int(model_config["edge_feat_dim"]) != expected_edge_dim:
        raise ValueError(
            f"Teacher edge feature dim={model_config['edge_feat_dim']} "
            f"but dataset dim={expected_edge_dim}"
        )
    teacher = DynamicSE3LiquidTeacher(**model_config)
    teacher.load_state_dict(checkpoint["model_state_dict"], strict=True)
    teacher.to(device)
    teacher.eval()
    for parameter in teacher.parameters():
        parameter.requires_grad_(False)
    return teacher


def move_batch(batch: dict[str, Any], device: torch.device) -> dict[str, Any]:
    moved = dict(batch)
    for key in [
        "node_feats",
        "coords",
        "edge_index",
        "edge_attr",
        "frame_batch",
        "atom_batch",
        "electronic_feats",
        "atom_counts",
        "trajectory_frame_ptr",
        "delta_times_ps",
    ]:
        moved[key] = batch[key].to(device)
    return moved


def symmetric_contrastive_loss(
    student_embeddings: torch.Tensor,
    teacher_embeddings: torch.Tensor,
    temperature: float,
) -> torch.Tensor:
    if student_embeddings.shape[0] <= 1:
        return student_embeddings.new_tensor(0.0)
    student_norm = F.normalize(student_embeddings, dim=-1)
    teacher_norm = F.normalize(teacher_embeddings.detach(), dim=-1)
    logits = student_norm @ teacher_norm.t() / max(float(temperature), 1e-6)
    labels = torch.arange(logits.shape[0], device=logits.device)
    return 0.5 * (
        F.cross_entropy(logits, labels)
        + F.cross_entropy(logits.t(), labels)
    )


def save_checkpoint(
    checkpoint_dir: str | Path,
    epoch: int,
    student: ChemBERTaDistillationStudent,
    optimizer: torch.optim.Optimizer,
    config: dict[str, Any],
    metrics: dict[str, float],
    scheduler: torch.optim.lr_scheduler.LRScheduler | None = None,
    scaler: torch.cuda.amp.GradScaler | None = None,
    keep_epoch_checkpoint: bool = False,
) -> None:
    checkpoint_dir = Path(checkpoint_dir)
    checkpoint_dir.mkdir(parents=True, exist_ok=True)
    payload = {
        "epoch": int(epoch),
        "encoder_type": "chemberta_3d_to_1d",
        "distillation_source": "dynamic_se3_liquid_teacher",
        "student_model_name": student.model_name,
        "student_model_state_dict": student.state_dict(),
        "student_state_dict": student.encoder_state_dict(),
        "optimizer_state_dict": optimizer.state_dict(),
        "config": config,
        "metrics": metrics,
    }
    if scheduler is not None:
        payload["scheduler_state_dict"] = scheduler.state_dict()
    if scaler is not None:
        payload["scaler_state_dict"] = scaler.state_dict()
    latest_path = checkpoint_dir / "latest.pt"
    temporary_path = checkpoint_dir / "latest.pt.tmp"
    torch.save(payload, temporary_path)
    temporary_path.replace(latest_path)
    if keep_epoch_checkpoint:
        torch.save(payload, checkpoint_dir / f"epoch_{epoch:03d}.pt")


def cosine_with_warmup(step: int, warmup_steps: int, total_steps: int) -> float:
    if step < warmup_steps:
        return float(step + 1) / max(warmup_steps, 1)
    progress = (step - warmup_steps) / max(total_steps - warmup_steps, 1)
    return 0.5 * (1.0 + math.cos(math.pi * min(max(progress, 0.0), 1.0)))


def train(config: dict[str, Any]) -> None:
    paths = config.get("paths", {})
    data_cfg = config.get("data", {})
    student_cfg = config.get("student_1d", {})
    loss_cfg = config.get("loss", {})
    train_cfg = config.get("train", {})

    set_seed(int(train_cfg.get("seed", 2026)))
    device = resolve_device(train_cfg.get("device", "auto"))
    dataset = build_dynamic_md_dataset(
        path=paths["dynamic_md_pt"],
        max_molecules=data_cfg.get("max_molecules"),
        max_frames=data_cfg.get("max_frames"),
        frame_stride=int(data_cfg.get("frame_stride", 1)),
        random_frame_crop=bool(data_cfg.get("random_frame_crop", True)),
        seed=int(train_cfg.get("seed", 2026)),
    )
    loader = DataLoader(
        dataset,
        batch_size=int(data_cfg.get("batch_size", 4)),
        shuffle=not isinstance(dataset, IterableDataset),
        num_workers=int(data_cfg.get("num_workers", 0)),
        collate_fn=collate_dynamic_md,
        pin_memory=device.type == "cuda",
    )

    teacher_checkpoint = load_checkpoint(paths["teacher_dynamic_checkpoint"])
    teacher = build_dynamic_teacher(teacher_checkpoint, dynamic_md_sample(dataset), device)
    student = ChemBERTaDistillationStudent(
        model_name=student_cfg.get(
            "model_name",
            "comld/models/DeepChemChemBERTa-77M-MLM",
        ),
    ).to(device)
    if teacher_checkpoint["model_config"]["output_dim"] != student.hidden_size:
        raise ValueError(
            f"3D teacher output dim={teacher_checkpoint['model_config']['output_dim']} "
            f"must equal ChemBERTa hidden size={student.hidden_size}"
        )

    optimizer = torch.optim.AdamW(
        student.parameters(),
        lr=float(train_cfg.get("lr", 2e-5)),
        weight_decay=float(train_cfg.get("weight_decay", 1e-2)),
    )
    epochs = int(train_cfg.get("epochs", 5))
    accumulation_steps = max(int(train_cfg.get("gradient_accumulation_steps", 1)), 1)
    updates_per_epoch = math.ceil(
        math.ceil(len(dataset) / int(data_cfg.get("batch_size", 4))) / accumulation_steps
    )
    total_updates = max(updates_per_epoch * epochs, 1)
    warmup_steps = int(train_cfg.get("warmup_steps", max(int(total_updates * 0.05), 1)))
    scheduler = torch.optim.lr_scheduler.LambdaLR(
        optimizer,
        lr_lambda=lambda step: cosine_with_warmup(step, warmup_steps, total_updates),
    )
    amp_enabled = bool(train_cfg.get("amp", True)) and device.type == "cuda"
    scaler = torch.cuda.amp.GradScaler(enabled=amp_enabled)
    resume_path = train_cfg.get(
        "resume_checkpoint",
        str(Path(paths.get("checkpoint_dir", "checkpoints/pretrain_zinc_dynamic_3d_to_1d")) / "latest.pt"),
    )
    resume = load_checkpoint(resume_path) if Path(resume_path).exists() else None
    start_epoch = 1
    if resume is not None:
        if resume.get("encoder_type") != "chemberta_3d_to_1d":
            raise ValueError(f"Invalid resume checkpoint type in {resume_path}")
        if "student_model_state_dict" in resume:
            student.load_state_dict(resume["student_model_state_dict"], strict=True)
        else:
            student.base_model.load_state_dict(resume["student_state_dict"], strict=True)
        optimizer.load_state_dict(resume["optimizer_state_dict"])
        if "scheduler_state_dict" in resume:
            scheduler.load_state_dict(resume["scheduler_state_dict"])
        if "scaler_state_dict" in resume:
            scaler.load_state_dict(resume["scaler_state_dict"])
        start_epoch = int(resume["epoch"]) + 1
        print(f"Resuming 3D-to-1D distillation from {resume_path} at epoch {start_epoch}")
    log_every = int(train_cfg.get("log_every", 20))
    save_every = int(train_cfg.get("save_every", 1))
    grad_clip = float(train_cfg.get("grad_clip", 1.0))
    lambda_atom = float(loss_cfg.get("lambda_atom", 1.0))
    lambda_mol = float(loss_cfg.get("lambda_mol", 1.0))
    lambda_contrastive = float(loss_cfg.get("lambda_contrastive", 0.1))
    lambda_mlm = float(loss_cfg.get("lambda_mlm", 0.1))
    mlm_probability = float(loss_cfg.get("mlm_probability", 0.15))
    temperature = float(loss_cfg.get("contrastive_temperature", 0.1))

    print(f"Device: {device}")
    print(f"Dynamic MD dataset: {len(dataset)} molecules from {paths['dynamic_md_pt']}")
    if isinstance(dataset, ShardedDynamicMDTrajectoryDataset):
        print(f"Streaming {len(dataset.shards)} trajectory shards (one shard in memory at a time)")
    print(f"3D teacher: {paths['teacher_dynamic_checkpoint']}")
    print(f"1D student: {student.model_name}")
    print("Direction: dynamic 3D (SE(3)+Liquid) -> 1D ChemBERTa")
    print(
        f"Optimization: amp={amp_enabled} accumulation={accumulation_steps} "
        f"warmup_steps={warmup_steps} total_updates≈{total_updates}"
    )

    for epoch in range(start_epoch, epochs + 1):
        if isinstance(dataset, ShardedDynamicMDTrajectoryDataset):
            dataset.set_epoch(epoch)
        student.train()
        totals = {
            "total": 0.0,
            "atom": 0.0,
            "mol": 0.0,
            "contrastive": 0.0,
            "mlm": 0.0,
        }
        steps = 0
        optimizer.zero_grad(set_to_none=True)
        progress = tqdm(loader, desc=f"Dynamic 3D->1D epoch {epoch}/{epochs}")
        for step, batch in enumerate(progress, start=1):
            batch = move_batch(batch, device)
            with torch.no_grad(), torch.autocast(
                device_type=device.type,
                dtype=torch.float16,
                enabled=amp_enabled,
            ):
                teacher_out = teacher(
                    node_feats=batch["node_feats"],
                    coords=batch["coords"],
                    edge_index=batch["edge_index"],
                    edge_attr=batch["edge_attr"],
                    frame_batch=batch["frame_batch"],
                    atom_batch=batch["atom_batch"],
                    trajectory_specs=batch["trajectory_specs"],
                    delta_times_ps=batch["delta_times_ps"],
                )
            with torch.autocast(device_type=device.type, dtype=torch.float16, enabled=amp_enabled):
                student_out = student(
                    smiles=batch["smiles"],
                    atom_counts=batch["atom_counts"],
                    device=device,
                    mlm_probability=mlm_probability,
                )

                atom_loss = F.mse_loss(
                    student_out["atom_embeddings"],
                    teacher_out["atom_embeddings"].detach(),
                )
                mol_loss = F.mse_loss(
                    student_out["mol_embeddings"],
                    teacher_out["mol_embeddings"].detach(),
                )
                contrastive = symmetric_contrastive_loss(
                    student_out["mol_embeddings"],
                    teacher_out["mol_embeddings"],
                    temperature=temperature,
                )
                total = (
                    lambda_atom * atom_loss
                    + lambda_mol * mol_loss
                    + lambda_contrastive * contrastive
                    + lambda_mlm * student_out["mlm_loss"]
                )

            scaler.scale(total / accumulation_steps).backward()
            should_update = step % accumulation_steps == 0
            if should_update:
                scaler.unscale_(optimizer)
                if grad_clip > 0:
                    torch.nn.utils.clip_grad_norm_(student.parameters(), grad_clip)
                scaler.step(optimizer)
                scaler.update()
                scheduler.step()
                optimizer.zero_grad(set_to_none=True)

            steps += 1
            values = {
                "total": total,
                "atom": atom_loss,
                "mol": mol_loss,
                "contrastive": contrastive,
                "mlm": student_out["mlm_loss"],
            }
            for key, value in values.items():
                totals[key] += float(value.detach().item())
            if step % log_every == 0:
                progress.set_postfix(
                    total=totals["total"] / steps,
                    atom=totals["atom"] / steps,
                    mol=totals["mol"] / steps,
                    ctr=totals["contrastive"] / steps,
                    mlm=totals["mlm"] / steps,
                    lr=optimizer.param_groups[0]["lr"],
                )

        if steps % accumulation_steps != 0:
            scaler.unscale_(optimizer)
            if grad_clip > 0:
                torch.nn.utils.clip_grad_norm_(student.parameters(), grad_clip)
            scaler.step(optimizer)
            scaler.update()
            scheduler.step()
            optimizer.zero_grad(set_to_none=True)

        metrics = {key: value / max(steps, 1) for key, value in totals.items()}
        print(
            f"Epoch {epoch}: total={metrics['total']:.6f} "
            f"atom={metrics['atom']:.6f} mol={metrics['mol']:.6f} "
            f"contrastive={metrics['contrastive']:.6f} mlm={metrics['mlm']:.6f}"
        )
        if save_every > 0 and epoch % save_every == 0:
            save_checkpoint(
                checkpoint_dir=paths.get(
                    "checkpoint_dir",
                    "checkpoints/pretrain_zinc_dynamic_3d_to_1d",
                ),
                epoch=epoch,
                student=student,
                optimizer=optimizer,
                config=config,
                metrics=metrics,
                scheduler=scheduler,
                scaler=scaler,
                keep_epoch_checkpoint=bool(train_cfg.get("keep_epoch_checkpoints", False)),
            )


def build_arg_parser() -> argparse.ArgumentParser:
    parser = argparse.ArgumentParser(
        description="Distill a dynamic SE(3)+Liquid 3D teacher into ChemBERTa."
    )
    parser.add_argument(
        "--config",
        default="configs/pretrain/pretrain_dynamic_3d_to_1d_zinc.yaml",
    )
    return parser


def main() -> None:
    args = build_arg_parser().parse_args()
    train(load_config(args.config))


if __name__ == "__main__":
    main()
```
