## Supplementary material for "PG-MLD: Physics-Guided Molecular Representation Learning via Dynamic 3D Trajectory Distillation": Code and Data: pretrain_dynamic_3d_to_1d_chemblm.py

```
from __future__ import annotations

import argparse

from comld.train.pretrain_dynamic_3d_to_1d_molformer import load_config, train


def build_arg_parser() -> argparse.ArgumentParser:
    parser = argparse.ArgumentParser(description="Distill a dynamic SE(3)+Liquid 3D teacher into ChemBLM.")
    parser.add_argument("--config", default="configs/pretrain/pretrain_dynamic_3d_to_1d_zinc_chemblm.yaml")
    return parser


def main() -> None:
    args = build_arg_parser().parse_args()
    config = load_config(args.config)
    config.setdefault("student_1d", {})
    config["student_1d"]["student_type"] = "chemblm"
    train(config)


if __name__ == "__main__":
    main()
```
