## Supplementary material for "PG-MLD: Physics-Guided Molecular Representation Learning via Dynamic 3D Trajectory Distillation": Code and Data: pretrain_static_3d_ablation.py

```
from __future__ import annotations

import argparse
from pathlib import Path
from typing import Any

from comld.train import pretrain_dynamic_3d as dynamic_pretraining


DEFAULT_CONFIG = "configs/ablation/md_value/pretrain_no_md_static_3d_zinc.yaml"


def load_config(path: str | Path) -> dict[str, Any]:
    return dynamic_pretraining.load_config(path)


def train(config: dict[str, Any]) -> None:
    """Pretrain the teacher on separately preprocessed static 3D views."""

    ablation_cfg = config.setdefault("ablation", {})
    ablation_cfg.update(
        {
            "name": "md_trajectory_value",
            "condition": "no_md",
            "trajectory_mode": "equilibrium_plus_cgn_can",
            "preserve_frame_count": True,
        }
    )

    print("Ablation: NO MD trajectory (equilibrium + static CGN/CAN views)")
    print("Static data are preprocessed separately; no MD coordinates are loaded")
    dynamic_pretraining.train(config)


def build_arg_parser() -> argparse.ArgumentParser:
    parser = argparse.ArgumentParser(
        description="Pretrain the no-MD equilibrium+CGN/CAN 3D teacher ablation."
    )
    parser.add_argument("--config", default=DEFAULT_CONFIG)
    return parser


def main() -> None:
    args = build_arg_parser().parse_args()
    train(load_config(args.config))


if __name__ == "__main__":
    main()
```
