## Supplementary material for "PG-MLD: Physics-Guided Molecular Representation Learning via Dynamic 3D Trajectory Distillation": Code and Data: pretrain_static_3d_to_1d_chemblm.py

```
from __future__ import annotations

import argparse
from pathlib import Path
from typing import Any

from comld.train import pretrain_dynamic_3d_to_1d_molformer as distillation


DEFAULT_CONFIG = "configs/ablation/md_value/distill_no_md_3d_to_chemblm.yaml"


def load_config(path: str | Path) -> dict[str, Any]:
    return distillation.load_config(path)


def train(config: dict[str, Any]) -> None:
    """Distill the no-MD static 3D teacher into ChemBLM."""

    student_type = str(config.get("student_1d", {}).get("student_type", "")).lower()
    if student_type != "chemblm":
        raise ValueError(
            "pretrain_static_3d_to_1d_chemblm requires student_1d.student_type=chemblm"
        )

    ablation_cfg = config.setdefault("ablation", {})
    ablation_cfg.update(
        {
            "name": "md_trajectory_value",
            "condition": "no_md",
            "trajectory_mode": "equilibrium_plus_cgn_can",
            "distillation_target": "chemblm",
        }
    )

    print("Ablation distillation: static no-MD 3D teacher -> 1D ChemBLM")
    distillation.train(config)


def build_arg_parser() -> argparse.ArgumentParser:
    parser = argparse.ArgumentParser(
        description="Distill the no-MD equilibrium+CGN/CAN 3D teacher into ChemBLM."
    )
    parser.add_argument("--config", default=DEFAULT_CONFIG)
    return parser


def main() -> None:
    args = build_arg_parser().parse_args()
    train(load_config(args.config))


if __name__ == "__main__":
    main()
```
