## Supplementary material for "PG-MLD: Physics-Guided Molecular Representation Learning via Dynamic 3D Trajectory Distillation": Code and Data: splits.py

```
from __future__ import annotations

from pathlib import Path
from typing import Sequence

import numpy as np
import pandas as pd


def make_scaffold(smiles: str, include_chirality: bool = False) -> str:
    from rdkit import Chem
    from rdkit.Chem.Scaffolds import MurckoScaffold

    mol = Chem.MolFromSmiles(smiles)
    if mol is None:
        return ""
    return MurckoScaffold.MurckoScaffoldSmiles(mol=mol, includeChirality=include_chirality)


def random_split_indices(
    n_items: int,
    train_frac: float,
    val_frac: float,
    seed: int,
) -> tuple[list[int], list[int], list[int]]:
    rng = np.random.default_rng(seed)
    indices = rng.permutation(n_items).tolist()
    n_train = int(round(n_items * train_frac))
    n_val = int(round(n_items * val_frac))
    n_train = min(max(n_train, 1), n_items)
    n_val = min(max(n_val, 1), max(n_items - n_train, 0))
    train_idx = indices[:n_train]
    val_idx = indices[n_train : n_train + n_val]
    test_idx = indices[n_train + n_val :]
    if not test_idx and val_idx:
        test_idx = [val_idx.pop()]
    return train_idx, val_idx, test_idx


def predefined_split_indices(
    split_dir: str | Path,
    source_indices: Sequence[int] | None = None,
) -> tuple[list[int], list[int], list[int]]:
    split_dir = Path(split_dir)
    split_files = {
        "train": split_dir / "train.csv.gz",
        "val": split_dir / "valid.csv.gz",
        "test": split_dir / "test.csv.gz",
    }
    missing = [str(path) for path in split_files.values() if not path.exists()]
    if missing:
        raise FileNotFoundError(f"Missing predefined split files: {missing}")

    raw_splits = {
        name: pd.read_csv(path, compression="gzip", header=None).iloc[:, 0].astype(int).tolist()
        for name, path in split_files.items()
    }
    split_sets = {name: set(indices) for name, indices in raw_splits.items()}
    if split_sets["train"] & split_sets["val"]:
        raise ValueError("Predefined train and validation splits overlap.")
    if split_sets["train"] & split_sets["test"]:
        raise ValueError("Predefined train and test splits overlap.")
    if split_sets["val"] & split_sets["test"]:
        raise ValueError("Predefined validation and test splits overlap.")

    if source_indices is None:
        return raw_splits["train"], raw_splits["val"], raw_splits["test"]

    source_to_dataset = {int(source_idx): dataset_idx for dataset_idx, source_idx in enumerate(source_indices)}
    raw_union = split_sets["train"] | split_sets["val"] | split_sets["test"]
    missing_dataset_rows = set(source_to_dataset) - raw_union
    if missing_dataset_rows:
        preview = sorted(missing_dataset_rows)[:10]
        raise ValueError(
            "Dataset rows are not covered by the predefined split indices. "
            f"First missing source indices: {preview}"
        )

    mapped = {
        name: [source_to_dataset[idx] for idx in indices if idx in source_to_dataset]
        for name, indices in raw_splits.items()
    }
    if any(not indices for indices in mapped.values()):
        raise ValueError("A predefined split became empty after mapping dataset rows.")
    return mapped["train"], mapped["val"], mapped["test"]


def _as_2d_float(values: Sequence[Sequence[float]] | np.ndarray | None) -> np.ndarray | None:
    if values is None:
        return None
    array = np.asarray(values, dtype=float)
    if array.ndim == 1:
        array = array.reshape(-1, 1)
    return array


def _flatten(groups: list[list[int]]) -> list[int]:
    return [idx for group in groups for idx in group]


def _build_split_from_group_order(
    scaffold_sets: list[list[int]],
    train_target: int,
    val_target: int,
) -> tuple[list[int], list[int], list[int]]:
    train_groups: list[list[int]] = []
    val_groups: list[list[int]] = []
    test_groups: list[list[int]] = []
    train_size = 0
    val_size = 0

    for group in scaffold_sets:
        if train_size + len(group) <= train_target or not train_groups:
            train_groups.append(group)
            train_size += len(group)
        elif val_size + len(group) <= val_target or not val_groups:
            val_groups.append(group)
            val_size += len(group)
        else:
            test_groups.append(group)

    if not val_groups and test_groups:
        val_groups.append(test_groups.pop(0))
    if not test_groups and val_groups:
        test_groups.append(val_groups.pop())
    if not train_groups and val_groups:
        train_groups.append(val_groups.pop())
    return _flatten(train_groups), _flatten(val_groups), _flatten(test_groups)


def _valid_class_tasks(labels: np.ndarray, masks: np.ndarray, indices: Sequence[int]) -> list[int]:
    if len(indices) == 0:
        return []
    split_labels = labels[list(indices)]
    split_masks = masks[list(indices)].astype(bool)
    valid_tasks: list[int] = []
    for task_idx in range(split_labels.shape[1]):
        task_valid = split_masks[:, task_idx] & np.isin(split_labels[:, task_idx], [0.0, 1.0])
        if task_valid.sum() == 0:
            continue
        unique = np.unique(split_labels[:, task_idx][task_valid].astype(int))
        if len(unique) >= 2:
            valid_tasks.append(task_idx)
    return valid_tasks


def _classification_balance_error(
    labels: np.ndarray,
    masks: np.ndarray,
    split_indices: Sequence[int],
    global_pos_frac: dict[int, float],
) -> float:
    if len(split_indices) == 0:
        return 10.0
    split_labels = labels[list(split_indices)]
    split_masks = masks[list(split_indices)].astype(bool)
    error = 0.0
    counted = 0
    for task_idx, global_frac in global_pos_frac.items():
        task_valid = split_masks[:, task_idx] & np.isin(split_labels[:, task_idx], [0.0, 1.0])
        if task_valid.sum() == 0:
            error += 2.0
            counted += 1
            continue
        split_frac = float(split_labels[:, task_idx][task_valid].mean())
        error += abs(split_frac - global_frac)
        counted += 1
    return error / max(counted, 1)


def _score_split(
    train_idx: list[int],
    val_idx: list[int],
    test_idx: list[int],
    targets: tuple[int, int, int],
    labels: np.ndarray | None,
    masks: np.ndarray | None,
    task_type: str | None,
) -> float:
    splits = [train_idx, val_idx, test_idx]
    size_error = sum(abs(len(indices) - target) / max(target, 1) for indices, target in zip(splits, targets))
    if task_type != "classification" or labels is None or masks is None:
        return size_error

    global_valid_tasks = _valid_class_tasks(labels, masks, range(labels.shape[0]))
    if not global_valid_tasks:
        return size_error + 1000.0

    global_pos_frac: dict[int, float] = {}
    for task_idx in global_valid_tasks:
        task_valid = masks[:, task_idx].astype(bool) & np.isin(labels[:, task_idx], [0.0, 1.0])
        global_pos_frac[task_idx] = float(labels[:, task_idx][task_valid].mean())

    coverage_penalty = 0.0
    for indices in [val_idx, test_idx]:
        if not _valid_class_tasks(labels, masks, indices):
            coverage_penalty += 1000.0

    balance_error = sum(
        _classification_balance_error(labels, masks, indices, global_pos_frac)
        for indices in [train_idx, val_idx, test_idx]
    )
    return size_error + coverage_penalty + balance_error


def scaffold_split_indices(
    smiles: Sequence[str],
    train_frac: float,
    val_frac: float,
    seed: int,
    task_type: str | None = None,
    labels: Sequence[Sequence[float]] | np.ndarray | None = None,
    masks: Sequence[Sequence[float]] | np.ndarray | None = None,
    max_attempts: int = 256,
) -> tuple[list[int], list[int], list[int]]:
    scaffold_to_indices: dict[str, list[int]] = {}
    for idx, smiles_value in enumerate(smiles):
        scaffold = make_scaffold(str(smiles_value)) or f"__empty_scaffold_{idx}"
        scaffold_to_indices.setdefault(scaffold, []).append(idx)

    scaffold_sets = sorted(scaffold_to_indices.values(), key=lambda indices: (-len(indices), indices[0]))
    n_items = len(smiles)
    train_target = int(round(n_items * train_frac))
    val_target = int(round(n_items * val_frac))
    test_target = max(n_items - train_target - val_target, 1)
    targets = (train_target, val_target, test_target)

    label_array = _as_2d_float(labels)
    mask_array = _as_2d_float(masks)
    if label_array is not None and mask_array is None:
        mask_array = np.ones_like(label_array, dtype=float)

    candidates: list[tuple[list[int], list[int], list[int]]] = [
        _build_split_from_group_order(scaffold_sets, train_target, val_target)
    ]
    if task_type == "classification" and len(scaffold_sets) > 3:
        rng = np.random.default_rng(seed)
        for _ in range(max_attempts):
            order = [list(group) for group in scaffold_sets]
            rng.shuffle(order)
            candidates.append(_build_split_from_group_order(order, train_target, val_target))

    return min(
        candidates,
        key=lambda candidate: _score_split(*candidate, targets, label_array, mask_array, task_type),
    )


def random_scaffold_split_indices(
    smiles: Sequence[str],
    train_frac: float,
    val_frac: float,
    seed: int,
) -> tuple[list[int], list[int], list[int]]:
    scaffold_to_indices: dict[str, list[int]] = {}
    for idx, smiles_value in enumerate(smiles):
        scaffold = make_scaffold(str(smiles_value)) or f"__empty_scaffold_{idx}"
        scaffold_to_indices.setdefault(scaffold, []).append(idx)

    scaffold_sets = [list(indices) for indices in scaffold_to_indices.values()]
    rng = np.random.default_rng(seed)
    rng.shuffle(scaffold_sets)

    n_items = len(smiles)
    train_target = int(round(n_items * train_frac))
    val_target = int(round(n_items * val_frac))
    return _build_split_from_group_order(scaffold_sets, train_target, val_target)


def classification_split_summary(
    labels: Sequence[Sequence[float]] | np.ndarray,
    masks: Sequence[Sequence[float]] | np.ndarray,
    train_idx: Sequence[int],
    val_idx: Sequence[int],
    test_idx: Sequence[int],
) -> str:
    label_array = _as_2d_float(labels)
    mask_array = _as_2d_float(masks)
    if label_array is None or mask_array is None:
        return ""
    parts = []
    for name, indices in [("train", train_idx), ("val", val_idx), ("test", test_idx)]:
        valid_tasks = _valid_class_tasks(label_array, mask_array, indices)
        parts.append(f"{name}_valid_auc_tasks={len(valid_tasks)}")
    return " ".join(parts)
```
