## Supplementary material for "PG-MLD: Physics-Guided Molecular Representation Learning via Dynamic 3D Trajectory Distillation": Code and Data: distill_no_md_3d_to_chemblm.yaml

```
paths:
  dynamic_md_pt: zinc/zinc15_250K_static_cgn_can_electronic/shard_*/static_cgn_can_electronic.pt
  teacher_dynamic_checkpoint: checkpoints/ablation/md_value/no_md_3d_teacher/latest.pt
  checkpoint_dir: checkpoints/ablation/md_value/no_md_3d_to_1d_chemblm

data:
  batch_size: 32
  num_workers: 4
  max_molecules:
  max_frames: 8
  frame_stride: 1
  random_frame_crop: true

student_1d:
  student_type: chemblm
  model_name: comld/models/ChemBLM
  trust_remote_code: false
  use_safetensors: false
  use_projection: true

loss:
  lambda_atom: 1.0
  lambda_mol: 1.0
  lambda_contrastive: 0.05
  contrastive_temperature: 0.2
  lambda_mlm: 0.0
  mlm_probability: 0.0

train:
  epochs: 10
  lr: 0.00002
  weight_decay: 0.01
  grad_clip: 0.5
  gradient_accumulation_steps: 1
  warmup_steps: 2000
  amp: false
  keep_epoch_checkpoints: false
  log_every: 100
  save_every: 1
  device: cuda
  seed: 2026

ablation:
  name: md_trajectory_value
  condition: no_md
  trajectory_mode: equilibrium_plus_cgn_can
  distillation_target: chemblm
```
