## Supplementary material for "PG-MLD: Physics-Guided Molecular Representation Learning via Dynamic 3D Trajectory Distillation": Code and Data: pretrain_no_md_static_3d_zinc.yaml

```
paths:
  # Produced by scripts/build_no_md_static_dataset.py.
  dynamic_md_pt: zinc/zinc15_250K_static_cgn_can_electronic/shard_*/static_cgn_can_electronic.pt
  checkpoint_dir: checkpoints/ablation/md_value/no_md_3d_teacher

data:
  batch_size: 4
  num_workers: 2
  max_molecules:
  max_frames: 8
  frame_stride: 1
  # Matches the full-MD config. Preprocessed static samples already have T=8,
  # so this option performs no crop in the no-MD condition.
  random_frame_crop: true

model:
  output_dim: 384
  hidden_dim: 384
  liquid_hidden_dim: 384
  num_layers: 4
  message_hidden_dim: 256
  num_heads: 8
  rbf_dim: 16
  radial_cutoff: 10.0
  coord_update_scale: 0.1
  dropout: 0.1
  electronic_dim: 2

noise:
  extra_cgn_sigma: 0.01
  electronic_mask_probability: 0.15

loss:
  lambda_denoise: 1.0
  lambda_distance: 0.1
  lambda_charge: 0.25
  lambda_temporal: 0.5
  lambda_consistency: 0.25

train:
  epochs: 10
  lr: 0.0002
  weight_decay: 0.01
  grad_clip: 5.0
  gradient_accumulation_steps: 2
  warmup_steps: 5000
  amp: true
  keep_epoch_checkpoints: false
  log_every: 100
  save_every: 1
  device: cuda
  seed: 2026

ablation:
  name: md_trajectory_value
  condition: no_md
  trajectory_mode: equilibrium_plus_cgn_can
  preserve_frame_count: true
```
