## Supplementary material for "PG-MLD: Physics-Guided Molecular Representation Learning via Dynamic 3D Trajectory Distillation": Code and Data: README.md

# MD trajectory value ablation

This ablation compares two conditions while keeping the teacher architecture,
frame count, training losses, optimizer, and pretraining data fixed.

- **with MD**: the existing ordered MD trajectory model
  (`checkpoints/pretrain_zinc_dynamic_3d_liquid/latest.pt`).
- **without MD**: a separately preprocessed sequence containing the minimized
  equilibrium conformer and static CGN/CAN views. No MD coordinate file is
  read. The views are cycled to the same `T=8` input length, retaining the same
  SE(3)+Liquid architecture and sequence length.

First run `scripts/build_no_md_static_dataset.py`. The no-MD teacher must also
receive this static dataset during 3D-to-1D distillation. Use
`pretrain_static_3d_to_1d_chemblm.py` for this condition.
