## Supplementary material for "PG-MLD: Physics-Guided Molecular Representation Learning via Dynamic 3D Trajectory Distillation": Code and Data: dynamic.yaml

```
experiment:
  name: ChemBLM
  variant: dynamic

paths:
  data_root: output/molecular/reg/moleculeace
  pretrained_checkpoint: checkpoints/pretrain_zinc_dynamic_3d_to_1d_chemblm/latest.pt
  checkpoint_root: checkpoints/cliff/chemblm/dynamic
  result_root: logs/cliff/chemblm/dynamic

data:
  dataset: CHEMBL2034_Ki
  smiles_col: smiles
  target_col: y [pEC50/pKi]
  cliff_col: cliff_mol
  split_col: split
  val_frac_from_train: 0.1
  validation_seed: 42
  batch_size: 32
  num_workers: 2

model:
  family: chemblm
  model_name: comld/models/ChemBLM
  trust_remote_code: false
  use_safetensors: false
  freeze_encoder: false

head:
  hidden_dim: 256
  dropout: 0.1

train:
  epochs: 60
  early_stopping_patience: 10
  lr: 0.00001
  weight_decay: 0.01
  grad_clip: 1.0
  gradient_accumulation_steps: 1
  warmup_ratio: 0.05
  amp: true
  log_every: 20
  device: auto
  seed: 42
```
