## Supplementary material for "PG-MLD: Physics-Guided Molecular Representation Learning via Dynamic 3D Trajectory Distillation": Code and Data: README.md

# MoleculeACE activity-cliff experiment

This experiment uses the official MoleculeACE CSV and its fixed `train`/`test`
assignment. A validation subset is created only from official training rows,
stratified by `cliff_mol`. The official test set is evaluated once after model
selection.

Reported primary metrics:

- `test.all_rmse`: RMSE on every official test molecule.
- `test.cliff_rmse`: RMSE on official test activity-cliff molecules.

Download and validate one official target:

```bash
python scripts/download_moleculeace.py --datasets CHEMBL2034_Ki

PYTHONPATH=. python -m comld.train.finetune_cliff_1d \
  --config configs/cliff/chemblm/original.yaml \
  --dataset CHEMBL2034_Ki \
  --seed 42 \
  --validate-only
```

Run original ChemBLM and PG-MLD-distilled ChemBLM:

```bash
TRANSFORMERS_OFFLINE=1 HF_HUB_OFFLINE=1 TOKENIZERS_PARALLELISM=false \
PYTHONPATH=. python -m comld.train.finetune_cliff_1d \
  --config configs/cliff/chemblm/original.yaml \
  --dataset CHEMBL2034_Ki \
  --seed 42

TRANSFORMERS_OFFLINE=1 HF_HUB_OFFLINE=1 TOKENIZERS_PARALLELISM=false \
PYTHONPATH=. python -m comld.train.finetune_cliff_1d \
  --config configs/cliff/chemblm/dynamic.yaml \
  --dataset CHEMBL2034_Ki \
  --seed 42
```

Use the same dataset and seed for both variants. For a final paper table, repeat
with seeds 42, 43, and 44 and report mean +/- standard deviation.
