## Supplementary material for "PG-MLD: Physics-Guided Molecular Representation Learning via Dynamic 3D Trajectory Distillation": Code and Data: finetune_esol_1d_random.yaml

```
paths:
  raw_csv: output/molecular/reg/esol/ogb_official/esol/mapping/mol.csv.gz
  split_dir:
  pretrained_checkpoint: checkpoints/pretrain_zinc_dynamic_3d_to_1d/latest.pt
  best_checkpoint: checkpoints/finetune_esol_1d_dynamic_random/best.pt

data:
  smiles_col: smiles
  label_col: measured log solubility in mols per litre
  task_type: regression
  batch_size: 32
  num_workers: 0
  split_method: random
  train_frac: 0.8
  val_frac: 0.1
  max_molecules:

model:
  model_name: comld/models/DeepChemChemBERTa-77M-MLM
  freeze_encoder: false

head:
  hidden_dim: 126
  dropout: 0.1

train:
  epochs: 100
  early_stopping_patience: 20
  lr: 0.00002
  weight_decay: 0.01
  grad_clip: 1.0
  log_every: 20
  device: auto
  seed: 2024
```
