## Supplementary material for "PG-MLD: Physics-Guided Molecular Representation Learning via Dynamic 3D Trajectory Distillation": Code and Data: finetune_bbbp_chemberta2_distilled.yaml

```
paths:
  raw_csv: output/molecular/cla/bbbp/ogb_official/bbbp/mapping/mol.csv.gz
  split_dir: output/molecular/cla/bbbp/ogb_official/bbbp/split/scaffold
  pretrained_checkpoint: checkpoints/pretrain_zinc_dynamic_3d_to_1d_chemberta2/latest.pt
  best_checkpoint: checkpoints/finetune_bbbp_chemberta2_distilled_scaffold/best.pt

data:
  smiles_col: smiles
  label_col: p_np
  task_type: classification
  batch_size: 32
  num_workers: 0
  split_method: scaffold
  train_frac: 0.8
  val_frac: 0.1
  max_molecules:

model:
  model_name: comld/models/DeepChemChemBERTa-77M-MTR
  freeze_encoder: false

head:
  hidden_dim: 256
  dropout: 0.1

train:
  epochs: 100
  early_stopping_patience: 20
  lr: 0.00001
  weight_decay: 0.01
  grad_clip: 1.0
  log_every: 20
  device: auto
  seed: 2026
```
