## Supplementary material for "PG-MLD: Physics-Guided Molecular Representation Learning via Dynamic 3D Trajectory Distillation": Code and Data: pretrain_dynamic_3d_to_1d_zinc.yaml

```
paths:
  dynamic_md_pt: zinc/zinc15_250K_openmm_full
  teacher_dynamic_checkpoint: checkpoints/pretrain_zinc_dynamic_3d_liquid/latest.pt
  checkpoint_dir: checkpoints/pretrain_zinc_dynamic_3d_to_1d

data:
  batch_size: 8
  num_workers: 2
  max_molecules:
  max_frames: 8
  frame_stride: 1
  random_frame_crop: true

student_1d:
  model_name: comld/models/DeepChemChemBERTa-77M-MLM

loss:
  lambda_atom: 1.0
  lambda_mol: 1.0
  lambda_contrastive: 0.1
  contrastive_temperature: 0.1
  lambda_mlm: 0.1
  mlm_probability: 0.15

train:
  epochs: 10
  lr: 0.00002
  weight_decay: 0.01
  grad_clip: 1.0
  gradient_accumulation_steps: 2
  warmup_steps: 2000
  amp: true
  keep_epoch_checkpoints: false
  log_every: 100
  save_every: 1
  device: cuda
  seed: 2026
```
