## Supplementary material for "PG-MLD: Physics-Guided Molecular Representation Learning via Dynamic 3D Trajectory Distillation": Code and Data: original.yaml

```
experiment:
  name: ChemBERTa
  variant: original

paths:
  raw_csv: output/molecular/reg/qm9/qm9.csv.gz
  uncharacterized_txt: output/molecular/reg/qm9/uncharacterized.txt
  pretrained_checkpoint:
  checkpoint_root: checkpoints/qm9/chemberta/original
  result_root: logs/qm9/chemberta/original

data:
  smiles_col: smiles
  source_energy_unit: hartree
  target: mu
  split_seed: 42
  train_size: 110000
  val_size: 10000
  batch_size: 64
  num_workers: 2

model:
  family: chemberta
  model_name: comld/models/DeepChemChemBERTa-77M-MLM
  freeze_encoder: false

head:
  hidden_dim: 256
  dropout: 0.1

train:
  epochs: 30
  early_stopping_patience: 8
  lr: 0.00002
  weight_decay: 0.01
  grad_clip: 1.0
  gradient_accumulation_steps: 1
  warmup_ratio: 0.05
  amp: true
  log_every: 100
  device: auto
  seed: 42
```
