## Supplementary material for "PG-MLD: Physics-Guided Molecular Representation Learning via Dynamic 3D Trajectory Distillation": Code and Data: README.md

# QM9 single-property regression

This directory contains a separate QM9 benchmark pipeline. It does not modify
the existing MoleculeNet classification/regression trainers.

## Protocol

- Standard 130,831 characterized QM9 molecules.
- One independently trained model per property.
- Fixed random split shared by every model: 110,000 train, 10,000 validation,
  and 10,831 test molecules (`seed=42`).
- Targets are standardized using training-set statistics only.
- Model selection uses validation MAE. The test set is evaluated once after
  loading the best validation checkpoint.
- `mu` is reported in D, `alpha` in `a0^3`, and energy targets in meV.

## Data

```bash
cd ~/CoMLD
wget -c https://ndownloader.figshare.com/files/3195404 \
  -O output/molecular/reg/qm9/uncharacterized.txt
```

The trainer reads `output/molecular/reg/qm9/qm9.csv.gz` directly, removes the
3,054 uncharacterized molecules using the list above, converts energy targets
from Hartree to eV, and creates the fixed split in memory. No conformer or
trajectory preprocessing is used for this downstream benchmark.

## Run one target

Original ChemBLM:

```bash
TRANSFORMERS_OFFLINE=1 HF_HUB_OFFLINE=1 TOKENIZERS_PARALLELISM=false \
PYTHONPATH=. python -m comld.train.finetune_qm9_1d \
  --config configs/qm9/chemblm/original.yaml \
  --target mu \
  --seed 42
```

PG-MLD-distilled ChemBLM:

```bash
TRANSFORMERS_OFFLINE=1 HF_HUB_OFFLINE=1 TOKENIZERS_PARALLELISM=false \
PYTHONPATH=. python -m comld.train.finetune_qm9_1d \
  --config configs/qm9/chemblm/dynamic.yaml \
  --target mu \
  --seed 42
```

Replace `mu` with one of:

`mu, alpha, homo, lumo, gap, zpve, u0, u, h`.

## Run the nine properties

```bash
PYTHONPATH=. python scripts/run_qm9.py \
  --config configs/qm9/chemblm/dynamic.yaml \
  --targets all \
  --seeds 42,43,44
```

Existing checkpoints are skipped unless `--overwrite` is supplied.

## Build the result table

```bash
PYTHONPATH=. python scripts/summarize_qm9.py
```

This reads real `result.json` files and writes:

- `output/qm9/qm9_mae_summary.csv`
- `output/qm9/qm9_mae_table.tex`

The LaTeX table marks the lowest mean MAE in bold and the second lowest with
an underline. It never invents missing results.
