## Supplementary material for "PG-MLD: Physics-Guided Molecular Representation Learning via Dynamic 3D Trajectory Distillation": Code and Data: README.md

# Config layout

```text
configs/
  pretrain/
    pretrain_dynamic_3d_zinc.yaml
    pretrain_dynamic_3d_to_1d_zinc.yaml
    pretrain_dynamic_3d_to_1d_zinc_chemberta2.yaml
    pretrain_dynamic_3d_to_1d_zinc_molformer.yaml
    pretrain_dynamic_3d_to_1d_zinc_chemblm.yaml

  finetune/
    scaffold/
      Original scaffold-split downstream configs.

    random/
      chemberta/
        ChemBERTa original/distilled random-split configs.

      chemberta2/
        ChemBERTa-2 random-split configs.

      molformer/
        MoLFormer random/random-scaffold configs.

    random_scaffold/
      SCAGE-style random scaffold split configs. Molecules are grouped by
      Murcko scaffold first, then scaffold groups are randomly shuffled by seed
      before train/validation/test assignment.
```

Recommended checks before running a downstream experiment:

```bash
grep -n "split_method" <config>
grep -n "pretrained_checkpoint" <config>
grep -n "model_name" <config>
grep -n "early_stopping_metric" <config>
```

For formal random-split comparison, logs should show:

```text
Split: method=random source=generated
Early stopping metric: val_primary
```

For SCAGE-style random scaffold comparison, logs should show:

```text
Split: method=random_scaffold source=generated
Early stopping metric: val_primary
```
