## Supplementary material for "PG-MLD: Physics-Guided Molecular Representation Learning via Dynamic 3D Trajectory Distillation": Code and Data: README.md

# ogbg-mol

## mol.csv.gz

The columns are organized as follows:

[task_1, ... task_n], SMILES, mol_id

n is the number of tasks, task_1,...,task_n represent different kinds of labels assigned to the molecule, and the SMILES string uniquely identifies the molecule.
i-th data object represents i-th molecule in mol.csv.gz.
