## Supplementary material for "PG-MLD: Physics-Guided Molecular Representation Learning via Dynamic 3D Trajectory Distillation": Code and Data: README.md

# PG-MLD

PG-MLD is a dynamic 3D physics-guided knowledge-distillation framework for
molecular language models. During training, a 3D teacher learns geometric,
conformational-dynamic, and local electronic information from molecular
trajectories. The learned information is transferred to a SMILES student so
that downstream inference requires only SMILES strings.

## Repository structure

- `comld/data`: molecular trajectory loading and featurization.
- `comld/models`: dynamic 3D teacher and molecular-language student modules.
- `comld/train`: pretraining, distillation, and downstream finetuning.
- `configs`: experiment configurations.
- `scripts`: data preparation and experiment launchers.
- `visualization`: PCA, SHAP, and representation analysis.

## External assets

Large files are intentionally not stored in this repository:

- molecular datasets and MD trajectories;
- pretrained language-model weights;
- PG-MLD pretraining and downstream checkpoints;
- logs, predictions, and generated figures.

Download or generate these assets separately and place them at the relative
paths specified in the YAML configurations. The data preparation scripts under
`scripts/` reproduce the expected project layout.

## Main training entry points

Dynamic 3D teacher pretraining:

```bash
PYTHONPATH=. python -m comld.train.pretrain_dynamic_3d \
  --config configs/pretrain/pretrain_dynamic_3d_zinc.yaml
```

Dynamic 3D-to-1D distillation:

```bash
PYTHONPATH=. python -m comld.train.pretrain_dynamic_3d_to_1d_chemblm \
  --config configs/pretrain/pretrain_dynamic_3d_to_1d_zinc_chemblm.yaml
```

MoleculeACE activity-cliff evaluation:

```bash
python scripts/download_moleculeace.py --datasets CHEMBL2034_Ki

PYTHONPATH=. python -m comld.train.finetune_cliff_1d \
  --config configs/cliff/chemblm/dynamic.yaml \
  --dataset CHEMBL2034_Ki \
  --seed 42
```

Further commands and dataset-specific settings are documented in the
corresponding configuration directories.
