## Supplementary material for "PG-MLD: Physics-Guided Molecular Representation Learning via Dynamic 3D Trajectory Distillation": Code and Data: build_no_md_static_dataset.py

```
from __future__ import annotations

import argparse
import math
from pathlib import Path
from typing import Any

import torch
from rdkit import Chem
from rdkit.Chem import AllChem
from tqdm import tqdm


def torch_load(path: Path) -> list[dict[str, Any]]:
    try:
        value = torch.load(path, map_location="cpu", weights_only=False)
    except TypeError:
        value = torch.load(path, map_location="cpu")
    if not isinstance(value, list):
        raise ValueError(f"Expected a list in {path}, got {type(value).__name__}")
    return value


def safe_float(value: str, default: float = 0.0) -> float:
    try:
        number = float(value)
    except (TypeError, ValueError):
        return default
    return number if math.isfinite(number) else default


def electronic_features(smiles: str, expected_symbols: list[str]) -> torch.Tensor:
    """Recompute formal/Gasteiger charges without reading an MD file."""

    mol = Chem.MolFromSmiles(smiles)
    if mol is None:
        raise ValueError(f"Invalid SMILES: {smiles}")
    mol = Chem.AddHs(mol)
    try:
        AllChem.ComputeGasteigerCharges(mol)
    except Exception:
        pass

    symbols = [atom.GetSymbol() for atom in mol.GetAtoms()]
    if symbols != expected_symbols:
        raise ValueError(
            f"Atom order mismatch for {smiles}: RDKit={symbols}, packed={expected_symbols}"
        )

    rows = []
    for atom in mol.GetAtoms():
        formal = float(atom.GetFormalCharge())
        gasteiger = safe_float(
            atom.GetProp("_GasteigerCharge")
            if atom.HasProp("_GasteigerCharge")
            else "0"
        )
        rows.append([formal, max(min(gasteiger, 2.0), -2.0)])
    return torch.tensor(rows, dtype=torch.float)


def fixed_view_sequence(
    equilibrium_coords: torch.Tensor,
    augmented_coords: torch.Tensor,
    target_frames: int,
) -> torch.Tensor:
    """Create exactly T static views without using any MD coordinates."""

    equilibrium_coords = torch.as_tensor(equilibrium_coords, dtype=torch.float)
    augmented_coords = torch.as_tensor(augmented_coords, dtype=torch.float)
    if equilibrium_coords.ndim != 2 or equilibrium_coords.shape[-1] != 3:
        raise ValueError(
            f"Expected equilibrium coordinates [N,3], got {tuple(equilibrium_coords.shape)}"
        )
    if augmented_coords.ndim == 2:
        augmented_coords = augmented_coords.unsqueeze(0)
    if augmented_coords.ndim != 3 or augmented_coords.shape[-1] != 3:
        raise ValueError(
            f"Expected augmented coordinates [V,N,3], got {tuple(augmented_coords.shape)}"
        )
    if equilibrium_coords.shape[0] != augmented_coords.shape[1]:
        raise ValueError("Equilibrium and augmented views have different atom counts")

    views = torch.cat([equilibrium_coords.unsqueeze(0), augmented_coords], dim=0)
    target_frames = max(int(target_frames), 1)
    repeats = math.ceil(target_frames / int(views.shape[0]))
    return views.repeat((repeats, 1, 1))[:target_frames].contiguous()


def build_static_item(
    base_item: dict[str, Any],
    augmented_item: dict[str, Any],
    target_frames: int,
    view_delta_ps: float,
) -> dict[str, Any]:
    row_idx = int(base_item["row_idx"])
    if int(augmented_item["row_idx"]) != row_idx:
        raise ValueError("base.pt and base_cgn_can.pt row indices are not aligned")
    smiles = str(base_item["smiles"])
    if str(augmented_item["smiles"]) != smiles:
        raise ValueError(f"SMILES mismatch at row {row_idx}")
    atom_symbols = [str(symbol) for symbol in base_item["atom_symbols"]]
    if [str(symbol) for symbol in augmented_item["atom_symbols"]] != atom_symbols:
        raise ValueError(f"Atom symbol mismatch at row {row_idx}")

    coords = fixed_view_sequence(
        equilibrium_coords=base_item["coords"],
        augmented_coords=augmented_item["coords"],
        target_frames=target_frames,
    )
    electronic = electronic_features(smiles, atom_symbols)
    if electronic.shape[0] != coords.shape[1]:
        raise ValueError(f"Charge/coordinate atom mismatch at row {row_idx}")

    timestep_fs = float(view_delta_ps) * 1000.0
    output = {
        "row_idx": row_idx,
        "smiles": smiles,
        "num_heavy_atoms": base_item.get("num_heavy_atoms"),
        "atom_symbols": atom_symbols,
        "coords": coords,
        "electronic_feats": electronic,
        "trajectory_electronic_feats": electronic.unsqueeze(0).repeat(
            coords.shape[0], 1, 1
        ),
        "formal_charge": electronic[:, 0].clone(),
        "gasteiger_charge": electronic[:, 1].clone(),
        # Numerical spacing for the retained Liquid integrator.  This is not
        # interpreted as physical MD time in the no-MD condition.
        "timestep_fs": timestep_fs,
        "report_interval": 1,
        "view_type": "equilibrium_plus_cgn_can_no_md",
    }
    for key in ("cgn_sigma", "can_max_angle_deg"):
        if key in augmented_item:
            output[key] = augmented_item[key]
    return output


def preprocess_shard(
    shard: Path,
    output_root: Path,
    target_frames: int,
    view_delta_ps: float,
    overwrite: bool,
) -> tuple[int, int]:
    base_path = shard / "base.pt"
    augmented_path = shard / "base_cgn_can.pt"
    if not base_path.exists() or not augmented_path.exists():
        raise FileNotFoundError(f"Missing base static files in {shard}")

    output_dir = output_root / shard.name
    output_path = output_dir / "static_cgn_can_electronic.pt"
    if output_path.exists() and not overwrite:
        existing = torch_load(output_path)
        print(f"Skip existing: {output_path} ({len(existing)} molecules)")
        return len(existing), 0

    base_items = torch_load(base_path)
    augmented_items = torch_load(augmented_path)
    augmented_by_row = {int(item["row_idx"]): item for item in augmented_items}

    output_items = []
    skipped = 0
    for base_item in base_items:
        row_idx = int(base_item["row_idx"])
        augmented_item = augmented_by_row.get(row_idx)
        if augmented_item is None:
            skipped += 1
            continue
        try:
            output_items.append(
                build_static_item(
                    base_item,
                    augmented_item,
                    target_frames=target_frames,
                    view_delta_ps=view_delta_ps,
                )
            )
        except (ValueError, RuntimeError) as exc:
            skipped += 1
            print(f"Skip row {row_idx}: {exc}")

    if not output_items:
        raise RuntimeError(f"No valid static molecules produced from {shard}")
    output_dir.mkdir(parents=True, exist_ok=True)
    temporary_path = output_path.with_suffix(".pt.tmp")
    torch.save(output_items, temporary_path)
    temporary_path.replace(output_path)
    print(f"Saved: {output_path} ({len(output_items)} molecules, skipped={skipped})")
    return len(output_items), skipped


def build_dataset(args: argparse.Namespace) -> None:
    input_root = Path(args.input_root)
    output_root = Path(args.output_root)
    shards = sorted(input_root.glob("shard_*"))
    shards = [shard for shard in shards if shard.is_dir()]
    if args.max_shards is not None:
        shards = shards[: int(args.max_shards)]
    if not shards:
        raise FileNotFoundError(f"No shard_* directories found under {input_root}")

    total = 0
    skipped = 0
    skipped_shards = 0
    for shard in tqdm(shards, desc="Build no-MD static shards"):
        try:
            count, shard_skipped = preprocess_shard(
                shard=shard,
                output_root=output_root,
                target_frames=args.target_frames,
                view_delta_ps=args.view_delta_ps,
                overwrite=args.overwrite,
            )
        except FileNotFoundError as exc:
            skipped_shards += 1
            print(f"Skip incomplete shard: {exc}")
            continue
        total += count
        skipped += shard_skipped

    completed_shards = len(shards) - skipped_shards
    print(f"Static no-MD dataset: {total} molecules in {completed_shards} shards")
    print(f"Skipped incomplete shards: {skipped_shards}")
    print(f"Skipped molecules: {skipped}")
    print(f"Output root: {output_root}")


def build_arg_parser() -> argparse.ArgumentParser:
    parser = argparse.ArgumentParser(
        description=(
            "Build an equilibrium+CGN/CAN static dataset for the no-MD ablation. "
            "No MD coordinate file is read."
        )
    )
    parser.add_argument(
        "--input-root",
        default="zinc/zinc15_250K_openmm_full",
    )
    parser.add_argument(
        "--output-root",
        default="zinc/zinc15_250K_static_cgn_can_electronic",
    )
    parser.add_argument("--target-frames", type=int, default=8)
    parser.add_argument("--view-delta-ps", type=float, default=0.1)
    parser.add_argument("--max-shards", type=int)
    parser.add_argument("--overwrite", action="store_true")
    return parser


def main() -> None:
    build_dataset(build_arg_parser().parse_args())


if __name__ == "__main__":
    main()
```
