## Supplementary material for "PG-MLD: Physics-Guided Molecular Representation Learning via Dynamic 3D Trajectory Distillation": Code and Data: download_moleculeace.py

```
from __future__ import annotations

import argparse
import csv
import shutil
import urllib.request
from pathlib import Path


DATASETS = (
    "CHEMBL1862_Ki",
    "CHEMBL1871_Ki",
    "CHEMBL2034_Ki",
    "CHEMBL2047_EC50",
    "CHEMBL204_Ki",
    "CHEMBL2147_Ki",
    "CHEMBL214_Ki",
    "CHEMBL218_EC50",
    "CHEMBL219_Ki",
    "CHEMBL228_Ki",
    "CHEMBL231_Ki",
    "CHEMBL233_Ki",
    "CHEMBL234_Ki",
    "CHEMBL235_EC50",
    "CHEMBL236_Ki",
    "CHEMBL237_EC50",
    "CHEMBL237_Ki",
    "CHEMBL238_Ki",
    "CHEMBL239_EC50",
    "CHEMBL244_Ki",
    "CHEMBL262_Ki",
    "CHEMBL264_Ki",
    "CHEMBL2835_Ki",
    "CHEMBL287_Ki",
    "CHEMBL2971_Ki",
    "CHEMBL3979_EC50",
    "CHEMBL4005_Ki",
    "CHEMBL4203_Ki",
    "CHEMBL4616_EC50",
    "CHEMBL4792_Ki",
)
BASE_URL = (
    "https://raw.githubusercontent.com/molML/MoleculeACE/main/"
    "MoleculeACE/Data/benchmark_data"
)
REQUIRED_COLUMNS = {"smiles", "cliff_mol", "split"}


def parse_datasets(values: list[str]) -> list[str]:
    if len(values) == 1 and values[0].strip().lower() == "all":
        return list(DATASETS)
    names: list[str] = []
    for value in values:
        names.extend(item.strip() for item in value.split(",") if item.strip())
    invalid = [name for name in names if name not in DATASETS]
    if invalid:
        raise ValueError(f"Unknown MoleculeACE datasets {invalid}.")
    return names


def validate_csv(path: Path) -> None:
    with path.open("r", encoding="utf-8", newline="") as handle:
        reader = csv.reader(handle)
        columns = set(next(reader))
    missing = REQUIRED_COLUMNS - columns
    if "y" not in columns and "y [pEC50/pKi]" not in columns:
        missing.add("y or y [pEC50/pKi]")
    if missing:
        raise ValueError(f"Downloaded file {path} is missing columns: {sorted(missing)}")


def download_dataset(name: str, output_dir: Path, overwrite: bool) -> Path:
    destination = output_dir / f"{name}.csv"
    if destination.exists() and not overwrite:
        validate_csv(destination)
        print(f"Already present: {destination}")
        return destination

    output_dir.mkdir(parents=True, exist_ok=True)
    temporary = destination.with_suffix(".csv.part")
    temporary.unlink(missing_ok=True)
    url = f"{BASE_URL}/{name}.csv"
    print(f"Downloading {url}")
    try:
        request = urllib.request.Request(url, headers={"User-Agent": "CoMLD-MoleculeACE/1.0"})
        with urllib.request.urlopen(request, timeout=120) as response, temporary.open("wb") as out:
            shutil.copyfileobj(response, out)
        validate_csv(temporary)
        temporary.replace(destination)
    except BaseException:
        temporary.unlink(missing_ok=True)
        raise
    print(f"Saved: {destination}")
    return destination


def main() -> None:
    parser = argparse.ArgumentParser(description="Download official MoleculeACE benchmark CSVs.")
    parser.add_argument(
        "--datasets",
        nargs="+",
        default=["CHEMBL2034_Ki"],
        help="Dataset names, comma-separated names, or 'all'.",
    )
    parser.add_argument(
        "--output-dir",
        default="output/molecular/reg/moleculeace",
    )
    parser.add_argument("--overwrite", action="store_true")
    args = parser.parse_args()

    output_dir = Path(args.output_dir)
    selected = parse_datasets(args.datasets)
    for dataset in selected:
        download_dataset(dataset, output_dir, overwrite=args.overwrite)
    print(f"MoleculeACE download complete: {len(selected)} dataset(s).")


if __name__ == "__main__":
    main()
```
