## Supplementary material for "PG-MLD: Physics-Guided Molecular Representation Learning via Dynamic 3D Trajectory Distillation": Code and Data: download_ogb_mol_datasets.py

```
from __future__ import annotations

import argparse
import shutil
import urllib.request
import zipfile
from pathlib import Path


OGB_CSV_MOL_URLS = {
    "bace": "http://snap.stanford.edu/ogb/data/graphproppred/csv_mol_download/bace.zip",
    "bbbp": "http://snap.stanford.edu/ogb/data/graphproppred/csv_mol_download/bbbp.zip",
    "clintox": "http://snap.stanford.edu/ogb/data/graphproppred/csv_mol_download/clintox.zip",
    "esol": "http://snap.stanford.edu/ogb/data/graphproppred/csv_mol_download/esol.zip",
    "freesolv": "http://snap.stanford.edu/ogb/data/graphproppred/csv_mol_download/freesolv.zip",
    "hiv": "http://snap.stanford.edu/ogb/data/graphproppred/csv_mol_download/hiv.zip",
    "lipophilicity": "http://snap.stanford.edu/ogb/data/graphproppred/csv_mol_download/lipophilicity.zip",
    "tox21": "http://snap.stanford.edu/ogb/data/graphproppred/csv_mol_download/tox21.zip",
    "toxcast": "http://snap.stanford.edu/ogb/data/graphproppred/csv_mol_download/toxcast.zip",
}

DATASET_GROUPS = {
    "bace": "cla",
    "bbbp": "cla",
    "clintox": "cla",
    "hiv": "cla",
    "tox21": "cla",
    "toxcast": "cla",
    "esol": "reg",
    "freesolv": "reg",
    "lipophilicity": "reg",
}


def download(url: str, out_path: Path) -> None:
    out_path.parent.mkdir(parents=True, exist_ok=True)
    if out_path.exists() and out_path.stat().st_size > 0:
        print(f"Using existing archive: {out_path}")
        return
    print(f"Downloading {url}")
    with urllib.request.urlopen(url) as response, out_path.open("wb") as handle:
        shutil.copyfileobj(response, handle)
    print(f"Saved: {out_path}")


def extract_dataset(dataset: str, archive_path: Path, out_root: Path) -> Path:
    dataset = dataset.lower()
    group = DATASET_GROUPS[dataset]
    extract_root = out_root / "molecular" / group / dataset / "ogb_official"
    extract_root.mkdir(parents=True, exist_ok=True)
    print(f"Extracting {archive_path} -> {extract_root}")
    with zipfile.ZipFile(archive_path) as zf:
        zf.extractall(extract_root)

    dataset_root = extract_root / dataset
    required = [
        dataset_root / "mapping" / "mol.csv.gz",
        dataset_root / "split" / "scaffold" / "train.csv.gz",
        dataset_root / "split" / "scaffold" / "valid.csv.gz",
        dataset_root / "split" / "scaffold" / "test.csv.gz",
    ]
    missing = [str(path) for path in required if not path.exists()]
    if missing:
        raise FileNotFoundError(
            f"Downloaded archive did not create expected OGB files for {dataset}. Missing: {missing}"
        )
    return dataset_root


def main() -> None:
    parser = argparse.ArgumentParser(description="Download official OGB csv_mol datasets.")
    parser.add_argument(
        "--datasets",
        nargs="+",
        default=["bace", "clintox"],
        choices=sorted(OGB_CSV_MOL_URLS),
        help="Datasets to download.",
    )
    parser.add_argument(
        "--all",
        action="store_true",
        help="Download all supported MoleculeNet OGB csv_mol datasets used by this project.",
    )
    parser.add_argument("--out-root", default="output", help="Root directory that contains molecular/cla.")
    parser.add_argument("--archive-dir", default="output/ogb_downloads", help="Where to store downloaded zip files.")
    args = parser.parse_args()

    out_root = Path(args.out_root)
    archive_dir = Path(args.archive_dir)
    datasets = sorted(OGB_CSV_MOL_URLS) if args.all else args.datasets
    for dataset in datasets:
        dataset = dataset.lower()
        archive_path = archive_dir / f"{dataset}.zip"
        download(OGB_CSV_MOL_URLS[dataset], archive_path)
        dataset_root = extract_dataset(dataset, archive_path, out_root)
        print(f"Ready: {dataset_root}")


if __name__ == "__main__":
    main()
```
