## Supplementary material for "PG-MLD: Physics-Guided Molecular Representation Learning via Dynamic 3D Trajectory Distillation": Code and Data: generate_openmm_md.py

```
from __future__ import annotations

import argparse
import random
from pathlib import Path

import numpy as np
from rdkit import Chem
from rdkit.Chem import AllChem
from rdkit.Chem import rdMolTransforms


def build_rdkit_mol(smiles: str, seed: int, max_attempts: int) -> Chem.Mol:
    mol = Chem.MolFromSmiles(smiles)
    if mol is None:
        raise ValueError(f"Invalid SMILES: {smiles}")
    mol = Chem.AddHs(mol)

    params = AllChem.ETKDGv3()
    params.randomSeed = int(seed)
    params.useRandomCoords = True
    params.numThreads = 0
    status = AllChem.EmbedMolecule(mol, params)
    if status != 0:
        status = AllChem.EmbedMolecule(mol, randomSeed=int(seed), maxAttempts=int(max_attempts), useRandomCoords=True)
    if status != 0:
        raise RuntimeError(f"RDKit conformer embedding failed for {smiles}")

    try:
        AllChem.MMFFOptimizeMolecule(mol, maxIters=500)
    except Exception:
        AllChem.UFFOptimizeMolecule(mol, maxIters=500)
    return mol


def safe_float(value: str, default: float = 0.0) -> float:
    try:
        number = float(value)
    except (TypeError, ValueError):
        return default
    if not np.isfinite(number):
        return default
    return float(number)


def atom_symbols(mol: Chem.Mol) -> list[str]:
    return [atom.GetSymbol() for atom in mol.GetAtoms()]


def compute_charge_features(mol: Chem.Mol) -> tuple[np.ndarray, np.ndarray, np.ndarray]:
    work = Chem.Mol(mol)
    try:
        AllChem.ComputeGasteigerCharges(work)
    except Exception:
        pass

    formal_charges = []
    gasteiger_charges = []
    for atom in work.GetAtoms():
        formal_charges.append(float(atom.GetFormalCharge()))
        gasteiger = safe_float(atom.GetProp("_GasteigerCharge") if atom.HasProp("_GasteigerCharge") else "0")
        gasteiger_charges.append(max(min(gasteiger, 2.0), -2.0))

    formal = np.asarray(formal_charges, dtype=np.float32)
    gasteiger = np.asarray(gasteiger_charges, dtype=np.float32)
    electronic = np.stack([formal, gasteiger], axis=-1).astype(np.float32)
    return electronic, formal, gasteiger


def save_charge_csv(
    path: str | Path,
    symbols: list[str],
    electronic_feats: np.ndarray,
) -> None:
    path = Path(path)
    lines = ["atom_index,element,formal_charge,gasteiger_charge"]
    for idx, (symbol, values) in enumerate(zip(symbols, electronic_feats), start=1):
        lines.append(f"{idx},{symbol},{float(values[0]):.6f},{float(values[1]):.6f}")
    path.write_text("\n".join(lines) + "\n", encoding="utf-8")


def assign_openff_partial_charges(off_mol, method: str):
    method = str(method).strip().lower()
    if method in {"default", "none", ""}:
        return None
    off_mol.assign_partial_charges(partial_charge_method=method)
    return [off_mol]


def rdkit_coords_angstrom(mol: Chem.Mol) -> np.ndarray:
    conf = mol.GetConformer()
    coords = []
    for atom_idx in range(mol.GetNumAtoms()):
        pos = conf.GetAtomPosition(atom_idx)
        coords.append([pos.x, pos.y, pos.z])
    return np.asarray(coords, dtype=np.float32)


def set_rdkit_coords_angstrom(mol: Chem.Mol, coords: np.ndarray) -> Chem.Mol:
    work = Chem.Mol(mol)
    conf = work.GetConformer()
    for atom_idx, xyz in enumerate(np.asarray(coords, dtype=float)):
        conf.SetAtomPosition(int(atom_idx), tuple(float(value) for value in xyz))
    return work


def make_cgn_views(coords: np.ndarray, num_views: int, sigma: float, seed: int) -> np.ndarray:
    coords = np.asarray(coords, dtype=np.float32)
    if num_views <= 0:
        return np.empty((0, *coords.shape), dtype=np.float32)
    rng = np.random.default_rng(int(seed))
    noise = rng.normal(loc=0.0, scale=float(sigma), size=(int(num_views), *coords.shape))
    return (coords[None, :, :] + noise).astype(np.float32)


def make_cgn_trajectory(coords: np.ndarray, sigma: float, seed: int) -> np.ndarray:
    coords = np.asarray(coords, dtype=np.float32)
    if sigma <= 0:
        return coords.copy()
    rng = np.random.default_rng(int(seed))
    noise = rng.normal(loc=0.0, scale=float(sigma), size=coords.shape)
    return (coords + noise).astype(np.float32)


def rotatable_bond_torsions(mol: Chem.Mol) -> list[tuple[int, int, int, int]]:
    torsions: list[tuple[int, int, int, int]] = []
    for bond in mol.GetBonds():
        if bond.GetBondType() != Chem.rdchem.BondType.SINGLE:
            continue
        if bond.IsInRing():
            continue

        begin = bond.GetBeginAtom()
        end = bond.GetEndAtom()
        if begin.GetAtomicNum() == 1 or end.GetAtomicNum() == 1:
            continue

        begin_neighbors = [atom.GetIdx() for atom in begin.GetNeighbors() if atom.GetIdx() != end.GetIdx()]
        end_neighbors = [atom.GetIdx() for atom in end.GetNeighbors() if atom.GetIdx() != begin.GetIdx()]
        if not begin_neighbors or not end_neighbors:
            continue

        torsions.append((begin_neighbors[0], begin.GetIdx(), end.GetIdx(), end_neighbors[0]))
    return torsions


def make_can_views(
    mol: Chem.Mol,
    base_coords: np.ndarray,
    num_views: int,
    max_angle_deg: float,
    seed: int,
) -> tuple[np.ndarray, int]:
    if num_views <= 0:
        return np.empty((0, mol.GetNumAtoms(), 3), dtype=np.float32), 0

    base_mol = set_rdkit_coords_angstrom(mol, base_coords)
    torsions = rotatable_bond_torsions(base_mol)
    views = []
    rng = random.Random(int(seed))
    for _ in range(int(num_views)):
        work = Chem.Mol(base_mol)
        conf = work.GetConformer()
        for i, j, k, l in torsions:
            current = rdMolTransforms.GetDihedralDeg(conf, int(i), int(j), int(k), int(l))
            delta = rng.uniform(-float(max_angle_deg), float(max_angle_deg))
            rdMolTransforms.SetDihedralDeg(conf, int(i), int(j), int(k), int(l), current + delta)
        views.append(rdkit_coords_angstrom(work))
    return np.stack(views, axis=0).astype(np.float32), len(torsions)


def make_can_trajectory(
    mol: Chem.Mol,
    trajectory_coords: np.ndarray,
    max_angle_deg: float,
    seed: int,
) -> tuple[np.ndarray, int]:
    trajectory_coords = np.asarray(trajectory_coords, dtype=np.float32)
    if trajectory_coords.shape[0] == 0:
        return trajectory_coords.copy(), 0

    probe_mol = set_rdkit_coords_angstrom(mol, trajectory_coords[0])
    torsions = rotatable_bond_torsions(probe_mol)
    rng = random.Random(int(seed))
    frames = []
    for frame_coords in trajectory_coords:
        work = set_rdkit_coords_angstrom(mol, frame_coords)
        conf = work.GetConformer()
        for i, j, k, l in torsions:
            current = rdMolTransforms.GetDihedralDeg(conf, int(i), int(j), int(k), int(l))
            delta = rng.uniform(-float(max_angle_deg), float(max_angle_deg))
            rdMolTransforms.SetDihedralDeg(conf, int(i), int(j), int(k), int(l), current + delta)
        frames.append(rdkit_coords_angstrom(work))
    return np.stack(frames, axis=0).astype(np.float32), len(torsions)


def make_cgn_can_trajectory(
    mol: Chem.Mol,
    trajectory_coords: np.ndarray,
    cgn_sigma: float,
    can_max_angle_deg: float,
    seed: int,
) -> tuple[np.ndarray, int]:
    can_coords, num_torsions = make_can_trajectory(
        mol=mol,
        trajectory_coords=trajectory_coords,
        max_angle_deg=can_max_angle_deg,
        seed=seed,
    )
    cgn_can_coords = make_cgn_trajectory(
        can_coords,
        sigma=cgn_sigma,
        seed=seed + 1,
    )
    return cgn_can_coords.astype(np.float32), num_torsions


def make_base_cgn_can_views(
    mol: Chem.Mol,
    base_coords: np.ndarray,
    num_views: int,
    cgn_sigma: float,
    can_max_angle_deg: float,
    seed: int,
) -> tuple[np.ndarray, int]:
    can_coords, num_torsions = make_can_views(
        mol=mol,
        base_coords=base_coords,
        num_views=num_views,
        max_angle_deg=can_max_angle_deg,
        seed=seed,
    )
    cgn_can_coords = make_cgn_trajectory(
        can_coords,
        sigma=cgn_sigma,
        seed=seed + 1,
    )
    return cgn_can_coords.astype(np.float32), num_torsions


def repeat_atom_features(features: np.ndarray, num_frames: int) -> np.ndarray:
    features = np.asarray(features, dtype=np.float32)
    if num_frames <= 0:
        return np.empty((0, *features.shape), dtype=np.float32)
    return np.repeat(features[None, :, :], int(num_frames), axis=0).astype(np.float32)


def run_openmm_md(
    smiles: str,
    out_dir: str | Path,
    equilibration_steps: int,
    steps: int,
    report_interval: int,
    seed: int,
    temperature_kelvin: float,
    timestep_fs: float,
    friction_per_ps: float,
    platform_name: str,
    cgn_views: int,
    cgn_sigma: float,
    trajectory_cgn: bool,
    can_views: int,
    can_max_angle_deg: float,
    trajectory_can: bool,
    trajectory_cgn_can: bool,
    partial_charge_method: str,
    save_intermediates: bool,
) -> None:
    try:
        import mdtraj as md
        import openmm
        import openmm.app as app
        import openmm.unit as unit
        from openff.toolkit.topology import Molecule
        from openff.toolkit.typing.engines.smirnoff import ForceField as OpenFFForceField
    except ImportError as exc:
        raise ImportError(
            "This script needs openmm, openff-toolkit, rdkit, mdtraj, and numpy. "
            "Install them in the same conda env before running."
        ) from exc

    out_dir = Path(out_dir)
    out_dir.mkdir(parents=True, exist_ok=True)

    print("Building RDKit conformer...", flush=True)
    rdkit_mol = build_rdkit_mol(smiles, seed=seed, max_attempts=100)
    print("Preparing OpenFF molecule and topology...", flush=True)
    off_mol = Molecule.from_rdkit(rdkit_mol, allow_undefined_stereo=True, hydrogens_are_explicit=True)
    charge_from_molecules = assign_openff_partial_charges(off_mol, partial_charge_method)
    off_topology = off_mol.to_topology()

    forcefield = OpenFFForceField("openff_unconstrained-2.1.0.offxml")
    print(f"Creating OpenMM system with partial_charge_method={partial_charge_method}...", flush=True)
    if charge_from_molecules is None:
        system = forcefield.create_openmm_system(off_topology)
    else:
        system = forcefield.create_openmm_system(
            off_topology,
            charge_from_molecules=charge_from_molecules,
        )

    topology = off_topology.to_openmm()
    positions = off_mol.conformers[0].to_openmm()

    integrator = openmm.LangevinMiddleIntegrator(
        temperature_kelvin * unit.kelvin,
        friction_per_ps / unit.picosecond,
        timestep_fs * unit.femtosecond,
    )
    integrator.setRandomNumberSeed(int(seed))

    if platform_name.lower() == "auto":
        platform = None
    else:
        platform = openmm.Platform.getPlatformByName(platform_name)

    if platform is None:
        simulation = app.Simulation(topology, system, integrator)
    else:
        simulation = app.Simulation(topology, system, integrator, platform)

    simulation.context.setPositions(positions)
    print("Minimizing energy...", flush=True)
    simulation.minimizeEnergy(maxIterations=1000)
    simulation.context.setVelocitiesToTemperature(
        temperature_kelvin * unit.kelvin,
        int(seed),
    )
    if int(equilibration_steps) > 0:
        print(f"Equilibrating: steps={equilibration_steps}...", flush=True)
        simulation.step(int(equilibration_steps))

    pdb_path = out_dir / "molecule.pdb"
    dcd_path = out_dir / "trajectory.dcd"
    log_path = out_dir / "md_log.txt"
    base_npy_path = out_dir / "base_coords_angstrom.npy"
    base_cgn_can_npy_path = out_dir / "base_cgn_can_coords_angstrom.npy"
    core_bundle_path = out_dir / "md_cgn_can_with_electronic.npz"
    npy_path = out_dir / "trajectory_coords_angstrom.npy"
    electronic_npy_path = out_dir / "electronic_feats.npy"
    formal_charge_npy_path = out_dir / "formal_charge.npy"
    gasteiger_charge_npy_path = out_dir / "gasteiger_charge.npy"
    cgn_electronic_npy_path = out_dir / "cgn_electronic_feats.npy"
    can_electronic_npy_path = out_dir / "can_electronic_feats.npy"
    trajectory_electronic_npy_path = out_dir / "trajectory_electronic_feats.npy"
    trajectory_cgn_electronic_npy_path = out_dir / "trajectory_cgn_electronic_feats.npy"
    trajectory_can_electronic_npy_path = out_dir / "trajectory_can_electronic_feats.npy"
    trajectory_cgn_can_electronic_npy_path = out_dir / "trajectory_cgn_can_electronic_feats.npy"
    charge_csv_path = out_dir / "atom_charges.csv"
    cgn_npy_path = out_dir / "cgn_coords_angstrom.npy"
    trajectory_cgn_npy_path = out_dir / "trajectory_cgn_coords_angstrom.npy"
    can_npy_path = out_dir / "can_coords_angstrom.npy"
    trajectory_can_npy_path = out_dir / "trajectory_can_coords_angstrom.npy"
    trajectory_cgn_can_npy_path = out_dir / "trajectory_cgn_can_coords_angstrom.npy"
    bundle_path = out_dir / "md_augmented_views.npz"
    generated_paths = [
        pdb_path,
        dcd_path,
        log_path,
        base_npy_path,
        base_cgn_can_npy_path,
        core_bundle_path,
        npy_path,
        electronic_npy_path,
        formal_charge_npy_path,
        gasteiger_charge_npy_path,
        cgn_electronic_npy_path,
        can_electronic_npy_path,
        trajectory_electronic_npy_path,
        trajectory_cgn_electronic_npy_path,
        trajectory_can_electronic_npy_path,
        trajectory_cgn_can_electronic_npy_path,
        charge_csv_path,
        cgn_npy_path,
        trajectory_cgn_npy_path,
        can_npy_path,
        trajectory_can_npy_path,
        trajectory_cgn_can_npy_path,
        bundle_path,
    ]
    for path in generated_paths:
        try:
            Path(path).unlink(missing_ok=True)
        except OSError:
            pass

    minimized_state = simulation.context.getState(getPositions=True)
    minimized_positions = minimized_state.getPositions(asNumpy=True).value_in_unit(unit.angstrom)
    base_coords_angstrom = np.asarray(minimized_positions, dtype=np.float32)
    np.save(base_npy_path, base_coords_angstrom)
    electronic_feats, formal_charge, gasteiger_charge = compute_charge_features(rdkit_mol)
    np.save(electronic_npy_path, electronic_feats)
    np.save(formal_charge_npy_path, formal_charge)
    np.save(gasteiger_charge_npy_path, gasteiger_charge)
    save_charge_csv(charge_csv_path, atom_symbols(rdkit_mol), electronic_feats)

    with pdb_path.open("w", encoding="utf-8") as handle:
        app.PDBFile.writeFile(topology, minimized_state.getPositions(), handle)

    simulation.reporters.append(app.DCDReporter(str(dcd_path), int(report_interval)))
    simulation.reporters.append(
        app.StateDataReporter(
            str(log_path),
            int(report_interval),
            step=True,
            time=True,
            potentialEnergy=True,
            temperature=True,
            speed=True,
        )
    )

    print(f"Running MD: steps={steps}, report_interval={report_interval}, platform={platform_name}...", flush=True)
    simulation.step(int(steps))
    print("Reading trajectory and writing augmented views...", flush=True)

    traj = md.load(str(dcd_path), top=str(pdb_path))
    coords_angstrom = traj.xyz * 10.0
    np.save(npy_path, coords_angstrom.astype(np.float32))
    trajectory_electronic_feats = repeat_atom_features(electronic_feats, coords_angstrom.shape[0])
    np.save(trajectory_electronic_npy_path, trajectory_electronic_feats)

    base_cgn_can_views = max(int(cgn_views), int(can_views), 1)
    base_cgn_can_coords, base_cgn_can_torsions = make_base_cgn_can_views(
        rdkit_mol,
        base_coords=base_coords_angstrom,
        num_views=base_cgn_can_views,
        cgn_sigma=float(cgn_sigma),
        can_max_angle_deg=float(can_max_angle_deg),
        seed=int(seed) + 606,
    )
    np.save(base_cgn_can_npy_path, base_cgn_can_coords)

    cgn_coords = make_cgn_views(
        base_coords_angstrom,
        num_views=int(cgn_views),
        sigma=float(cgn_sigma),
        seed=int(seed) + 101,
    )
    np.save(cgn_npy_path, cgn_coords)
    cgn_electronic_feats = repeat_atom_features(electronic_feats, cgn_coords.shape[0])
    np.save(cgn_electronic_npy_path, cgn_electronic_feats)

    if trajectory_cgn:
        trajectory_cgn_coords = make_cgn_trajectory(
            coords_angstrom,
            sigma=float(cgn_sigma),
            seed=int(seed) + 202,
        )
        np.save(trajectory_cgn_npy_path, trajectory_cgn_coords)
        trajectory_cgn_electronic_feats = repeat_atom_features(electronic_feats, trajectory_cgn_coords.shape[0])
        np.save(trajectory_cgn_electronic_npy_path, trajectory_cgn_electronic_feats)
    else:
        trajectory_cgn_coords = np.empty((0, *coords_angstrom.shape[1:]), dtype=np.float32)
        trajectory_cgn_electronic_feats = repeat_atom_features(electronic_feats, 0)

    can_coords, num_torsions = make_can_views(
        rdkit_mol,
        base_coords=base_coords_angstrom,
        num_views=int(can_views),
        max_angle_deg=float(can_max_angle_deg),
        seed=int(seed) + 303,
    )
    np.save(can_npy_path, can_coords)
    can_electronic_feats = repeat_atom_features(electronic_feats, can_coords.shape[0])
    np.save(can_electronic_npy_path, can_electronic_feats)

    if trajectory_can:
        trajectory_can_coords, trajectory_can_torsions = make_can_trajectory(
            rdkit_mol,
            trajectory_coords=coords_angstrom,
            max_angle_deg=float(can_max_angle_deg),
            seed=int(seed) + 404,
        )
        np.save(trajectory_can_npy_path, trajectory_can_coords)
        trajectory_can_electronic_feats = repeat_atom_features(electronic_feats, trajectory_can_coords.shape[0])
        np.save(trajectory_can_electronic_npy_path, trajectory_can_electronic_feats)
    else:
        trajectory_can_coords = np.empty((0, *coords_angstrom.shape[1:]), dtype=np.float32)
        trajectory_can_torsions = num_torsions
        trajectory_can_electronic_feats = repeat_atom_features(electronic_feats, 0)

    if trajectory_cgn_can:
        trajectory_cgn_can_coords, trajectory_cgn_can_torsions = make_cgn_can_trajectory(
            rdkit_mol,
            trajectory_coords=coords_angstrom,
            cgn_sigma=float(cgn_sigma),
            can_max_angle_deg=float(can_max_angle_deg),
            seed=int(seed) + 505,
        )
        np.save(trajectory_cgn_can_npy_path, trajectory_cgn_can_coords)
        trajectory_cgn_can_electronic_feats = repeat_atom_features(electronic_feats, trajectory_cgn_can_coords.shape[0])
        np.save(trajectory_cgn_can_electronic_npy_path, trajectory_cgn_can_electronic_feats)
    else:
        trajectory_cgn_can_coords = np.empty((0, *coords_angstrom.shape[1:]), dtype=np.float32)
        trajectory_cgn_can_torsions = num_torsions
        trajectory_cgn_can_electronic_feats = repeat_atom_features(electronic_feats, 0)

    np.savez_compressed(
        bundle_path,
        base_coords_angstrom=base_coords_angstrom,
        electronic_feats=electronic_feats,
        formal_charge=formal_charge,
        gasteiger_charge=gasteiger_charge,
        trajectory_coords_angstrom=coords_angstrom.astype(np.float32),
        trajectory_electronic_feats=trajectory_electronic_feats,
        cgn_coords_angstrom=cgn_coords,
        cgn_electronic_feats=cgn_electronic_feats,
        trajectory_cgn_coords_angstrom=trajectory_cgn_coords,
        trajectory_cgn_electronic_feats=trajectory_cgn_electronic_feats,
        can_coords_angstrom=can_coords,
        can_electronic_feats=can_electronic_feats,
        trajectory_can_coords_angstrom=trajectory_can_coords,
        trajectory_can_electronic_feats=trajectory_can_electronic_feats,
        trajectory_cgn_can_coords_angstrom=trajectory_cgn_can_coords,
        trajectory_cgn_can_electronic_feats=trajectory_cgn_can_electronic_feats,
    )

    np.savez_compressed(
        core_bundle_path,
        smiles=np.asarray(smiles),
        atom_symbols=np.asarray(atom_symbols(rdkit_mol)),
        base_coords_angstrom=base_coords_angstrom,
        base_cgn_can_coords_angstrom=base_cgn_can_coords,
        trajectory_cgn_can_coords_angstrom=trajectory_cgn_can_coords,
        electronic_feats=electronic_feats,
        trajectory_cgn_can_electronic_feats=trajectory_cgn_can_electronic_feats,
        formal_charge=formal_charge,
        gasteiger_charge=gasteiger_charge,
        cgn_sigma=np.asarray(float(cgn_sigma), dtype=np.float32),
        can_max_angle_deg=np.asarray(float(can_max_angle_deg), dtype=np.float32),
        timestep_fs=np.asarray(float(timestep_fs), dtype=np.float32),
        report_interval=np.asarray(int(report_interval), dtype=np.int32),
    )

    if not save_intermediates:
        intermediate_paths = [
            pdb_path,
            dcd_path,
            log_path,
            npy_path,
            electronic_npy_path,
            formal_charge_npy_path,
            gasteiger_charge_npy_path,
            cgn_electronic_npy_path,
            can_electronic_npy_path,
            trajectory_electronic_npy_path,
            trajectory_cgn_electronic_npy_path,
            trajectory_can_electronic_npy_path,
            trajectory_cgn_can_electronic_npy_path,
            charge_csv_path,
            cgn_npy_path,
            trajectory_cgn_npy_path,
            can_npy_path,
            trajectory_can_npy_path,
            trajectory_cgn_can_npy_path,
            bundle_path,
        ]
        for path in intermediate_paths:
            try:
                Path(path).unlink(missing_ok=True)
            except OSError:
                pass

    print(f"SMILES: {smiles}")
    print(f"Saved base coords: {base_npy_path}")
    print(f"base coords shape: {base_coords_angstrom.shape}  # [num_atoms, 3]")
    print(f"Saved base CGN+CAN coords: {base_cgn_can_npy_path}")
    print(
        f"base CGN+CAN shape: {base_cgn_can_coords.shape}  # [num_views, num_atoms, 3], "
        f"rotatable_torsions={base_cgn_can_torsions}"
    )
    print(f"Saved core MD-CGN-CAN bundle: {core_bundle_path}")
    print(
        f"bundle trajectory CGN+CAN shape: {trajectory_cgn_can_coords.shape}  # [num_frames, num_atoms, 3]"
    )
    print(
        f"bundle trajectory electronic shape: {trajectory_cgn_can_electronic_feats.shape} "
        f"# [num_frames, num_atoms, 2]"
    )
    if not save_intermediates:
        print("Saved only 3 core files. Use --save-intermediates to keep PDB/DCD/log and separate ablation arrays.")
        return

    print(f"Saved PDB: {pdb_path}")
    print(f"Saved DCD: {dcd_path}")
    print(f"Saved log: {log_path}")
    print(f"Saved electronic feats: {electronic_npy_path}")
    print(f"electronic feats shape: {electronic_feats.shape}  # [num_atoms, 2] = [formal_charge, Gasteiger_charge]")
    print(f"Saved trajectory electronic feats: {trajectory_electronic_npy_path}")
    print(f"trajectory electronic feats shape: {trajectory_electronic_feats.shape}  # [num_frames, num_atoms, 2]")
    print(f"Saved formal charge: {formal_charge_npy_path}")
    print(f"Saved Gasteiger charge: {gasteiger_charge_npy_path}")
    print(f"Saved charge CSV: {charge_csv_path}")
    print(f"Saved coords: {npy_path}")
    print(f"coords shape: {coords_angstrom.shape}  # [num_frames, num_atoms, 3], unit=Angstrom")
    print(f"Saved CGN coords: {cgn_npy_path}")
    print(f"CGN shape: {cgn_coords.shape}  # [num_cgn_views, num_atoms, 3], sigma={cgn_sigma} Angstrom")
    print(f"Saved CGN electronic feats: {cgn_electronic_npy_path}")
    print(f"CGN electronic feats shape: {cgn_electronic_feats.shape}  # [num_cgn_views, num_atoms, 2]")
    if trajectory_cgn:
        print(f"Saved trajectory CGN coords: {trajectory_cgn_npy_path}")
        print(f"trajectory CGN shape: {trajectory_cgn_coords.shape}  # [num_frames, num_atoms, 3]")
        print(f"Saved trajectory CGN electronic feats: {trajectory_cgn_electronic_npy_path}")
    print(f"Saved CAN coords: {can_npy_path}")
    print(
        f"CAN shape: {can_coords.shape}  # [num_can_views, num_atoms, 3], "
        f"rotatable_torsions={num_torsions}, max_angle={can_max_angle_deg} deg"
    )
    print(f"Saved CAN electronic feats: {can_electronic_npy_path}")
    print(f"CAN electronic feats shape: {can_electronic_feats.shape}  # [num_can_views, num_atoms, 2]")
    if trajectory_can:
        print(f"Saved trajectory CAN coords: {trajectory_can_npy_path}")
        print(
            f"trajectory CAN shape: {trajectory_can_coords.shape}  # [num_frames, num_atoms, 3], "
            f"rotatable_torsions={trajectory_can_torsions}"
        )
        print(f"Saved trajectory CAN electronic feats: {trajectory_can_electronic_npy_path}")
    if trajectory_cgn_can:
        print(f"Saved trajectory CGN+CAN coords: {trajectory_cgn_can_npy_path}")
        print(
            f"trajectory CGN+CAN shape: {trajectory_cgn_can_coords.shape}  # [num_frames, num_atoms, 3], "
            f"rotatable_torsions={trajectory_cgn_can_torsions}, sigma={cgn_sigma} Angstrom"
        )
        print(f"Saved trajectory CGN+CAN electronic feats: {trajectory_cgn_can_electronic_npy_path}")
        print(
            f"trajectory CGN+CAN electronic feats shape: {trajectory_cgn_can_electronic_feats.shape} "
            f" # [num_frames, num_atoms, 2]"
        )
    print(f"Saved bundled arrays: {bundle_path}")


def build_arg_parser() -> argparse.ArgumentParser:
    parser = argparse.ArgumentParser(description="Generate a small-molecule OpenMM MD trajectory from SMILES.")
    parser.add_argument("--smiles", default="CCO", help="Input molecule SMILES.")
    parser.add_argument("--out-dir", default="md_test/ethanol", help="Output directory.")
    parser.add_argument("--steps", type=int, default=10000, help="Total MD steps.")
    parser.add_argument(
        "--equilibration-steps",
        type=int,
        default=1000,
        help="Thermalization steps run before trajectory reporters are attached.",
    )
    parser.add_argument("--report-interval", type=int, default=100, help="Save one frame every N steps.")
    parser.add_argument("--seed", type=int, default=2026)
    parser.add_argument("--temperature-k", type=float, default=300.0)
    parser.add_argument("--timestep-fs", type=float, default=1.0)
    parser.add_argument("--friction-per-ps", type=float, default=1.0)
    parser.add_argument("--platform", default="auto", help="auto, CPU, CUDA, OpenCL, etc.")
    parser.add_argument(
        "--partial-charge-method",
        default="gasteiger",
        choices=["gasteiger", "mmff94", "formal_charge", "zeros", "am1bcc", "default"],
        help=(
            "Partial charges used by OpenFF parameterization. "
            "gasteiger is fastest and avoids AmberTools AM1-BCC failures; "
            "am1bcc/default may be slower and can call antechamber/sqm."
        ),
    )
    parser.add_argument("--cgn-views", type=int, default=4, help="Number of Gaussian-noise conformer views from the minimized structure.")
    parser.add_argument("--cgn-sigma", type=float, default=0.02, help="CGN Gaussian sigma in Angstrom.")
    parser.add_argument(
        "--no-trajectory-cgn",
        action="store_true",
        help="Disable per-frame CGN noise on the saved MD trajectory.",
    )
    parser.add_argument("--can-views", type=int, default=4, help="Number of torsion-angle-noise conformer views.")
    parser.add_argument("--can-max-angle-deg", type=float, default=30.0, help="Maximum absolute torsion perturbation angle.")
    parser.add_argument(
        "--no-trajectory-can",
        action="store_true",
        help="Disable per-frame CAN torsion noise on the saved MD trajectory.",
    )
    parser.add_argument(
        "--no-trajectory-cgn-can",
        action="store_true",
        help="Disable per-frame combined CAN followed by CGN noise on the saved MD trajectory.",
    )
    parser.add_argument(
        "--save-intermediates",
        action="store_true",
        help="Keep PDB/DCD/log and separate CAN/CGN/trajectory arrays. By default only three core output files are kept.",
    )
    return parser


def main() -> None:
    args = build_arg_parser().parse_args()
    run_openmm_md(
        smiles=args.smiles,
        out_dir=args.out_dir,
        equilibration_steps=args.equilibration_steps,
        steps=args.steps,
        report_interval=args.report_interval,
        seed=args.seed,
        temperature_kelvin=args.temperature_k,
        timestep_fs=args.timestep_fs,
        friction_per_ps=args.friction_per_ps,
        platform_name=args.platform,
        cgn_views=args.cgn_views,
        cgn_sigma=args.cgn_sigma,
        trajectory_cgn=not args.no_trajectory_cgn,
        can_views=args.can_views,
        can_max_angle_deg=args.can_max_angle_deg,
        trajectory_can=not args.no_trajectory_can,
        trajectory_cgn_can=not args.no_trajectory_cgn_can,
        partial_charge_method=args.partial_charge_method,
        save_intermediates=args.save_intermediates,
    )


if __name__ == "__main__":
    main()
```
