## Supplementary material for "PG-MLD: Physics-Guided Molecular Representation Learning via Dynamic 3D Trajectory Distillation": Code and Data: plot_molecular_views.py

```
from __future__ import annotations

import argparse
from pathlib import Path
from typing import Sequence

import matplotlib

matplotlib.use("Agg")
import matplotlib.pyplot as plt
import numpy as np
from matplotlib import patheffects
from mpl_toolkits.mplot3d import Axes3D  # noqa: F401
from rdkit import Chem
from rdkit.Chem import AllChem, rdMolTransforms


ATOM_COLORS = {
    "H": "#E8E8E8",
    "C": "#6F767D",
    "N": "#4169E1",
    "O": "#E53935",
    "F": "#42C95A",
    "Cl": "#31C84B",
    "Br": "#8B3A2B",
    "I": "#7B4AB5",
    "P": "#F28C28",
    "S": "#E6C229",
}

ATOM_RADII = {
    "H": 22,
    "C": 58,
    "N": 62,
    "O": 64,
    "F": 60,
    "Cl": 82,
    "Br": 88,
    "I": 96,
    "P": 78,
    "S": 78,
}


def build_molecule(smiles: str, seed: int = 2026) -> Chem.Mol:
    mol = Chem.MolFromSmiles(smiles)
    if mol is None:
        raise ValueError(f"Invalid SMILES: {smiles}")
    mol = Chem.AddHs(mol)
    params = AllChem.ETKDGv3()
    params.randomSeed = int(seed)
    params.useRandomCoords = True
    if AllChem.EmbedMolecule(mol, params) != 0:
        raise RuntimeError(f"RDKit embedding failed for: {smiles}")
    try:
        AllChem.MMFFOptimizeMolecule(mol, maxIters=500)
    except Exception:
        AllChem.UFFOptimizeMolecule(mol, maxIters=500)
    return mol


def coordinates(mol: Chem.Mol) -> np.ndarray:
    conf = mol.GetConformer()
    return np.asarray(
        [
            [
                conf.GetAtomPosition(i).x,
                conf.GetAtomPosition(i).y,
                conf.GetAtomPosition(i).z,
            ]
            for i in range(mol.GetNumAtoms())
        ],
        dtype=np.float32,
    )


def set_coordinates(mol: Chem.Mol, coords: np.ndarray) -> Chem.Mol:
    work = Chem.Mol(mol)
    conf = work.GetConformer()
    for atom_idx, xyz in enumerate(np.asarray(coords, dtype=float)):
        conf.SetAtomPosition(atom_idx, tuple(float(value) for value in xyz))
    return work


def rotatable_torsions(mol: Chem.Mol) -> list[tuple[int, int, int, int]]:
    torsions: list[tuple[int, int, int, int]] = []
    for bond in mol.GetBonds():
        if bond.GetBondType() != Chem.rdchem.BondType.SINGLE or bond.IsInRing():
            continue
        begin, end = bond.GetBeginAtom(), bond.GetEndAtom()
        if begin.GetAtomicNum() == 1 or end.GetAtomicNum() == 1:
            continue
        left = [a.GetIdx() for a in begin.GetNeighbors() if a.GetIdx() != end.GetIdx()]
        right = [a.GetIdx() for a in end.GetNeighbors() if a.GetIdx() != begin.GetIdx()]
        if left and right:
            torsions.append((left[0], begin.GetIdx(), end.GetIdx(), right[0]))
    return torsions


def make_cgn(coords: np.ndarray, sigma: float, seed: int) -> np.ndarray:
    rng = np.random.default_rng(seed)
    return np.asarray(coords) + rng.normal(0.0, sigma, size=np.asarray(coords).shape)


def make_can(
    mol: Chem.Mol,
    coords: np.ndarray,
    max_angle_deg: float,
    seed: int,
) -> np.ndarray:
    work = set_coordinates(mol, coords)
    conf = work.GetConformer()
    rng = np.random.default_rng(seed)
    for i, j, k, l in rotatable_torsions(work):
        angle = rdMolTransforms.GetDihedralDeg(conf, i, j, k, l)
        delta = rng.uniform(-max_angle_deg, max_angle_deg)
        rdMolTransforms.SetDihedralDeg(conf, i, j, k, l, angle + float(delta))
    return coordinates(work)


def make_synthetic_trajectory(
    mol: Chem.Mol,
    base_coords: np.ndarray,
    num_frames: int,
    max_angle_deg: float,
    sigma: float,
    seed: int,
) -> np.ndarray:
    frames = []
    for frame_idx in range(num_frames):
        can = make_can(
            mol,
            base_coords,
            max_angle_deg=max_angle_deg,
            seed=seed + frame_idx * 17,
        )
        frames.append(make_cgn(can, sigma=sigma, seed=seed + frame_idx * 31))
    return np.stack(frames).astype(np.float32)


def load_trajectory(path: str | Path, item_index: int = 0) -> tuple[str | None, np.ndarray]:
    path = Path(path)
    if path.suffix.lower() == ".npy":
        return None, np.load(path, allow_pickle=False)
    if path.suffix.lower() == ".npz":
        with np.load(path, allow_pickle=False) as data:
            preferred = [
                "trajectory_coords_angstrom",
                "trajectory_cgn_can_coords_angstrom",
                "coords",
            ]
            key = next((candidate for candidate in preferred if candidate in data), None)
            if key is None:
                raise KeyError(f"No trajectory array found in {path}. Keys: {list(data.keys())}")
            return None, np.asarray(data[key])
    if path.suffix.lower() == ".pt":
        import torch

        try:
            items = torch.load(path, map_location="cpu", weights_only=False)
        except TypeError:
            items = torch.load(path, map_location="cpu")
        item = items[item_index] if isinstance(items, list) else items
        coords = item["coords"]
        if hasattr(coords, "detach"):
            coords = coords.detach().cpu().numpy()
        return str(item.get("smiles", "")) or None, np.asarray(coords)
    raise ValueError(f"Unsupported trajectory file: {path}")


def normalization_reference(coords: np.ndarray) -> tuple[np.ndarray, float]:
    coords = np.asarray(coords, dtype=float)
    center = coords.mean(axis=0)
    scale = np.max(np.linalg.norm(coords - center, axis=1))
    return center, max(float(scale), 1e-6)


def normalize_coordinates(
    coords: np.ndarray,
    center: np.ndarray | None = None,
    scale: float | None = None,
) -> np.ndarray:
    coords = np.asarray(coords, dtype=float)
    if center is None or scale is None:
        center, scale = normalization_reference(coords)
    return (coords - np.asarray(center)) / max(float(scale), 1e-6)


def draw_bond(ax, p1: np.ndarray, p2: np.ndarray, color: str = "#9AA0A6") -> None:
    ax.plot(
        [p1[0], p2[0]],
        [p1[1], p2[1]],
        [p1[2], p2[2]],
        color=color,
        linewidth=2.2,
        solid_capstyle="round",
        alpha=0.95,
        zorder=1,
    )


def draw_molecule(
    ax,
    mol: Chem.Mol,
    coords: np.ndarray,
    elevation: float = 18,
    azimuth: float = -68,
    center: np.ndarray | None = None,
    scale: float | None = None,
    alpha: float = 1.0,
) -> None:
    xyz = normalize_coordinates(coords, center=center, scale=scale)
    for bond in mol.GetBonds():
        begin, end = bond.GetBeginAtomIdx(), bond.GetEndAtomIdx()
        ax.plot(
            [xyz[begin, 0], xyz[end, 0]],
            [xyz[begin, 1], xyz[end, 1]],
            [xyz[begin, 2], xyz[end, 2]],
            color="#9AA0A6",
            linewidth=2.2,
            solid_capstyle="round",
            alpha=0.95 * alpha,
            zorder=1,
        )

    # Plot larger atoms later to obtain cleaner ball-and-stick occlusion.
    order = sorted(range(mol.GetNumAtoms()), key=lambda i: mol.GetAtomWithIdx(i).GetAtomicNum())
    for atom_idx in order:
        symbol = mol.GetAtomWithIdx(atom_idx).GetSymbol()
        color = ATOM_COLORS.get(symbol, "#C77CFF")
        size = ATOM_RADII.get(symbol, 65)
        point = xyz[atom_idx]
        artist = ax.scatter(
            point[0],
            point[1],
            point[2],
            s=size,
            c=color,
            edgecolors="#454545",
            linewidths=0.35,
            depthshade=True,
            alpha=alpha,
            zorder=2,
        )
        artist.set_path_effects([patheffects.withStroke(linewidth=0.25, foreground="white")])

    ax.view_init(elev=elevation, azim=azimuth)
    ax.set_xlim(-1.15, 1.15)
    ax.set_ylim(-1.15, 1.15)
    ax.set_zlim(-1.15, 1.15)
    ax.set_box_aspect((1, 1, 1))
    ax.set_axis_off()
    ax.set_facecolor("white")


def select_frame_indices(num_frames: int, panels: int) -> list[int]:
    if num_frames <= panels:
        return list(range(num_frames))
    return np.linspace(0, num_frames - 1, panels, dtype=int).tolist()


def save_figure(fig, output: str | Path, dpi: int) -> list[Path]:
    output = Path(output)
    output.parent.mkdir(parents=True, exist_ok=True)
    stem = output.with_suffix("")
    paths = [stem.with_suffix(".png"), stem.with_suffix(".pdf"), stem.with_suffix(".svg")]
    fig.savefig(paths[0], dpi=dpi, bbox_inches="tight", pad_inches=0.04, facecolor="white")
    fig.savefig(paths[1], bbox_inches="tight", pad_inches=0.04, facecolor="white")
    fig.savefig(paths[2], bbox_inches="tight", pad_inches=0.04, facecolor="white")
    return paths


def plot_views(
    mol: Chem.Mol,
    eq: np.ndarray,
    cgn: np.ndarray,
    can: np.ndarray,
    output: str | Path,
    dpi: int,
    cgn_sigma: float,
    cgn_display_scale: float,
) -> list[Path]:
    plt.rcParams.update({"font.family": "serif", "mathtext.fontset": "stix"})
    fig = plt.figure(figsize=(9.0, 3.2), constrained_layout=True)
    labels = [
        (r"$X^{\mathrm{eq}}$", "equilibrium"),
        (
            r"$X^{\mathrm{CGN}}$",
            rf"coord.-noise, $\sigma={cgn_sigma:.2f}\,\AA$",
        ),
        (r"$X^{\mathrm{CAN}}$", "torsion-pert."),
    ]
    common_center, common_scale = normalization_reference(eq)
    for panel_idx, (coords, (math_label, subtitle)) in enumerate(zip([eq, cgn, can], labels), start=1):
        ax = fig.add_subplot(1, 3, panel_idx, projection="3d")
        if panel_idx == 2:
            # Show the true equilibrium structure as a faint reference. The
            # displacement arrows are enlarged only for legibility.
            draw_molecule(
                ax,
                mol,
                eq,
                center=common_center,
                scale=common_scale,
                alpha=0.16,
            )
            draw_molecule(
                ax,
                mol,
                cgn,
                center=common_center,
                scale=common_scale,
            )
            eq_xyz = normalize_coordinates(eq, common_center, common_scale)
            cgn_xyz = normalize_coordinates(cgn, common_center, common_scale)
            displacement = (cgn_xyz - eq_xyz) * float(cgn_display_scale)
            for atom_idx, start in enumerate(eq_xyz):
                delta = displacement[atom_idx]
                ax.quiver(
                    start[0],
                    start[1],
                    start[2],
                    delta[0],
                    delta[1],
                    delta[2],
                    color="#F28C28",
                    linewidth=1.0,
                    arrow_length_ratio=0.35,
                    alpha=0.85,
                    normalize=False,
                )
        else:
            draw_molecule(
                ax,
                mol,
                coords,
                center=common_center,
                scale=common_scale,
            )
        ax.text2D(
            0.5,
            1.02,
            math_label,
            transform=ax.transAxes,
            ha="center",
            va="bottom",
            fontsize=18,
            fontweight="bold",
        )
        ax.text2D(
            0.5,
            0.95,
            subtitle,
            transform=ax.transAxes,
            ha="center",
            va="bottom",
            fontsize=12,
        )
        if panel_idx == 2 and cgn_display_scale != 1.0:
            ax.text2D(
                0.5,
                0.895,
                rf"orange displacement arrows $\times{cgn_display_scale:g}$",
                transform=ax.transAxes,
                ha="center",
                va="bottom",
                fontsize=9,
                color="#C86800",
            )
    return save_figure(fig, output, dpi)


def plot_trajectory(
    mol: Chem.Mol,
    trajectory: np.ndarray,
    output: str | Path,
    panels: int,
    dpi: int,
) -> list[Path]:
    plt.rcParams.update({"font.family": "serif", "mathtext.fontset": "stix"})
    frame_indices = select_frame_indices(len(trajectory), panels)
    fig = plt.figure(figsize=(2.7 * panels, 2.65), constrained_layout=True)
    for panel_idx, frame_idx in enumerate(frame_indices, start=1):
        ax = fig.add_subplot(1, panels, panel_idx, projection="3d")
        draw_molecule(ax, mol, trajectory[frame_idx])
        if panel_idx == 1:
            label = r"$t_1$"
        elif panel_idx == panels:
            label = r"$t_T$"
        else:
            label = rf"$t_{{{panel_idx}}}$"
        ax.text2D(
            0.5,
            -0.015,
            label,
            transform=ax.transAxes,
            ha="center",
            va="top",
            fontsize=17,
            fontweight="bold",
        )
        if panel_idx < panels:
            fig.text(
                panel_idx / panels,
                0.50,
                r"$\longrightarrow$" if panel_idx != panels - 1 else r"$\cdots$",
                ha="center",
                va="center",
                fontsize=22,
            )
    return save_figure(fig, output, dpi)


def build_arg_parser() -> argparse.ArgumentParser:
    parser = argparse.ArgumentParser(
        description="Generate publication-quality molecular trajectory and EQ/CGN/CAN panels."
    )
    parser.add_argument(
        "--smiles",
        help="SMILES used to define atoms and bonds. Optional when a packed .pt item contains SMILES.",
    )
    parser.add_argument(
        "--mode",
        choices=["views", "trajectory", "both"],
        default="both",
    )
    parser.add_argument("--trajectory", help="Optional .pt/.npy/.npz trajectory file.")
    parser.add_argument("--item-index", type=int, default=0, help="Molecule index for a packed .pt file.")
    parser.add_argument("--output-dir", default="output/figures/molecular_views")
    parser.add_argument("--prefix", default="molecule")
    parser.add_argument("--panels", type=int, default=4)
    parser.add_argument("--cgn-sigma", type=float, default=0.02)
    parser.add_argument(
        "--cgn-display-scale",
        type=float,
        default=10.0,
        help="Visual-only scale factor for CGN displacement arrows; coordinates remain unchanged.",
    )
    parser.add_argument("--can-angle", type=float, default=30.0)
    parser.add_argument("--seed", type=int, default=2026)
    parser.add_argument("--dpi", type=int, default=600)
    return parser


def main() -> None:
    args = build_arg_parser().parse_args()
    trajectory = None
    trajectory_smiles = None
    if args.trajectory:
        trajectory_smiles, trajectory = load_trajectory(args.trajectory, item_index=args.item_index)
    smiles = args.smiles or trajectory_smiles
    if not smiles:
        raise ValueError("--smiles is required unless --trajectory is a packed .pt containing SMILES.")

    mol = build_molecule(smiles, seed=args.seed)
    eq = coordinates(mol)
    cgn = make_cgn(eq, sigma=args.cgn_sigma, seed=args.seed + 1)
    can = make_can(mol, eq, max_angle_deg=args.can_angle, seed=args.seed + 2)

    if args.trajectory:
        if trajectory_smiles and args.smiles and trajectory_smiles != args.smiles:
            raise ValueError(
                f"Provided SMILES does not match packed trajectory item: {trajectory_smiles}"
            )
        if trajectory.shape[1] != mol.GetNumAtoms():
            heavy_mol = Chem.RemoveHs(mol)
            if trajectory.shape[1] == heavy_mol.GetNumAtoms():
                mol = heavy_mol
                eq = coordinates(mol)
                cgn = make_cgn(eq, sigma=args.cgn_sigma, seed=args.seed + 1)
                can = make_can(mol, eq, max_angle_deg=args.can_angle, seed=args.seed + 2)
            else:
                raise ValueError(
                    f"Trajectory atoms={trajectory.shape[1]}, molecule atoms={mol.GetNumAtoms()}"
                )
    else:
        trajectory = make_synthetic_trajectory(
            mol,
            eq,
            num_frames=max(args.panels, 4),
            max_angle_deg=args.can_angle,
            sigma=args.cgn_sigma,
            seed=args.seed + 10,
        )

    output_dir = Path(args.output_dir)
    generated: list[Path] = []
    if args.mode in {"views", "both"}:
        generated += plot_views(
            mol,
            eq,
            cgn,
            can,
            output_dir / f"{args.prefix}_eq_cgn_can.png",
            dpi=args.dpi,
            cgn_sigma=args.cgn_sigma,
            cgn_display_scale=args.cgn_display_scale,
        )
    if args.mode in {"trajectory", "both"}:
        generated += plot_trajectory(
            mol,
            trajectory,
            output_dir / f"{args.prefix}_trajectory.png",
            panels=args.panels,
            dpi=args.dpi,
        )
    print("Generated:")
    for path in generated:
        print(f"  {path}")


if __name__ == "__main__":
    main()
```
