## Supplementary material for "PG-MLD: Physics-Guided Molecular Representation Learning via Dynamic 3D Trajectory Distillation": Code and Data: prepare_qm9.py

```
from __future__ import annotations

import argparse
import json
import shutil
import urllib.request
import zipfile
from pathlib import Path

import numpy as np
import pandas as pd
from tqdm import tqdm


QM9_ARCHIVE_URL = (
    "https://deepchemdata.s3-us-west-1.amazonaws.com/datasets/molnet_publish/qm9.zip"
)
QM9_UNCHARACTERIZED_URL = "https://ndownloader.figshare.com/files/3195404"
HARTREE_TO_EV = 27.211386246
TARGET_COLUMNS = ("mu", "alpha", "homo", "lumo", "gap", "zpve", "u0", "u", "h")
SOURCE_ALIASES = {
    "mu": ("mu",),
    "alpha": ("alpha",),
    "homo": ("homo",),
    "lumo": ("lumo",),
    "gap": ("gap",),
    "zpve": ("zpve",),
    "u0": ("u0", "U0"),
    "u": ("u298", "U"),
    "h": ("h298", "H"),
}
ENERGY_TARGETS = ("homo", "lumo", "gap", "zpve", "u0", "u", "h")


def download(url: str, destination: Path) -> None:
    destination.parent.mkdir(parents=True, exist_ok=True)
    temporary = destination.with_suffix(destination.suffix + ".part")
    print(f"Downloading {url}")
    try:
        with urllib.request.urlopen(url) as response, temporary.open("wb") as handle:
            total = int(response.headers.get("Content-Length", "0") or 0)
            with tqdm(total=total, unit="B", unit_scale=True, desc=destination.name) as progress:
                while True:
                    block = response.read(1024 * 1024)
                    if not block:
                        break
                    handle.write(block)
                    progress.update(len(block))
        temporary.replace(destination)
    except BaseException:
        temporary.unlink(missing_ok=True)
        raise


def ensure_raw_files(raw_dir: Path, allow_download: bool) -> tuple[Path, Path, Path]:
    archive = raw_dir / "qm9.zip"
    sdf_path = raw_dir / "gdb9.sdf"
    target_csv = raw_dir / "gdb9.sdf.csv"
    uncharacterized = raw_dir / "uncharacterized.txt"
    raw_dir.mkdir(parents=True, exist_ok=True)

    if not sdf_path.exists() or not target_csv.exists():
        if not archive.exists():
            if not allow_download:
                raise FileNotFoundError(
                    f"Missing {archive}. Re-run without --no-download or place the official archive there."
                )
            download(QM9_ARCHIVE_URL, archive)
        print(f"Extracting {archive} -> {raw_dir}")
        with zipfile.ZipFile(archive) as handle:
            wanted = {"gdb9.sdf", "gdb9.sdf.csv"}
            members = {Path(name).name: name for name in handle.namelist()}
            missing = wanted.difference(members)
            if missing:
                raise ValueError(f"QM9 archive is missing {sorted(missing)}")
            for filename in wanted:
                source = handle.open(members[filename])
                with source, (raw_dir / filename).open("wb") as destination:
                    shutil.copyfileobj(source, destination)

    if not uncharacterized.exists():
        if not allow_download:
            raise FileNotFoundError(
                f"Missing {uncharacterized}. Re-run without --no-download or provide the official file."
            )
        download(QM9_UNCHARACTERIZED_URL, uncharacterized)
    return sdf_path, target_csv, uncharacterized


def uncharacterized_indices(path: Path) -> set[int]:
    lines = path.read_text(encoding="utf-8", errors="replace").splitlines()
    # This follows the official PyG QM9 preprocessing: data rows occupy [9:-2],
    # and molecule identifiers in the file are one-based.
    indices: set[int] = set()
    for line in lines[9:-2]:
        fields = line.split()
        if not fields:
            continue
        try:
            indices.add(int(fields[0]) - 1)
        except ValueError:
            continue
    if len(indices) != 3054:
        raise ValueError(
            f"Expected 3054 uncharacterized QM9 molecules, parsed {len(indices)} from {path}."
        )
    return indices


def resolve_source_columns(frame: pd.DataFrame) -> dict[str, str]:
    lower_to_original = {str(column).lower(): str(column) for column in frame.columns}
    resolved: dict[str, str] = {}
    for canonical, aliases in SOURCE_ALIASES.items():
        for alias in aliases:
            source = lower_to_original.get(alias.lower())
            if source is not None:
                resolved[canonical] = source
                break
        if canonical not in resolved:
            raise ValueError(
                f"Cannot find source column for {canonical!r}. Available columns: {list(frame.columns)}"
            )
    return resolved


def smiles_from_sdf(sdf_path: Path, skip: set[int], expected_rows: int) -> tuple[list[int], list[str]]:
    try:
        from rdkit import Chem, RDLogger
    except ImportError as exc:
        raise ImportError("RDKit is required to extract canonical SMILES from the official QM9 SDF.") from exc

    RDLogger.DisableLog("rdApp.*")
    supplier = Chem.SDMolSupplier(str(sdf_path), removeHs=False, sanitize=True)
    kept_indices: list[int] = []
    smiles: list[str] = []
    for index, molecule in enumerate(tqdm(supplier, total=expected_rows, desc="Canonicalize QM9 SMILES")):
        if index in skip:
            continue
        if molecule is None:
            raise ValueError(f"RDKit failed to parse characterized QM9 molecule at source index {index}.")
        heavy_atom_molecule = Chem.RemoveHs(molecule)
        canonical = Chem.MolToSmiles(heavy_atom_molecule, isomericSmiles=True, canonical=True)
        if not canonical:
            raise ValueError(f"Empty canonical SMILES at source index {index}.")
        kept_indices.append(index)
        smiles.append(canonical)

    if len(kept_indices) != expected_rows - len(skip):
        raise ValueError(
            f"Expected {expected_rows - len(skip)} characterized molecules, got {len(kept_indices)}."
        )
    return kept_indices, smiles


def make_fixed_split(
    count: int,
    train_size: int,
    val_size: int,
    seed: int,
) -> tuple[np.ndarray, np.ndarray, np.ndarray]:
    if train_size <= 0 or val_size <= 0 or train_size + val_size >= count:
        raise ValueError(
            f"Invalid split sizes: count={count}, train={train_size}, val={val_size}"
        )
    # RandomState matches the long-standing sklearn/DimeNet QM9 split convention.
    permutation = np.random.RandomState(seed).permutation(count)
    train = permutation[:train_size].astype(np.int64)
    val = permutation[train_size : train_size + val_size].astype(np.int64)
    test = permutation[train_size + val_size :].astype(np.int64)
    return train, val, test


def atomic_csv_gzip(frame: pd.DataFrame, destination: Path) -> None:
    destination.parent.mkdir(parents=True, exist_ok=True)
    temporary = destination.with_suffix(destination.suffix + ".tmp")
    try:
        frame.to_csv(temporary, index=False, compression="gzip")
        temporary.replace(destination)
    except BaseException:
        temporary.unlink(missing_ok=True)
        raise


def atomic_npz(destination: Path, **arrays: np.ndarray) -> None:
    destination.parent.mkdir(parents=True, exist_ok=True)
    temporary = destination.with_suffix(destination.suffix + ".tmp")
    try:
        with temporary.open("wb") as handle:
            np.savez_compressed(handle, **arrays)
        temporary.replace(destination)
    except BaseException:
        temporary.unlink(missing_ok=True)
        raise


def prepare(args: argparse.Namespace) -> None:
    raw_dir = Path(args.raw_dir)
    sdf_path, target_csv, uncharacterized_path = ensure_raw_files(
        raw_dir,
        allow_download=not args.no_download,
    )

    source = pd.read_csv(target_csv)
    resolved = resolve_source_columns(source)
    skip = uncharacterized_indices(uncharacterized_path)
    source_indices, smiles = smiles_from_sdf(sdf_path, skip, expected_rows=len(source))

    selected = source.iloc[source_indices].reset_index(drop=True)
    output = pd.DataFrame(
        {
            "source_index": np.asarray(source_indices, dtype=np.int64),
            "mol_id": selected.iloc[:, 0].astype(str),
            "smiles": smiles,
        }
    )
    for target in TARGET_COLUMNS:
        values = pd.to_numeric(selected[resolved[target]], errors="coerce").to_numpy(dtype=np.float64)
        if target in ENERGY_TARGETS:
            values = values * HARTREE_TO_EV
        if not np.isfinite(values).all():
            bad = int((~np.isfinite(values)).sum())
            raise ValueError(f"Target {target} contains {bad} non-finite values.")
        output[target] = values

    if len(output) != 130831:
        raise ValueError(
            f"Expected the standard 130831 characterized QM9 molecules, got {len(output)}."
        )

    output_path = Path(args.output)
    split_path = Path(args.split_output)
    atomic_csv_gzip(output, output_path)
    train, val, test = make_fixed_split(
        len(output),
        train_size=args.train_size,
        val_size=args.val_size,
        seed=args.seed,
    )
    atomic_npz(split_path, train=train, val=val, test=test)

    metadata = {
        "source_archive_url": QM9_ARCHIVE_URL,
        "uncharacterized_url": QM9_UNCHARACTERIZED_URL,
        "molecules": len(output),
        "excluded_uncharacterized": len(skip),
        "targets": list(TARGET_COLUMNS),
        "stored_units": {
            "mu": "D",
            "alpha": "a0^3",
            "homo": "eV",
            "lumo": "eV",
            "gap": "eV",
            "zpve": "eV",
            "u0": "eV",
            "u": "eV",
            "h": "eV",
        },
        "split": {
            "method": "fixed_random",
            "seed": args.seed,
            "train": len(train),
            "val": len(val),
            "test": len(test),
        },
    }
    metadata_path = output_path.with_suffix("").with_suffix(".metadata.json")
    metadata_path.write_text(json.dumps(metadata, indent=2), encoding="utf-8")

    print(f"Saved QM9 CSV: {output_path}")
    print(f"Saved fixed split: {split_path}")
    print(f"Saved metadata: {metadata_path}")
    print(
        f"QM9: total={len(output)} train={len(train)} val={len(val)} test={len(test)} "
        f"seed={args.seed}"
    )
    print("Stored energy units: eV; evaluation reports HOMO/LUMO/Gap/ZPVE/U0/U/H in meV.")


def build_arg_parser() -> argparse.ArgumentParser:
    parser = argparse.ArgumentParser(
        description="Prepare the standard characterized QM9 SMILES regression benchmark."
    )
    parser.add_argument("--raw-dir", default="data/qm9/raw")
    parser.add_argument("--output", default="output/molecular/reg/qm9/qm9.csv.gz")
    parser.add_argument(
        "--split-output",
        default="output/molecular/reg/qm9/split/random_seed42_110k_10k.npz",
    )
    parser.add_argument("--train-size", type=int, default=110000)
    parser.add_argument("--val-size", type=int, default=10000)
    parser.add_argument("--seed", type=int, default=42)
    parser.add_argument(
        "--no-download",
        action="store_true",
        help="Require the official raw files to already exist in --raw-dir.",
    )
    return parser


def main() -> None:
    prepare(build_arg_parser().parse_args())


if __name__ == "__main__":
    main()
```
