## Supplementary material for "PG-MLD: Physics-Guided Molecular Representation Learning via Dynamic 3D Trajectory Distillation": Code and Data: preprocess_zinc_openmm_md.py

```
from __future__ import annotations

import argparse
import contextlib
import csv
import os
import signal
import sys
import tempfile
import time
import traceback
from pathlib import Path
from typing import Iterator

try:
    from tqdm import tqdm
except ImportError:
    def tqdm(iterable, total=None, desc=None):
        return iterable


CORE_FILES = (
    "base_coords_angstrom.npy",
    "base_cgn_can_coords_angstrom.npy",
    "md_cgn_can_with_electronic.npz",
)


@contextlib.contextmanager
def time_limit(seconds: float | None):
    if seconds is None or float(seconds) <= 0 or not hasattr(signal, "SIGALRM"):
        yield
        return

    def _raise_timeout(signum, frame):
        raise TimeoutError(f"OpenMM MD generation exceeded {seconds} seconds")

    old_handler = signal.getsignal(signal.SIGALRM)
    try:
        signal.signal(signal.SIGALRM, _raise_timeout)
        signal.setitimer(signal.ITIMER_REAL, float(seconds))
        yield
    finally:
        signal.setitimer(signal.ITIMER_REAL, 0)
        signal.signal(signal.SIGALRM, old_handler)


@contextlib.contextmanager
def maybe_silence(verbose: bool):
    if verbose:
        yield
        return
    with open(os.devnull, "w", encoding="utf-8") as devnull:
        stdout_fd = os.dup(1)
        stderr_fd = os.dup(2)
        try:
            sys.stdout.flush()
            sys.stderr.flush()
            os.dup2(devnull.fileno(), 1)
            os.dup2(devnull.fileno(), 2)
            with contextlib.redirect_stdout(devnull), contextlib.redirect_stderr(devnull):
                yield
        finally:
            sys.stdout.flush()
            sys.stderr.flush()
            os.dup2(stdout_fd, 1)
            os.dup2(stderr_fd, 2)
            os.close(stdout_fd)
            os.close(stderr_fd)


def iter_smiles(
    csv_path: Path,
    smiles_col: str,
    start_index: int = 0,
    max_molecules: int | None = None,
) -> Iterator[tuple[int, str]]:
    start_index = max(int(start_index), 0)
    yielded = 0
    with csv_path.open("r", encoding="utf-8-sig", newline="") as handle:
        reader = csv.DictReader(handle)
        if reader.fieldnames is None or smiles_col not in reader.fieldnames:
            raise ValueError(f"Column '{smiles_col}' not found in {csv_path}. Columns: {reader.fieldnames}")
        for row_idx, row in enumerate(reader):
            if row_idx < start_index:
                continue
            if max_molecules is not None and yielded >= int(max_molecules):
                break
            smiles = str(row.get(smiles_col, "")).strip()
            if smiles:
                yielded += 1
                yield row_idx, smiles


def heavy_atom_count(smiles: str) -> int | None:
    from rdkit import Chem

    mol = Chem.MolFromSmiles(smiles)
    if mol is None:
        return None
    return int(mol.GetNumHeavyAtoms())


def load_run_openmm_md():
    try:
        from generate_openmm_md import run_openmm_md
    except ImportError:
        from scripts.generate_openmm_md import run_openmm_md
    return run_openmm_md


def molecule_out_dir(out_root: Path, row_idx: int) -> Path:
    return out_root / f"{int(row_idx):06d}"


def has_core_outputs(out_dir: Path) -> bool:
    return all((out_dir / name).exists() for name in CORE_FILES)


def default_pack_paths(out_root: Path) -> dict[str, Path]:
    return {
        "base": out_root / "base.pt",
        "base_cgn_can": out_root / "base_cgn_can.pt",
        "md_cgn_can_electronic": out_root / "md_cgn_can_electronic.pt",
    }


def load_packed_views(out_dir: Path, record: dict) -> tuple[dict, dict, dict]:
    import numpy as np
    import torch

    base_coords = np.load(out_dir / "base_coords_angstrom.npy", allow_pickle=False)
    base_cgn_can_coords = np.load(out_dir / "base_cgn_can_coords_angstrom.npy", allow_pickle=False)
    with np.load(out_dir / "md_cgn_can_with_electronic.npz", allow_pickle=False) as bundle:
        common = {
            "row_idx": int(record["row_idx"]),
            "smiles": str(record.get("smiles", "")),
            "num_heavy_atoms": (
                int(record["num_heavy_atoms"]) if str(record.get("num_heavy_atoms", "")).strip() else None
            ),
            "atom_symbols": [str(value) for value in bundle["atom_symbols"].tolist()],
        }
        base_item = {
            **common,
            "coords": torch.tensor(base_coords, dtype=torch.float),
        }
        base_cgn_can_item = {
            **common,
            "coords": torch.tensor(base_cgn_can_coords, dtype=torch.float),
        }
        md_item = {
            **common,
            "coords": torch.tensor(bundle["trajectory_cgn_can_coords_angstrom"], dtype=torch.float),
            "electronic_feats": torch.tensor(bundle["electronic_feats"], dtype=torch.float),
            "trajectory_electronic_feats": torch.tensor(bundle["trajectory_cgn_can_electronic_feats"], dtype=torch.float),
            "formal_charge": torch.tensor(bundle["formal_charge"], dtype=torch.float),
            "gasteiger_charge": torch.tensor(bundle["gasteiger_charge"], dtype=torch.float),
        }
        if "cgn_sigma" in bundle:
            base_cgn_can_item["cgn_sigma"] = float(bundle["cgn_sigma"])
            md_item["cgn_sigma"] = float(bundle["cgn_sigma"])
        if "can_max_angle_deg" in bundle:
            base_cgn_can_item["can_max_angle_deg"] = float(bundle["can_max_angle_deg"])
            md_item["can_max_angle_deg"] = float(bundle["can_max_angle_deg"])
        if "timestep_fs" in bundle:
            md_item["timestep_fs"] = float(bundle["timestep_fs"])
        if "report_interval" in bundle:
            md_item["report_interval"] = int(bundle["report_interval"])
    return base_item, base_cgn_can_item, md_item


def write_summary_header(path: Path) -> None:
    with path.open("w", encoding="utf-8", newline="") as handle:
        writer = csv.DictWriter(
            handle,
            fieldnames=[
                "row_idx",
                "status",
                "smiles",
                "out_dir",
                "num_heavy_atoms",
                "seconds",
                "reason",
            ],
        )
        writer.writeheader()


def append_summary(path: Path, record: dict) -> None:
    with path.open("a", encoding="utf-8", newline="") as handle:
        writer = csv.DictWriter(
            handle,
            fieldnames=[
                "row_idx",
                "status",
                "smiles",
                "out_dir",
                "num_heavy_atoms",
                "seconds",
                "reason",
            ],
        )
        writer.writerow(record)


def write_run_report(
    path: Path,
    out_root: Path,
    summary_path: Path,
    pack_paths: dict[str, Path],
    packed_count: int,
    ok_count: int,
    skipped_count: int,
    failed_count: int,
    total_count: int,
    started_at: float,
    finished_at: float,
) -> None:
    elapsed = finished_at - started_at
    lines = [
        "ZINC OpenMM-MD preprocessing report",
        f"output_root: {out_root}",
        f"summary_csv: {summary_path}",
        f"base_pt: {pack_paths['base']}",
        f"base_cgn_can_pt: {pack_paths['base_cgn_can']}",
        f"md_cgn_can_electronic_pt: {pack_paths['md_cgn_can_electronic']}",
        f"packed_count: {packed_count}",
        f"total: {total_count}",
        f"ok: {ok_count}",
        f"skipped: {skipped_count}",
        f"failed: {failed_count}",
        f"elapsed_seconds: {elapsed:.3f}",
    ]
    path.write_text("\n".join(lines) + "\n", encoding="utf-8")


def read_summary_rows(summary_path: Path) -> list[dict[str, str]]:
    with summary_path.open("r", encoding="utf-8-sig", newline="") as handle:
        return list(csv.DictReader(handle))


def save_three_pt(
    pack_paths: dict[str, Path],
    base_items: list[dict],
    base_cgn_can_items: list[dict],
    md_items: list[dict],
) -> int:
    import torch

    for path in pack_paths.values():
        path.parent.mkdir(parents=True, exist_ok=True)
    torch.save(base_items, pack_paths["base"])
    torch.save(base_cgn_can_items, pack_paths["base_cgn_can"])
    torch.save(md_items, pack_paths["md_cgn_can_electronic"])
    return len(md_items)


def pack_successful_outputs(out_root: Path, summary_path: Path, pack_paths: dict[str, Path]) -> int:
    base_items = []
    base_cgn_can_items = []
    md_items = []
    for row in read_summary_rows(summary_path):
        if row.get("status") not in {"ok", "skipped"}:
            continue
        out_dir = Path(row.get("out_dir", ""))
        if not out_dir.is_absolute():
            out_dir = Path(out_dir)
        if not has_core_outputs(out_dir):
            continue

        base_item, base_cgn_can_item, md_item = load_packed_views(out_dir, row)
        base_item["out_dir"] = str(out_dir)
        base_cgn_can_item["out_dir"] = str(out_dir)
        md_item["out_dir"] = str(out_dir)
        base_items.append(base_item)
        base_cgn_can_items.append(base_cgn_can_item)
        md_items.append(md_item)

    return save_three_pt(pack_paths, base_items, base_cgn_can_items, md_items)


def preprocess_zinc_openmm_md(
    csv_path: str | Path,
    out_root: str | Path,
    smiles_col: str = "smiles",
    start_index: int = 0,
    max_molecules: int | None = 1000,
    max_atoms: int | None = 80,
    molecule_timeout_sec: float | None = 180.0,
    equilibration_steps: int = 1000,
    steps: int = 1000,
    report_interval: int = 100,
    platform: str = "OpenCL",
    seed: int = 2026,
    temperature_kelvin: float = 300.0,
    timestep_fs: float = 1.0,
    friction_per_ps: float = 1.0,
    partial_charge_method: str = "gasteiger",
    cgn_views: int = 4,
    cgn_sigma: float = 0.02,
    can_views: int = 4,
    can_max_angle_deg: float = 30.0,
    keep_folders: bool = False,
    force: bool = False,
    save_intermediates: bool = False,
    verbose: bool = False,
) -> None:
    csv_path = Path(csv_path)
    out_root = Path(out_root)
    out_root.mkdir(parents=True, exist_ok=True)
    summary_path = out_root / "summary.csv"
    report_path = out_root / "run_report.txt"
    pack_paths = default_pack_paths(out_root)
    write_summary_header(summary_path)

    rows = list(
        iter_smiles(
            csv_path,
            smiles_col=smiles_col,
            start_index=start_index,
            max_molecules=max_molecules,
        )
    )
    ok_count = 0
    skipped_count = 0
    failed_count = 0
    base_items = []
    base_cgn_can_items = []
    md_items = []
    started_at = time.time()
    run_openmm_md = load_run_openmm_md()

    for row_idx, smiles in tqdm(rows, total=len(rows), desc="ZINC OpenMM MD"):
        start = time.time()
        out_dir = molecule_out_dir(out_root, row_idx) if keep_folders else None
        if out_dir is not None:
            out_dir.mkdir(parents=True, exist_ok=True)
        num_heavy_atoms = heavy_atom_count(smiles)

        record = {
            "row_idx": row_idx,
            "status": "",
            "smiles": smiles,
            "out_dir": str(out_dir) if out_dir is not None else "",
            "num_heavy_atoms": "" if num_heavy_atoms is None else num_heavy_atoms,
            "seconds": "",
            "reason": "",
        }

        if num_heavy_atoms is None:
            failed_count += 1
            record.update(status="failed", reason="invalid_smiles", seconds=f"{time.time() - start:.3f}")
            append_summary(summary_path, record)
            continue

        if max_atoms is not None and num_heavy_atoms > int(max_atoms):
            skipped_count += 1
            record.update(status="skipped", reason=f"heavy_atoms>{max_atoms}", seconds=f"{time.time() - start:.3f}")
            append_summary(summary_path, record)
            continue

        if out_dir is not None and has_core_outputs(out_dir) and not force:
            skipped_count += 1
            record.update(status="skipped", reason="existing_core_outputs", seconds=f"{time.time() - start:.3f}")
            append_summary(summary_path, record)
            continue

        try:
            if out_dir is None:
                with tempfile.TemporaryDirectory(prefix=f"zinc_md_{int(row_idx):06d}_") as tmp_dir:
                    tmp_out_dir = Path(tmp_dir)
                    with time_limit(molecule_timeout_sec), maybe_silence(verbose):
                        run_openmm_md(
                            smiles=smiles,
                            out_dir=tmp_out_dir,
                            equilibration_steps=equilibration_steps,
                            steps=steps,
                            report_interval=report_interval,
                            seed=int(seed) + int(row_idx),
                            temperature_kelvin=temperature_kelvin,
                            timestep_fs=timestep_fs,
                            friction_per_ps=friction_per_ps,
                            platform_name=platform,
                            cgn_views=cgn_views,
                            cgn_sigma=cgn_sigma,
                            trajectory_cgn=True,
                            can_views=can_views,
                            can_max_angle_deg=can_max_angle_deg,
                            trajectory_can=True,
                            trajectory_cgn_can=True,
                            partial_charge_method=partial_charge_method,
                            save_intermediates=save_intermediates,
                        )
                    if not has_core_outputs(tmp_out_dir):
                        raise RuntimeError("missing_core_outputs_after_generation")
                    base_item, base_cgn_can_item, md_item = load_packed_views(tmp_out_dir, record)
                    base_items.append(base_item)
                    base_cgn_can_items.append(base_cgn_can_item)
                    md_items.append(md_item)
            else:
                with time_limit(molecule_timeout_sec), maybe_silence(verbose):
                    run_openmm_md(
                        smiles=smiles,
                        out_dir=out_dir,
                        equilibration_steps=equilibration_steps,
                        steps=steps,
                        report_interval=report_interval,
                        seed=int(seed) + int(row_idx),
                        temperature_kelvin=temperature_kelvin,
                        timestep_fs=timestep_fs,
                        friction_per_ps=friction_per_ps,
                        platform_name=platform,
                        cgn_views=cgn_views,
                        cgn_sigma=cgn_sigma,
                        trajectory_cgn=True,
                        can_views=can_views,
                        can_max_angle_deg=can_max_angle_deg,
                        trajectory_can=True,
                        trajectory_cgn_can=True,
                        partial_charge_method=partial_charge_method,
                        save_intermediates=save_intermediates,
                    )
                if not has_core_outputs(out_dir):
                    raise RuntimeError("missing_core_outputs_after_generation")
            ok_count += 1
            record.update(status="ok", seconds=f"{time.time() - start:.3f}")
        except KeyboardInterrupt:
            raise
        except Exception as exc:
            failed_count += 1
            reason = repr(exc)
            if verbose:
                reason = reason + " | " + traceback.format_exc().splitlines()[-1]
            record.update(status="failed", reason=reason, seconds=f"{time.time() - start:.3f}")

        append_summary(summary_path, record)

    finished_at = time.time()
    if keep_folders:
        packed_count = pack_successful_outputs(out_root, summary_path, pack_paths)
    else:
        packed_count = save_three_pt(pack_paths, base_items, base_cgn_can_items, md_items)
    write_run_report(
        report_path,
        out_root=out_root,
        summary_path=summary_path,
        pack_paths=pack_paths,
        packed_count=packed_count,
        ok_count=ok_count,
        skipped_count=skipped_count,
        failed_count=failed_count,
        total_count=len(rows),
        started_at=started_at,
        finished_at=finished_at,
    )
    print(f"Done: ok={ok_count}, skipped={skipped_count}, failed={failed_count}, total={len(rows)}")
    print(f"Report: {report_path}")
    print(f"Summary: {summary_path}")
    print(f"Packed: {packed_count} molecules")
    print(f"  base: {pack_paths['base']}")
    print(f"  base_cgn_can: {pack_paths['base_cgn_can']}")
    print(f"  md_cgn_can_electronic: {pack_paths['md_cgn_can_electronic']}")


def build_arg_parser() -> argparse.ArgumentParser:
    parser = argparse.ArgumentParser(description="Batch-generate real OpenMM MD + CAN + CGN files for ZINC SMILES.")
    parser.add_argument("--csv", default="zinc/zinc15_250K.csv")
    parser.add_argument("--out-root", default="zinc/zinc15_250K_openmm_md_1000")
    parser.add_argument("--smiles-col", default="smiles")
    parser.add_argument("--start-index", type=int, default=0)
    parser.add_argument("--max-molecules", type=int, default=1000)
    parser.add_argument("--max-atoms", type=int, default=80)
    parser.add_argument("--molecule-timeout-sec", type=float, default=180.0)
    parser.add_argument("--equilibration-steps", type=int, default=1000)
    parser.add_argument("--steps", type=int, default=1000)
    parser.add_argument("--report-interval", type=int, default=100)
    parser.add_argument("--platform", default="OpenCL")
    parser.add_argument("--seed", type=int, default=2026)
    parser.add_argument("--temperature-k", type=float, default=300.0)
    parser.add_argument("--timestep-fs", type=float, default=1.0)
    parser.add_argument("--friction-per-ps", type=float, default=1.0)
    parser.add_argument(
        "--partial-charge-method",
        default="gasteiger",
        choices=["gasteiger", "mmff94", "formal_charge", "zeros", "am1bcc", "default"],
    )
    parser.add_argument("--cgn-views", type=int, default=4)
    parser.add_argument("--cgn-sigma", type=float, default=0.02)
    parser.add_argument("--can-views", type=int, default=4)
    parser.add_argument("--can-max-angle-deg", type=float, default=30.0)
    parser.add_argument("--keep-folders", action="store_true", help="Keep one output folder per molecule. Default uses temporary folders and only saves packed .pt.")
    parser.add_argument("--force", action="store_true")
    parser.add_argument("--save-intermediates", action="store_true")
    parser.add_argument("--verbose", action="store_true")
    return parser


def main() -> None:
    args = build_arg_parser().parse_args()
    preprocess_zinc_openmm_md(
        csv_path=args.csv,
        out_root=args.out_root,
        smiles_col=args.smiles_col,
        start_index=args.start_index,
        max_molecules=args.max_molecules,
        max_atoms=args.max_atoms,
        molecule_timeout_sec=args.molecule_timeout_sec,
        equilibration_steps=args.equilibration_steps,
        steps=args.steps,
        report_interval=args.report_interval,
        platform=args.platform,
        seed=args.seed,
        temperature_kelvin=args.temperature_k,
        timestep_fs=args.timestep_fs,
        friction_per_ps=args.friction_per_ps,
        partial_charge_method=args.partial_charge_method,
        cgn_views=args.cgn_views,
        cgn_sigma=args.cgn_sigma,
        can_views=args.can_views,
        can_max_angle_deg=args.can_max_angle_deg,
        keep_folders=args.keep_folders,
        force=args.force,
        save_intermediates=args.save_intermediates,
        verbose=args.verbose,
    )


if __name__ == "__main__":
    main()
```
