## Supplementary material for "PG-MLD: Physics-Guided Molecular Representation Learning via Dynamic 3D Trajectory Distillation": Code and Data: run_all_chemblm_pca.py

```
from __future__ import annotations

import argparse
import csv
import os
import subprocess
import sys
from pathlib import Path


PROJECT_ROOT = Path(__file__).resolve().parents[1]
VISUALIZER = PROJECT_ROOT / "visualization" / "t-sne.py"

TASKS = {
    "bbbp": {
        "dataset": "BBBP",
        "config": "configs/finetune/chemblm/dynamic_random/bbbp.yaml",
        "target": "p_np",
        "type": "classification",
    },
    "bace": {
        "dataset": "BACE",
        "config": "configs/finetune/chemblm/dynamic_random/bace.yaml",
        "target": "Class",
        "type": "classification",
    },
    "clintox": {
        "dataset": "ClinTox",
        "config": "configs/finetune/chemblm/dynamic_random/clintox.yaml",
        "target": "CT_TOX",
        "type": "classification",
    },
    "tox21": {
        "dataset": "Tox21",
        "config": "configs/finetune/chemblm/dynamic_random/tox21.yaml",
        "target": "SR-p53",
        "type": "classification",
    },
    "toxcast": {
        "dataset": "ToxCast",
        "config": "configs/finetune/chemblm/dynamic_random/toxcast.yaml",
        "target": "TOX21_TR_LUC_GH3_Antagonist",
        "type": "classification",
    },
    "hiv": {
        "dataset": "HIV",
        "config": "configs/finetune/chemblm/dynamic_random/hiv.yaml",
        "target": "HIV_active",
        "type": "classification",
    },
}


def read_best_checkpoint(config_path: Path) -> Path:
    """Read paths.best_checkpoint without adding a PyYAML dependency."""
    in_paths = False
    for raw_line in config_path.read_text(encoding="utf-8").splitlines():
        stripped = raw_line.strip()
        if raw_line and not raw_line[0].isspace() and stripped.endswith(":"):
            in_paths = stripped == "paths:"
            continue
        if in_paths and stripped.startswith("best_checkpoint:"):
            value = stripped.split(":", 1)[1].strip().strip("'\"")
            if not value or value.lower() in {"null", "none", "~"}:
                raise ValueError(f"paths.best_checkpoint is empty in {config_path}")
            checkpoint = Path(value)
            return checkpoint if checkpoint.is_absolute() else PROJECT_ROOT / checkpoint
    raise ValueError(f"paths.best_checkpoint was not found in {config_path}")


def parse_args() -> argparse.Namespace:
    parser = argparse.ArgumentParser(description="Run ChemBLM PCA representation plots for molecular benchmarks.")
    parser.add_argument("--tasks", nargs="+", choices=list(TASKS), default=list(TASKS))
    parser.add_argument("--output-root", default="output/figures/pca_chemblm_all")
    parser.add_argument("--subset", default="all", choices=["train", "val", "test", "all"])
    parser.add_argument("--feature", default="penultimate", choices=["encoder", "penultimate", "logits"])
    parser.add_argument("--seed", type=int, default=2026)
    parser.add_argument("--batch-size", type=int, default=64)
    parser.add_argument("--max-points", type=int, default=0)
    parser.add_argument("--continue-on-error", action="store_true")
    return parser.parse_args()


def main() -> None:
    args = parse_args()
    output_root = PROJECT_ROOT / args.output_root
    log_dir = output_root / "logs"
    log_dir.mkdir(parents=True, exist_ok=True)
    rows: list[dict[str, object]] = []

    environment = os.environ.copy()
    environment.update(
        {
            "TRANSFORMERS_OFFLINE": "1",
            "HF_HUB_OFFLINE": "1",
            "TOKENIZERS_PARALLELISM": "false",
            "PYTHONPATH": str(PROJECT_ROOT),
        }
    )

    for task_key in args.tasks:
        spec = TASKS[task_key]
        task_output = output_root / task_key
        config_path = PROJECT_ROOT / spec["config"]
        log_path = log_dir / f"{task_key}.log"

        if not config_path.is_file():
            message = f"Config not found: {config_path}"
            print(f"\n===== {task_key}: skipped =====\n{message}", flush=True)
            rows.append(
                {
                    "task": task_key,
                    "target": spec["target"],
                    "task_type": spec["type"],
                    "status": "missing_config",
                    "return_code": 2,
                    "output_dir": str(task_output.relative_to(PROJECT_ROOT)),
                    "log": str(log_path.relative_to(PROJECT_ROOT)),
                    "message": message,
                }
            )
            if not args.continue_on_error:
                break
            continue

        try:
            checkpoint_path = read_best_checkpoint(config_path)
        except ValueError as error:
            checkpoint_path = None
            message = str(error)
        else:
            message = ""

        if checkpoint_path is None or not checkpoint_path.is_file():
            if checkpoint_path is not None:
                message = f"Checkpoint not found: {checkpoint_path}"
            print(f"\n===== {task_key}: skipped =====\n{message}", flush=True)
            rows.append(
                {
                    "task": task_key,
                    "target": spec["target"],
                    "task_type": spec["type"],
                    "status": "missing_checkpoint",
                    "return_code": 2,
                    "output_dir": str(task_output.relative_to(PROJECT_ROOT)),
                    "log": str(log_path.relative_to(PROJECT_ROOT)),
                    "message": message,
                }
            )
            if not args.continue_on_error:
                break
            continue

        command = [
            sys.executable,
            str(VISUALIZER),
            "--config",
            str(config_path),
            "--checkpoint",
            str(checkpoint_path),
            "--output-dir",
            str(task_output),
            "--dataset-name",
            str(spec["dataset"]),
            "--task-name",
            str(spec["target"]),
            "--subset",
            args.subset,
            "--feature",
            args.feature,
            "--backend",
            "pca",
            "--seed",
            str(args.seed),
            "--batch-size",
            str(args.batch_size),
        ]
        if spec["type"] == "classification":
            command.append("--balance")
        if args.max_points > 0:
            command.extend(["--max-points", str(args.max_points)])

        print(f"\n===== {task_key}: {spec['target']} =====", flush=True)
        print(" ".join(command), flush=True)
        with log_path.open("w", encoding="utf-8") as log_handle:
            process = subprocess.Popen(
                command,
                cwd=PROJECT_ROOT,
                env=environment,
                stdout=subprocess.PIPE,
                stderr=subprocess.STDOUT,
                text=True,
                encoding="utf-8",
                errors ="replace",
                bufsize=1,
            )
            assert process.stdout is not None
            for line in process.stdout:
                print(line, end="")
                log_handle.write(line)
            return_code = process.wait()

        rows.append(
            {
                "task": task_key,
                "target": spec["target"],
                "task_type": spec["type"],
                "status": "ok" if return_code == 0 else "failed",
                "return_code": return_code,
                "output_dir": str(task_output.relative_to(PROJECT_ROOT)),
                "log": str(log_path.relative_to(PROJECT_ROOT)),
                "message": "",
            }
        )
        if return_code != 0 and not args.continue_on_error:
            break

    summary_path = output_root / "summary.csv"
    with summary_path.open("w", newline="", encoding="utf-8") as handle:
        writer = csv.DictWriter(handle, fieldnames=list(rows[0]) if rows else ["task", "status"])
        writer.writeheader()
        writer.writerows(rows)
    print(f"\nSummary: {summary_path}")


if __name__ == "__main__":
    main()
```
