## Supplementary material for "PG-MLD: Physics-Guided Molecular Representation Learning via Dynamic 3D Trajectory Distillation": Code and Data: run_all_finetune_1d.py

```
from __future__ import annotations

import argparse
import csv
import json
import re
import subprocess
import sys
from datetime import datetime
from pathlib import Path


TASK_CONFIGS = [
    ("bbbp", "configs/finetune/scaffold/finetune_bbbp_1d.yaml"),
    ("bace", "configs/finetune/scaffold/finetune_bace_1d.yaml"),
    ("clintox", "configs/finetune/scaffold/finetune_clintox_1d.yaml"),
    ("hiv", "configs/finetune/scaffold/finetune_hiv_1d.yaml"),
    ("tox21", "configs/finetune/scaffold/finetune_tox21_1d.yaml"),
    ("toxcast", "configs/finetune/scaffold/finetune_toxcast_1d.yaml"),
    ("esol", "configs/finetune/scaffold/finetune_esol_1d.yaml"),
    ("freesolv", "configs/finetune/scaffold/finetune_freesolv_1d.yaml"),
    ("lipophilicity", "configs/finetune/scaffold/finetune_lipophilicity_1d.yaml"),
]

EPOCH_RE = re.compile(
    r"^Epoch\s+(?P<epoch>\d+):\s+train_loss=(?P<train_loss>[-+0-9.eE]+)\s+"
    r"val=(?P<val>.*?)\s+test=(?P<test>.*?)\s+best_epoch=(?P<best_epoch>\d+)\s*$"
)
METRIC_RE = re.compile(r"([A-Za-z_][A-Za-z0-9_]*)=([-+0-9.eE]+|nan|NaN|inf|-inf)")


def parse_metrics(text: str) -> dict[str, float]:
    metrics: dict[str, float] = {}
    for key, value in METRIC_RE.findall(text):
        try:
            metrics[key] = float(value)
        except ValueError:
            pass
    return metrics


def parse_best_from_log(log_path: Path) -> dict[str, object]:
    best_row: dict[str, object] | None = None
    best_epoch_seen = 0
    last_epoch_row: dict[str, object] | None = None

    with log_path.open("r", encoding="utf-8", errors="replace") as handle:
        for line in handle:
            match = EPOCH_RE.match(line.strip())
            if not match:
                continue
            epoch = int(match.group("epoch"))
            best_epoch = int(match.group("best_epoch"))
            row: dict[str, object] = {
                "epoch": epoch,
                "train_loss": float(match.group("train_loss")),
                "best_epoch_reported": best_epoch,
            }
            for prefix, blob in [("val", match.group("val")), ("test", match.group("test"))]:
                for key, value in parse_metrics(blob).items():
                    row[f"{prefix}_{key}"] = value
            last_epoch_row = row

            if best_epoch > best_epoch_seen:
                best_epoch_seen = best_epoch
                best_row = row

    if best_row is None:
        return {
            "status": "no_epoch_metrics_found",
            "log": str(log_path),
        }
    best_row["status"] = "ok"
    best_row["log"] = str(log_path)
    if last_epoch_row is not None:
        best_row["last_epoch"] = last_epoch_row.get("epoch")
    return best_row


def run_task(task: str, config: str, log_path: Path, split_method: str) -> int:
    command = [
        sys.executable,
        "-m",
        "comld.train.finetune_mol_1d",
        "--config",
        config,
        "--split-method",
        split_method,
    ]
    print(f"\n===== Running {task}: {' '.join(command)} =====", flush=True)
    log_path.parent.mkdir(parents=True, exist_ok=True)
    with log_path.open("w", encoding="utf-8") as log:
        log.write(f"# task: {task}\n")
        log.write(f"# config: {config}\n")
        log.write(f"# started_at: {datetime.now().isoformat(timespec='seconds')}\n")
        log.write(f"# command: {' '.join(command)}\n\n")
        log.flush()
        process = subprocess.Popen(
            command,
            stdout=subprocess.PIPE,
            stderr=subprocess.STDOUT,
            text=True,
            bufsize=1,
        )
        assert process.stdout is not None
        for line in process.stdout:
            print(line, end="", flush=True)
            log.write(line)
        return_code = process.wait()
        log.write(f"\n# finished_at: {datetime.now().isoformat(timespec='seconds')}\n")
        log.write(f"# return_code: {return_code}\n")
    return return_code


def main() -> None:
    parser = argparse.ArgumentParser(description="Run all 1D finetuning tasks and summarize best metrics.")
    parser.add_argument("--tasks", nargs="+", choices=[name for name, _ in TASK_CONFIGS], help="Tasks to run.")
    parser.add_argument("--split-method", default="scaffold", choices=["random", "scaffold"])
    parser.add_argument("--log-dir", default="logs/finetune_1d_all")
    parser.add_argument("--summary-csv", default="logs/finetune_1d_all/summary.csv")
    parser.add_argument("--summary-json", default="logs/finetune_1d_all/summary.json")
    parser.add_argument("--continue-on-error", action="store_true")
    args = parser.parse_args()

    selected = {task for task in args.tasks} if args.tasks else None
    configs = [(task, config) for task, config in TASK_CONFIGS if selected is None or task in selected]

    log_dir = Path(args.log_dir)
    results: list[dict[str, object]] = []
    for task, config in configs:
        log_path = log_dir / f"{task}.log"
        if not Path(config).exists():
            row = {"task": task, "config": config, "status": "missing_config", "log": str(log_path)}
            results.append(row)
            print(f"Missing config for {task}: {config}", flush=True)
            if not args.continue_on_error:
                break
            continue

        return_code = run_task(task, config, log_path, args.split_method)
        row = {"task": task, "config": config, "return_code": return_code}
        row.update(parse_best_from_log(log_path))
        results.append(row)
        if return_code != 0 and not args.continue_on_error:
            print(f"Stopping because {task} failed with return_code={return_code}", flush=True)
            break

    summary_csv = Path(args.summary_csv)
    summary_json = Path(args.summary_json)
    summary_csv.parent.mkdir(parents=True, exist_ok=True)
    all_keys = sorted({key for row in results for key in row})
    preferred = [
        "task",
        "status",
        "config",
        "return_code",
        "epoch",
        "best_epoch_reported",
        "last_epoch",
        "train_loss",
        "val_roc_auc",
        "test_roc_auc",
        "val_accuracy",
        "test_accuracy",
        "val_rmse",
        "test_rmse",
        "val_mae",
        "test_mae",
        "val_loss",
        "test_loss",
        "log",
    ]
    fieldnames = [key for key in preferred if key in all_keys] + [key for key in all_keys if key not in preferred]
    with summary_csv.open("w", newline="", encoding="utf-8") as handle:
        writer = csv.DictWriter(handle, fieldnames=fieldnames)
        writer.writeheader()
        writer.writerows(results)
    with summary_json.open("w", encoding="utf-8") as handle:
        json.dump(results, handle, ensure_ascii=False, indent=2)

    print(f"\nSummary CSV: {summary_csv}", flush=True)
    print(f"Summary JSON: {summary_json}", flush=True)


if __name__ == "__main__":
    main()
```
