## Supplementary material for "PG-MLD: Physics-Guided Molecular Representation Learning via Dynamic 3D Trajectory Distillation": Code and Data: run_chemberta_random_suite.sh

```
#!/usr/bin/env bash
set -u

cd "$(dirname "$0")/.."

export TRANSFORMERS_OFFLINE=1
export HF_HUB_OFFLINE=1
export TOKENIZERS_PARALLELISM=false

mkdir -p logs/chemberta_random_suite

configs=(
  configs/finetune/random/chemberta/finetune_bace_chemberta_random.yaml
  configs/finetune/random/chemberta/finetune_bace_chemberta_distilled_random.yaml
  configs/finetune/random/chemberta/finetune_clintox_chemberta_random.yaml
  configs/finetune/random/chemberta/finetune_clintox_chemberta_distilled_random.yaml
  configs/finetune/random/chemberta/finetune_hiv_chemberta_random.yaml
  configs/finetune/random/chemberta/finetune_hiv_chemberta_distilled_random.yaml
  configs/finetune/random/chemberta/finetune_tox21_chemberta_random.yaml
  configs/finetune/random/chemberta/finetune_tox21_chemberta_distilled_random.yaml
  configs/finetune/random/chemberta/finetune_toxcast_chemberta_random.yaml
  configs/finetune/random/chemberta/finetune_toxcast_chemberta_distilled_random.yaml
)

for cfg in "${configs[@]}"; do
  name="$(basename "${cfg%.yaml}")"
  echo "===== Running ${name} ====="
  python -m comld.train.finetune_bbbp_1d \
    --config "${cfg}" \
    --split-method random \
    2>&1 | tee "logs/chemberta_random_suite/${name}.log"
done
```
