## Supplementary material for "PG-MLD: Physics-Guided Molecular Representation Learning via Dynamic 3D Trajectory Distillation": Code and Data: run_chemblm_tasks.py

```
from __future__ import annotations

import argparse
import csv
import json
import os
import re
import subprocess
import sys
from datetime import datetime
from pathlib import Path


PROJECT_ROOT = Path(__file__).resolve().parents[1]
TASKS = ("bbbp", "bace", "clintox", "tox21", "toxcast", "hiv")
VARIANTS = ("random", "dynamic_random")
EPOCH_RE = re.compile(
    r"^Epoch\s+(?P<epoch>\d+):\s+train_loss=(?P<train_loss>[-+0-9.eE]+)\s+"
    r"val=(?P<val>.*?)\s+test=(?P<test>.*?)\s+best_epoch=(?P<best_epoch>\d+)\s*$"
)
METRIC_RE = re.compile(r"([A-Za-z_][A-Za-z0-9_]*)=([-+0-9.eE]+|nan|NaN|inf|-inf)")


def parse_args() -> argparse.Namespace:
    parser = argparse.ArgumentParser(
        description="Run clean ChemBLM random and dynamic_random classification experiments."
    )
    parser.add_argument("--tasks", nargs="+", choices=TASKS, default=list(TASKS))
    parser.add_argument("--variants", nargs="+", choices=VARIANTS, default=list(VARIANTS))
    parser.add_argument("--continue-on-error", action="store_true")
    parser.add_argument(
        "--overwrite",
        action="store_true",
        help="Rerun even when the target best.pt already exists.",
    )
    return parser.parse_args()


def config_path(task: str, variant: str) -> Path:
    return PROJECT_ROOT / "configs" / "finetune" / "chemblm" / variant / f"{task}.yaml"


def checkpoint_path(task: str, variant: str) -> Path:
    return PROJECT_ROOT / "checkpoints" / task / "chemblm" / variant / "best.pt"


def log_path(task: str, variant: str) -> Path:
    return PROJECT_ROOT / "logs" / task / "chemblm" / f"{variant}.log"


def parse_metrics(text: str) -> dict[str, float]:
    result: dict[str, float] = {}
    for key, value in METRIC_RE.findall(text):
        try:
            result[key] = float(value)
        except ValueError:
            continue
    return result


def parse_best_from_log(path: Path) -> dict[str, object]:
    best_epoch_seen = 0
    best_row: dict[str, object] | None = None
    last_epoch = 0
    with path.open("r", encoding="utf-8", errors="replace") as handle:
        for line in handle:
            match = EPOCH_RE.match(line.strip())
            if match is None:
                continue
            epoch = int(match.group("epoch"))
            reported_best = int(match.group("best_epoch"))
            last_epoch = epoch
            if reported_best <= best_epoch_seen:
                continue
            best_epoch_seen = reported_best
            row: dict[str, object] = {
                "best_epoch": reported_best,
                "train_loss_at_best": float(match.group("train_loss")),
            }
            for prefix, blob in (("val", match.group("val")), ("test", match.group("test"))):
                for name, value in parse_metrics(blob).items():
                    row[f"{prefix}_{name}"] = value
            best_row = row
    if best_row is None:
        return {"parse_status": "no_epoch_metrics", "last_epoch": last_epoch}
    best_row["parse_status"] = "ok"
    best_row["last_epoch"] = last_epoch
    return best_row


def main() -> None:
    args = parse_args()
    environment = os.environ.copy()
    environment.update(
        {
            "TRANSFORMERS_OFFLINE": "1",
            "HF_HUB_OFFLINE": "1",
            "TOKENIZERS_PARALLELISM": "false",
            "PYTHONPATH": str(PROJECT_ROOT),
        }
    )
    distilled_checkpoint = (
        PROJECT_ROOT / "checkpoints" / "pretrain_zinc_dynamic_3d_to_1d_chemblm" / "latest.pt"
    )
    results: list[dict[str, object]] = []
    stop = False

    for task in args.tasks:
        for variant in args.variants:
            cfg = config_path(task, variant)
            best = checkpoint_path(task, variant)
            log = log_path(task, variant)
            base_row: dict[str, object] = {
                "task": task,
                "model": "chemblm",
                "variant": variant,
                "config": str(cfg.relative_to(PROJECT_ROOT)),
                "checkpoint": str(best.relative_to(PROJECT_ROOT)),
                "log": str(log.relative_to(PROJECT_ROOT)),
            }

            if not cfg.is_file():
                print(f"Missing config: {cfg}", flush=True)
                results.append({**base_row, "status": "missing_config", "return_code": 2})
                stop = not args.continue_on_error
                if stop:
                    break
                continue
            if variant == "dynamic_random" and not distilled_checkpoint.is_file():
                print(f"Missing distilled checkpoint: {distilled_checkpoint}", flush=True)
                results.append({**base_row, "status": "missing_pretrained_checkpoint", "return_code": 2})
                stop = not args.continue_on_error
                if stop:
                    break
                continue
            if best.is_file() and not args.overwrite:
                print(f"Skip existing checkpoint: {best}", flush=True)
                results.append({**base_row, "status": "skipped_existing", "return_code": 0})
                continue

            command = [
                sys.executable,
                "-m",
                "comld.train.finetune_chemblm_1d",
                "--config",
                str(cfg),
                "--split-method",
                "random",
            ]
            print(f"\n===== {task} / chemblm / {variant} =====", flush=True)
            print(" ".join(command), flush=True)
            log.parent.mkdir(parents=True, exist_ok=True)
            with log.open("w", encoding="utf-8") as handle:
                handle.write(f"# started_at: {datetime.now().isoformat(timespec='seconds')}\n")
                handle.write(f"# command: {' '.join(command)}\n\n")
                handle.flush()
                process = subprocess.Popen(
                    command,
                    cwd=PROJECT_ROOT,
                    env=environment,
                    stdout=subprocess.PIPE,
                    stderr=subprocess.STDOUT,
                    text=True,
                    encoding="utf-8",
                    errors="replace",
                    bufsize=1,
                )
                assert process.stdout is not None
                for line in process.stdout:
                    print(line, end="", flush=True)
                    handle.write(line)
                return_code = process.wait()
                handle.write(f"\n# return_code: {return_code}\n")
                handle.write(f"# finished_at: {datetime.now().isoformat(timespec='seconds')}\n")

            row = {
                **base_row,
                "status": "ok" if return_code == 0 else "failed",
                "return_code": return_code,
            }
            row.update(parse_best_from_log(log))
            results.append(row)
            if return_code != 0 and not args.continue_on_error:
                stop = True
                break
        if stop:
            break

    summary_dir = PROJECT_ROOT / "logs" / "chemblm"
    summary_dir.mkdir(parents=True, exist_ok=True)
    summary_csv = summary_dir / "summary.csv"
    summary_json = summary_dir / "summary.json"

    existing_rows: list[dict[str, object]] = []
    if summary_json.is_file():
        try:
            loaded = json.loads(summary_json.read_text(encoding="utf-8"))
            if isinstance(loaded, list):
                existing_rows = [row for row in loaded if isinstance(row, dict)]
        except (OSError, json.JSONDecodeError):
            existing_rows = []

    merged: dict[tuple[str, str, str], dict[str, object]] = {}
    for row in existing_rows:
        key = (str(row.get("task", "")), str(row.get("model", "")), str(row.get("variant", "")))
        merged[key] = row
    for row in results:
        key = (str(row.get("task", "")), str(row.get("model", "")), str(row.get("variant", "")))
        if row.get("status") == "skipped_existing" and key in merged:
            continue
        merged[key] = row
    task_order = {name: index for index, name in enumerate(TASKS)}
    variant_order = {name: index for index, name in enumerate(VARIANTS)}
    results = sorted(
        merged.values(),
        key=lambda row: (
            task_order.get(str(row.get("task", "")), len(task_order)),
            variant_order.get(str(row.get("variant", "")), len(variant_order)),
        ),
    )

    all_keys = sorted({key for row in results for key in row})
    preferred = [
        "task",
        "model",
        "variant",
        "status",
        "return_code",
        "best_epoch",
        "last_epoch",
        "val_roc_auc",
        "test_roc_auc",
        "val_micro_roc_auc",
        "test_micro_roc_auc",
        "val_accuracy",
        "test_accuracy",
        "val_loss",
        "test_loss",
        "config",
        "checkpoint",
        "log",
    ]
    fieldnames = [key for key in preferred if key in all_keys]
    fieldnames.extend(key for key in all_keys if key not in fieldnames)
    with summary_csv.open("w", newline="", encoding="utf-8") as handle:
        writer = csv.DictWriter(handle, fieldnames=fieldnames)
        writer.writeheader()
        writer.writerows(results)
    with summary_json.open("w", encoding="utf-8") as handle:
        json.dump(results, handle, ensure_ascii=False, indent=2)
    print(f"\nSummary CSV: {summary_csv}", flush=True)
    print(f"Summary JSON: {summary_json}", flush=True)


if __name__ == "__main__":
    main()
```
