## Supplementary material for "PG-MLD: Physics-Guided Molecular Representation Learning via Dynamic 3D Trajectory Distillation": Code and Data: run_qm9.py

```
from __future__ import annotations

import argparse
import subprocess
import sys
from pathlib import Path


TARGETS = ("mu", "alpha", "homo", "lumo", "gap", "zpve", "u0", "u", "h")


def parse_targets(value: str) -> list[str]:
    if value.strip().lower() == "all":
        return list(TARGETS)
    targets = [item.strip().lower() for item in value.split(",") if item.strip()]
    invalid = [target for target in targets if target not in TARGETS]
    if invalid:
        raise ValueError(f"Unsupported targets {invalid}; choose from {TARGETS} or use all.")
    return targets


def parse_seeds(value: str) -> list[int]:
    seeds = [int(item.strip()) for item in value.split(",") if item.strip()]
    if not seeds:
        raise ValueError("At least one seed is required.")
    return seeds


def main() -> None:
    parser = argparse.ArgumentParser(
        description="Run one or more single-property QM9 experiments from one model config."
    )
    parser.add_argument("--config", required=True)
    parser.add_argument("--targets", default="all")
    parser.add_argument("--seeds", default="42,43,44")
    parser.add_argument("--overwrite", action="store_true")
    args = parser.parse_args()

    config_path = Path(args.config)
    if not config_path.exists():
        raise FileNotFoundError(config_path)

    import yaml

    config = yaml.safe_load(config_path.read_text(encoding="utf-8")) or {}
    checkpoint_root = Path(config.get("paths", {}).get("checkpoint_root", "checkpoints/qm9"))
    targets = parse_targets(args.targets)
    seeds = parse_seeds(args.seeds)

    failures = 0
    for target in targets:
        for seed in seeds:
            checkpoint = checkpoint_root / target / f"seed_{seed}" / "best.pt"
            if checkpoint.exists() and not args.overwrite:
                print(f"Skip existing checkpoint: {checkpoint}")
                continue
            command = [
                sys.executable,
                "-m",
                "comld.train.finetune_qm9_1d",
                "--config",
                str(config_path),
                "--target",
                target,
                "--seed",
                str(seed),
            ]
            print("\n===== " + " ".join(command) + " =====", flush=True)
            completed = subprocess.run(command, check=False)
            if completed.returncode != 0:
                failures += 1
                print(f"FAILED: target={target} seed={seed} return_code={completed.returncode}")
                break

    if failures:
        raise SystemExit(f"{failures} QM9 target(s) failed.")
    print("QM9 runs completed. Use scripts/summarize_qm9.py to build the MAE table.")


if __name__ == "__main__":
    main()
```
