## Supplementary material for "PG-MLD: Physics-Guided Molecular Representation Learning via Dynamic 3D Trajectory Distillation": Code and Data: sample_pubchem_smiles.py

```
from __future__ import annotations

import argparse
import csv
import gzip
import random
from pathlib import Path
from typing import Any

from tqdm import tqdm


def parse_pubchem_line(line: str) -> tuple[str, str] | None:
    line = line.strip()
    if not line:
        return None
    parts = line.split()
    if len(parts) < 2:
        return None
    cid = parts[0]
    smiles = parts[1]
    return cid, smiles


def descriptors(smiles: str) -> dict[str, Any] | None:
    from rdkit import Chem
    from rdkit.Chem import Crippen, Descriptors, Lipinski, rdMolDescriptors

    mol = Chem.MolFromSmiles(smiles)
    if mol is None:
        return None
    heavy_atoms = int(mol.GetNumHeavyAtoms())
    if heavy_atoms <= 1:
        return None

    return {
        "smiles": Chem.MolToSmiles(mol),
        "mol_weight": float(Descriptors.MolWt(mol)),
        "logp": float(Crippen.MolLogP(mol)),
        "tpsa": float(rdMolDescriptors.CalcTPSA(mol)),
        "heavy_atom_count": heavy_atoms,
        "ring_count": int(rdMolDescriptors.CalcNumRings(mol)),
        "aromatic_ring_count": int(rdMolDescriptors.CalcNumAromaticRings(mol)),
        "rotatable_bonds": int(Lipinski.NumRotatableBonds(mol)),
        "hbd": int(Lipinski.NumHDonors(mol)),
        "hba": int(Lipinski.NumHAcceptors(mol)),
    }


def bin_index(value: float, low: float, high: float, bins: int) -> int | None:
    if value < low or value > high:
        return None
    if value == high:
        return bins - 1
    width = (high - low) / bins
    return int((value - low) / width)


def balanced_bins(items_by_bin: list[list[dict[str, Any]]], target: int, seed: int) -> list[dict[str, Any]]:
    rng = random.Random(seed)
    non_empty = [bucket for bucket in items_by_bin if bucket]
    if not non_empty:
        return []
    per_bin = max(1, target // len(non_empty))

    chosen: list[dict[str, Any]] = []
    leftovers: list[dict[str, Any]] = []
    for bucket in non_empty:
        rng.shuffle(bucket)
        chosen.extend(bucket[:per_bin])
        leftovers.extend(bucket[per_bin:])

    if len(chosen) < target and leftovers:
        rng.shuffle(leftovers)
        chosen.extend(leftovers[: target - len(chosen)])
    if len(chosen) > target:
        rng.shuffle(chosen)
        chosen = chosen[:target]
    rng.shuffle(chosen)
    return chosen


def main() -> None:
    parser = argparse.ArgumentParser(
        description=(
            "Sample PubChem CID-SMILES.gz into a visualization-friendly CSV. "
            "The default strategy stratifies molecules by molecular weight so "
            "PCA color maps have a wider descriptor range."
        )
    )
    parser.add_argument("--input", required=True, help="PubChem CID-SMILES.gz")
    parser.add_argument("--out", required=True)
    parser.add_argument("--target", type=int, default=50000)
    parser.add_argument("--seed", type=int, default=2026)
    parser.add_argument("--max-lines", type=int, default=3000000, help="0 means scan the whole file.")
    parser.add_argument("--mw-min", type=float, default=80.0)
    parser.add_argument("--mw-max", type=float, default=700.0)
    parser.add_argument("--bins", type=int, default=50)
    parser.add_argument(
        "--per-bin-cap",
        type=int,
        default=3000,
        help="Reservoir cap per molecular-weight bin while streaming.",
    )
    args = parser.parse_args()

    rng = random.Random(args.seed)
    input_path = Path(args.input)
    out_path = Path(args.out)
    out_path.parent.mkdir(parents=True, exist_ok=True)

    buckets: list[list[dict[str, Any]]] = [[] for _ in range(int(args.bins))]
    seen_by_bin = [0 for _ in range(int(args.bins))]
    scanned = 0
    valid = 0
    in_range = 0

    with gzip.open(input_path, "rt", encoding="utf-8", errors="ignore") as handle:
        progress_total = args.max_lines if args.max_lines and args.max_lines > 0 else None
        for line in tqdm(handle, total=progress_total, desc="Scan PubChem"):
            if args.max_lines and scanned >= int(args.max_lines):
                break
            scanned += 1
            parsed = parse_pubchem_line(line)
            if parsed is None:
                continue
            cid, smiles = parsed
            desc = descriptors(smiles)
            if desc is None:
                continue
            valid += 1
            b = bin_index(desc["mol_weight"], float(args.mw_min), float(args.mw_max), int(args.bins))
            if b is None:
                continue
            in_range += 1
            row = {"cid": cid, **desc}

            seen_by_bin[b] += 1
            bucket = buckets[b]
            if len(bucket) < int(args.per_bin_cap):
                bucket.append(row)
            else:
                # Reservoir sampling inside each MW bin.
                j = rng.randrange(seen_by_bin[b])
                if j < int(args.per_bin_cap):
                    bucket[j] = row

    chosen = balanced_bins(buckets, target=int(args.target), seed=int(args.seed))
    if not chosen:
        raise RuntimeError("No molecules selected. Try wider --mw-min/--mw-max or larger --max-lines.")

    fieldnames = [
        "cid",
        "smiles",
        "mol_weight",
        "logp",
        "tpsa",
        "heavy_atom_count",
        "ring_count",
        "aromatic_ring_count",
        "rotatable_bonds",
        "hbd",
        "hba",
    ]
    with open(out_path, "w", newline="", encoding="utf-8") as handle:
        writer = csv.DictWriter(handle, fieldnames=fieldnames)
        writer.writeheader()
        writer.writerows(chosen)

    weights = [float(row["mol_weight"]) for row in chosen]
    print(f"Scanned lines: {scanned}")
    print(f"Valid RDKit molecules: {valid}")
    print(f"In MW range [{args.mw_min}, {args.mw_max}]: {in_range}")
    print(f"Selected molecules: {len(chosen)}")
    print(f"MW range selected: min={min(weights):.3f} median={sorted(weights)[len(weights)//2]:.3f} max={max(weights):.3f}")
    print(f"Saved: {out_path}")


if __name__ == "__main__":
    main()
```
