## Supplementary material for "PG-MLD: Physics-Guided Molecular Representation Learning via Dynamic 3D Trajectory Distillation": Code and Data: summarize_qm9.py

```
from __future__ import annotations

import argparse
import csv
import json
from collections import defaultdict
from pathlib import Path
from statistics import mean, stdev
from typing import Any


TARGETS = ("mu", "alpha", "homo", "lumo", "gap", "zpve", "u0", "u", "h")
TARGET_HEADERS = {
    "mu": r"$\mu$ (D)",
    "alpha": r"$\alpha$ ($a_0^3$)",
    "homo": "HOMO (meV)",
    "lumo": "LUMO (meV)",
    "gap": "Gap (meV)",
    "zpve": "ZPVE (meV)",
    "u0": r"$U_0$ (meV)",
    "u": r"$U$ (meV)",
    "h": r"$H$ (meV)",
}
DECIMALS = {
    "mu": 3,
    "alpha": 3,
    "homo": 1,
    "lumo": 1,
    "gap": 1,
    "zpve": 2,
    "u0": 2,
    "u": 2,
    "h": 2,
}


def latex_escape(value: str) -> str:
    return (
        value.replace("\\", r"\textbackslash{}")
        .replace("&", r"\&")
        .replace("%", r"\%")
        .replace("_", r"\_")
    )


def format_value(target: str, average: float, deviation: float) -> str:
    digits = DECIMALS[target]
    return f"{average:.{digits}f}" + r"$\pm$" + f"{deviation:.{digits}f}"


def collect(root: Path) -> list[dict[str, Any]]:
    records: list[dict[str, Any]] = []
    for path in sorted(root.rglob("result.json")):
        try:
            record = json.loads(path.read_text(encoding="utf-8"))
            target = str(record["target"])
            value = float(record["test"]["mae"])
        except (KeyError, TypeError, ValueError, json.JSONDecodeError) as exc:
            print(f"Skip invalid result {path}: {exc}")
            continue
        if target not in TARGETS:
            continue
        record["_path"] = str(path)
        record["_mae"] = value
        records.append(record)
    return records


def aggregate(records: list[dict[str, Any]]) -> dict[tuple[str, str, str], tuple[float, float, int]]:
    grouped: dict[tuple[str, str, str], list[float]] = defaultdict(list)
    for record in records:
        key = (
            str(record.get("experiment", record.get("model_family", "model"))),
            str(record.get("variant", "original")),
            str(record["target"]),
        )
        grouped[key].append(float(record["_mae"]))

    result: dict[tuple[str, str, str], tuple[float, float, int]] = {}
    for key, values in grouped.items():
        result[key] = (
            mean(values),
            stdev(values) if len(values) > 1 else 0.0,
            len(values),
        )
    return result


def display_name(experiment: str, variant: str) -> str:
    if variant.lower() in {"dynamic", "dynamic_random", "distilled", "pg-mld"}:
        return f"{experiment} + PG-MLD"
    return experiment


def highlight_cells(
    aggregate_values: dict[tuple[str, str, str], tuple[float, float, int]],
) -> dict[tuple[str, str, str], str]:
    marks: dict[tuple[str, str, str], str] = {}
    for target in TARGETS:
        candidates = sorted(
            (
                (stats[0], key)
                for key, stats in aggregate_values.items()
                if key[2] == target
            ),
            key=lambda item: item[0],
        )
        if candidates:
            marks[candidates[0][1]] = "best"
        if len(candidates) > 1:
            marks[candidates[1][1]] = "second"
    return marks


def write_csv(
    destination: Path,
    aggregate_values: dict[tuple[str, str, str], tuple[float, float, int]],
) -> None:
    rows = sorted({(key[0], key[1]) for key in aggregate_values})
    destination.parent.mkdir(parents=True, exist_ok=True)
    with destination.open("w", newline="", encoding="utf-8-sig") as handle:
        writer = csv.writer(handle)
        writer.writerow(["model", "variant", *TARGETS])
        for experiment, variant in rows:
            row: list[Any] = [experiment, variant]
            for target in TARGETS:
                stats = aggregate_values.get((experiment, variant, target))
                row.append("" if stats is None else format_value(target, stats[0], stats[1]))
            writer.writerow(row)


def write_latex(
    destination: Path,
    aggregate_values: dict[tuple[str, str, str], tuple[float, float, int]],
    highlight: bool,
) -> None:
    rows = sorted({(key[0], key[1]) for key in aggregate_values})
    marks = highlight_cells(aggregate_values) if highlight else {}
    column_spec = "l" + "c" * len(TARGETS)
    lines = [
        r"\begin{tabular}{" + column_spec + "}",
        r"\toprule",
        "Model & " + " & ".join(TARGET_HEADERS[target] for target in TARGETS) + r" \\",
        r"\midrule",
    ]
    for experiment, variant in rows:
        cells = [latex_escape(display_name(experiment, variant))]
        for target in TARGETS:
            key = (experiment, variant, target)
            stats = aggregate_values.get(key)
            if stats is None:
                cells.append("--")
                continue
            value = format_value(target, stats[0], stats[1])
            if marks.get(key) == "best":
                value = r"\textbf{" + value + "}"
            elif marks.get(key) == "second":
                value = r"\underline{" + value + "}"
            cells.append(value)
        lines.append(" & ".join(cells) + r" \\")
    lines.extend([r"\bottomrule", r"\end{tabular}", ""])
    destination.parent.mkdir(parents=True, exist_ok=True)
    destination.write_text("\n".join(lines), encoding="utf-8")


def main() -> None:
    parser = argparse.ArgumentParser(description="Aggregate real QM9 test MAEs into CSV and LaTeX.")
    parser.add_argument("--root", default="logs/qm9")
    parser.add_argument("--output-dir", default="output/qm9")
    parser.add_argument("--no-highlight", action="store_true")
    args = parser.parse_args()

    records = collect(Path(args.root))
    if not records:
        raise FileNotFoundError(f"No valid result.json files found under {args.root}")
    values = aggregate(records)
    output_dir = Path(args.output_dir)
    csv_path = output_dir / "qm9_mae_summary.csv"
    latex_path = output_dir / "qm9_mae_table.tex"
    write_csv(csv_path, values)
    write_latex(latex_path, values, highlight=not args.no_highlight)
    print(f"Results: {len(records)} runs, {len(values)} model-target aggregates")
    print(f"Saved CSV: {csv_path}")
    print(f"Saved LaTeX: {latex_path}")


if __name__ == "__main__":
    main()
```
