## Supplementary material for "PG-MLD: Physics-Guided Molecular Representation Learning via Dynamic 3D Trajectory Distillation": Code and Data: plot_pca_figure5.py

```
from __future__ import annotations

import argparse
from dataclasses import dataclass
from pathlib import Path

import matplotlib as mpl
import matplotlib.pyplot as plt
from matplotlib.ticker import MaxNLocator
import numpy as np


FIGURE_WIDTH_IN = 3.25
FIGURE_HEIGHT_IN = 2.55
OUTPUT_DPI = 330

CLASS_COLORS = {0: "#3E7454", 1: "#0A48AB"}
FONT_FAMILY = "Arial"


@dataclass(frozen=True)
class PanelData:
    dataset: str
    variant: str
    coordinates: np.ndarray
    labels: np.ndarray
    dbi: float
    pca_overlap: float
    source: Path


def configure_matplotlib() -> None:
    mpl.rcParams.update(
        {
            "font.family": "sans-serif",
            "font.sans-serif": ["Arial", "Helvetica", "DejaVu Sans"],
            "axes.unicode_minus": False,
            "pdf.fonttype": 42,
            "ps.fonttype": 42,
            "svg.fonttype": "none",
            "figure.facecolor": "white",
            "axes.facecolor": "white",
        }
    )


def load_panel(path: Path, dataset: str, variant: str) -> PanelData:
    if not path.is_file():
        raise FileNotFoundError(f"PCA cache not found: {path}")
    cache = np.load(path, allow_pickle=True)
    required = {"tsne", "labels", "dbi_original", "dbi_tsne", "backend"}
    missing = sorted(required.difference(cache.files))
    if missing:
        raise ValueError(f"Missing arrays in {path}: {missing}")

    backend = str(np.asarray(cache["backend"]).item()).lower()
    if backend != "pca":
        raise ValueError(f"Expected cached PCA coordinates in {path}, got backend={backend!r}")

    coordinates = np.asarray(cache["tsne"], dtype=np.float64)
    labels = np.asarray(cache["labels"], dtype=np.int64).reshape(-1)
    if coordinates.ndim != 2 or coordinates.shape[1] != 2:
        raise ValueError(f"Expected [N, 2] coordinates in {path}, got {coordinates.shape}")
    if len(coordinates) != len(labels):
        raise ValueError(f"Coordinate/label length mismatch in {path}")

    valid = np.isfinite(coordinates).all(axis=1) & np.isin(labels, [0, 1])
    removed = int((~valid).sum())
    if removed:
        print(f"{path}: removed {removed} invalid cached points.")
    coordinates = coordinates[valid]
    labels = labels[valid]
    if len(coordinates) == 0:
        raise ValueError(f"No valid PCA points remain in {path}")

    return PanelData(
        dataset=dataset,
        variant=variant,
        coordinates=coordinates,
        labels=labels,
        dbi=float(np.asarray(cache["dbi_original"]).item()),
        pca_overlap=float(np.asarray(cache["dbi_tsne"]).item()),
        source=path,
    )


def data_limits(arrays: list[np.ndarray], margin: float = 0.045) -> tuple[tuple[float, float], tuple[float, float]]:
    merged = np.concatenate(arrays, axis=0)
    xmin, xmax = float(merged[:, 0].min()), float(merged[:, 0].max())
    ymin, ymax = float(merged[:, 1].min()), float(merged[:, 1].max())
    xpad = max((xmax - xmin) * margin, 1e-6)
    ypad = max((ymax - ymin) * margin, 1e-6)
    return (xmin - xpad, xmax + xpad), (ymin - ypad, ymax + ypad)


def major_ticks(limits: tuple[float, float]) -> np.ndarray:
    locator = MaxNLocator(nbins=3, steps=[1, 2, 2.5, 5, 10])
    ticks = locator.tick_values(*limits)
    return ticks[(ticks >= limits[0]) & (ticks <= limits[1])]


def _numpy_gaussian_kde(samples: np.ndarray, query: np.ndarray) -> np.ndarray:
    samples = np.asarray(samples, dtype=np.float64)
    query = np.asarray(query, dtype=np.float64)
    if samples.ndim == 1:
        samples = samples[None, :]
    if query.ndim == 1:
        query = query[None, :]
    dimension, count = samples.shape
    if count < max(3, dimension + 1):
        raise ValueError(f"Too few samples for {dimension}D KDE: {count}")

    covariance = np.atleast_2d(np.cov(samples, bias=False))
    factor = count ** (-1.0 / (dimension + 4.0))
    covariance = covariance * factor**2
    scale = max(float(np.trace(covariance)) / max(dimension, 1), 1.0)
    covariance += np.eye(dimension) * scale * 1e-8
    inverse = np.linalg.inv(covariance)
    determinant = max(float(np.linalg.det(covariance)), 1e-24)
    normalizer = np.sqrt((2.0 * np.pi) ** dimension * determinant)

    result = np.empty(query.shape[1], dtype=np.float64)
    sample_rows = samples.T
    for start in range(0, query.shape[1], 2048):
        stop = min(start + 2048, query.shape[1])
        delta = query[:, start:stop].T[:, None, :] - sample_rows[None, :, :]
        exponent = np.einsum("mni,ij,mnj->mn", delta, inverse, delta, optimize=True)
        result[start:stop] = np.exp(-0.5 * exponent).mean(axis=1) / normalizer
    return result


def kde_density(samples: np.ndarray, query: np.ndarray) -> np.ndarray:
    try:
        from scipy.stats import gaussian_kde

        return np.asarray(gaussian_kde(samples)(query), dtype=np.float64)
    except Exception as exc:
        if not getattr(kde_density, "_fallback_reported", False):
            print(f"SciPy gaussian_kde unavailable; using NumPy fallback: {exc}")
            kde_density._fallback_reported = True
        return _numpy_gaussian_kde(samples, query)


def density_levels(density: np.ndarray) -> np.ndarray:
    positive = density[np.isfinite(density) & (density > 0)]
    if positive.size == 0:
        return np.asarray([])
    return np.unique(np.quantile(positive, [0.58, 0.72, 0.84, 0.92, 0.97]))


def draw_2d_density(
    axis: plt.Axes,
    coordinates: np.ndarray,
    labels: np.ndarray,
    xlim: tuple[float, float],
    ylim: tuple[float, float],
) -> None:
    xgrid = np.linspace(*xlim, 120)
    ygrid = np.linspace(*ylim, 120)
    xx, yy = np.meshgrid(xgrid, ygrid)
    query = np.vstack([xx.ravel(), yy.ravel()])
    for cls in (0, 1):
        points = coordinates[labels == cls]
        if len(points) < 5:
            continue
        try:
            density = kde_density(points.T, query).reshape(xx.shape)
            levels = density_levels(density)
            if len(levels) < 2:
                continue
            axis.contourf(
                xx,
                yy,
                density,
                levels=levels,
                colors=[CLASS_COLORS[cls]],
                alpha=0.055,
                antialiased=True,
                zorder=1,
            )
            axis.contour(
                xx,
                yy,
                density,
                levels=levels,
                colors=[CLASS_COLORS[cls]],
                linewidths=0.8,
                alpha=0.14,
                zorder=2,
            )
        except Exception as exc:
            print(f"Skipping {cls=} 2D density: {exc}")


def draw_marginals(
    top_axis: plt.Axes,
    right_axis: plt.Axes,
    coordinates: np.ndarray,
    labels: np.ndarray,
    xlim: tuple[float, float],
    ylim: tuple[float, float],
) -> None:
    xgrid = np.linspace(*xlim, 320)
    ygrid = np.linspace(*ylim, 320)
    top_max = 0.0
    right_max = 0.0
    for cls in (0, 1):
        points = coordinates[labels == cls]
        try:
            xdensity = kde_density(points[:, 0], xgrid)
            top_axis.plot(xgrid, xdensity, color=CLASS_COLORS[cls], linewidth=0.8)
            top_axis.fill_between(xgrid, 0, xdensity, color=CLASS_COLORS[cls], alpha=0.12)
            top_max = max(top_max, float(xdensity.max()))
        except Exception as exc:
            print(f"Skipping {cls=} top density: {exc}")
        try:
            ydensity = kde_density(points[:, 1], ygrid)
            right_axis.plot(ydensity, ygrid, color=CLASS_COLORS[cls], linewidth=0.8)
            right_axis.fill_betweenx(ygrid, 0, ydensity, color=CLASS_COLORS[cls], alpha=0.12)
            right_max = max(right_max, float(ydensity.max()))
        except Exception as exc:
            print(f"Skipping {cls=} right density: {exc}")

    top_axis.set_xlim(xlim)
    top_axis.set_ylim(0, top_max * 1.06 if top_max else 1.0)
    right_axis.set_ylim(ylim)
    right_axis.set_xlim(0, right_max * 1.06 if right_max else 1.0)
    for axis in (top_axis, right_axis):
        axis.set_xticks([])
        axis.set_yticks([])
        for spine in axis.spines.values():
            spine.set_visible(False)


def style_main_axis(
    axis: plt.Axes,
    xlim: tuple[float, float],
    ylim: tuple[float, float],
    show_ylabel: bool,
    show_xlabel: bool,
) -> None:
    axis.set_xlim(xlim)
    axis.set_ylim(ylim)
    axis.set_xticks(major_ticks(xlim))
    axis.set_yticks(major_ticks(ylim))
    axis.tick_params(axis="both", labelsize=5.2, width=0.8, length=1.8, pad=0.8)
    for label in [*axis.get_xticklabels(), *axis.get_yticklabels()]:
        label.set_fontfamily(FONT_FAMILY)
        label.set_fontweight("normal")

    if show_ylabel:
        axis.set_ylabel("PCA 2", fontsize=6.2, fontfamily=FONT_FAMILY, labelpad=0.8)
    else:
        axis.tick_params(axis="y", labelleft=False)
    if show_xlabel:
        axis.set_xlabel("PCA 1", fontsize=6.2, fontfamily=FONT_FAMILY, labelpad=1.0)
    else:
        axis.tick_params(axis="x", labelbottom=False)

    axis.grid(True, color="#D9DEE3", linewidth=0.45, alpha=0.50, zorder=0)
    axis.spines["top"].set_visible(False)
    axis.spines["right"].set_visible(False)
    axis.spines["left"].set_linewidth(0.8)
    axis.spines["bottom"].set_linewidth(0.8)


def add_panel(
    fig: plt.Figure,
    panel: PanelData,
    left: float,
    bottom: float,
    main_width: float,
    main_height: float,
    top_height: float,
    right_width: float,
    marginal_gap: float,
    xlim: tuple[float, float],
    ylim: tuple[float, float],
    show_ylabel: bool,
    show_xlabel: bool,
) -> None:
    def rect(x: float, y: float, width: float, height: float) -> list[float]:
        return [
            x / FIGURE_WIDTH_IN,
            y / FIGURE_HEIGHT_IN,
            width / FIGURE_WIDTH_IN,
            height / FIGURE_HEIGHT_IN,
        ]

    main_axis = fig.add_axes(rect(left, bottom, main_width, main_height))
    top_axis = fig.add_axes(
        rect(left, bottom + main_height + marginal_gap, main_width, top_height)
    )
    right_axis = fig.add_axes(
        rect(left + main_width + marginal_gap, bottom, right_width, main_height)
    )

    draw_2d_density(main_axis, panel.coordinates, panel.labels, xlim, ylim)
    for cls in (0, 1):
        points = panel.coordinates[panel.labels == cls]
        main_axis.scatter(
            points[:, 0],
            points[:, 1],
            s=3.0,
            color=CLASS_COLORS[cls],
            alpha=0.72,
            edgecolors="none",
            rasterized=True,
            zorder=4,
        )
    style_main_axis(main_axis, xlim, ylim, show_ylabel, show_xlabel)
    draw_marginals(top_axis, right_axis, panel.coordinates, panel.labels, xlim, ylim)


def render_figure(
    panels: dict[tuple[str, str], PanelData],
    output_dir: Path,
) -> list[Path]:
    configure_matplotlib()
    fig = plt.figure(figsize=(FIGURE_WIDTH_IN, FIGURE_HEIGHT_IN), facecolor="white")

    left_margin = 0.285
    right_margin = 0.025
    column_gap = 0.235
    panel_width = (FIGURE_WIDTH_IN - left_margin - right_margin - column_gap) / 2.0
    marginal_gap = 0.018
    right_kde_width = 0.145
    main_width = panel_width - marginal_gap - right_kde_width

    bottom_margin = 0.220
    top_margin = 0.020
    row_gap = 0.080
    row_height = (FIGURE_HEIGHT_IN - bottom_margin - top_margin - row_gap) / 2.0
    main_height = 0.765
    top_kde_height = 0.100
    title_band = row_height - main_height - marginal_gap - top_kde_height

    column_lefts = [
        left_margin,
        left_margin + panel_width + column_gap,
    ]
    row_bottoms = [
        bottom_margin + row_height + row_gap,
        bottom_margin,
    ]

    row_specs = [
        ("BACE", row_bottoms[0]),
        ("BBBP", row_bottoms[1]),
    ]
    row_limits = {
        dataset: data_limits(
            [
                panels[(dataset, "LLM")].coordinates,
                panels[(dataset, "PG-MLD")].coordinates,
            ]
        )
        for dataset, _ in row_specs
    }

    for row_index, (dataset, row_bottom) in enumerate(row_specs):
        xlim, ylim = row_limits[dataset]
        for column_index, variant in enumerate(("LLM", "PG-MLD")):
            panel = panels[(dataset, variant)]
            add_panel(
                fig=fig,
                panel=panel,
                left=column_lefts[column_index],
                bottom=row_bottom,
                main_width=main_width,
                main_height=main_height,
                top_height=top_kde_height,
                right_width=right_kde_width,
                marginal_gap=marginal_gap,
                xlim=xlim,
                ylim=ylim,
                show_ylabel=column_index == 0,
                show_xlabel=row_index == 1,
            )

            center_x = (
                column_lefts[column_index] + 0.5 * main_width
            ) / FIGURE_WIDTH_IN
            title_top = (
                row_bottom
                + main_height
                + marginal_gap
                + top_kde_height
                + title_band
            ) / FIGURE_HEIGHT_IN
            model_title = (
                f"{dataset} LLM"
                if variant == "LLM"
                else "PG-MLD distilled"
            )
            fig.text(
                center_x,
                title_top - 0.004,
                model_title,
                ha="center",
                va="top",
                fontsize=9.5,
                fontweight="bold",
                fontfamily=FONT_FAMILY,
            )
            fig.text(
                center_x,
                title_top - 0.050,
                f"DBI {panel.dbi:.2f} | PCA {panel.pca_overlap:.2f}",
                ha="center",
                va="top",
                fontsize=9,
                fontweight="normal",
                fontfamily=FONT_FAMILY,
            )

    output_dir.mkdir(parents=True, exist_ok=True)
    base = output_dir / "figure5_pca_comparison"
    paths = [base.with_suffix(".png")]
    fig.savefig(paths[0], dpi=OUTPUT_DPI, facecolor="white")
    plt.close(fig)
    return paths


def build_parser() -> argparse.ArgumentParser:
    parser = argparse.ArgumentParser(
        description="Replot cached BACE/BBBP PCA coordinates as AAAI Figure 5."
    )
    parser.add_argument(
        "--bace-original",
        default="output/figures/pca_chemblm_original/bace/bace_chemblm_all_penultimate_balanced.npz",
    )
    parser.add_argument(
        "--bace-distilled",
        default="output/figures/pca_chemblm_all/bace/bace_chemblm_all_penultimate_balanced.npz",
    )
    parser.add_argument(
        "--bbbp-original",
        default="output/figures/pca_chemblm_original/bbbp/bbbp_chemblm_all_penultimate_balanced.npz",
    )
    parser.add_argument(
        "--bbbp-distilled",
        default="output/figures/pca_chemblm_all/bbbp/bbbp_chemblm_all_penultimate_balanced.npz",
    )
    parser.add_argument(
        "--output-dir",
        default="output/figures/pca_figure5_publication",
    )
    return parser


def main() -> None:
    args = build_parser().parse_args()
    panels = {
        ("BACE", "LLM"): load_panel(Path(args.bace_original), "BACE", "LLM"),
        ("BACE", "PG-MLD"): load_panel(
            Path(args.bace_distilled), "BACE", "PG-MLD"
        ),
        ("BBBP", "LLM"): load_panel(Path(args.bbbp_original), "BBBP", "LLM"),
        ("BBBP", "PG-MLD"): load_panel(
            Path(args.bbbp_distilled), "BBBP", "PG-MLD"
        ),
    }
    paths = render_figure(panels, Path(args.output_dir))

    for panel in panels.values():
        print(
            f"{panel.dataset} {panel.variant}: "
            f"points={len(panel.coordinates)} "
            f"DBI={panel.dbi:.12g} PCA-overlap={panel.pca_overlap:.12g} "
            f"source={panel.source}"
        )
    for path in paths:
        print(f"Saved: {path}")
    print("Cached PCA coordinates were plotted directly; PCA was not recomputed.")


if __name__ == "__main__":
    main()
```
