## Supplementary material for "PG-MLD: Physics-Guided Molecular Representation Learning via Dynamic 3D Trajectory Distillation": Code and Data: plot_shap_figure4.py

```
from __future__ import annotations

import argparse
from pathlib import Path

import matplotlib as mpl
import matplotlib.pyplot as plt
import numpy as np
import pandas as pd


ORIGINAL_VARIANT = "Original ChemBLM"
DISTILLED_VARIANT = "PG-MLD distilled"
DISPLAY_NAMES = {
    ORIGINAL_VARIANT: "Original LLM",
    DISTILLED_VARIANT: "PG-MLD distilled",
}
NEGATIVE_COLOR = "#0A48AB"
POSITIVE_COLOR = "#E37019"
FONT_FAMILY = "Arial"

FIGURE_WIDTH_IN = 3.25
FIGURE_HEIGHT_IN = 820.0 / 330.0
OUTPUT_DPI = 330
TOKEN_FONT_SIZE = 5.2


def configure_matplotlib() -> None:
    mpl.rcParams.update(
        {
            "font.family": "sans-serif",
            "font.sans-serif": ["Arial", "Helvetica", "DejaVu Sans"],
            "axes.unicode_minus": False,
            "pdf.fonttype": 42,
            "ps.fonttype": 42,
            "svg.fonttype": "none",
            "figure.facecolor": "white",
            "axes.facecolor": "white",
        }
    )


def load_cached_explanations(
    csv_path: Path,
    sample_id: int,
    top_k: int,
) -> tuple[list[str], dict[str, np.ndarray], int, dict[str, float]]:
    if not csv_path.is_file():
        raise FileNotFoundError(f"Cached token SHAP CSV not found: {csv_path}")

    frame = pd.read_csv(csv_path)
    required = {
        "variant",
        "dataset_index",
        "true_label",
        "predicted_probability",
        "token_index",
        "token",
        "normalized_shap_value",
    }
    missing = sorted(required.difference(frame.columns))
    if missing:
        raise ValueError(f"Missing columns in {csv_path}: {missing}")

    sample = frame.loc[frame["dataset_index"].astype(int) == int(sample_id)].copy()
    if sample.empty:
        available = sorted(frame["dataset_index"].astype(int).unique().tolist())
        raise ValueError(
            f"Sample {sample_id} is absent from {csv_path}. "
            f"Available sample IDs include: {available[:20]}"
        )

    variants = [ORIGINAL_VARIANT, DISTILLED_VARIANT]
    ordered: dict[str, pd.DataFrame] = {}
    for variant in variants:
        subset = sample.loc[sample["variant"] == variant].sort_values("token_index")
        if subset.empty:
            raise ValueError(f"Variant {variant!r} is absent for sample {sample_id}")
        if subset["token_index"].duplicated().any():
            raise ValueError(f"Duplicate token indices for {variant!r}, sample {sample_id}")
        ordered[variant] = subset.reset_index(drop=True)

    reference_indices = ordered[ORIGINAL_VARIANT]["token_index"].astype(int).tolist()
    reference_tokens = ordered[ORIGINAL_VARIANT]["token"].astype(str).tolist()
    for variant in variants[1:]:
        indices = ordered[variant]["token_index"].astype(int).tolist()
        tokens = ordered[variant]["token"].astype(str).tolist()
        if indices != reference_indices or tokens != reference_tokens:
            raise ValueError(
                "Original and distilled cached explanations do not have identical "
                "token indices and token order."
            )

    all_values = {
        variant: ordered[variant]["normalized_shap_value"].astype(float).to_numpy()
        for variant in variants
    }
    max_abs = np.maximum(
        np.abs(all_values[ORIGINAL_VARIANT]),
        np.abs(all_values[DISTILLED_VARIANT]),
    )
    keep = min(int(top_k), len(reference_tokens))
    chosen = np.argsort(max_abs)[-keep:]
    chosen = chosen[np.argsort(max_abs[chosen])]

    labels = [reference_tokens[index] for index in chosen]
    selected_values = {variant: values[chosen] for variant, values in all_values.items()}

    labels_by_variant = {
        variant: ordered[variant]["true_label"].astype(int).unique().tolist()
        for variant in variants
    }
    if any(len(values) != 1 for values in labels_by_variant.values()):
        raise ValueError(f"Inconsistent true labels for sample {sample_id}: {labels_by_variant}")
    true_labels = {values[0] for values in labels_by_variant.values()}
    if len(true_labels) != 1:
        raise ValueError(f"Variants disagree on the true label: {labels_by_variant}")
    true_label = int(next(iter(true_labels)))

    probabilities = {}
    for variant in variants:
        values = ordered[variant]["predicted_probability"].astype(float).unique()
        if len(values) != 1:
            raise ValueError(f"Inconsistent prediction probability for {variant!r}")
        probabilities[variant] = float(values[0])

    return labels, selected_values, true_label, probabilities


def shared_axis_limits(values: dict[str, np.ndarray]) -> tuple[tuple[float, float], np.ndarray]:
    merged = np.concatenate(list(values.values()))
    lower = min(0.0, float(np.min(merged)))
    upper = max(0.0, float(np.max(merged)))
    span = max(upper - lower, 1e-6)
    limits = (lower - 0.055 * span, upper + 0.055 * span)

    return limits, np.asarray([0.0], dtype=float)


def draw_panel(
    axis: plt.Axes,
    values: np.ndarray,
    tokens: list[str],
    xlim: tuple[float, float],
    xticks: np.ndarray,
) -> None:
    y = np.arange(len(tokens))
    colors = [POSITIVE_COLOR if value >= 0 else NEGATIVE_COLOR for value in values]
    axis.barh(y, values, height=0.58, color=colors, edgecolor="none", zorder=3)
    axis.axvline(0.0, color="#555B61", linewidth=0.8, zorder=4)
    axis.set_xlim(xlim)
    axis.set_xticks([])
    axis.set_yticks(y, tokens)

    axis.grid(False)
    axis.set_axisbelow(True)
    axis.spines["top"].set_visible(False)
    axis.spines["right"].set_visible(False)
    axis.spines["left"].set_linewidth(0.8)
    axis.spines["bottom"].set_visible(True)
    axis.spines["bottom"].set_linewidth(0.8)
    axis.tick_params(axis="x", bottom=False, labelbottom=False)
    axis.tick_params(
        axis="y",
        labelsize=TOKEN_FONT_SIZE,
        width=0.8,
        length=1.8,
        pad=1.0,
    )
    axis.set_ylim(-0.65, len(tokens) - 0.35)
    for label in [*axis.get_xticklabels(), *axis.get_yticklabels()]:
        label.set_fontfamily(FONT_FAMILY)
        label.set_fontweight("normal")


def plot_figure4(
    tokens: list[str],
    values: dict[str, np.ndarray],
    true_label: int,
    probabilities: dict[str, float],
    output_dir: Path,
    sample_id: int,
) -> list[Path]:
    xlim, xticks = shared_axis_limits(values)
    fig = plt.figure(figsize=(FIGURE_WIDTH_IN, FIGURE_HEIGHT_IN), facecolor="white")

    left_margin_in = 0.240
    right_margin_in = 0.170
    panel_gap_in = 0.380
    bottom_margin_in = 0.225
    panel_top_in = 2.100
    panel_height_in = panel_top_in - bottom_margin_in
    panel_width_in = (
        FIGURE_WIDTH_IN - left_margin_in - right_margin_in - panel_gap_in
    ) / 2.0
    panel_lefts = [
        left_margin_in,
        left_margin_in + panel_width_in + panel_gap_in,
    ]

    variants = [ORIGINAL_VARIANT, DISTILLED_VARIANT]
    for left_in, variant in zip(panel_lefts, variants):
        axis = fig.add_axes(
            [
                left_in / FIGURE_WIDTH_IN,
                bottom_margin_in / FIGURE_HEIGHT_IN,
                panel_width_in / FIGURE_WIDTH_IN,
                panel_height_in / FIGURE_HEIGHT_IN,
            ]
        )
        draw_panel(axis, values[variant], tokens, xlim, xticks)
        center_x = (left_in + 0.5 * panel_width_in) / FIGURE_WIDTH_IN
        fig.text(
            center_x,
            0.975,
            DISPLAY_NAMES[variant],
            ha="center",
            va="top",
            fontsize=9.5,
            fontweight="bold",
            fontfamily=FONT_FAMILY,
            linespacing=0.92,
        )
        fig.text(
            center_x,
            0.900,
            f"true={true_label}, p(class 1)={probabilities[variant]:.3f}",
            ha="center",
            va="top",
            fontsize=9,
            fontweight="normal",
            fontfamily=FONT_FAMILY,
        )

    output_dir.mkdir(parents=True, exist_ok=True)
    base = output_dir / f"figure4_sample_{sample_id:05d}"
    paths = [base.with_suffix(".png")]
    fig.savefig(paths[0], dpi=OUTPUT_DPI, facecolor="white")
    plt.close(fig)
    return paths


def build_parser() -> argparse.ArgumentParser:
    parser = argparse.ArgumentParser(
        description=(
            "Replot cached ChemBLM token SHAP values as an AAAI single-column "
            "two-panel figure without recomputing SHAP."
        )
    )
    parser.add_argument(
        "--input-csv",
        default="output/figures/shap/bace/chemblm/comparison_normalized/token_shap_values.csv",
    )
    parser.add_argument("--sample-id", type=int, default=962)
    parser.add_argument("--top-k", type=int, default=20)
    parser.add_argument(
        "--output-dir",
        default="output/figures/shap/bace/chemblm/figure4_publication",
    )
    return parser


def main() -> None:
    args = build_parser().parse_args()
    configure_matplotlib()
    input_csv = Path(args.input_csv)
    tokens, values, true_label, probabilities = load_cached_explanations(
        input_csv,
        sample_id=args.sample_id,
        top_k=args.top_k,
    )
    paths = plot_figure4(
        tokens=tokens,
        values=values,
        true_label=true_label,
        probabilities=probabilities,
        output_dir=Path(args.output_dir),
        sample_id=args.sample_id,
    )

    print(f"Source CSV: {input_csv}")
    print(f"Sample: {args.sample_id}")
    print(f"True label: {true_label}")
    print(f"Tokens plotted ({len(tokens)}): {tokens}")
    for variant in (ORIGINAL_VARIANT, DISTILLED_VARIANT):
        print(f"{DISPLAY_NAMES[variant]} probability: {probabilities[variant]:.12g}")
    for path in paths:
        print(f"Saved: {path}")
    print("SHAP values were loaded from cache; no SHAP computation was performed.")


if __name__ == "__main__":
    main()
```
