## Supplementary material for "PG-MLD: Physics-Guided Molecular Representation Learning via Dynamic 3D Trajectory Distillation": Code and Data: representation_neighbor_comparison.py

```
from __future__ import annotations

import argparse
import csv
import sys
from pathlib import Path
from typing import Any

PROJECT_ROOT = Path(__file__).resolve().parents[1]
if str(PROJECT_ROOT) not in sys.path:
    sys.path.insert(0, str(PROJECT_ROOT))

import matplotlib as mpl
import matplotlib.pyplot as plt
from matplotlib.patches import FancyBboxPatch
import numpy as np
import pandas as pd


ORIGINAL_COLOR = "#E37019"
DISTILLED_COLOR = "#7D5AA9"
PLOT_FONT_FAMILY = "Arial"

# Fixed high-resolution publication canvas: exactly 4569 x 2000 px at 600 dpi.
PUBLICATION_DPI = 600
FIGURE_WIDTH_IN = 4569.0 / PUBLICATION_DPI
FIGURE_HEIGHT_IN = 2000.0 / PUBLICATION_DPI

# Four strictly aligned, equal-sized cards in each of two rows.
HORIZONTAL_MARGIN_IN = 0.060
QUERY_RETRIEVAL_GAP_IN = 0.200
NEIGHBOR_CARD_GAP_IN = 0.055
CARD_WIDTH_IN = (
    FIGURE_WIDTH_IN
    - 2.0 * HORIZONTAL_MARGIN_IN
    - QUERY_RETRIEVAL_GAP_IN
    - 2.0 * NEIGHBOR_CARD_GAP_IN
) / 4.0
CARD_HEIGHT_IN = 1.380
BOTTOM_ROW_Y_IN = 0.040
TOP_ROW_Y_IN = 1.600


def configure_matplotlib() -> None:
    mpl.rcParams.update(
        {
            "font.family": "sans-serif",
            "font.sans-serif": ["Arial", "Helvetica", "DejaVu Sans"],
            "axes.unicode_minus": False,
            "pdf.fonttype": 42,
            "ps.fonttype": 42,
            "svg.fonttype": "none",
            "figure.facecolor": "white",
            "axes.facecolor": "white",
        }
    )


def load_embeddings(path: str | Path) -> tuple[np.ndarray, list[str] | None]:
    path = Path(path)
    if not path.is_file():
        raise FileNotFoundError(f"Embedding file not found: {path}")
    with np.load(path, allow_pickle=False) as archive:
        if "embeddings" not in archive:
            raise KeyError(f"{path} does not contain an 'embeddings' array")
        embeddings = np.asarray(archive["embeddings"], dtype=np.float32)
        embedded_smiles = archive["smiles"].astype(str).tolist() if "smiles" in archive else None
    if embeddings.ndim != 2:
        raise ValueError(f"Expected [N, D] embeddings in {path}, got {embeddings.shape}")
    if not np.isfinite(embeddings).all():
        raise ValueError(f"Embeddings contain NaN or infinity: {path}")
    return embeddings, embedded_smiles


def load_smiles(path: str | Path, smiles_col: str) -> list[str]:
    path = Path(path)
    if not path.is_file():
        raise FileNotFoundError(f"Metadata CSV not found: {path}")
    frame = pd.read_csv(path)
    candidates = [smiles_col, "canonical_smiles", "smiles", "input_smiles"]
    column = next((name for name in candidates if name and name in frame.columns), None)
    if column is None:
        raise ValueError(f"No SMILES column found in {path}. Columns: {list(frame.columns)}")
    return frame[column].astype(str).tolist()


def l2_normalize(embeddings: np.ndarray) -> np.ndarray:
    norms = np.linalg.norm(embeddings, axis=1, keepdims=True)
    return embeddings / np.maximum(norms, 1e-12)


def nearest_neighbors(normalized: np.ndarray, query_index: int, top_k: int) -> tuple[list[int], list[float]]:
    similarities = normalized @ normalized[query_index]
    order = np.argsort(-similarities)
    selected = [int(index) for index in order if int(index) != int(query_index)][: int(top_k)]
    return selected, [float(similarities[index]) for index in selected]


def fingerprint_similarities(query_smiles: str, neighbor_smiles: list[str]) -> tuple[list[float], list[float]]:
    from rdkit import Chem, DataStructs

    query_mol = Chem.MolFromSmiles(query_smiles)
    if query_mol is None:
        raise ValueError(f"RDKit could not parse query SMILES: {query_smiles}")

    try:
        from rdkit.Chem import rdFingerprintGenerator

        morgan = rdFingerprintGenerator.GetMorganGenerator(radius=2, fpSize=2048)
        query_ecfp = morgan.GetFingerprint(query_mol)
        make_ecfp = morgan.GetFingerprint
    except (ImportError, AttributeError):
        from rdkit.Chem import AllChem

        query_ecfp = AllChem.GetMorganFingerprintAsBitVect(query_mol, radius=2, nBits=2048)
        make_ecfp = lambda mol: AllChem.GetMorganFingerprintAsBitVect(mol, radius=2, nBits=2048)

    query_rdk = Chem.RDKFingerprint(query_mol)
    ecfp_values: list[float] = []
    rdk_values: list[float] = []
    for smiles in neighbor_smiles:
        mol = Chem.MolFromSmiles(smiles)
        if mol is None:
            ecfp_values.append(float("nan"))
            rdk_values.append(float("nan"))
            continue
        ecfp_values.append(float(DataStructs.TanimotoSimilarity(query_ecfp, make_ecfp(mol))))
        rdk_values.append(float(DataStructs.TanimotoSimilarity(query_rdk, Chem.RDKFingerprint(mol))))
    return ecfp_values, rdk_values


def molecule_image(smiles: str, size: tuple[int, int] = (1100, 700)) -> Any:
    from rdkit import Chem
    from rdkit.Chem import Draw, rdDepictor

    mol = Chem.MolFromSmiles(smiles)
    if mol is None:
        raise ValueError(f"RDKit could not parse SMILES: {smiles}")
    rdDepictor.Compute2DCoords(mol)
    options = Draw.MolDrawOptions()
    options.padding = 0.08
    options.bondLineWidth = 6.0
    if hasattr(options, "additionalAtomLabelPadding"):
        options.additionalAtomLabelPadding = 0.08
    return Draw.MolToImage(mol, size=size, fitImage=True, options=options)


def style_card(axis: plt.Axes, color: str, linewidth: float = 0.9) -> None:
    axis.set_xticks([])
    axis.set_yticks([])
    for spine in axis.spines.values():
        spine.set_visible(False)
    axis.set_facecolor("white")
    rounded_border = FancyBboxPatch(
        (0.0, 0.0),
        1.0,
        1.0,
        boxstyle="round,pad=0.006,rounding_size=0.025",
        transform=axis.transAxes,
        facecolor="none",
        edgecolor=color,
        linewidth=linewidth,
        clip_on=False,
        zorder=10,
    )
    axis.add_patch(rounded_border)


def draw_molecule_in_card(axis: plt.Axes, image: Any, reserve_metrics: bool = False) -> None:
    # Retrieved molecules reserve an independent bottom band for Cos/ECFP.
    bounds = [0.040, 0.105, 0.92, 0.82]
    if reserve_metrics:
        bounds = [0.045, 0.205, 0.91, 0.75]
    image_axis = axis.inset_axes(bounds, zorder=1)
    image_axis.imshow(image)
    image_axis.set_axis_off()


def draw_compact_arrow(fig: plt.Figure, row_bottom_in: float, color: str) -> None:
    """Draw a small Query-to-retrieval arrow without allocating a wide column."""
    arrow_width_in = QUERY_RETRIEVAL_GAP_IN - 0.030
    arrow_axis = add_physical_axis(
        fig,
        FIGURE_WIDTH_IN,
        FIGURE_HEIGHT_IN,
        HORIZONTAL_MARGIN_IN + CARD_WIDTH_IN + 0.015,
        row_bottom_in + CARD_HEIGHT_IN * 0.50 - 0.090,
        arrow_width_in,
        0.180,
    )
    arrow_axis.set_axis_off()
    arrow_axis.annotate(
        "",
        xy=(0.94, 0.50),
        xytext=(0.06, 0.50),
        xycoords="axes fraction",
        arrowprops={
            "arrowstyle": "-|>",
            "color": color,
            "linewidth": 1.15,
            "mutation_scale": 10.0,
            "shrinkA": 0.0,
            "shrinkB": 0.0,
        },
    )


def add_physical_axis(
    fig: plt.Figure,
    figure_width_in: float,
    figure_height_in: float,
    left_in: float,
    bottom_in: float,
    width_in: float,
    height_in: float,
) -> plt.Axes:
    """Create an axis whose dimensions are specified in physical inches."""
    return fig.add_axes(
        [
            left_in / figure_width_in,
            bottom_in / figure_height_in,
            width_in / figure_width_in,
            height_in / figure_height_in,
        ]
    )


def draw_publication_row(
    fig: plt.Figure,
    row_bottom_in: float,
    row_label: str,
    color: str,
    query_image: Any,
    smiles: list[str],
    result: dict[str, Any],
) -> None:
    card_contents: list[tuple[str, Any, float | None, float | None]] = [
        ("Query", query_image, None, None)
    ]
    card_contents.extend(
        (
            f"Top-{rank}",
            molecule_image(smiles[neighbor_index]),
            float(cosine),
            float(ecfp),
        )
        for rank, (neighbor_index, cosine, ecfp) in enumerate(
            zip(result["indices"], result["cosine"], result["ecfp"]),
            start=1,
        )
    )
    draw_compact_arrow(fig, row_bottom_in, color)

    for column, (card_label, image, cosine, ecfp) in enumerate(card_contents):
        if column == 0:
            left_in = HORIZONTAL_MARGIN_IN
        else:
            left_in = (
                HORIZONTAL_MARGIN_IN
                + CARD_WIDTH_IN
                + QUERY_RETRIEVAL_GAP_IN
                + (column - 1) * (CARD_WIDTH_IN + NEIGHBOR_CARD_GAP_IN)
            )
        axis = add_physical_axis(
            fig,
            FIGURE_WIDTH_IN,
            FIGURE_HEIGHT_IN,
            left_in,
            row_bottom_in,
            CARD_WIDTH_IN,
            CARD_HEIGHT_IN,
        )
        style_card(axis, color, linewidth=0.9)
        if column == 0:
            draw_molecule_in_card(axis, image, reserve_metrics=False)
            axis.set_title(
                row_label,
                color="black",
                fontsize=9.5,
                fontweight="bold",
                fontfamily=PLOT_FONT_FAMILY,
                pad=3.0,
            )
            axis.text(
                0.5,
                0.026,
                card_label,
                transform=axis.transAxes,
                ha="center",
                va="bottom",
                fontsize=9,
                fontweight="bold",
                fontfamily=PLOT_FONT_FAMILY,
                zorder=20,
                bbox={
                    "boxstyle": "square,pad=0.08",
                    "facecolor": "white",
                    "edgecolor": "none",
                    "alpha": 0.94,
                },
            )
            continue

        draw_molecule_in_card(axis, image, reserve_metrics=True)
        axis.set_title(
            card_label,
            color=color,
            fontsize=9,
            fontweight="bold",
            fontfamily=PLOT_FONT_FAMILY,
            pad=3.0,
        )
        axis.plot(
            [0.025, 0.975],
            [0.205, 0.205],
            transform=axis.transAxes,
            color=mpl.colors.to_rgba(color, 0.45),
            linewidth=0.65,
            zorder=12,
            solid_capstyle="round",
        )
        axis.text(
            0.045,
            0.092,
            f"Cos = {cosine:.3f}\nECFP = {ecfp:.3f}",
            transform=axis.transAxes,
            ha="left",
            va="center",
            fontsize=9,
            linespacing=1.05,
            fontfamily=PLOT_FONT_FAMILY,
            zorder=20,
        )


def plot_comparison(
    query_index: int,
    smiles: list[str],
    results: dict[str, dict[str, Any]],
    output_dir: Path,
    original_color: str,
    distilled_color: str,
) -> list[Path]:
    top_k = len(next(iter(results.values()))["indices"])
    if top_k != 3:
        raise ValueError(f"The AAAI publication layout requires exactly three neighbors, got {top_k}.")

    variant_specs = [
        ("Original ChemBLM", "Original LLM", original_color),
        ("PG-MLD distilled", "PG-MLD distilled", distilled_color),
    ]
    query_image = molecule_image(smiles[query_index])
    output_dir.mkdir(parents=True, exist_ok=True)
    fig = plt.figure(figsize=(FIGURE_WIDTH_IN, FIGURE_HEIGHT_IN), facecolor="white")
    for variant, row_label, color in variant_specs:
        row_bottom_in = TOP_ROW_Y_IN if variant == "Original ChemBLM" else BOTTOM_ROW_Y_IN
        draw_publication_row(
            fig=fig,
            row_bottom_in=row_bottom_in,
            row_label=row_label,
            color=color,
            query_image=query_image,
            smiles=smiles,
            result=results[variant],
        )

    base = output_dir / f"neighbor_comparison_query_{query_index:05d}"
    paths = [base.with_suffix(".png")]
    # Manual physical placement already removes excess whitespace. Avoid a
    # tight bounding box so the requested 6.75-inch canvas stays exact.
    fig.savefig(paths[0], dpi=PUBLICATION_DPI, facecolor="white")
    plt.close(fig)
    return paths


def save_table(
    path: Path,
    query_index: int,
    smiles: list[str],
    results: dict[str, dict[str, Any]],
) -> None:
    path.parent.mkdir(parents=True, exist_ok=True)
    with path.open("w", newline="", encoding="utf-8") as handle:
        writer = csv.writer(handle)
        writer.writerow(
            ["variant", "query_index", "query_smiles", "rank", "neighbor_index", "neighbor_smiles", "cosine", "ecfp_tanimoto", "rdk_tanimoto"]
        )
        for variant, result in results.items():
            for rank, (index, cosine, ecfp, rdk) in enumerate(
                zip(result["indices"], result["cosine"], result["ecfp"], result["rdk"]),
                start=1,
            ):
                writer.writerow(
                    [variant, query_index, smiles[query_index], rank, index, smiles[index], cosine, ecfp, rdk]
                )


def validate_color(value: str) -> str:
    try:
        mpl.colors.to_rgba(value)
    except ValueError as exc:
        raise argparse.ArgumentTypeError(f"Invalid Matplotlib color: {value}") from exc
    return value


def main() -> None:
    parser = argparse.ArgumentParser(
        description="Compare original ChemBLM and PG-MLD nearest-neighbor retrieval using fixed embeddings."
    )
    parser.add_argument("--original-npz", required=True)
    parser.add_argument("--distilled-npz", required=True)
    parser.add_argument("--csv", required=True, help="Metadata CSV aligned with both embedding arrays.")
    parser.add_argument("--smiles-col", default="canonical_smiles")
    parser.add_argument("--query-indices", required=True, help="Comma-separated row indices, e.g. 100,1000,5000")
    parser.add_argument("--top-k", type=int, default=3)
    parser.add_argument("--original-color", type=validate_color, default=ORIGINAL_COLOR)
    parser.add_argument("--distilled-color", type=validate_color, default=DISTILLED_COLOR)
    parser.add_argument("--output-dir", default="output/figures/representation_neighbor_comparison")
    args = parser.parse_args()

    configure_matplotlib()
    original, original_smiles = load_embeddings(args.original_npz)
    distilled, distilled_smiles = load_embeddings(args.distilled_npz)
    smiles = load_smiles(args.csv, args.smiles_col)
    if original.shape[0] != distilled.shape[0] or original.shape[0] != len(smiles):
        raise ValueError(
            "Original embeddings, distilled embeddings, and metadata must contain the same number of molecules: "
            f"original={len(original)}, distilled={len(distilled)}, metadata={len(smiles)}"
        )
    for name, embedded_smiles in (
        ("original", original_smiles),
        ("distilled", distilled_smiles),
    ):
        if embedded_smiles is None:
            print(f"Warning: {name} NPZ has no SMILES array; only row-count alignment can be checked.")
        elif len(embedded_smiles) != len(smiles):
            raise ValueError(
                f"{name} NPZ SMILES count ({len(embedded_smiles)}) does not match metadata ({len(smiles)})."
            )
        elif embedded_smiles != smiles:
            mismatch = next(
                index
                for index, (embedded, metadata) in enumerate(zip(embedded_smiles, smiles))
                if embedded != metadata
            )
            raise ValueError(
                f"{name} embeddings are not aligned with metadata at row {mismatch}: "
                f"NPZ={embedded_smiles[mismatch]!r}, CSV={smiles[mismatch]!r}"
            )
    if args.top_k != 3:
        raise ValueError("This AAAI publication layout requires --top-k 3.")

    normalized = {
        "Original ChemBLM": l2_normalize(original),
        "PG-MLD distilled": l2_normalize(distilled),
    }
    output_dir = Path(args.output_dir)
    query_indices = [int(value.strip()) for value in args.query_indices.split(",") if value.strip()]
    if not query_indices:
        raise ValueError("--query-indices must contain at least one integer")
    aggregate_rows: list[dict[str, Any]] = []
    for query_index in query_indices:
        if query_index < 0 or query_index >= len(smiles):
            raise IndexError(f"Query index out of range: {query_index}; molecules={len(smiles)}")
        results: dict[str, dict[str, Any]] = {}
        for variant, embeddings in normalized.items():
            indices, cosine = nearest_neighbors(embeddings, query_index, args.top_k)
            ecfp, rdk = fingerprint_similarities(smiles[query_index], [smiles[index] for index in indices])
            results[variant] = {"indices": indices, "cosine": cosine, "ecfp": ecfp, "rdk": rdk}
            for rank, (neighbor_index, cosine_value, ecfp_value, rdk_value) in enumerate(
                zip(indices, cosine, ecfp, rdk),
                start=1,
            ):
                aggregate_rows.append(
                    {
                        "variant": variant,
                        "query_index": query_index,
                        "rank": rank,
                        "neighbor_index": neighbor_index,
                        "cosine": cosine_value,
                        "ecfp_tanimoto": ecfp_value,
                        "rdk_tanimoto": rdk_value,
                    }
                )

        table_path = output_dir / f"neighbor_comparison_query_{query_index:05d}.csv"
        save_table(table_path, query_index, smiles, results)
        paths = plot_comparison(
            query_index=query_index,
            smiles=smiles,
            results=results,
            output_dir=output_dir,
            original_color=args.original_color,
            distilled_color=args.distilled_color,
        )
        print(f"Query {query_index}: {smiles[query_index]}")
        print(f"Saved comparison table: {table_path}")
        for path in paths:
            print(f"Saved comparison figure: {path}")

    aggregate_frame = pd.DataFrame(aggregate_rows)
    aggregate_path = output_dir / "neighbor_comparison_all.csv"
    aggregate_frame.to_csv(aggregate_path, index=False)
    summary_rows: list[dict[str, Any]] = []
    for variant, group in aggregate_frame.groupby("variant", sort=False):
        summary_rows.append(
            {
                "variant": variant,
                "queries": int(group["query_index"].nunique()),
                "retrieved_neighbors": int(len(group)),
                "mean_ecfp_tanimoto": float(group["ecfp_tanimoto"].mean()),
                "std_ecfp_tanimoto": float(group["ecfp_tanimoto"].std(ddof=0)),
                "mean_rdk_tanimoto": float(group["rdk_tanimoto"].mean()),
                "std_rdk_tanimoto": float(group["rdk_tanimoto"].std(ddof=0)),
            }
        )
    summary_path = output_dir / "neighbor_comparison_summary.csv"
    pd.DataFrame(summary_rows).to_csv(summary_path, index=False)
    print(f"Saved aggregate retrieval table: {aggregate_path}")
    print(f"Saved retrieval summary: {summary_path}")
    for row in summary_rows:
        print(
            f"{row['variant']}: ECFP={row['mean_ecfp_tanimoto']:.4f}±{row['std_ecfp_tanimoto']:.4f}, "
            f"RDK={row['mean_rdk_tanimoto']:.4f}±{row['std_rdk_tanimoto']:.4f}"
        )


if __name__ == "__main__":
    main()
```
