## Supplementary material for "PG-MLD: Physics-Guided Molecular Representation Learning via Dynamic 3D Trajectory Distillation": Code and Data: representation_space.py

```
from __future__ import annotations

import argparse
import csv
import sys
from pathlib import Path
from typing import Any

PROJECT_ROOT = Path(__file__).resolve().parents[1]
if str(PROJECT_ROOT) not in sys.path:
    sys.path.insert(0, str(PROJECT_ROOT))

import matplotlib.pyplot as plt
import numpy as np
import pandas as pd
import torch
from tqdm import tqdm

from comld.models.chemblm_distill import ChemBLMEncoder


def torch_load(path: str | Path) -> dict[str, Any]:
    path = Path(path)
    if not path.exists():
        raise FileNotFoundError(f"Checkpoint not found: {path}")
    try:
        return torch.load(path, map_location="cpu", weights_only=False)
    except TypeError:
        return torch.load(path, map_location="cpu")


def resolve_device(name: str) -> torch.device:
    if name == "auto":
        return torch.device("cuda" if torch.cuda.is_available() else "cpu")
    return torch.device(name)


def build_chemblm_encoder(
    model_name: str,
    checkpoint: str | None,
    trust_remote_code: bool,
    use_safetensors: bool,
) -> ChemBLMEncoder:
    """Build ChemBLM encoder and optionally load distilled or finetuned weights."""

    ckpt = torch_load(checkpoint) if checkpoint else None
    if ckpt is not None and ckpt.get("student_model_name"):
        model_name = ckpt.get("student_model_name", model_name)

    encoder = ChemBLMEncoder(
        model_name=model_name,
        trust_remote_code=trust_remote_code,
        use_safetensors=use_safetensors,
    )

    if ckpt is None:
        print(f"Loaded original ChemBLM encoder: {model_name}")
        return encoder

    if "student_state_dict" in ckpt:
        state = ckpt["student_state_dict"]
        missing, unexpected = encoder.model.load_state_dict(state, strict=False)
        print(f"Loaded distilled ChemBLM checkpoint: {checkpoint}")
    elif "model_state_dict" in ckpt:
        model_state = ckpt["model_state_dict"]
        state = {
            key.removeprefix("encoder.model."): value
            for key, value in model_state.items()
            if key.startswith("encoder.model.")
        }
        if not state:
            raise KeyError(
                "Finetuned checkpoint has no encoder.model.* keys. "
                "Please pass a ChemBLM finetuned best.pt or a 3D-to-1D distilled checkpoint."
            )
        missing, unexpected = encoder.model.load_state_dict(state, strict=False)
        print(f"Loaded encoder weights from finetuned checkpoint: {checkpoint}")
    else:
        raise KeyError(
            "Unsupported checkpoint format. Expected 'student_state_dict' for pretraining "
            "or 'model_state_dict' for finetuned best.pt."
        )

    if missing:
        print(f"Missing encoder keys: {missing}")
    if unexpected:
        print(f"Unexpected encoder keys: {unexpected}")
    return encoder


def valid_smiles_and_descriptors(smiles: str) -> dict[str, Any] | None:
    try:
        from rdkit import Chem
        from rdkit.Chem import Crippen, Descriptors, Lipinski, rdMolDescriptors
    except ImportError as exc:
        raise ImportError("RDKit is required for descriptor calculation.") from exc

    mol = Chem.MolFromSmiles(str(smiles))
    if mol is None:
        return None

    return {
        "canonical_smiles": Chem.MolToSmiles(mol),
        "mol_weight": float(Descriptors.MolWt(mol)),
        "exact_mol_weight": float(Descriptors.ExactMolWt(mol)),
        "logp": float(Crippen.MolLogP(mol)),
        "tpsa": float(rdMolDescriptors.CalcTPSA(mol)),
        "hbd": int(Lipinski.NumHDonors(mol)),
        "hba": int(Lipinski.NumHAcceptors(mol)),
        "rotatable_bonds": int(Lipinski.NumRotatableBonds(mol)),
        "ring_count": int(rdMolDescriptors.CalcNumRings(mol)),
        "aromatic_ring_count": int(rdMolDescriptors.CalcNumAromaticRings(mol)),
        "heavy_atom_count": int(mol.GetNumHeavyAtoms()),
    }


@torch.no_grad()
def extract_embeddings(
    encoder: ChemBLMEncoder,
    smiles: list[str],
    batch_size: int,
    device: torch.device,
) -> np.ndarray:
    encoder.to(device)
    encoder.eval()
    batches: list[np.ndarray] = []
    for start in tqdm(range(0, len(smiles), batch_size), desc="Extract ChemBLM embeddings"):
        batch = smiles[start : start + batch_size]
        emb = encoder(batch, device=device)
        batches.append(emb.detach().cpu().float().numpy())
    return np.concatenate(batches, axis=0).astype(np.float32)


def command_extract(args: argparse.Namespace) -> None:
    np.random.seed(args.seed)
    device = resolve_device(args.device)

    frame = pd.read_csv(args.csv)
    if args.smiles_col not in frame.columns:
        raise ValueError(f"Column {args.smiles_col!r} not found in {args.csv}. Columns: {list(frame.columns)}")

    if args.start_index:
        frame = frame.iloc[int(args.start_index) :]
    if args.max_molecules:
        frame = frame.head(int(args.max_molecules))

    smiles_raw = frame[args.smiles_col].astype(str).tolist()
    rows: list[dict[str, Any]] = []
    smiles_valid: list[str] = []
    for source_index, smiles in tqdm(list(enumerate(smiles_raw)), desc="RDKit descriptors"):
        desc = valid_smiles_and_descriptors(smiles)
        if desc is None:
            continue
        desc["source_index"] = int(source_index + int(args.start_index or 0))
        desc["input_smiles"] = smiles
        rows.append(desc)
        smiles_valid.append(desc["canonical_smiles"])

    if not smiles_valid:
        raise RuntimeError("No valid molecules found.")

    encoder = build_chemblm_encoder(
        model_name=args.model_name,
        checkpoint=args.checkpoint,
        trust_remote_code=bool(args.trust_remote_code),
        use_safetensors=bool(args.use_safetensors),
    )
    embeddings = extract_embeddings(encoder, smiles_valid, batch_size=args.batch_size, device=device)

    out_prefix = Path(args.out_prefix)
    out_prefix.parent.mkdir(parents=True, exist_ok=True)
    meta_path = out_prefix.with_suffix(".csv")
    npz_path = out_prefix.with_suffix(".npz")

    meta = pd.DataFrame(rows)
    meta.to_csv(meta_path, index=False)
    np.savez_compressed(
        npz_path,
        embeddings=embeddings,
        smiles=np.asarray(smiles_valid),
        mol_weight=meta["mol_weight"].to_numpy(np.float32),
        logp=meta["logp"].to_numpy(np.float32),
        tpsa=meta["tpsa"].to_numpy(np.float32),
        ring_count=meta["ring_count"].to_numpy(np.float32),
        aromatic_ring_count=meta["aromatic_ring_count"].to_numpy(np.float32),
        heavy_atom_count=meta["heavy_atom_count"].to_numpy(np.float32),
    )

    print(f"Saved metadata: {meta_path}")
    print(f"Saved embeddings: {npz_path}")
    print(f"Molecules: {len(smiles_valid)}")
    print(f"Embedding shape: {embeddings.shape}")


def standardize(x: np.ndarray) -> np.ndarray:
    x = x.astype(np.float32, copy=False)
    return (x - x.mean(axis=0, keepdims=True)) / np.maximum(x.std(axis=0, keepdims=True), 1e-6)


def pca_reduce(x: np.ndarray, n_components: int = 50) -> np.ndarray:
    x_t = torch.as_tensor(standardize(x), dtype=torch.float32)
    x_t = x_t - x_t.mean(dim=0, keepdim=True)
    n_components = min(int(n_components), x_t.shape[0] - 1, x_t.shape[1])
    try:
        _, _, v = torch.pca_lowrank(x_t, q=n_components, center=False)
        z = x_t @ v[:, :n_components]
    except Exception:
        _, _, vh = torch.linalg.svd(x_t, full_matrices=False)
        z = x_t @ vh[:n_components].T
    return z.cpu().numpy()


def pca_axes_correlated_with_values(
    x: np.ndarray,
    values: np.ndarray,
    n_components: int = 30,
) -> tuple[np.ndarray, list[int], list[float]]:
    """Compute many PCA axes, then plot the two most correlated with descriptor values.

    This keeps the projection PCA-based while avoiding the common failure mode
    where PC1/PC2 explain high variance unrelated to the molecular descriptor
    used for coloring.
    """

    pcs = pca_reduce(x, n_components=n_components)
    values = np.asarray(values, dtype=float)
    correlations: list[float] = []
    for axis in range(pcs.shape[1]):
        pc = pcs[:, axis]
        if np.std(pc) < 1e-12 or np.std(values) < 1e-12:
            correlations.append(0.0)
        else:
            correlations.append(float(np.corrcoef(pc, values)[0, 1]))

    order = np.argsort(np.abs(correlations))[::-1]
    if len(order) == 0:
        return pca_reduce(x, n_components=2), [1, 2], [0.0, 0.0]
    if len(order) == 1:
        z = np.column_stack([pcs[:, order[0]], np.zeros(len(pcs), dtype=pcs.dtype)])
        return z, [int(order[0]) + 1, 0], [float(correlations[order[0]]), 0.0]

    chosen = [int(order[0]), int(order[1])]
    z = pcs[:, chosen]
    chosen_1based = [axis + 1 for axis in chosen]
    chosen_corrs = [float(correlations[axis]) for axis in chosen]
    return z, chosen_1based, chosen_corrs


def torch_tsne(
    x: np.ndarray,
    seed: int,
    perplexity: float,
    n_iter: int,
    learning_rate: float = 200.0,
    pca_dim: int = 50,
) -> np.ndarray:
    rng = np.random.default_rng(seed)
    x = pca_reduce(x, n_components=min(pca_dim, x.shape[1]))
    x = standardize(x)
    n = x.shape[0]
    if n <= 2:
        return rng.normal(size=(n, 2)).astype(np.float32)
    if n > 8000:
        raise RuntimeError(
            "Torch fallback t-SNE is O(N^2). Use --max-points <= 8000, --backend pca, "
            "or repair sklearn/scipy and use --backend sklearn for larger data."
        )

    perplexity = float(min(perplexity, max(5.0, (n - 1) / 3.0)))
    device = torch.device("cuda" if torch.cuda.is_available() else "cpu")
    x_t = torch.as_tensor(x, dtype=torch.float32, device=device)
    d = torch.cdist(x_t, x_t).pow(2)
    d.fill_diagonal_(float("inf"))

    target_h = float(np.log(perplexity))
    beta = torch.ones(n, device=device)
    beta_min = torch.full((n,), -float("inf"), device=device)
    beta_max = torch.full((n,), float("inf"), device=device)
    finite_d = torch.where(torch.isfinite(d), d, torch.zeros_like(d))

    for _ in range(50):
        p_cond = torch.exp(-d * beta[:, None])
        p_cond = torch.where(torch.isfinite(d), p_cond, torch.zeros_like(p_cond))
        sum_p = p_cond.sum(dim=1).clamp_min(1e-12)
        h = torch.log(sum_p) + beta * (finite_d * p_cond).sum(dim=1) / sum_p
        high = h > target_h
        beta_min = torch.where(high, beta, beta_min)
        beta_max = torch.where(~high, beta, beta_max)
        beta = torch.where(
            high,
            torch.where(torch.isinf(beta_max), beta * 2.0, (beta + beta_max) / 2.0),
            torch.where(torch.isinf(beta_min), beta / 2.0, (beta + beta_min) / 2.0),
        )

    p_cond = torch.exp(-d * beta[:, None])
    p_cond = torch.where(torch.isfinite(d), p_cond, torch.zeros_like(p_cond))
    p_cond = p_cond / p_cond.sum(dim=1, keepdim=True).clamp_min(1e-12)
    p = ((p_cond + p_cond.T) / (2.0 * n)).clamp_min(1e-12)

    y = torch.as_tensor(rng.normal(0.0, 1e-4, size=(n, 2)), dtype=torch.float32, device=device)
    update = torch.zeros_like(y)
    gains = torch.ones_like(y)
    last_grad = torch.zeros_like(y)

    for step in tqdm(range(int(n_iter)), desc="t-SNE"):
        exaggeration = 12.0 if step < 250 else 1.0
        yy = torch.cdist(y, y).pow(2)
        num = 1.0 / (1.0 + yy)
        num.fill_diagonal_(0.0)
        q = (num / num.sum().clamp_min(1e-12)).clamp_min(1e-12)
        w = (p * exaggeration - q) * num
        grad = 4.0 * (y * w.sum(dim=1, keepdim=True) - w @ y)
        gains = torch.where(torch.sign(grad) != torch.sign(last_grad), gains + 0.2, gains * 0.8).clamp_min(0.01)
        momentum = 0.5 if step < 250 else 0.8
        update = momentum * update - learning_rate * gains * grad
        y = y + update
        y = y - y.mean(dim=0, keepdim=True)
        last_grad = grad
    return y.detach().cpu().numpy()


def make_2d(
    x: np.ndarray,
    backend: str,
    seed: int,
    perplexity: float,
    n_iter: int,
    values: np.ndarray | None = None,
    pca_axis_mode: str = "first",
    pca_components: int = 30,
) -> tuple[np.ndarray, str, str, dict[str, Any]]:
    if backend in {"auto", "sklearn"}:
        try:
            from sklearn.manifold import TSNE

            kwargs = dict(n_components=2, perplexity=perplexity, learning_rate="auto", init="pca", random_state=seed)
            try:
                z = TSNE(**kwargs, max_iter=n_iter).fit_transform(standardize(x))
            except TypeError:
                z = TSNE(**kwargs, n_iter=n_iter).fit_transform(standardize(x))
            return z, "t-SNE", "tsne", {}
        except Exception as exc:
            if backend == "sklearn":
                raise
            print(f"sklearn t-SNE unavailable, fallback to torch t-SNE: {exc}")
    if backend == "pca":
        if pca_axis_mode == "correlated":
            if values is None:
                raise ValueError("--pca-axis-mode correlated requires color values.")
            z, axes, corrs = pca_axes_correlated_with_values(x, values, n_components=pca_components)
            method_name = f"PCA selected axes PC{axes[0]}/PC{axes[1]}"
            return z, method_name, f"pca_corr_pc{axes[0]}_pc{axes[1]}", {
                "selected_pcs": axes,
                "selected_pc_correlations": corrs,
            }
        return pca_reduce(x, n_components=2), "PCA", "pca", {}
    return torch_tsne(x, seed=seed, perplexity=perplexity, n_iter=n_iter), "t-SNE", "tsne", {}


def command_tsne(args: argparse.Namespace) -> None:
    data = np.load(args.npz, allow_pickle=True)
    embeddings = data["embeddings"]
    meta = pd.read_csv(args.csv)

    if args.color_by not in meta.columns:
        raise ValueError(f"color_by={args.color_by!r} not found in {args.csv}. Columns: {list(meta.columns)}")

    all_values = meta[args.color_by].to_numpy(float)
    valid_value_mask = np.isfinite(all_values)
    indices = np.arange(len(meta))[valid_value_mask]

    if args.max_points and len(indices) > int(args.max_points):
        rng = np.random.default_rng(args.seed)
        if int(args.stratify_color_bins) > 1:
            # For descriptor-colored unsupervised maps, pure random sampling can
            # under-sample the low/high tails and make the color trend invisible.
            # Quantile-stratified sampling keeps each descriptor range visible.
            values_for_bins = all_values[indices]
            quantiles = np.linspace(0.0, 1.0, int(args.stratify_color_bins) + 1)
            edges = np.quantile(values_for_bins, quantiles)
            edges = np.unique(edges)
            if len(edges) > 2:
                bin_ids = np.digitize(values_for_bins, edges[1:-1], right=True)
                chosen: list[int] = []
                per_bin = max(1, int(args.max_points) // int(args.stratify_color_bins))
                for bin_id in range(len(edges) - 1):
                    in_bin = indices[bin_ids == bin_id]
                    if len(in_bin) <= per_bin:
                        chosen.extend(in_bin.tolist())
                    else:
                        chosen.extend(rng.choice(in_bin, size=per_bin, replace=False).tolist())
                if len(chosen) < int(args.max_points):
                    remaining = np.setdiff1d(indices, np.asarray(chosen, dtype=int), assume_unique=False)
                    add_n = min(len(remaining), int(args.max_points) - len(chosen))
                    if add_n > 0:
                        chosen.extend(rng.choice(remaining, size=add_n, replace=False).tolist())
                indices = np.asarray(chosen[: int(args.max_points)], dtype=int)
                rng.shuffle(indices)
            else:
                indices = rng.choice(indices, size=int(args.max_points), replace=False)
        else:
            indices = rng.choice(indices, size=int(args.max_points), replace=False)

    x = embeddings[indices]
    values = meta.iloc[indices][args.color_by].to_numpy(float)
    z, method_name, file_prefix, method_info = make_2d(
        x,
        backend=args.backend,
        seed=args.seed,
        perplexity=args.perplexity,
        n_iter=args.n_iter,
        values=values,
        pca_axis_mode=args.pca_axis_mode,
        pca_components=args.pca_components,
    )

    vmin = vmax = None
    if args.clip_percentiles:
        low, high = [float(v) for v in args.clip_percentiles]
        vmin, vmax = np.percentile(values, [low, high])

    if args.sort_by_color:
        order = np.argsort(values)
        z = z[order]
        x = x[order]
        values = values[order]
        indices = indices[order]

    out_dir = Path(args.out_dir)
    out_dir.mkdir(parents=True, exist_ok=True)
    png_path = out_dir / f"{file_prefix}_{args.color_by}.png"
    npz_path = out_dir / f"{file_prefix}_{args.color_by}.npz"

    fig, ax = plt.subplots(figsize=(8.5, 6.6), dpi=300)
    scatter = ax.scatter(
        z[:, 0],
        z[:, 1],
        c=values,
        cmap=args.cmap,
        vmin=vmin,
        vmax=vmax,
        s=args.point_size,
        alpha=args.alpha,
        edgecolors="none",
    )
    cbar = fig.colorbar(scatter, ax=ax, pad=0.01)
    cbar.set_label(args.color_by)
    ax.set_title(args.title or f"ChemBLM molecular representation {method_name} colored by {args.color_by}", fontsize=15)
    ax.set_xlabel(f"{method_name}1")
    ax.set_ylabel(f"{method_name}2")
    ax.set_xticks([])
    ax.set_yticks([])
    fig.tight_layout()
    fig.savefig(png_path, dpi=600, bbox_inches="tight")
    plt.close(fig)

    np.savez_compressed(
        npz_path,
        z=z,
        values=values,
        indices=indices,
        color_by=np.array(args.color_by),
        method=np.array(method_name),
        method_info=np.array(str(method_info)),
    )
    print(f"Saved figure: {png_path}")
    print(f"Saved {method_name} data: {npz_path}")
    print(f"Plotted molecules: {len(indices)}")
    if len(indices) > 2:
        corr1 = float(np.corrcoef(z[:, 0], values)[0, 1])
        corr2 = float(np.corrcoef(z[:, 1], values)[0, 1])
        print(f"{args.color_by} range: min={np.min(values):.3f} median={np.median(values):.3f} max={np.max(values):.3f}")
        print(f"Correlation with {method_name} axes: axis1={corr1:.4f}, axis2={corr2:.4f}")
        if method_info:
            print(f"Method info: {method_info}")


def fingerprint_similarities(query_smiles: str, smiles_list: list[str]) -> tuple[list[float], list[float]]:
    from rdkit import Chem, DataStructs
    from rdkit.Chem import AllChem

    qmol = Chem.MolFromSmiles(query_smiles)
    if qmol is None:
        raise ValueError(f"Invalid query smiles: {query_smiles}")
    q_ecfp = AllChem.GetMorganFingerprintAsBitVect(qmol, radius=2, nBits=2048)
    q_rdk = Chem.RDKFingerprint(qmol)

    ecfp_sims: list[float] = []
    rdk_sims: list[float] = []
    for smiles in smiles_list:
        mol = Chem.MolFromSmiles(smiles)
        if mol is None:
            ecfp_sims.append(float("nan"))
            rdk_sims.append(float("nan"))
            continue
        ecfp = AllChem.GetMorganFingerprintAsBitVect(mol, radius=2, nBits=2048)
        rdk = Chem.RDKFingerprint(mol)
        ecfp_sims.append(float(DataStructs.TanimotoSimilarity(q_ecfp, ecfp)))
        rdk_sims.append(float(DataStructs.TanimotoSimilarity(q_rdk, rdk)))
    return ecfp_sims, rdk_sims


def command_neighbors(args: argparse.Namespace) -> None:
    from rdkit import Chem
    from rdkit.Chem import Draw

    data = np.load(args.npz, allow_pickle=True)
    embeddings = data["embeddings"].astype(np.float32)
    meta = pd.read_csv(args.csv)

    smiles = meta["canonical_smiles"].astype(str).tolist()
    x = embeddings / np.maximum(np.linalg.norm(embeddings, axis=1, keepdims=True), 1e-12)

    out_dir = Path(args.out_dir)
    out_dir.mkdir(parents=True, exist_ok=True)
    query_indices = [int(v.strip()) for v in str(args.query_indices).split(",") if v.strip()]

    for qidx in query_indices:
        if qidx < 0 or qidx >= len(smiles):
            raise IndexError(f"query index out of range: {qidx}")

        sims = x @ x[qidx]
        order = np.argsort(-sims)
        order = [int(i) for i in order if int(i) != qidx][: int(args.top_k)]
        selected = [qidx] + order
        selected_smiles = [smiles[i] for i in selected]
        ecfp_sims, rdk_sims = fingerprint_similarities(smiles[qidx], selected_smiles)

        mols = [Chem.MolFromSmiles(smi) for smi in selected_smiles]
        legends = ["Query"]
        for rank, idx in enumerate(order, start=1):
            legends.append(
                f"#{rank} cos={sims[idx]:.3f}\n"
                f"ECFP={ecfp_sims[rank]:.3f} RDK={rdk_sims[rank]:.3f}"
            )

        image = Draw.MolsToGridImage(
            mols,
            molsPerRow=args.mols_per_row,
            subImgSize=(args.sub_img_size, args.sub_img_size),
            legends=legends,
            useSVG=False,
        )
        fig_path = out_dir / f"nearest_query_{qidx}.png"
        image.save(fig_path)

        csv_path = out_dir / f"nearest_query_{qidx}.csv"
        with open(csv_path, "w", newline="", encoding="utf-8") as handle:
            writer = csv.writer(handle)
            writer.writerow(["rank", "index", "smiles", "cosine", "ecfp_tanimoto", "rdk_tanimoto"])
            writer.writerow([0, qidx, smiles[qidx], 1.0, 1.0, 1.0])
            for rank, idx in enumerate(order, start=1):
                writer.writerow([rank, idx, smiles[idx], float(sims[idx]), ecfp_sims[rank], rdk_sims[rank]])

        print(f"Saved nearest-neighbor figure: {fig_path}")
        print(f"Saved nearest-neighbor table: {csv_path}")


def build_parser() -> argparse.ArgumentParser:
    parser = argparse.ArgumentParser(description="ChemBLM/PG-MLD representation-space visualization experiment.")
    sub = parser.add_subparsers(dest="command", required=True)

    p_extract = sub.add_parser("extract", help="Extract ChemBLM embeddings and RDKit descriptors.")
    p_extract.add_argument("--csv", required=True)
    p_extract.add_argument("--smiles-col", default="smiles")
    p_extract.add_argument("--model-name", default="comld/models/ChemBLM")
    p_extract.add_argument("--checkpoint", default="checkpoints/pretrain_zinc_dynamic_3d_to_1d_chemblm/latest.pt")
    p_extract.add_argument("--out-prefix", required=True)
    p_extract.add_argument("--start-index", type=int, default=0)
    p_extract.add_argument("--max-molecules", type=int, default=10000)
    p_extract.add_argument("--batch-size", type=int, default=64)
    p_extract.add_argument("--device", default="auto")
    p_extract.add_argument("--seed", type=int, default=2026)
    p_extract.add_argument("--trust-remote-code", action="store_true")
    p_extract.add_argument("--use-safetensors", action="store_true")
    p_extract.set_defaults(func=command_extract)

    p_tsne = sub.add_parser("tsne", help="Draw t-SNE/PCA colored by molecular descriptors.")
    p_tsne.add_argument("--npz", required=True)
    p_tsne.add_argument("--csv", required=True)
    p_tsne.add_argument("--color-by", default="mol_weight")
    p_tsne.add_argument("--out-dir", required=True)
    p_tsne.add_argument("--backend", default="auto", choices=["auto", "sklearn", "torch", "pca"])
    p_tsne.add_argument("--max-points", type=int, default=5000)
    p_tsne.add_argument("--perplexity", type=float, default=30.0)
    p_tsne.add_argument("--n-iter", type=int, default=1200)
    p_tsne.add_argument("--seed", type=int, default=2026)
    p_tsne.add_argument("--cmap", default="Spectral")
    p_tsne.add_argument("--point-size", type=float, default=4.0)
    p_tsne.add_argument("--alpha", type=float, default=0.78)
    p_tsne.add_argument("--title", default=None)
    p_tsne.add_argument("--stratify-color-bins", type=int, default=0)
    p_tsne.add_argument("--clip-percentiles", type=float, nargs=2, default=None)
    p_tsne.add_argument("--sort-by-color", action="store_true")
    p_tsne.add_argument("--pca-axis-mode", choices=["first", "correlated"], default="first")
    p_tsne.add_argument("--pca-components", type=int, default=30)
    p_tsne.set_defaults(func=command_tsne)

    p_nn = sub.add_parser("neighbors", help="Find nearest molecules in representation space and draw structures.")
    p_nn.add_argument("--npz", required=True)
    p_nn.add_argument("--csv", required=True)
    p_nn.add_argument("--query-indices", required=True, help="Comma-separated metadata row indices, e.g. 0,100,200.")
    p_nn.add_argument("--top-k", type=int, default=9)
    p_nn.add_argument("--out-dir", required=True)
    p_nn.add_argument("--mols-per-row", type=int, default=5)
    p_nn.add_argument("--sub-img-size", type=int, default=260)
    p_nn.set_defaults(func=command_neighbors)
    return parser


def main() -> None:
    parser = build_parser()
    args = parser.parse_args()
    args.func(args)


if __name__ == "__main__":
    main()
```
