## Supplementary material for "PG-MLD: Physics-Guided Molecular Representation Learning via Dynamic 3D Trajectory Distillation": Code and Data: shap_chemblm.py

```
from __future__ import annotations

import argparse
import csv
import sys
from collections import defaultdict
from pathlib import Path
from typing import Any, Sequence

PROJECT_ROOT = Path(__file__).resolve().parents[1]
if str(PROJECT_ROOT) not in sys.path:
    sys.path.insert(0, str(PROJECT_ROOT))

import matplotlib as mpl
import matplotlib.pyplot as plt
import numpy as np
import torch
from tqdm import tqdm

from comld.models.chemberta_teacher import find_smiles_atom_spans, mean_pool_tokens
from comld.models.molformer_distill import attention_mean_pool
from comld.train.finetune_1d import (
    SmilesTaskDataset,
    load_checkpoint,
    load_config,
    resolve_device,
    set_seed,
    split_dataset_indices,
)
from comld.train.finetune_chemblm_1d import build_model as build_chemblm_model


POSITIVE_COLOR = "#E37019"
NEGATIVE_COLOR = "#0A48AB"
ORIGINAL_COLOR = "#7C8793"
DISTILLED_COLOR = "#3E7454"

# Compact SCI style shared with visualization/t-sne.py.  These dimensions
# describe the central bar-chart axes, not the complete exported canvas.
CM_TO_INCH = 1.0 / 2.54
TOKEN_PANEL_WIDTH_IN = 4.24 * CM_TO_INCH
TOKEN_PANEL_HEIGHT_IN = 5.71 * CM_TO_INCH
PLOT_FONT_FAMILY = "Arial"
AXIS_LINEWIDTH = 0.45
GRID_LINEWIDTH = 0.25
TICK_LINEWIDTH = 0.40


def configure_matplotlib() -> None:
    mpl.rcParams.update(
        {
            "font.family": "sans-serif",
            "font.sans-serif": ["Arial", "Helvetica", "DejaVu Sans"],
            "axes.unicode_minus": False,
            "pdf.fonttype": 42,
            "ps.fonttype": 42,
            "svg.fonttype": "none",
            "figure.facecolor": "white",
            "axes.facecolor": "white",
        }
    )


def torch_load(path: str | Path) -> dict[str, Any]:
    path = Path(path)
    if not path.is_file():
        raise FileNotFoundError(f"Checkpoint not found: {path}")
    try:
        return torch.load(path, map_location="cpu", weights_only=False)
    except TypeError:
        return torch.load(path, map_location="cpu")


def load_best_model(
    config: dict[str, Any],
    checkpoint_path: str | Path,
    output_dim: int,
    device: torch.device,
) -> torch.nn.Module:
    pretrained = load_checkpoint(config.get("paths", {}).get("pretrained_checkpoint"))
    model = build_chemblm_model(config, checkpoint=pretrained, output_dim=output_dim)
    checkpoint = torch_load(checkpoint_path)
    state = checkpoint.get("model_state_dict")
    if state is None:
        raise KeyError(f"{checkpoint_path} is not a finetuned ChemBLM best.pt checkpoint.")
    missing, unexpected = model.load_state_dict(state, strict=False)
    if missing:
        print(f"Missing checkpoint keys: {missing}")
    if unexpected:
        print(f"Unexpected checkpoint keys: {unexpected}")
    model.to(device).eval()
    print(f"Loaded checkpoint: {checkpoint_path} (epoch={checkpoint.get('epoch', 'unknown')})")
    return model


def repeat_inputs(inputs: dict[str, torch.Tensor], count: int) -> dict[str, torch.Tensor]:
    return {key: value.repeat(count, *([1] * (value.ndim - 1))) for key, value in inputs.items()}


@torch.no_grad()
def probabilities_from_ids(
    model: torch.nn.Module,
    model_inputs: dict[str, torch.Tensor],
    offsets: torch.Tensor | None,
    task_index: int,
) -> torch.Tensor:
    output = model.encoder.model(**model_inputs)
    hidden = output.last_hidden_state if hasattr(output, "last_hidden_state") else output[0]
    attention_mask = model_inputs["attention_mask"]
    if offsets is not None:
        pooled = mean_pool_tokens(hidden, attention_mask, offsets)
    else:
        pooled = attention_mean_pool(hidden, attention_mask)
    logits = model.head(pooled)
    return torch.sigmoid(logits[:, task_index])


def tokenize_for_shap(
    model: torch.nn.Module,
    smiles: str,
    device: torch.device,
) -> tuple[dict[str, torch.Tensor], torch.Tensor | None, list[int], list[str], list[tuple[int, int]]]:
    tokenizer = model.encoder.tokenizer
    kwargs: dict[str, Any] = {
        "truncation": True,
        "return_tensors": "pt",
        "return_special_tokens_mask": True,
    }
    if getattr(tokenizer, "is_fast", False):
        kwargs["return_offsets_mapping"] = True
    encoded = tokenizer(smiles, **kwargs)
    offsets = encoded.pop("offset_mapping", None)
    special_mask = encoded.pop("special_tokens_mask")[0].bool()
    attention = encoded["attention_mask"][0].bool()
    input_ids_cpu = encoded["input_ids"][0]
    positions = torch.where(attention & ~special_mask)[0].tolist()
    if not positions:
        raise RuntimeError(f"Tokenizer produced no explainable tokens for SMILES: {smiles}")

    token_strings = tokenizer.convert_ids_to_tokens(input_ids_cpu[positions].tolist())
    spans: list[tuple[int, int]] = []
    if offsets is not None:
        offsets_cpu = offsets[0]
        spans = [(int(offsets_cpu[pos, 0]), int(offsets_cpu[pos, 1])) for pos in positions]
        display = []
        for token, (start, end) in zip(token_strings, spans):
            fragment = smiles[start:end]
            display.append(fragment if fragment else token)
        token_strings = display
    else:
        spans = [(-1, -1) for _ in positions]

    model_inputs = {key: value.to(device) for key, value in encoded.items()}
    if offsets is not None:
        offsets = offsets.to(device)
    return model_inputs, offsets, positions, token_strings, spans


@torch.no_grad()
def permutation_shap(
    model: torch.nn.Module,
    smiles: str,
    task_index: int,
    permutations: int,
    inference_batch_size: int,
    seed: int,
    device: torch.device,
) -> dict[str, Any]:
    model_inputs, offsets, positions, tokens, spans = tokenize_for_shap(model, smiles, device)
    tokenizer = model.encoder.tokenizer
    mask_token_id = tokenizer.mask_token_id
    if mask_token_id is None:
        mask_token_id = tokenizer.unk_token_id
    if mask_token_id is None:
        raise ValueError("ChemBLM tokenizer has neither mask_token_id nor unk_token_id.")

    original_ids = model_inputs["input_ids"][0].clone()
    baseline_ids = original_ids.clone()
    baseline_ids[positions] = int(mask_token_id)
    n_features = len(positions)
    rng = np.random.default_rng(seed)

    coalitions: list[np.ndarray] = []
    orders: list[np.ndarray] = []
    for _ in range(max(1, int(permutations))):
        order = rng.permutation(n_features)
        orders.append(order)
        state = np.zeros(n_features, dtype=bool)
        coalitions.append(state.copy())
        for feature_index in order:
            state[feature_index] = True
            coalitions.append(state.copy())
    coalition_matrix = np.stack(coalitions, axis=0)

    values: list[torch.Tensor] = []
    for start in range(0, len(coalition_matrix), max(1, int(inference_batch_size))):
        batch_coalitions = coalition_matrix[start : start + int(inference_batch_size)]
        count = len(batch_coalitions)
        batch_inputs = repeat_inputs(model_inputs, count)
        batch_ids = baseline_ids.unsqueeze(0).repeat(count, 1)
        for row, included in enumerate(batch_coalitions):
            active = [positions[idx] for idx in np.flatnonzero(included)]
            if active:
                batch_ids[row, active] = original_ids[active]
        batch_inputs["input_ids"] = batch_ids
        batch_offsets = offsets.repeat(count, 1, 1) if offsets is not None else None
        values.append(probabilities_from_ids(model, batch_inputs, batch_offsets, task_index).cpu())
    coalition_values = torch.cat(values).numpy()

    shap_values = np.zeros(n_features, dtype=np.float64)
    stride = n_features + 1
    for permutation_index, order in enumerate(orders):
        sequence_values = coalition_values[permutation_index * stride : (permutation_index + 1) * stride]
        for step, feature_index in enumerate(order):
            shap_values[feature_index] += float(sequence_values[step + 1] - sequence_values[step])
    shap_values /= max(1, len(orders))

    baseline_probability = float(np.mean(coalition_values[::stride]))
    full_probability = float(np.mean(coalition_values[stride - 1 :: stride]))
    reconstruction = baseline_probability + float(shap_values.sum())
    return {
        "smiles": smiles,
        "tokens": tokens,
        "spans": spans,
        "shap_values": shap_values,
        "baseline_probability": baseline_probability,
        "full_probability": full_probability,
        "reconstruction_error": abs(full_probability - reconstruction),
    }


def select_samples(
    dataset: SmilesTaskDataset,
    indices: Sequence[int],
    task_index: int,
    num_samples: int,
    seed: int,
) -> list[int]:
    valid = [idx for idx in indices if float(dataset.masks[idx][task_index]) > 0.5]
    if not valid:
        raise RuntimeError("No valid labels are available for the selected task and subset.")
    rng = np.random.default_rng(seed)
    by_class = {
        cls: [idx for idx in valid if int(round(float(dataset.labels[idx][task_index]))) == cls]
        for cls in (0, 1)
    }
    target = min(max(1, int(num_samples)), len(valid))
    selected: list[int] = []
    per_class = target // 2
    for cls in (0, 1):
        pool = np.asarray(by_class[cls], dtype=int)
        take = min(per_class, len(pool))
        if take:
            selected.extend(rng.choice(pool, size=take, replace=False).tolist())
    remaining = [idx for idx in valid if idx not in set(selected)]
    if len(selected) < target and remaining:
        selected.extend(
            rng.choice(np.asarray(remaining), size=min(target - len(selected), len(remaining)), replace=False).tolist()
        )
    return selected


def save_csv(path: Path, rows: list[dict[str, Any]]) -> None:
    path.parent.mkdir(parents=True, exist_ok=True)
    if not rows:
        return
    with path.open("w", newline="", encoding="utf-8") as handle:
        writer = csv.DictWriter(handle, fieldnames=list(rows[0]))
        writer.writeheader()
        writer.writerows(rows)


def save_figure(fig: plt.Figure, base: Path) -> list[Path]:
    base.parent.mkdir(parents=True, exist_ok=True)
    paths = [base.with_suffix(suffix) for suffix in (".png", ".pdf", ".svg")]
    fig.savefig(paths[0], dpi=600, bbox_inches="tight", pad_inches=0.04, facecolor="white")
    fig.savefig(paths[1], bbox_inches="tight", pad_inches=0.04, facecolor="white")
    fig.savefig(paths[2], bbox_inches="tight", pad_inches=0.04, facecolor="white")
    plt.close(fig)
    return paths


def normalize_shap_values(values: np.ndarray) -> np.ndarray:
    values = np.asarray(values, dtype=np.float64)
    denominator = float(np.abs(values).sum())
    if denominator <= 1e-12:
        return np.zeros_like(values)
    return values / denominator


def token_shap_to_atoms(
    smiles: str,
    token_spans: Sequence[tuple[int, int]],
    token_values: np.ndarray,
) -> tuple[Any, np.ndarray]:
    """Distribute signed token SHAP values to overlapping or nearest atoms."""
    from rdkit import Chem
    from rdkit.Chem import rdDepictor

    mol = Chem.MolFromSmiles(smiles)
    if mol is None:
        raise ValueError(f"RDKit could not parse SMILES: {smiles}")
    if not any(start >= 0 and end > start for start, end in token_spans):
        raise RuntimeError(
            "The tokenizer did not provide character offsets, so token SHAP values "
            "cannot be mapped to atoms reliably. Use a fast tokenizer."
        )
    rdDepictor.Compute2DCoords(mol)
    atom_spans = find_smiles_atom_spans(smiles)
    if len(atom_spans) != mol.GetNumAtoms():
        raise RuntimeError(
            f"SMILES atom-span count ({len(atom_spans)}) does not match RDKit atom count "
            f"({mol.GetNumAtoms()}) for {smiles}"
        )

    weights = np.zeros(mol.GetNumAtoms(), dtype=np.float64)
    for (token_start, token_end), value in zip(token_spans, token_values):
        if token_start < 0 or token_end <= token_start:
            continue
        recipients = [
            atom_index
            for atom_index, (atom_start, atom_end) in enumerate(atom_spans)
            if atom_start < token_end and atom_end > token_start
        ]
        if not recipients:
            left = [
                (token_start - atom_end, atom_index)
                for atom_index, (_, atom_end) in enumerate(atom_spans)
                if atom_end <= token_start
            ]
            right = [
                (atom_start - token_end, atom_index)
                for atom_index, (atom_start, _) in enumerate(atom_spans)
                if atom_start >= token_end
            ]
            if left:
                recipients.append(min(left)[1])
            if right:
                recipients.append(min(right)[1])
            recipients = list(dict.fromkeys(recipients))
        if recipients:
            share = float(value) / len(recipients)
            weights[recipients] += share
    return mol, weights


def render_atom_similarity_map(
    mol: Any,
    weights: np.ndarray,
    base: Path,
    scale: float,
    width: int = 520,
    height: int = 360,
) -> list[Path]:
    from rdkit.Chem.Draw import SimilarityMaps, rdMolDraw2D

    base.parent.mkdir(parents=True, exist_ok=True)
    color_map = mpl.colors.LinearSegmentedColormap.from_list(
        "shap_blue_white_red",
        [NEGATIVE_COLOR, "#FFFFFF", POSITIVE_COLOR],
    )
    common_kwargs = {
        "colorMap": color_map,
        "scale": max(float(scale), 1e-8),
        "contourLines": 10,
        "alpha": 0.55,
    }

    svg_path = base.with_suffix(".svg")
    svg_drawer = rdMolDraw2D.MolDraw2DSVG(int(width), int(height))
    SimilarityMaps.GetSimilarityMapFromWeights(mol, weights.tolist(), svg_drawer, **common_kwargs)
    svg_drawer.FinishDrawing()
    svg_path.write_text(svg_drawer.GetDrawingText(), encoding="utf-8")

    png_path = base.with_suffix(".png")
    png_drawer = rdMolDraw2D.MolDraw2DCairo(int(width), int(height))
    SimilarityMaps.GetSimilarityMapFromWeights(mol, weights.tolist(), png_drawer, **common_kwargs)
    png_drawer.FinishDrawing()
    png_path.write_bytes(png_drawer.GetDrawingText())
    return [png_path, svg_path]


def safe_variant_name(name: str) -> str:
    return "_".join(part for part in "".join(char.lower() if char.isalnum() else " " for char in name).split())


def plot_atom_comparison(
    sample_id: int,
    label: int,
    explanations: dict[str, dict[str, Any]],
    output_dir: Path,
) -> tuple[list[Path], list[dict[str, Any]]]:
    atom_data: dict[str, tuple[Any, np.ndarray]] = {}
    for variant, explanation in explanations.items():
        normalized = explanation["normalized_shap_values"]
        atom_data[variant] = token_shap_to_atoms(
            explanation["smiles"],
            explanation["spans"],
            normalized,
        )
    shared_scale = max(
        max(float(np.abs(weights).max()), 1e-8)
        for _, weights in atom_data.values()
    )

    saved: list[Path] = []
    atom_rows: list[dict[str, Any]] = []
    png_paths: list[tuple[str, Path]] = []
    for variant, (mol, weights) in atom_data.items():
        base = output_dir / f"sample_{sample_id:05d}_{safe_variant_name(variant)}_atom_shap"
        variant_paths = render_atom_similarity_map(mol, weights, base, shared_scale)
        saved.extend(variant_paths)
        png_paths.append((variant, variant_paths[0]))
        for atom_index, weight in enumerate(weights):
            atom = mol.GetAtomWithIdx(atom_index)
            atom_rows.append(
                {
                    "variant": variant,
                    "source_index": sample_id,
                    "true_label": label,
                    "atom_index": atom_index,
                    "atom_symbol": atom.GetSymbol(),
                    "normalized_atom_shap": float(weight),
                    "abs_normalized_atom_shap": abs(float(weight)),
                    "smiles": explanations[variant]["smiles"],
                }
            )

    fig, axes = plt.subplots(1, len(png_paths), figsize=(3.7 * len(png_paths), 3.1))
    axes_array = np.atleast_1d(axes)
    for axis, (variant, png_path) in zip(axes_array, png_paths):
        axis.imshow(plt.imread(png_path))
        axis.set_title(variant, fontsize=9, fontweight="semibold")
        axis.axis("off")
    scalar = mpl.cm.ScalarMappable(
        norm=mpl.colors.Normalize(vmin=-shared_scale, vmax=shared_scale),
        cmap=mpl.colors.LinearSegmentedColormap.from_list(
            "shap_blue_white_red_bar",
            [NEGATIVE_COLOR, "#FFFFFF", POSITIVE_COLOR],
        ),
    )
    colorbar = fig.colorbar(scalar, ax=list(axes_array), fraction=0.025, pad=0.02)
    colorbar.set_label("Normalized atom SHAP", fontsize=8)
    colorbar.ax.tick_params(labelsize=7)
    fig.suptitle(f"Sample {sample_id} | true label={label}", fontsize=10, fontweight="semibold")
    fig.subplots_adjust(top=0.84, wspace=0.06)
    saved.extend(save_figure(fig, output_dir / f"sample_{sample_id:05d}_atom_shap_comparison"))
    return saved, atom_rows


def plot_local_comparison(
    sample_id: int,
    label: int,
    explanations: dict[str, dict[str, Any]],
    output_dir: Path,
    top_k: int,
) -> list[Path]:
    variants = list(explanations)
    reference = explanations[variants[0]]
    tokens = reference["tokens"]
    max_abs = np.zeros(len(tokens), dtype=np.float64)
    for explanation in explanations.values():
        if explanation["tokens"] == tokens:
            max_abs = np.maximum(max_abs, np.abs(explanation["normalized_shap_values"]))
    chosen = np.argsort(max_abs)[-min(int(top_k), len(tokens)) :]
    chosen = chosen[np.argsort(max_abs[chosen])]
    labels = [tokens[idx] for idx in chosen]

    all_values = np.concatenate(
        [np.asarray(explanation["normalized_shap_values"])[chosen] for explanation in explanations.values()]
    )
    lower = min(0.0, float(np.min(all_values)))
    upper = max(0.0, float(np.max(all_values)))
    span = max(upper - lower, 1e-3)
    shared_xlim = (lower - 0.05 * span, upper + 0.05 * span)

    # Physical margins are kept separate from the exact 4.24 cm x 5.71 cm
    # central panel so each exported model panel has the same geometry.
    left_margin_in = 0.42
    right_margin_in = 0.05
    bottom_margin_in = 0.28
    title_band_in = 0.38
    top_margin_in = 0.03
    figure_width_in = left_margin_in + TOKEN_PANEL_WIDTH_IN + right_margin_in
    figure_height_in = bottom_margin_in + TOKEN_PANEL_HEIGHT_IN + title_band_in + top_margin_in

    saved: list[Path] = []
    for variant in variants:
        explanation = explanations[variant]
        values = explanation["normalized_shap_values"][chosen]
        colors = [POSITIVE_COLOR if value >= 0 else NEGATIVE_COLOR for value in values]
        fig = plt.figure(figsize=(figure_width_in, figure_height_in), facecolor="white")
        axis = fig.add_axes(
            [
                left_margin_in / figure_width_in,
                bottom_margin_in / figure_height_in,
                TOKEN_PANEL_WIDTH_IN / figure_width_in,
                TOKEN_PANEL_HEIGHT_IN / figure_height_in,
            ]
        )
        axis.barh(np.arange(len(chosen)), values, color=colors, alpha=0.86, height=0.68)
        axis.axvline(0, color="#60666C", linewidth=AXIS_LINEWIDTH, zorder=1)
        axis.set_xlim(shared_xlim)
        axis.set_yticks(np.arange(len(chosen)), labels)
        axis.set_xlabel(
            "Normalized SHAP value",
            fontsize=6,
            fontweight="normal",
            fontfamily=PLOT_FONT_FAMILY,
            labelpad=1.5,
        )
        axis.grid(axis="x", color="#D9DEE3", linewidth=GRID_LINEWIDTH, alpha=0.45)
        axis.set_axisbelow(True)
        axis.spines["top"].set_visible(False)
        axis.spines["right"].set_visible(False)
        axis.spines["left"].set_linewidth(AXIS_LINEWIDTH)
        axis.spines["bottom"].set_linewidth(AXIS_LINEWIDTH)
        axis.tick_params(
            axis="both",
            labelsize=4,
            width=TICK_LINEWIDTH,
            length=1.5,
            pad=1.2,
        )
        for tick_label in [*axis.get_xticklabels(), *axis.get_yticklabels()]:
            tick_label.set_fontfamily(PLOT_FONT_FAMILY)
            tick_label.set_fontweight("normal")

        title_x = (left_margin_in + 0.5 * TOKEN_PANEL_WIDTH_IN) / figure_width_in
        fig.text(
            title_x,
            (figure_height_in - top_margin_in) / figure_height_in,
            variant,
            ha="center",
            va="top",
            fontsize=7,
            fontweight="bold",
            fontfamily=PLOT_FONT_FAMILY,
        )
        fig.text(
            title_x,
            (figure_height_in - 0.17) / figure_height_in,
            f"Sample {sample_id} | true={label}, p(class 1)={explanation['full_probability']:.3f}",
            ha="center",
            va="top",
            fontsize=6,
            fontweight="normal",
            fontfamily=PLOT_FONT_FAMILY,
        )
        variant_slug = safe_variant_name(variant)
        saved.extend(
            save_figure(
                fig,
                output_dir / f"sample_{sample_id:05d}_{variant_slug}_token_shap",
            )
        )
    return saved


def aggregate_global(token_rows: list[dict[str, Any]]) -> list[dict[str, Any]]:
    grouped_raw: dict[tuple[str, str], list[float]] = defaultdict(list)
    grouped_normalized: dict[tuple[str, str], list[float]] = defaultdict(list)
    for row in token_rows:
        key = (str(row["variant"]), str(row["token"]))
        grouped_raw[key].append(abs(float(row["shap_value"])))
        grouped_normalized[key].append(abs(float(row["normalized_shap_value"])))
    output = []
    for (variant, token), values in grouped_raw.items():
        normalized_values = grouped_normalized[(variant, token)]
        output.append(
            {
                "variant": variant,
                "token": token,
                "count": len(values),
                "mean_abs_shap": float(np.mean(values)),
                "std_abs_shap": float(np.std(values)),
                "mean_normalized_abs_shap": float(np.mean(normalized_values)),
                "std_normalized_abs_shap": float(np.std(normalized_values)),
            }
        )
    return output


def plot_global_summary(
    rows: list[dict[str, Any]],
    output_dir: Path,
    top_k: int,
    min_token_count: int,
) -> list[Path]:
    variants = list(dict.fromkeys(str(row["variant"]) for row in rows))
    lookup = {
        (str(row["variant"]), str(row["token"])): float(row["mean_normalized_abs_shap"])
        for row in rows
    }
    token_scores: dict[str, float] = defaultdict(float)
    for row in rows:
        if int(row["count"]) >= int(min_token_count):
            token_scores[str(row["token"])] = max(
                token_scores[str(row["token"])],
                float(row["mean_normalized_abs_shap"]),
            )
    tokens = sorted(token_scores, key=token_scores.get, reverse=True)[: int(top_k)]
    tokens = list(reversed(tokens))
    if not tokens:
        raise RuntimeError(
            f"No token appears at least {min_token_count} times. "
            "Lower --min-token-count or explain more molecules."
        )
    y = np.arange(len(tokens))
    width = 0.72 / max(1, len(variants))
    fig, axis = plt.subplots(figsize=(5.4, max(3.2, 0.25 * len(tokens))))
    for variant_index, variant in enumerate(variants):
        offset = (variant_index - (len(variants) - 1) / 2) * width
        values = [lookup.get((variant, token), 0.0) for token in tokens]
        color = DISTILLED_COLOR if "distill" in variant.lower() else ORIGINAL_COLOR
        axis.barh(y + offset, values, height=width * 0.9, label=variant, color=color)
    axis.set_yticks(y, tokens)
    axis.set_xlabel("Mean normalized |SHAP value|", fontsize=9)
    axis.set_title("Global normalized ChemBLM token importance", fontsize=11, fontweight="semibold")
    axis.legend(frameon=False, fontsize=8)
    axis.grid(axis="x", color="#D9DEE3", linewidth=0.5, alpha=0.55)
    axis.set_axisbelow(True)
    axis.spines["top"].set_visible(False)
    axis.spines["right"].set_visible(False)
    axis.tick_params(labelsize=8)
    return save_figure(fig, output_dir / "global_mean_normalized_abs_shap")


def subset_indices(
    dataset: SmilesTaskDataset,
    config: dict[str, Any],
    subset: str,
    split_method_override: str | None,
    seed: int,
) -> list[int]:
    paths = config.get("paths", {})
    data_cfg = config.get("data", {})
    method = split_method_override or data_cfg.get("split_method", "random")
    train_idx, val_idx, test_idx = split_dataset_indices(
        dataset,
        method=method,
        train_frac=float(data_cfg.get("train_frac", 0.8)),
        val_frac=float(data_cfg.get("val_frac", 0.1)),
        seed=seed,
        task_type="classification",
        split_dir=paths.get("split_dir"),
    )
    if subset == "train":
        return train_idx
    if subset == "val":
        return val_idx
    if subset == "test":
        return test_idx
    return train_idx + val_idx + test_idx


def main() -> None:
    parser = argparse.ArgumentParser(
        description="Token-level permutation SHAP for original and 3D-distilled ChemBLM classifiers."
    )
    parser.add_argument("--config", required=True, help="Primary/distilled finetuning YAML.")
    parser.add_argument("--checkpoint", default=None, help="Primary finetuned best.pt.")
    parser.add_argument("--reference-config", default=None, help="Optional original ChemBLM YAML.")
    parser.add_argument("--reference-checkpoint", default=None, help="Optional original ChemBLM best.pt.")
    parser.add_argument("--task-name", default=None)
    parser.add_argument("--subset", choices=["train", "val", "test", "all"], default="test")
    parser.add_argument("--split-method", choices=["random", "random_scaffold", "scaffold"], default=None)
    parser.add_argument("--num-samples", type=int, default=20)
    parser.add_argument("--permutations", type=int, default=32)
    parser.add_argument("--inference-batch-size", type=int, default=128)
    parser.add_argument("--plot-samples", type=int, default=6)
    parser.add_argument("--top-k-local", type=int, default=20)
    parser.add_argument("--top-k-global", type=int, default=20)
    parser.add_argument(
        "--min-token-count",
        type=int,
        default=5,
        help="Minimum number of token occurrences required before a token is shown globally.",
    )
    parser.add_argument("--seed", type=int, default=None)
    parser.add_argument("--device", default=None)
    parser.add_argument("--output-dir", default="output/figures/shap/chemblm")
    args = parser.parse_args()

    configure_matplotlib()
    config = load_config(args.config)
    paths = config.get("paths", {})
    data_cfg = config.get("data", {})
    train_cfg = config.get("train", {})
    if str(data_cfg.get("task_type", "classification")).lower() != "classification":
        raise ValueError("This SHAP script currently supports classification tasks only.")
    checkpoint_path = args.checkpoint or paths.get("best_checkpoint")
    if not checkpoint_path:
        raise ValueError("Primary finetuned checkpoint is missing.")

    seed = int(args.seed if args.seed is not None else train_cfg.get("seed", 2026))
    primary_split_seed = int(train_cfg.get("seed", 2026))
    set_seed(seed)
    device = resolve_device(args.device or train_cfg.get("device", "auto"))
    label_cols = data_cfg.get("label_cols", data_cfg.get("label_col"))
    dataset = SmilesTaskDataset(
        raw_csv=paths["raw_csv"],
        smiles_col=data_cfg.get("smiles_col", "smiles"),
        label_cols=label_cols,
        task_type="classification",
        max_molecules=data_cfg.get("max_molecules"),
    )
    task_name = args.task_name or dataset.label_cols[0]
    if task_name not in dataset.label_cols:
        raise ValueError(f"Unknown task {task_name!r}; available tasks: {dataset.label_cols}")
    task_index = dataset.label_cols.index(task_name)

    variants: list[tuple[str, dict[str, Any], str | Path]] = []
    reference_config: dict[str, Any] | None = None
    if args.reference_config or args.reference_checkpoint:
        if not args.reference_config:
            raise ValueError("--reference-config is required when --reference-checkpoint is used.")
        reference_config = load_config(args.reference_config)
        reference_checkpoint = args.reference_checkpoint or reference_config.get("paths", {}).get("best_checkpoint")
        if not reference_checkpoint:
            raise ValueError("Reference finetuned checkpoint is missing.")
        variants.append(("Original ChemBLM", reference_config, reference_checkpoint))
    variants.append(("PG-MLD distilled", config, checkpoint_path))

    indices = subset_indices(dataset, config, args.subset, args.split_method, primary_split_seed)
    print(f"Primary {args.subset} split: seed={primary_split_seed}, molecules={len(indices)}")
    if reference_config is not None:
        reference_paths = reference_config.get("paths", {})
        if Path(reference_paths.get("raw_csv", "")) != Path(paths["raw_csv"]):
            raise ValueError("Primary and reference configs must use the same raw_csv for paired SHAP comparison.")
        reference_split_seed = int(reference_config.get("train", {}).get("seed", 2026))
        reference_indices = subset_indices(
            dataset,
            reference_config,
            args.subset,
            args.split_method,
            reference_split_seed,
        )
        common = sorted(set(indices).intersection(reference_indices))
        print(
            f"Reference {args.subset} split: seed={reference_split_seed}, molecules={len(reference_indices)}; "
            f"common molecules={len(common)}"
        )
        if primary_split_seed != reference_split_seed:
            print(
                "Warning: checkpoint split seeds differ. SHAP uses only their common subset; "
                "retrain with identical seeds for the strongest controlled comparison."
            )
        indices = common
        if not indices:
            raise RuntimeError("Primary and reference subsets have no molecules in common.")

    selected = select_samples(dataset, indices, task_index, args.num_samples, seed)
    print(f"Selected {len(selected)} molecules for task={task_name} with sampling seed={seed}")

    models = {
        name: load_best_model(variant_config, checkpoint, len(dataset.label_cols), device)
        for name, variant_config, checkpoint in variants
    }
    output_dir = Path(args.output_dir)
    output_dir.mkdir(parents=True, exist_ok=True)
    token_rows: list[dict[str, Any]] = []
    sample_rows: list[dict[str, Any]] = []
    sample_explanations: dict[int, dict[str, dict[str, Any]]] = defaultdict(dict)

    for variant_name, _, _ in variants:
        model = models[variant_name]
        progress = tqdm(selected, desc=f"Permutation SHAP: {variant_name}")
        for dataset_index in progress:
            smiles = dataset.smiles[dataset_index]
            label = int(round(float(dataset.labels[dataset_index][task_index])))
            explanation = permutation_shap(
                model=model,
                smiles=smiles,
                task_index=task_index,
                permutations=args.permutations,
                inference_batch_size=args.inference_batch_size,
                # A paired comparison must use identical coalition orders for
                # the original and distilled models.
                seed=seed + dataset_index,
                device=device,
            )
            explanation["normalized_shap_values"] = normalize_shap_values(
                explanation["shap_values"]
            )
            sample_explanations[dataset_index][variant_name] = explanation
            sample_rows.append(
                {
                    "variant": variant_name,
                    "dataset_index": dataset_index,
                    "source_index": dataset.source_indices[dataset_index],
                    "task": task_name,
                    "true_label": label,
                    "predicted_probability": explanation["full_probability"],
                    "baseline_probability": explanation["baseline_probability"],
                    "shap_sum": float(np.sum(explanation["shap_values"])),
                    "shap_abs_sum": float(np.abs(explanation["shap_values"]).sum()),
                    "normalized_shap_abs_sum": float(
                        np.abs(explanation["normalized_shap_values"]).sum()
                    ),
                    "reconstruction_error": explanation["reconstruction_error"],
                    "smiles": smiles,
                }
            )
            for token_index, (token, span, value, normalized_value) in enumerate(
                zip(
                    explanation["tokens"],
                    explanation["spans"],
                    explanation["shap_values"],
                    explanation["normalized_shap_values"],
                )
            ):
                token_rows.append(
                    {
                        "variant": variant_name,
                        "dataset_index": dataset_index,
                        "source_index": dataset.source_indices[dataset_index],
                        "task": task_name,
                        "true_label": label,
                        "predicted_probability": explanation["full_probability"],
                        "token_index": token_index,
                        "token": token,
                        "char_start": span[0],
                        "char_end": span[1],
                        "shap_value": float(value),
                        "abs_shap_value": abs(float(value)),
                        "normalized_shap_value": float(normalized_value),
                        "normalized_abs_shap_value": abs(float(normalized_value)),
                        "smiles": smiles,
                    }
                )

    global_rows = aggregate_global(token_rows)
    save_csv(output_dir / "token_shap_values.csv", token_rows)
    save_csv(output_dir / "sample_predictions.csv", sample_rows)
    save_csv(output_dir / "global_token_importance.csv", global_rows)
    global_paths = plot_global_summary(
        global_rows,
        output_dir,
        args.top_k_global,
        args.min_token_count,
    )

    local_paths: list[Path] = []
    atom_rows: list[dict[str, Any]] = []
    for dataset_index in selected[: max(0, int(args.plot_samples))]:
        label = int(round(float(dataset.labels[dataset_index][task_index])))
        local_paths.extend(
            plot_local_comparison(
                sample_id=dataset.source_indices[dataset_index],
                label=label,
                explanations=sample_explanations[dataset_index],
                output_dir=output_dir / "local",
                top_k=args.top_k_local,
            )
        )
        try:
            atom_paths, sample_atom_rows = plot_atom_comparison(
                sample_id=dataset.source_indices[dataset_index],
                label=label,
                explanations=sample_explanations[dataset_index],
                output_dir=output_dir / "atom",
            )
            local_paths.extend(atom_paths)
            atom_rows.extend(sample_atom_rows)
        except Exception as exc:
            print(
                f"Warning: skipped atom-level SHAP heatmap for source_index="
                f"{dataset.source_indices[dataset_index]}: {exc}"
            )
    save_csv(output_dir / "atom_shap_values.csv", atom_rows)

    max_error = max(float(row["reconstruction_error"]) for row in sample_rows)
    print(f"Saved token SHAP CSV: {output_dir / 'token_shap_values.csv'}")
    print(f"Saved sample summary: {output_dir / 'sample_predictions.csv'}")
    print(f"Saved global summary: {output_dir / 'global_token_importance.csv'}")
    if atom_rows:
        print(f"Saved atom-level SHAP CSV: {output_dir / 'atom_shap_values.csv'}")
    for path in [*global_paths, *local_paths]:
        print(f"Saved figure: {path}")
    print(f"Maximum SHAP local-accuracy reconstruction error: {max_error:.8f}")


if __name__ == "__main__":
    main()
```
