## Supplementary material for "PG-MLD: Physics-Guided Molecular Representation Learning via Dynamic 3D Trajectory Distillation": Code and Data: t-sne.py

```
from __future__ import annotations

import argparse
import subprocess
import sys
from pathlib import Path
from typing import Any

PROJECT_ROOT = Path(__file__).resolve().parents[1]
if str(PROJECT_ROOT) not in sys.path:
    sys.path.insert(0, str(PROJECT_ROOT))

import matplotlib as mpl
import matplotlib.pyplot as plt
import numpy as np
import torch
from matplotlib import font_manager
from matplotlib.gridspec import GridSpec
from torch.utils.data import DataLoader, Subset
from tqdm import tqdm

from comld.train.finetune_1d import (
    SmilesTaskDataset,
    collate_smiles,
    load_checkpoint,
    load_config,
    resolve_device,
    set_seed,
    split_dataset_indices,
)
from comld.train.finetune_chemblm_1d import build_model as build_chemblm_model


CLASS_COLORS = {0: "#0A48AB", 1: "#3E7454"}
EDGE_COLORS = {0: "#0A48AB", 1: "#3E7454"}

# Compact SCI figure style. Sizes are in Matplotlib points unless noted.
SCATTER_SIZE = 2.5
SCATTER_EDGE_LINEWIDTH = 0.1
MARGINAL_KDE_LINEWIDTH = 0.35
KDE_CONTOUR_LINEWIDTH = 0.25
AXIS_LINEWIDTH = 0.45
GRID_LINEWIDTH = 0.25
TICK_LINEWIDTH = 0.40

# Exact physical size of the central scatter panel (not the whole figure).
CM_TO_INCH = 1.0 / 2.54
MAIN_PANEL_WIDTH_IN = 3.67 * CM_TO_INCH
MAIN_PANEL_HEIGHT_IN = 2.50 * CM_TO_INCH
PLOT_FONT_FAMILY = "Arial"


def register_arial_font() -> str:
    """Register system Arial files explicitly for isolated Conda environments."""
    try:
        font_manager.findfont(
            font_manager.FontProperties(family=["Arial"]),
            fallback_to_default=False,
        )
        return "Arial"
    except ValueError:
        pass

    candidates: list[Path] = []
    try:
        result = subprocess.run(
            ["fc-match", "-f", "%{file}\n", "Arial"],
            check=False,
            capture_output=True,
            text=True,
            timeout=10,
        )
        matched = Path(result.stdout.strip().splitlines()[0]) if result.stdout.strip() else None
        if matched is not None and matched.is_file():
            candidates.append(matched)
            candidates.extend(
                path
                for path in matched.parent.iterdir()
                if path.is_file()
                and "arial" in path.name.lower()
                and path.suffix.lower() in {".ttf", ".otf", ".ttc"}
            )
    except (OSError, subprocess.SubprocessError):
        pass

    registered_names: set[str] = set()
    for path in dict.fromkeys(candidates):
        try:
            font_manager.fontManager.addfont(str(path))
            registered_names.add(font_manager.FontProperties(fname=str(path)).get_name())
        except (OSError, RuntimeError, ValueError):
            continue

    if any(name.casefold() == "arial" for name in registered_names):
        print(f"Registered Arial directly for Matplotlib: {candidates[0]}")
        return "Arial"

    print("Warning: Arial could not be registered; falling back to DejaVu Sans.")
    return "DejaVu Sans"


def configure_matplotlib() -> None:
    global PLOT_FONT_FAMILY
    PLOT_FONT_FAMILY = register_arial_font()
    mpl.rcParams.update(
        {
            "font.family": "sans-serif",
            "font.sans-serif": [PLOT_FONT_FAMILY, "Helvetica", "DejaVu Sans"],
            "axes.unicode_minus": False,
            "pdf.fonttype": 42,
            "ps.fonttype": 42,
            "svg.fonttype": "none",
            "figure.facecolor": "white",
            "axes.facecolor": "white",
        }
    )


def torch_load(path: str | Path) -> dict[str, Any]:
    path = Path(path)
    if not path.exists():
        raise FileNotFoundError(f"Checkpoint not found: {path}")
    try:
        return torch.load(path, map_location="cpu", weights_only=False)
    except TypeError:
        return torch.load(path, map_location="cpu")


def build_best_chemblm_model(config: dict[str, Any], best_checkpoint: str | Path, output_dim: int) -> torch.nn.Module:
    pretrain_ckpt = load_checkpoint(config.get("paths", {}).get("pretrained_checkpoint"))
    model = build_chemblm_model(config, checkpoint=pretrain_ckpt, output_dim=output_dim)

    best = torch_load(best_checkpoint)
    state = best.get("model_state_dict")
    if state is None:
        raise KeyError(
            f"{best_checkpoint} has no 'model_state_dict'. "
            "Please use a finetuned best.pt, not a pretraining checkpoint."
        )
    missing, unexpected = model.load_state_dict(state, strict=False)
    if missing:
        print(f"Missing keys when loading best checkpoint: {missing}")
    if unexpected:
        print(f"Unexpected keys when loading best checkpoint: {unexpected}")
    print(f"Loaded finetuned best checkpoint: {best_checkpoint}")
    if "epoch" in best:
        print(f"Best checkpoint epoch: {best['epoch']}")
    return model


@torch.no_grad()
def extract_features(
    model: torch.nn.Module,
    loader: DataLoader,
    device: torch.device,
    feature: str,
) -> tuple[np.ndarray, np.ndarray, np.ndarray, list[str]]:
    model.eval()
    features: list[torch.Tensor] = []
    labels: list[torch.Tensor] = []
    masks: list[torch.Tensor] = []
    smiles_all: list[str] = []

    for batch in tqdm(loader, desc=f"Extract {feature} features"):
        smiles = batch["smiles"]
        h = model.encoder(smiles, device=device)

        if feature == "encoder":
            z = h
        elif feature == "penultimate":
            layers = list(model.head.children())
            if len(layers) < 2:
                raise ValueError("Model head is too short; cannot extract penultimate representation.")
            z = h
            for layer in layers[:-1]:
                z = layer(z)
        elif feature == "logits":
            z = model.head(h)
        else:
            raise ValueError(f"Unknown feature={feature!r}. Use encoder, penultimate, or logits.")

        features.append(z.detach().cpu().float())
        labels.append(batch["labels"].detach().cpu().float())
        masks.append(batch["label_mask"].detach().cpu().float())
        smiles_all.extend(smiles)

    return (
        torch.cat(features, dim=0).numpy(),
        torch.cat(labels, dim=0).numpy(),
        torch.cat(masks, dim=0).numpy(),
        smiles_all,
    )


def standardize(x: np.ndarray) -> np.ndarray:
    x = x.astype(np.float32, copy=False)
    return (x - x.mean(axis=0, keepdims=True)) / np.maximum(x.std(axis=0, keepdims=True), 1e-6)


def pca_reduce(x: np.ndarray, n_components: int = 50) -> np.ndarray:
    x_t = torch.as_tensor(standardize(x), dtype=torch.float32)
    x_t = x_t - x_t.mean(dim=0, keepdim=True)
    n_components = min(int(n_components), x_t.shape[0] - 1, x_t.shape[1])
    try:
        _, _, v = torch.pca_lowrank(x_t, q=n_components, center=False)
        z = x_t @ v[:, :n_components]
    except Exception:
        _, _, vh = torch.linalg.svd(x_t, full_matrices=False)
        z = x_t @ vh[:n_components].T
    return z.cpu().numpy()


def torch_tsne(
    x: np.ndarray,
    seed: int,
    perplexity: float = 30.0,
    n_iter: int = 1200,
    learning_rate: float = 200.0,
    pca_dim: int = 50,
) -> np.ndarray:
    """Dependency-light t-SNE implementation for broken scipy/sklearn envs."""

    rng = np.random.default_rng(seed)
    x = pca_reduce(x, n_components=min(pca_dim, x.shape[1]))
    x = standardize(x)
    n = x.shape[0]
    if n <= 2:
        return rng.normal(size=(n, 2)).astype(np.float32)

    perplexity = float(min(perplexity, max(5.0, (n - 1) / 3.0)))
    device = torch.device("cuda" if torch.cuda.is_available() else "cpu")
    x_t = torch.as_tensor(x, dtype=torch.float32, device=device)

    d = torch.cdist(x_t, x_t).pow(2)
    d.fill_diagonal_(float("inf"))

    target_h = np.log(perplexity)
    beta = torch.ones(n, device=device)
    beta_min = torch.full((n,), -float("inf"), device=device)
    beta_max = torch.full((n,), float("inf"), device=device)

    finite_d = torch.where(torch.isfinite(d), d, torch.zeros_like(d))
    for _ in range(50):
        p_cond = torch.exp(-d * beta[:, None])
        p_cond = torch.where(torch.isfinite(d), p_cond, torch.zeros_like(p_cond))
        sum_p = p_cond.sum(dim=1).clamp_min(1e-12)
        h = torch.log(sum_p) + beta * (finite_d * p_cond).sum(dim=1) / sum_p
        high = h > target_h
        beta_min = torch.where(high, beta, beta_min)
        beta_max = torch.where(~high, beta, beta_max)
        beta = torch.where(
            high,
            torch.where(torch.isinf(beta_max), beta * 2.0, (beta + beta_max) / 2.0),
            torch.where(torch.isinf(beta_min), beta / 2.0, (beta + beta_min) / 2.0),
        )

    p_cond = torch.exp(-d * beta[:, None])
    p_cond = torch.where(torch.isfinite(d), p_cond, torch.zeros_like(p_cond))
    p_cond = p_cond / p_cond.sum(dim=1, keepdim=True).clamp_min(1e-12)
    p = ((p_cond + p_cond.T) / (2.0 * n)).clamp_min(1e-12)

    y = torch.as_tensor(rng.normal(0.0, 1e-4, size=(n, 2)), dtype=torch.float32, device=device)
    update = torch.zeros_like(y)
    gains = torch.ones_like(y)
    last_grad = torch.zeros_like(y)

    for step in tqdm(range(int(n_iter)), desc="t-SNE"):
        exaggeration = 12.0 if step < 250 else 1.0
        yy = torch.cdist(y, y).pow(2)
        num = 1.0 / (1.0 + yy)
        num.fill_diagonal_(0.0)
        q = (num / num.sum().clamp_min(1e-12)).clamp_min(1e-12)
        w = (p * exaggeration - q) * num
        grad = 4.0 * (y * w.sum(dim=1, keepdim=True) - w @ y)

        gains = torch.where(torch.sign(grad) != torch.sign(last_grad), gains + 0.2, gains * 0.8).clamp_min(0.01)
        momentum = 0.5 if step < 250 else 0.8
        update = momentum * update - learning_rate * gains * grad
        y = y + update
        y = y - y.mean(dim=0, keepdim=True)
        last_grad = grad

    return y.detach().cpu().numpy()


def make_2d(x: np.ndarray, backend: str, seed: int, perplexity: float, n_iter: int) -> np.ndarray:
    if backend in {"auto", "sklearn"}:
        try:
            from sklearn.manifold import TSNE

            kwargs = dict(
                n_components=2,
                perplexity=perplexity,
                learning_rate="auto",
                init="pca",
                random_state=seed,
            )
            try:
                return TSNE(**kwargs, max_iter=n_iter).fit_transform(standardize(x))
            except TypeError:
                return TSNE(**kwargs, n_iter=n_iter).fit_transform(standardize(x))
        except Exception as exc:
            if backend == "sklearn":
                raise
            print(f"sklearn t-SNE unavailable, fallback to torch t-SNE: {exc}")
    if backend == "pca":
        return pca_reduce(x, n_components=2)
    return torch_tsne(x, seed=seed, perplexity=perplexity, n_iter=n_iter)


def select_binary_task(
    x: np.ndarray,
    labels: np.ndarray,
    masks: np.ndarray,
    smiles: list[str],
    task_names: list[str],
    task_name: str,
    balance: bool,
    seed: int,
) -> tuple[np.ndarray, np.ndarray, list[str]]:
    if task_name not in task_names:
        raise ValueError(f"Task {task_name!r} not found. Available tasks: {task_names}")

    j = task_names.index(task_name)
    y = labels[:, j]
    valid = masks[:, j] > 0
    idx0 = np.flatnonzero(valid & (y < 0.5))
    idx1 = np.flatnonzero(valid & (y > 0.5))
    if len(idx0) == 0 or len(idx1) == 0:
        raise ValueError(f"Task {task_name!r} is not binary after masking: class0={len(idx0)}, class1={len(idx1)}")

    if balance:
        rng = np.random.default_rng(seed)
        n = min(len(idx0), len(idx1))
        keep = np.concatenate(
            [
                rng.choice(idx0, size=n, replace=False),
                rng.choice(idx1, size=n, replace=False),
            ]
        )
        rng.shuffle(keep)
    else:
        keep = np.concatenate([idx0, idx1])

    y_binary = (labels[keep, j] > 0.5).astype(int)
    smiles_keep = [smiles[int(i)] for i in keep]
    return x[keep], y_binary, smiles_keep


def select_regression_task(
    x: np.ndarray,
    labels: np.ndarray,
    masks: np.ndarray,
    smiles: list[str],
    task_names: list[str],
    task_name: str,
) -> tuple[np.ndarray, np.ndarray, list[str]]:
    if task_name not in task_names:
        raise ValueError(f"Task {task_name!r} not found. Available tasks: {task_names}")
    j = task_names.index(task_name)
    values = labels[:, j].astype(np.float64, copy=False)
    keep = np.flatnonzero((masks[:, j] > 0) & np.isfinite(values))
    if len(keep) < 3:
        raise ValueError(f"Task {task_name!r} has only {len(keep)} valid regression targets.")
    return x[keep], values[keep], [smiles[int(i)] for i in keep]


def davies_bouldin_index(x: np.ndarray, y: np.ndarray) -> float:
    classes = np.unique(y)
    if len(classes) < 2:
        return float("nan")
    x = x.astype(np.float64, copy=False)
    centroids = []
    scatters = []
    for cls in classes:
        pts = x[y == cls]
        center = pts.mean(axis=0)
        centroids.append(center)
        scatters.append(np.linalg.norm(pts - center, axis=1).mean())
    centroids = np.stack(centroids, axis=0)
    scatters = np.asarray(scatters)

    scores = []
    for i in range(len(classes)):
        ratios = []
        for j in range(len(classes)):
            if i == j:
                continue
            dist = np.linalg.norm(centroids[i] - centroids[j])
            ratios.append((scatters[i] + scatters[j]) / max(dist, 1e-12))
        scores.append(max(ratios))
    return float(np.mean(scores))


def data_limits(z: np.ndarray, margin_ratio: float = 0.05) -> tuple[tuple[float, float], tuple[float, float]]:
    xmin, xmax = float(z[:, 0].min()), float(z[:, 0].max())
    ymin, ymax = float(z[:, 1].min()), float(z[:, 1].max())
    xpad = (xmax - xmin) * margin_ratio if xmax > xmin else 1.0
    ypad = (ymax - ymin) * margin_ratio if ymax > ymin else 1.0
    return (xmin - xpad, xmax + xpad), (ymin - ypad, ymax + ypad)


def _numpy_gaussian_kde(samples: np.ndarray, query: np.ndarray) -> np.ndarray:
    """Small SciPy-compatible fallback used only when scipy.stats is unavailable."""

    samples = np.asarray(samples, dtype=np.float64)
    query = np.asarray(query, dtype=np.float64)
    if samples.ndim == 1:
        samples = samples[None, :]
    if query.ndim == 1:
        query = query[None, :]
    dim, count = samples.shape
    if count < max(3, dim + 1):
        raise ValueError(f"Too few samples for {dim}D KDE: {count}")

    covariance = np.atleast_2d(np.cov(samples, bias=False))
    factor = count ** (-1.0 / (dim + 4.0))
    covariance = covariance * factor**2
    scale = max(float(np.trace(covariance)) / max(dim, 1), 1.0)
    covariance = covariance + np.eye(dim) * scale * 1e-8
    inverse = np.linalg.inv(covariance)
    determinant = max(float(np.linalg.det(covariance)), 1e-24)
    normalizer = np.sqrt((2.0 * np.pi) ** dim * determinant)

    result = np.empty(query.shape[1], dtype=np.float64)
    sample_rows = samples.T
    for start in range(0, query.shape[1], 4096):
        stop = min(start + 4096, query.shape[1])
        differences = query[:, start:stop].T[:, None, :] - sample_rows[None, :, :]
        exponent = np.einsum("mni,ij,mnj->mn", differences, inverse, differences, optimize=True)
        result[start:stop] = np.exp(-0.5 * exponent).mean(axis=1) / normalizer
    return result


def gaussian_kde_density(samples: np.ndarray, query: np.ndarray) -> np.ndarray:
    samples = np.asarray(samples, dtype=np.float64)
    query = np.asarray(query, dtype=np.float64)
    try:
        from scipy.stats import gaussian_kde

        return np.asarray(gaussian_kde(samples)(query), dtype=np.float64)
    except Exception as exc:
        if not getattr(gaussian_kde_density, "_fallback_reported", False):
            print(f"SciPy gaussian_kde unavailable; using NumPy KDE fallback: {exc}")
            gaussian_kde_density._fallback_reported = True
        return _numpy_gaussian_kde(samples, query)


def kde_levels(density: np.ndarray) -> np.ndarray:
    positive = density[np.isfinite(density) & (density > 0)]
    if positive.size == 0:
        return np.array([])
    return np.unique(np.quantile(positive, (0.55, 0.70, 0.82, 0.90, 0.96)))


def draw_2d_kde(
    ax: plt.Axes,
    z: np.ndarray,
    y: np.ndarray,
    xlim: tuple[float, float],
    ylim: tuple[float, float],
) -> None:
    xs = np.linspace(*xlim, 190)
    ys = np.linspace(*ylim, 190)
    xx, yy = np.meshgrid(xs, ys)
    grid = np.vstack([xx.ravel(), yy.ravel()])

    for cls in (0, 1):
        points = z[y == cls]
        if len(points) < 5:
            print(f"Skipping class {cls} 2D KDE: only {len(points)} samples.")
            continue
        try:
            density = gaussian_kde_density(points.T, grid).reshape(xx.shape)
            levels = kde_levels(density)
            if len(levels) < 2:
                continue
            ax.contourf(
                xx,
                yy,
                density,
                levels=levels,
                colors=[CLASS_COLORS[cls]],
                alpha=0.075,
                antialiased=True,
                zorder=1,
            )
            ax.contour(
                xx,
                yy,
                density,
                levels=levels,
                colors=[CLASS_COLORS[cls]],
                linewidths=KDE_CONTOUR_LINEWIDTH,
                alpha=0.18,
                zorder=2,
            )
        except Exception as exc:
            print(f"Skipping class {cls} 2D KDE: {exc}")


def draw_marginal_kde(
    ax_top: plt.Axes,
    ax_right: plt.Axes,
    z: np.ndarray,
    y: np.ndarray,
    xlim: tuple[float, float],
    ylim: tuple[float, float],
) -> None:
    x_grid = np.linspace(*xlim, 500)
    y_grid = np.linspace(*ylim, 500)
    top_max = 0.0
    right_max = 0.0

    for cls in (0, 1):
        try:
            x_density = gaussian_kde_density(z[y == cls, 0], x_grid)
            top_max = max(top_max, float(x_density.max()))
            ax_top.plot(x_grid, x_density, color=CLASS_COLORS[cls], linewidth=MARGINAL_KDE_LINEWIDTH)
            ax_top.fill_between(x_grid, 0, x_density, color=CLASS_COLORS[cls], alpha=0.12)
        except Exception as exc:
            print(f"Skipping class {cls} top KDE: {exc}")
        try:
            y_density = gaussian_kde_density(z[y == cls, 1], y_grid)
            right_max = max(right_max, float(y_density.max()))
            ax_right.plot(y_density, y_grid, color=CLASS_COLORS[cls], linewidth=MARGINAL_KDE_LINEWIDTH)
            ax_right.fill_betweenx(y_grid, 0, y_density, color=CLASS_COLORS[cls], alpha=0.12)
        except Exception as exc:
            print(f"Skipping class {cls} right KDE: {exc}")

    ax_top.set_xlim(xlim)
    ax_top.set_ylim(0, top_max * 1.08 if top_max > 0 else 1.0)
    ax_right.set_ylim(ylim)
    ax_right.set_xlim(0, right_max * 1.08 if right_max > 0 else 1.0)
    for marginal in (ax_top, ax_right):
        marginal.set_xticks([])
        marginal.set_yticks([])
        for spine in marginal.spines.values():
            spine.set_visible(False)


def plot_tsne(
    z: np.ndarray,
    y: np.ndarray,
    output_path: str | Path,
    title: str,
    dbi_original: float,
    dbi_tsne: float,
    class0_name: str = "Class 0",
    class1_name: str = "Class 1",
    show_legend: bool = False,
    method_name: str = "t-SNE",
) -> list[Path]:
    configure_matplotlib()
    z = np.asarray(z, dtype=np.float64)
    y = np.asarray(y, dtype=int)
    valid = np.isfinite(z).all(axis=1) & np.isin(y, [0, 1])
    removed = int((~valid).sum())
    if removed:
        print(f"Removed invalid t-SNE samples before plotting: {removed}")
    z, y = z[valid], y[valid]
    if len(z) == 0:
        raise RuntimeError("No valid t-SNE coordinates remain for plotting.")

    xlim, ylim = data_limits(z)

    # The outer canvas is only a carrier for titles and marginal KDEs. The
    # central axes are positioned in physical units so their exported size is
    # exactly 3.67 cm x 2.50 cm.
    left_margin_in = 0.34
    bottom_margin_in = 0.28
    marginal_gap_in = 0.035
    top_kde_height_in = 0.24
    right_kde_width_in = 0.24
    title_band_in = 0.30
    right_margin_in = 0.05
    top_margin_in = 0.03
    figure_width_in = (
        left_margin_in
        + MAIN_PANEL_WIDTH_IN
        + marginal_gap_in
        + right_kde_width_in
        + right_margin_in
    )
    figure_height_in = (
        bottom_margin_in
        + MAIN_PANEL_HEIGHT_IN
        + marginal_gap_in
        + top_kde_height_in
        + title_band_in
        + top_margin_in
    )
    fig = plt.figure(figsize=(figure_width_in, figure_height_in), facecolor="white")

    def axes_rect(left: float, bottom: float, width: float, height: float) -> list[float]:
        return [
            left / figure_width_in,
            bottom / figure_height_in,
            width / figure_width_in,
            height / figure_height_in,
        ]

    ax = fig.add_axes(
        axes_rect(left_margin_in, bottom_margin_in, MAIN_PANEL_WIDTH_IN, MAIN_PANEL_HEIGHT_IN)
    )
    ax_top = fig.add_axes(
        axes_rect(
            left_margin_in,
            bottom_margin_in + MAIN_PANEL_HEIGHT_IN + marginal_gap_in,
            MAIN_PANEL_WIDTH_IN,
            top_kde_height_in,
        )
    )
    ax_right = fig.add_axes(
        axes_rect(
            left_margin_in + MAIN_PANEL_WIDTH_IN + marginal_gap_in,
            bottom_margin_in,
            right_kde_width_in,
            MAIN_PANEL_HEIGHT_IN,
        )
    )

    draw_marginal_kde(ax_top, ax_right, z, y, xlim, ylim)
    draw_2d_kde(ax, z, y, xlim, ylim)

    names = {0: class0_name, 1: class1_name}
    for cls in (0, 1):
        mask = y == cls
        ax.scatter(
            z[mask, 0],
            z[mask, 1],
            s=SCATTER_SIZE,
            c=CLASS_COLORS[cls],
            alpha=0.72,
            linewidth=SCATTER_EDGE_LINEWIDTH,
            edgecolors=EDGE_COLORS[cls],
            rasterized=True,
            zorder=4,
            label=names[cls],
        )

    if show_legend:
        legend = ax.legend(
            loc="upper right",
            frameon=True,
            framealpha=0.92,
            facecolor="white",
            edgecolor="#D3D7DB",
            fontsize=5,
        )
        legend.get_frame().set_linewidth(0.4)

    ax.set_xlim(xlim)
    ax.set_ylim(ylim)
    ax.set_xlabel(
        f"{method_name} 1",
        fontsize=6,
        fontweight="normal",
        fontfamily=PLOT_FONT_FAMILY,
        labelpad=1.5,
    )
    ax.set_ylabel(
        f"{method_name} 2",
        fontsize=6,
        fontweight="normal",
        fontfamily=PLOT_FONT_FAMILY,
        labelpad=1.5,
    )
    ax.tick_params(
        axis="both",
        labelsize=4,
        width=TICK_LINEWIDTH,
        length=1.5,
        pad=1.2,
    )
    for tick_label in [*ax.get_xticklabels(), *ax.get_yticklabels()]:
        tick_label.set_fontfamily(PLOT_FONT_FAMILY)
        tick_label.set_fontweight("normal")
    ax.spines["top"].set_visible(False)
    ax.spines["right"].set_visible(False)
    ax.spines["left"].set_linewidth(AXIS_LINEWIDTH)
    ax.spines["bottom"].set_linewidth(AXIS_LINEWIDTH)
    ax.grid(True, color="#D9DEE3", linewidth=GRID_LINEWIDTH, alpha=0.45)
    ax.set_axisbelow(True)

    title_x = (left_margin_in + 0.5 * MAIN_PANEL_WIDTH_IN) / figure_width_in
    fig.text(
        title_x,
        (figure_height_in - top_margin_in) / figure_height_in,
        title,
        ha="center",
        va="top",
        fontsize=7,
        fontweight="bold",
        fontfamily=PLOT_FONT_FAMILY,
    )
    fig.text(
        title_x,
        (figure_height_in - 0.17) / figure_height_in,
        f"DBI (original): {dbi_original:.2f}, {method_name}: {dbi_tsne:.2f}",
        ha="center",
        va="top",
        fontsize=6,
        fontweight="normal",
        fontfamily=PLOT_FONT_FAMILY,
    )

    base = Path(output_path).with_suffix("")
    base.parent.mkdir(parents=True, exist_ok=True)
    saved = [base.with_suffix(ext) for ext in (".png", ".pdf", ".svg")]
    fig.savefig(saved[0], dpi=600, bbox_inches="tight", pad_inches=0.04, facecolor="white")
    fig.savefig(saved[1], bbox_inches="tight", pad_inches=0.04, facecolor="white")
    fig.savefig(saved[2], bbox_inches="tight", pad_inches=0.04, facecolor="white")
    plt.close(fig)
    return saved


def pearson_correlation(a: np.ndarray, b: np.ndarray) -> float:
    a = np.asarray(a, dtype=np.float64)
    b = np.asarray(b, dtype=np.float64)
    if len(a) < 2 or np.std(a) < 1e-12 or np.std(b) < 1e-12:
        return float("nan")
    return float(np.corrcoef(a, b)[0, 1])


def draw_global_kde(
    ax: plt.Axes,
    ax_top: plt.Axes,
    ax_right: plt.Axes,
    z: np.ndarray,
    xlim: tuple[float, float],
    ylim: tuple[float, float],
) -> None:
    color = CLASS_COLORS[0]
    xs = np.linspace(*xlim, 190)
    ys = np.linspace(*ylim, 190)
    xx, yy = np.meshgrid(xs, ys)
    try:
        density = gaussian_kde_density(z.T, np.vstack([xx.ravel(), yy.ravel()])).reshape(xx.shape)
        levels = kde_levels(density)
        if len(levels) >= 2:
            ax.contourf(xx, yy, density, levels=levels, colors=[color], alpha=0.06, zorder=1)
            ax.contour(xx, yy, density, levels=levels, colors=[color], linewidths=0.7, alpha=0.17, zorder=2)
    except Exception as exc:
        print(f"Skipping global 2D KDE: {exc}")

    x_grid = np.linspace(*xlim, 500)
    y_grid = np.linspace(*ylim, 500)
    try:
        dx = gaussian_kde_density(z[:, 0], x_grid)
        ax_top.plot(x_grid, dx, color=color, linewidth=MARGINAL_KDE_LINEWIDTH)
        ax_top.fill_between(x_grid, 0, dx, color=color, alpha=0.12)
        ax_top.set_ylim(0, float(dx.max()) * 1.08)
    except Exception as exc:
        print(f"Skipping global top KDE: {exc}")
        ax_top.set_ylim(0, 1)
    try:
        dy = gaussian_kde_density(z[:, 1], y_grid)
        ax_right.plot(dy, y_grid, color=color, linewidth=MARGINAL_KDE_LINEWIDTH)
        ax_right.fill_betweenx(y_grid, 0, dy, color=color, alpha=0.12)
        ax_right.set_xlim(0, float(dy.max()) * 1.08)
    except Exception as exc:
        print(f"Skipping global right KDE: {exc}")
        ax_right.set_xlim(0, 1)

    ax_top.set_xlim(xlim)
    ax_right.set_ylim(ylim)
    for marginal in (ax_top, ax_right):
        marginal.set_xticks([])
        marginal.set_yticks([])
        for spine in marginal.spines.values():
            spine.set_visible(False)


def plot_regression_embedding(
    z: np.ndarray,
    values: np.ndarray,
    output_path: str | Path,
    title: str,
    target_name: str,
    method_name: str,
) -> tuple[list[Path], float, float]:
    configure_matplotlib()
    z = np.asarray(z, dtype=np.float64)
    values = np.asarray(values, dtype=np.float64)
    valid = np.isfinite(z).all(axis=1) & np.isfinite(values)
    removed = int((~valid).sum())
    if removed:
        print(f"Removed invalid regression samples before plotting: {removed}")
    z, values = z[valid], values[valid]
    if len(z) == 0:
        raise RuntimeError("No valid regression samples remain for plotting.")

    xlim, ylim = data_limits(z)
    fig = plt.figure(figsize=(7.2, 7.2), facecolor="white")
    grid = GridSpec(
        2,
        2,
        figure=fig,
        width_ratios=[0.85, 0.15],
        height_ratios=[0.15, 0.85],
        wspace=0.03,
        hspace=0.03,
        left=0.10,
        right=0.90,
        bottom=0.08,
        top=0.88,
    )
    ax_top = fig.add_subplot(grid[0, 0])
    ax = fig.add_subplot(grid[1, 0])
    ax_right = fig.add_subplot(grid[1, 1])
    draw_global_kde(ax, ax_top, ax_right, z, xlim, ylim)

    points = ax.scatter(
        z[:, 0],
        z[:, 1],
        c=values,
        cmap="viridis",
        s=SCATTER_SIZE,
        alpha=0.76,
        linewidth=0.25,
        edgecolors="white",
        rasterized=True,
        zorder=4,
    )
    ax.set_xlim(xlim)
    ax.set_ylim(ylim)
    ax.set_box_aspect(1)
    ax.set_xlabel(f"{method_name} 1", fontsize=11)
    ax.set_ylabel(f"{method_name} 2", fontsize=11)
    ax.tick_params(axis="both", labelsize=9.5, width=0.8, length=3.5)
    ax.spines["top"].set_visible(False)
    ax.spines["right"].set_visible(False)
    ax.spines["left"].set_linewidth(0.8)
    ax.spines["bottom"].set_linewidth(0.8)
    ax.grid(True, color="#D9DEE3", linewidth=0.55, alpha=0.45)
    ax.set_axisbelow(True)

    corr1 = pearson_correlation(z[:, 0], values)
    corr2 = pearson_correlation(z[:, 1], values)
    fig.text(0.5, 0.982, title, ha="center", va="top", fontsize=15, fontweight="semibold")
    fig.text(
        0.5,
        0.936,
        f"{target_name}: r({method_name} 1)={corr1:.2f}, r({method_name} 2)={corr2:.2f}",
        ha="center",
        va="top",
        fontsize=11.5,
    )
    cbar_axis = fig.add_axes([0.945, 0.17, 0.014, 0.62])
    colorbar = fig.colorbar(points, cax=cbar_axis)
    colorbar.set_label(target_name, fontsize=9.5)
    colorbar.ax.tick_params(labelsize=8.5, width=0.6, length=2.5)
    colorbar.outline.set_linewidth(0.6)

    base = Path(output_path).with_suffix("")
    base.parent.mkdir(parents=True, exist_ok=True)
    saved = [base.with_suffix(ext) for ext in (".png", ".pdf", ".svg")]
    fig.savefig(saved[0], dpi=600, facecolor="white")
    fig.savefig(saved[1], facecolor="white")
    fig.savefig(saved[2], facecolor="white")
    plt.close(fig)
    return saved, corr1, corr2


def main() -> None:
    parser = argparse.ArgumentParser(description="t-SNE visualization from a finetuned ChemBLM best.pt.")
    parser.add_argument("--config", required=True)
    parser.add_argument("--checkpoint", default=None, help="Finetuned best.pt; defaults to paths.best_checkpoint in YAML.")
    parser.add_argument("--output-dir", default="output/figures/tsne")
    parser.add_argument("--dataset-name", default="BBBP")
    parser.add_argument("--task-name", default=None)
    parser.add_argument("--subset", default="test", choices=["train", "val", "test", "all"])
    parser.add_argument("--split-method", default=None, choices=["random", "random_scaffold", "scaffold"])
    parser.add_argument("--feature", default="penultimate", choices=["encoder", "penultimate", "logits"])
    parser.add_argument("--balance", action="store_true", help="Balance class 0/1 before t-SNE.")
    parser.add_argument("--max-points", type=int, default=0, help="0 means no random subsampling after class filtering.")
    parser.add_argument("--batch-size", type=int, default=64)
    parser.add_argument("--backend", default="auto", choices=["auto", "sklearn", "torch", "pca"])
    parser.add_argument("--perplexity", type=float, default=30.0)
    parser.add_argument("--n-iter", type=int, default=1200)
    parser.add_argument("--seed", type=int, default=None)
    parser.add_argument("--device", default=None)
    parser.add_argument("--class0-name", default="Class 0")
    parser.add_argument("--class1-name", default="Class 1")
    parser.add_argument("--show-legend", action="store_true", help="Show the Class 0/Class 1 legend (hidden by default).")
    args = parser.parse_args()

    config = load_config(args.config)
    paths = config.get("paths", {})
    data_cfg = config.get("data", {})
    train_cfg = config.get("train", {})
    checkpoint_path = args.checkpoint or paths.get("best_checkpoint")
    if not checkpoint_path:
        raise ValueError("No finetuned checkpoint supplied and paths.best_checkpoint is empty in the config.")

    seed = int(args.seed if args.seed is not None else train_cfg.get("seed", 2026))
    set_seed(seed)
    device = resolve_device(args.device or train_cfg.get("device", "auto"))

    label_cols = data_cfg.get("label_cols", data_cfg.get("label_col"))
    dataset = SmilesTaskDataset(
        raw_csv=paths["raw_csv"],
        smiles_col=data_cfg.get("smiles_col", "smiles"),
        label_cols=label_cols,
        task_type=data_cfg.get("task_type", "classification"),
        max_molecules=data_cfg.get("max_molecules"),
    )
    task_name = args.task_name or dataset.label_cols[0]

    split_method = args.split_method or data_cfg.get("split_method", "random")
    train_idx, val_idx, test_idx = split_dataset_indices(
        dataset=dataset,
        method=split_method,
        train_frac=float(data_cfg.get("train_frac", 0.8)),
        val_frac=float(data_cfg.get("val_frac", 0.1)),
        seed=seed,
        task_type=data_cfg.get("task_type", "classification"),
        split_dir=paths.get("split_dir"),
    )
    if args.subset == "train":
        indices = train_idx
    elif args.subset == "val":
        indices = val_idx
    elif args.subset == "test":
        indices = test_idx
    else:
        indices = train_idx + val_idx + test_idx

    loader = DataLoader(
        Subset(dataset, indices),
        batch_size=int(args.batch_size),
        shuffle=False,
        num_workers=0,
        collate_fn=collate_smiles,
    )

    model = build_best_chemblm_model(config, checkpoint_path, output_dim=len(dataset.label_cols)).to(device)
    x, labels, masks, smiles = extract_features(model, loader, device, feature=args.feature)
    task_type = str(data_cfg.get("task_type", "classification")).lower()
    if task_type == "regression":
        if args.balance:
            print("Ignoring --balance for regression task.")
        x, y, smiles = select_regression_task(
            x=x,
            labels=labels,
            masks=masks,
            smiles=smiles,
            task_names=dataset.label_cols,
            task_name=task_name,
        )
    else:
        x, y, smiles = select_binary_task(
            x=x,
            labels=labels,
            masks=masks,
            smiles=smiles,
            task_names=dataset.label_cols,
            task_name=task_name,
            balance=bool(args.balance),
            seed=seed,
        )

    if args.max_points and len(y) > int(args.max_points):
        rng = np.random.default_rng(seed)
        keep = rng.choice(np.arange(len(y)), size=int(args.max_points), replace=False)
        x = x[keep]
        y = y[keep]
        smiles = [smiles[int(i)] for i in keep]

    z = make_2d(x, backend=args.backend, seed=seed, perplexity=args.perplexity, n_iter=args.n_iter)
    method_name = "PCA" if args.backend == "pca" else "t-SNE"

    output_dir = Path(args.output_dir)
    output_dir.mkdir(parents=True, exist_ok=True)
    stem = f"{args.dataset_name.lower()}_chemblm_{args.subset}_{args.feature}"
    if args.balance:
        stem += "_balanced"

    if task_type == "regression":
        dbi_original = float("nan")
        dbi_tsne = float("nan")
        saved_figures, corr1, corr2 = plot_regression_embedding(
            z=z,
            values=y,
            output_path=output_dir / f"{stem}.png",
            title=f"{args.dataset_name} ChemBLM",
            target_name=task_name,
            method_name=method_name,
        )
    else:
        dbi_original = davies_bouldin_index(standardize(x), y)
        dbi_tsne = davies_bouldin_index(z, y)
        corr1 = float("nan")
        corr2 = float("nan")
        saved_figures = plot_tsne(
            z=z,
            y=y,
            output_path=output_dir / f"{stem}.png",
            title=f"{args.dataset_name} ChemBLM",
            dbi_original=dbi_original,
            dbi_tsne=dbi_tsne,
            class0_name=args.class0_name,
            class1_name=args.class1_name,
            show_legend=bool(args.show_legend),
            method_name=method_name,
        )
    np.savez_compressed(
        output_dir / f"{stem}.npz",
        tsne=z,
        features=x,
        labels=y,
        smiles=np.array(smiles),
        task_name=np.array(task_name),
        dbi_original=np.array(dbi_original),
        dbi_tsne=np.array(dbi_tsne),
        correlation_axis1=np.array(corr1),
        correlation_axis2=np.array(corr2),
        task_type=np.array(task_type),
        backend=np.array(args.backend),
        seed=np.array(seed),
        perplexity=np.array(args.perplexity),
        n_iter=np.array(args.n_iter),
    )

    for saved_path in saved_figures:
        print(f"Saved: {saved_path}")
    if task_type == "regression":
        print(f"Samples: total={len(y)} target_min={float(np.min(y)):.4f} target_max={float(np.max(y)):.4f}")
    else:
        print(f"Samples: total={len(y)} class0={(y == 0).sum()} class1={(y == 1).sum()}")
    print(f"Feature: {args.feature}")
    print(f"Reduction: {method_name} (backend={args.backend})")
    if task_type == "regression":
        print(f"Correlation: {method_name}1={corr1:.4f}, {method_name}2={corr2:.4f}")
    else:
        print(f"DBI original={dbi_original:.4f}, DBI {method_name}={dbi_tsne:.4f}")


if __name__ == "__main__":
    main()
```
